# Ticks convert pathogenic *Coxiella* into endosymbionts

**DOI:** 10.1101/2020.12.29.424491

**Authors:** Amanda E. Brenner, Sebastián Muñoz-Leal, Madhur Sachan, Marcelo B. Labruna, Rahul Raghavan

**Affiliations:** Department of Biology and Center for Life in Extreme Environments, Portland State University, Portland, Oregon, 97201, USA; Departamento de Patología y Medicina Preventiva, Facultad de Ciencias Veterinarias, Universidad de Concepción, Chillán, Ñuble, Chile; Departamento de Medicina Veterinária Preventiva e Saúde Animal, Faculdade de Medicina Veterinária e Zootecnia, Universidade de São Paulo, São Paulo, Brazil

## Abstract

Both symbiotic and pathogenic bacteria in the family Coxiellaceae cause morbidity and mortality in humans and animals. For instance, *Coxiella*-like endosymbionts (CLEs) improve the reproductive success of ticks — a major disease vector, while *Coxiella burnetii* is the etiological agent of human Q fever and uncharacterized coxiellae cause infections in both animals and humans. To better understand the evolution of pathogenesis and symbiosis in this group of intracellular bacteria, we sequenced the genome of a CLE present in the soft tick *Ornithodoros amblus* (CLEOA) and compared it to the genomes of other bacteria in the order Legionellales. Our analyses confirmed that CLEOA is more closely related to *C. burnetii*, the human pathogen, than to CLEs in hard ticks, and showed that most clades of CLEs contain both endosymbionts and pathogens, indicating that several CLE lineages have evolved independently from pathogenic *Coxiella*. We also determined that the last common ancestor of CLEOA and *C. burnetii* was equipped to infect macrophages, and that even though HGT contributed significantly to the evolution of *C. burnetii*, most acquisition events occurred primarily in ancestors predating the CLEOA-*C. burnetii* divergence. These discoveries clarify the evolution of *C. burnetii*, which previously was assumed to have emerged when an avirulent tick endosymbiont recently gained virulence factors via HGT. Finally, we identified several metabolic pathways, including heme biosynthesis, that are likely critical to the intracellular growth of the human pathogen but not the tick symbiont, and show that the use of heme analogs is a promising approach to controlling *C. burnetii* infections.

## INTRODUCTION

A bacterium’s genome size and gene content signal both the degree of its dependence on the host and the length of the bacterium-host relationship. For example, a bacterium that has established a long-term, obligate symbiosis would have a tiny genome filled with protein-coding genes (Wernegreen et al. 2000). Conversely, the genome of a bacterium that is in early stages of symbiosis is usually large and contains numerous pseudogenized genes, which, as the relationship progresses, would eventually be lost, resulting in a tiny genome (Moran 2002; McCutcheon and Moran 2011). The genomes of *Coxiella*-like endosymbionts (CLEs) found in ticks (Acari: Ixodida) fall into both categories: Some ticks, e.g., *Amblyomma americanum* and *A. sculptum*, contain small-genomed CLEs (~0.6 Mbp) that have few pseudogenes, indicating that they represent an ancient lineage of tick endosymbionts (Smith et al. 2015). In contrast, CLEs in *Rhipicephalus turanicus* (CRt), and *R. sanguineus* have large genomes (~1.7 Mbp) filled with pseudogenes, denoting that the bacteria are in early stages of symbioses (Gottlieb et al. 2015; Tsementzi et al. 2018). While most ticks contain CLEs, a few have *Francisella*-like endosymbionts (FLEs) instead (Gerhart et al. 2016, 2018; Duron et al. 2017). All FLEs studied to date have large genomes (~1.5 Mbp) with hundreds of pseudogenes, including inactivated virulence genes, indicating that FLEs evolved recently from pathogenic ancestors (Gerhart et al. 2016, 2018; Duron et al. 2018b). Irrespective of their age, CLEs and FLEs improve the reproductive fitness of their hosts by likely providing metabolites missing in vertebrate blood, ticks’ sole nutritional source (Gottlieb et al. 2015; Smith et al. 2015; Gerhart et al. 2016, 2018; Duron et al. 2017, 2018b; Tsementzi et al. 2018).

*C. burnetii*, the causative agent of human Q fever, has also been detected in ticks; in fact, the intracellular pathogen was first isolated from hard ticks *Dermacentor andersoni* and *Haemaphysalis humerosa* (Cox 1938; Smith and Derrick 1940). In addition, transstadial transmission and fecal excretion of *C. burnetii* occur in laboratory-raised ticks (Eldin et al. 2017; Körner et al. 2020). However, it is not clear whether ticks play any meaningful role in the natural spread of *C. burnetii* (Duron et al. 2015b); instead, Q fever generally occurs following inhalation of *C. burnetii*-contaminated aerosols originating from infected farm animals (Maurin and Raoult 1999; Eldin et al. 2017). Within the human lungs, *C. burnetii* infects alveolar macrophages and generates a large replicative vacuole, termed the *Coxiella*-containing vacuole (CCV), by subverting host responses through a Dot/Icm Type IVB secretion system (T4BSS). This secretion system is essential to the pathogenicity of both *C. burnetii* and *Legionella pneumophila*, the two established pathogens in the order Legionellales (Segal et al. 1999; Chen et al. 2010; Beare et al. 2011; Newton et al. 2014; Burstein et al. 2016). Genes for T4BSS, which evolved from conjugation machinery, have spread across the bacterial kingdom via horizontal gene transfer (HGT), a process through which organisms gain foreign genes, allowing them to quickly adapt to a new environment (Ochman et al. 2000; Lerat et al. 2005).

The closest relatives of *C. burnetii* are CLEs present in ticks (Almeida et al. 2012; Duron et al. 2015a; Smith et al. 2015), leading to the notion that the human pathogen emerged when an avirulent tick endosymbiont gained pathogenicity genes, probably via HGT (Duron et al. 2015a; Gerhart et al. 2016). Contrary to this hypothesis, by sequencing the genome of a CLE in *Ornithodoros amblus* (henceforth referred to as CLEOA), we show that a common virulent ancestor gave rise to both *C. burnetii* and CLEOA. The potentially-pathogenic ancestor contained genes for most virulence factors, including T4BSS, indicating that the erstwhile bacterium was likely capable of infecting mammalian macrophages. In CLEOA, homologs of most virulence-associated genes have been rendered non-functional, but its genome is enriched for B vitamin and cofactor biosynthesis pathways, suggesting that a virulent bacterium has morphed into a nutrient-provisioning tick endosymbiont. Similar to this pathogen-to-endosymbiont transformation, we found that several other tick endosymbionts likely evolved from pathogenic ancestors, indicating that the blood-sucking arthropod is proficient at “domesticating” virulent bacteria into endosymbionts. Finally, by inhibiting *C. burnetii* growth using a synthetic analog of heme, a metabolite produced by *C. burnetii* but not by CLEOA, we demonstrate how knowledge gained through comparative genomics could be applied to developing novel strategies to control Q fever, which is difficult to treat with currently available antibiotics.

## RESULTS

### CLEOA arose from a pathogenic ancestor

Phylogenetic trees based mainly on 16S rDNA have previously indicated that the closest relatives of *C. burnetii* are CLEs present in *Orniothodoros* and *Argas* soft ticks (family Argasidae) (Almeida et al. 2012; Duron et al. 2015a; Smith et al. 2015); however, all CLE genomes available to date are from CLEs in hard ticks (family Ixodidae) (Gottlieb et al. 2015; Smith et al. 2015; Tsementzi et al. 2018; Guizzo et al. 2017), stymieing earlier efforts to understand *C. burnetii* evolution. Here, by sequencing the first soft-tick CLE genome, we were able to build a more definitive phylogenomic tree, which confirmed that CLEOA is a sister taxon of *C. burnetii* (Fig. 1; Supplemental Table S1). In addition, the presence of pseudogenized T4BSS genes in CLEOA indicates that the tick-symbiont evolved from a pathogenic ancestor with a functional T4BSS (Fig. 2; Supplemental Table S2).

**Figure 1.**
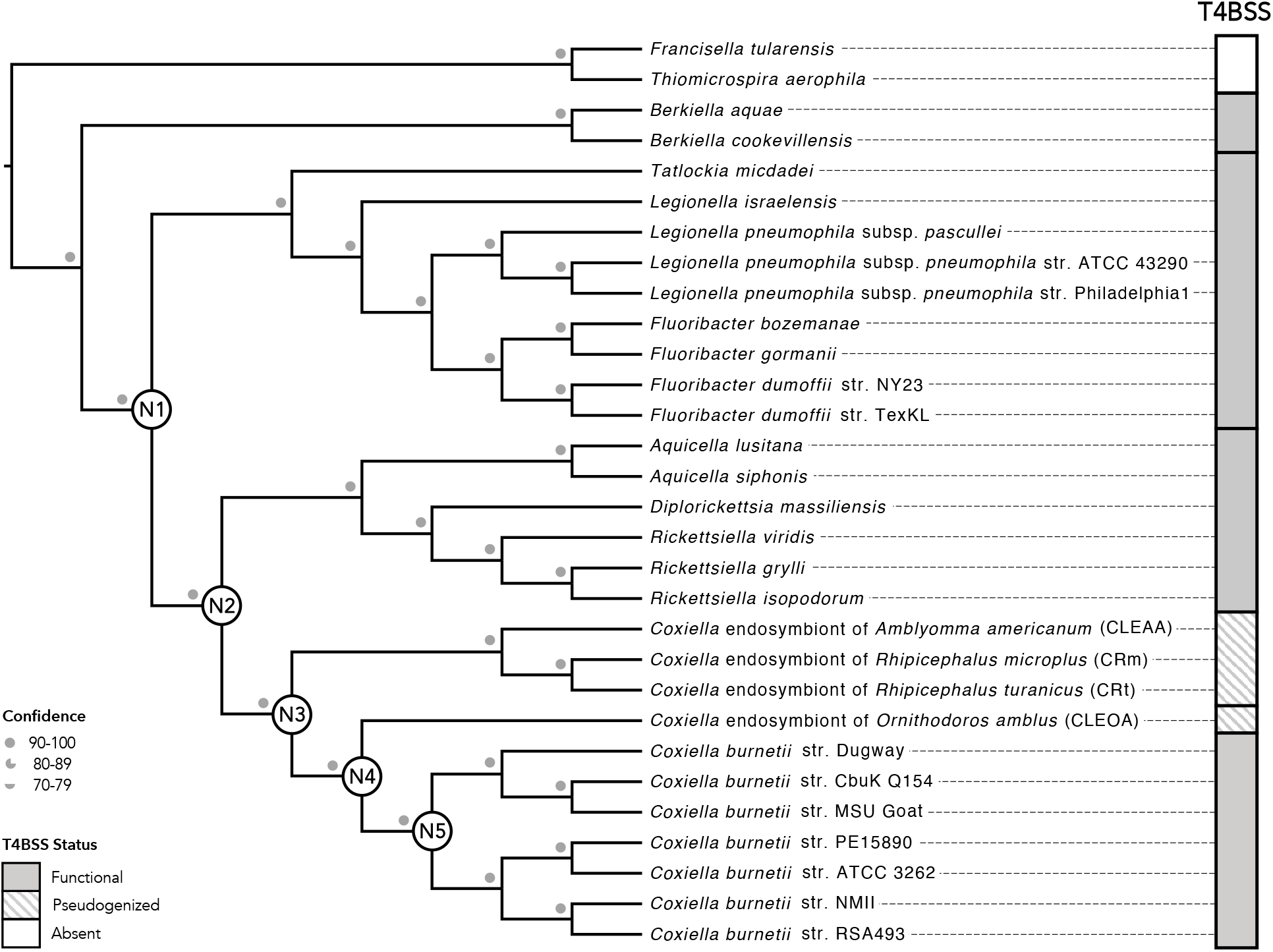
CLEOA is the closet relative of *C. burnetii*. Maximum likelihood and Bayesian trees built using 117 single-copy protein-coding genes were combined to generate the shown phylogenomic tree. Bootstrap support and posterior probabilities agreed at all branchpoints and are depicted as a single confidence value. The Dot/Icm Type IVB secretion system (T4BSS), which is critical to pathogenesis, is found in all members of the order Legionellales, but has been pseudogenized in CLEs. Nodes N1-N5 mark major branching points in the evolution of *C. burnetii*.

**Figure 2.**
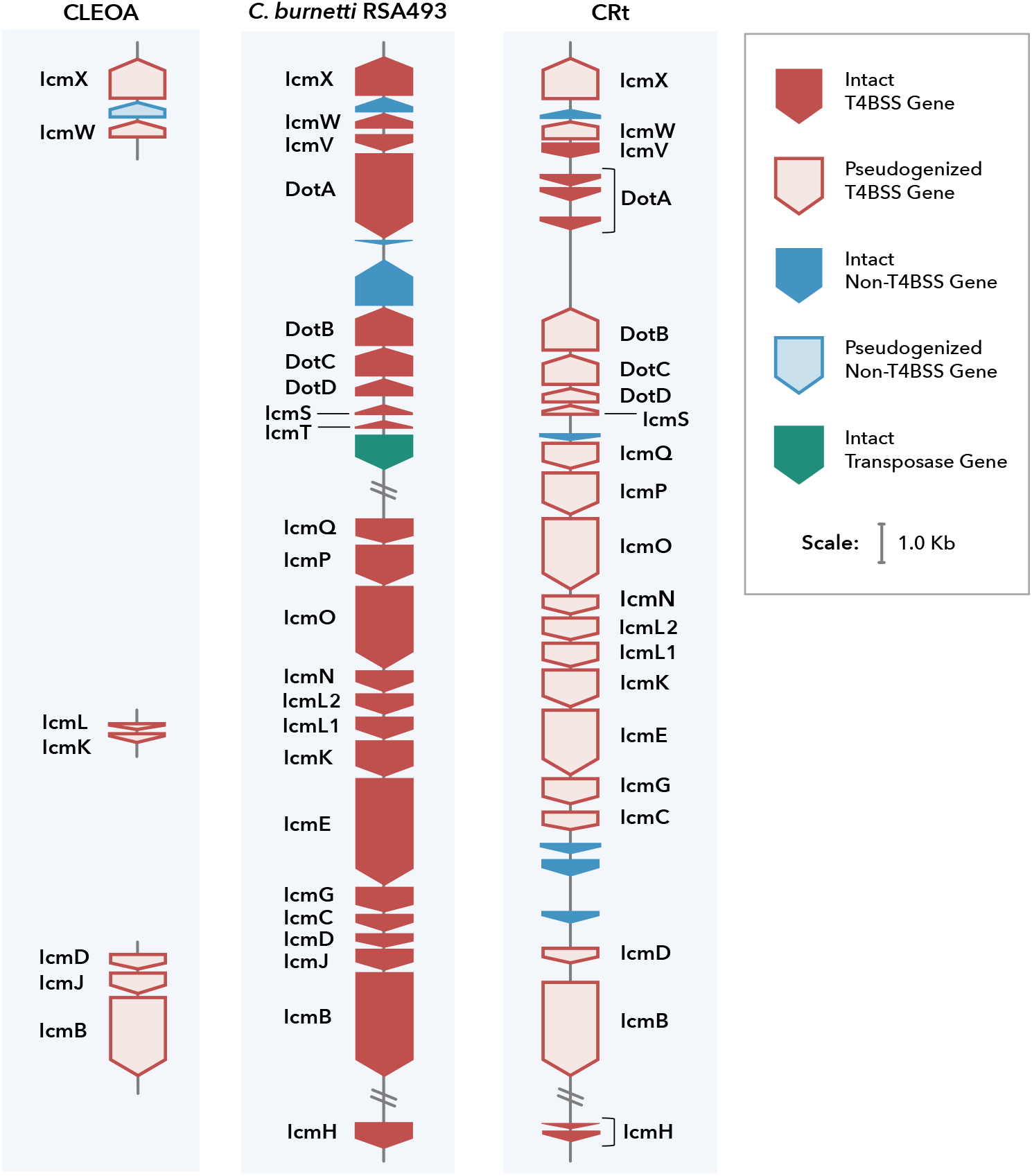
CLEs contain nonfunctional T4BSS. Comparison of T4BSS genes in CLEOA, *C. burnetii* RSA493 and CRt indicate that the secretion system has been rendered nonfunctional in tick endosymbionts. Filled blocks represent intact genes. Outlined blocks represent pseudogenized genes.

To resolve the ancestry of pathogenicity in Legionellales we determined the prevalence of T4BSS, which is an essential virulence factors in this group of bacteria that includes human pathogens (*C. burnetii* and *L. pneumophila*), opportunistic pathogens (*Rickettsiella*), and symbionts (CLEs). Our analyses revealed that the secretion system is intact in all members of this order with the exception of CLEs, which only contained remnants of the T4BSS (Figs. 1, 2). The most parsimonious explanation for this phyletic pattern is that T4BSS was present in the common ancestor of all Legionellales and was later lost in lineages that gave rise to CLEs, including CLEOA.

### Multiple CLEs have evolved from pathogens

*Coxiella* detected in ticks are classified into four clades, three of which contain intermingled pathogens and endosymbionts (Fig. 3; Supplemental Table S3; Duron et al. 2015a). Clade A includes *C. burnetii* — the human pathogen, and CLEOA, which arose from a pathogenic ancestor, as discussed above. In Clade B, CLEs of *Haemaphysalis* ticks are present along with a presumably pathogenic *Coxiella* that caused horse infection (Seo et al. 2016). Clade C has CRt, a pathogen-derived endosymbiont, along with strains that caused opportunistic human skin infections (Gottlieb et al. 2015; Angelakis et al. 2016; Guimard et al. 2017; Tsementzi et al. 2018; Ben-Yosef et al. 2020). Only Clade D, which contains small-genomed CLEs (e.g., CLEAA), has no known pathogenic representatives. This phylogenetic pattern of endosymbionts clustering with pathogens indicate that, similar to the pathogenic ancestry of CLEOA and CRt, CLEs of several other ticks have also evolved from pathogenic coxiellae. Thus, based on phylogenetic and T4BSS distribution patterns, we surmise that *Coxiella* strains that infect vertebrates (e.g., humans, horses, birds) and invertebrates (e.g., crayfish) are widespread across the globe (Fig. 3), and ticks have converted several of them into beneficial endosymbionts.

**Figure 3.**
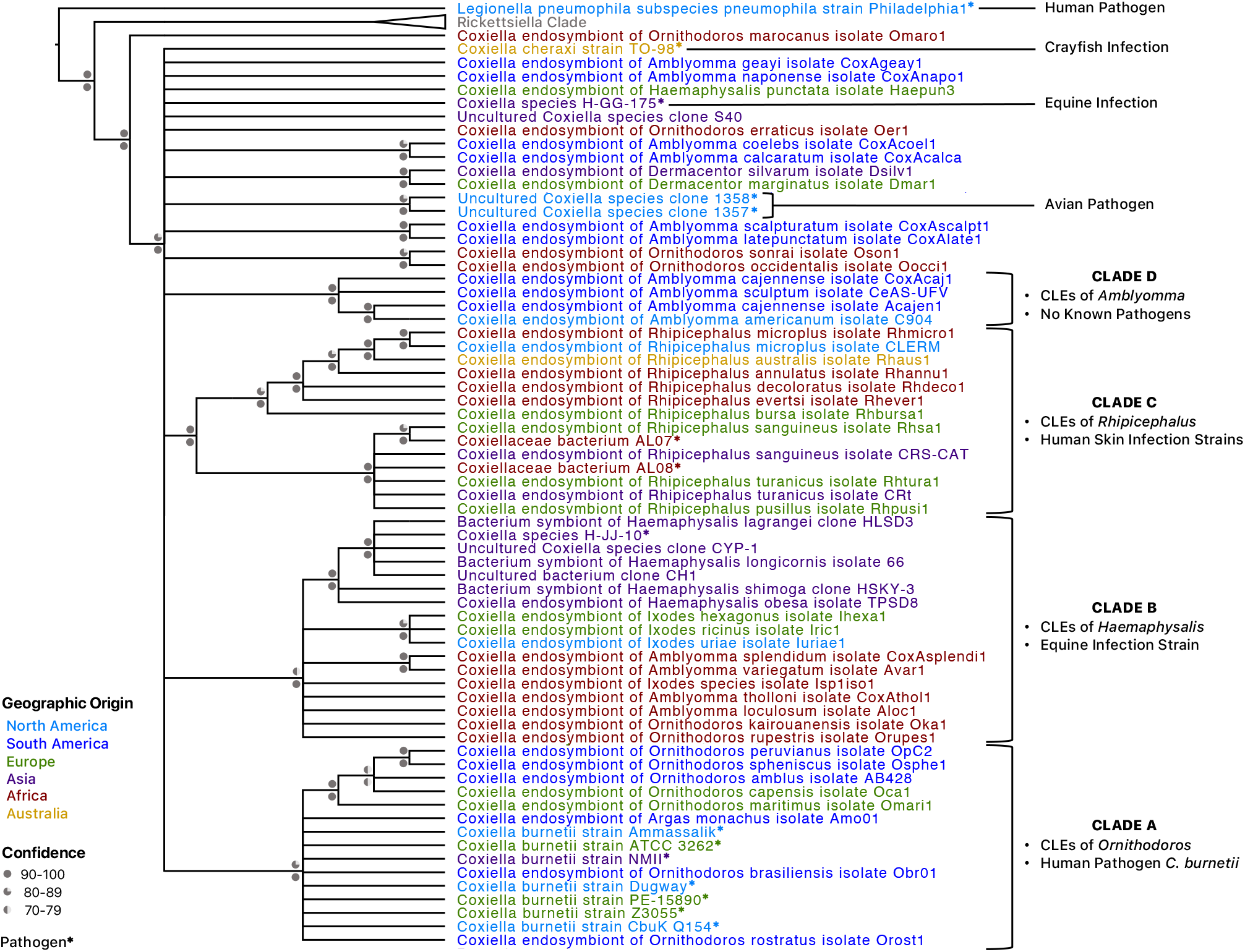
CLE clades contain both tick endosymbionts and pathogens. A 16S rDNA-based phylogenetic tree is shown. Bootstrap support and posterior probabilities are labeled above and below branchpoints, respectively. Nodes with ≤70% bootstrap support were collapsed to polytomies. Taxa colors represent the continent from which the host was derived. Established pathogens are marked with asterisks. Clades A-D were originally defined by Duron et al. 2015a.

### HGT was a major contributor to gene accumulation in *C. burnetii*’s ancestors

In order to better understand the evolution of *C. burnetii*, we traced the ancestry of its protein-coding genes by determining whether their orthologs — either functional or pseudogenized — were present in other Legionellales members. Out of 1,530 protein-coding genes whose ancestries we could trace, 790 were deemed to be ancestral, meaning it was present in the ancestor that diverged from *Legionella* (Node 1), and an additional 585 genes originated in Nodes 2-4 (Fig. 4; Supplemental Table S4). This data demonstrates that the common ancestor of *C. burnetii* and CLEOA contained most of the genes, including virulence factors, present in *C. burnetii*, and was hence well equipped to infect mammals.

**Figure 4.**
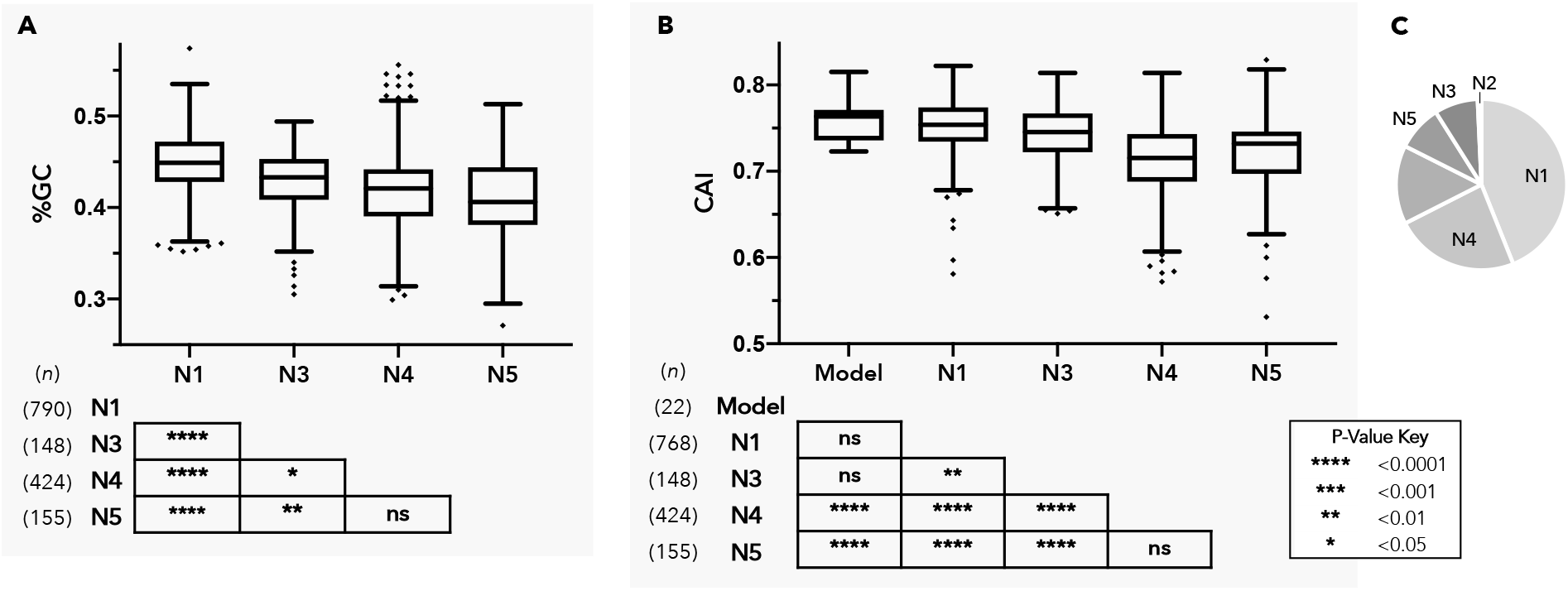
HGT was a major contributor to gene accumulation in *C. burnetii*’s ancestors. Nodes N1-N5 (as labeled in Fig. 1) represent major branchpoints in the evolution of *C. burnetii*. (A) and (B) depict %GC and Codon Adaptation Index (CAI) distributions, respectively, for genes originating at each node. Boxes illustrate each distribution’s interquartile range while the black line dividing the box represents the median. Whiskers represent minimum and maximum values, excluding outliers (black diamonds) which were determined using the Tukey method. P-values shown in tables are for pairwise T-tests (pooled SD, BKY adjusted). Genes gained at N2 were excluded due to small sample size (*n*=13). (C) *C. burnetii* genome composition based on nodes of gene origin: N1: 43.9%, N2: 0.7%, N3: 8.2%, N4: 22.6%, N5: 8.6%. The unlabeled portion represents potentially spurious genes (*n*=224) as well as genes with undefined nodes of origin (*n*=44).

A major impediment to unspooling the evolutionary history of *C. burnetii* is the sparse availability of Coxiellaceae genomes, which makes it difficult to ascertain whether genes were gained by *C. burnetii*’s ancestors at a Nodes 2-5 or were instead lost in other bacteria represented at each node. To overcome this difficulty, we calculated each *C. burnetii* gene’s nucleotide composition (%GC) and Codon Adaptation Index (CAI), two measures known to distinguish foreign-origin genes from ancestral ones (Fig. 4; Supplemental Table S5; Sharp and Li 1987; Lawrence and Ochman 1997; Jansen et al. 2003; Raghavan et al. 2012). Both %GC and CAI values for genes that originated in Nodes 3-5 were significantly different from those of ancestral (Node 1) genes, indicating that a considerable portion of these genes were acquired horizontally. [Node 2 genes were excluded from this analysis due to small sample size (*n*=13).] Interestingly, %GC and CAI values for Node 5 genes were not significantly different from those gained at Node 4, suggesting that many of the genes currently found only in the human pathogen were present in the common ancestor of *C. burnetii* and CLEOA, and were later lost in the tick endosymbiont. However, it is clear that HGT has contributed to the accumulation of genes at Node 5 as well because 88 out of 155 genes in this category showed phylogenetic patterns consistent with HGT (Supplemental Fig. S1, Table S6). Cumulatively, our results validate the important role HGT has played in assembling *C. burnetii*’s protein repertoire (Moses et al. 2017), and show that this process occurred principally in ancestors that preceded the *C. burnetii*-CLEOA split.

### CLEOA potentially provides *O. amblus* with nutrients missing in vertebrate blood

Similar to other hematophagic organisms (Duron and Gottlieb 2020), ticks likely obtain nutrients missing in blood from endosymbiotic bacteria such as CLEs and FLEs (Gottlieb et al. 2015; Smith et al. 2015; Gerhart et al. 2016, 2018; Duron et al. 2018b; Tsementzi et al. 2018). An analysis of the CLEOA genome revealed that a large number of genes have been inactivated in the endosymbiont as part of its adaptation to an obligately host-dependent lifestyle (Table 1). Nevertheless, the bacterium has retained complete pathways for the synthesis of several vitamins and cofactors, including those for the biosynthesis of biotin (vitamin B7), folic acid (B9), riboflavin (B2), FAD, CoA, and lipoic acid. These pathways are also conserved in all CLEs and FLEs (Fig. 5), suggesting that CLEOA and other tick endosymbionts provide one or more of these metabolites to their respective hosts.

**Table 1.** Genome characteristics of CLEOA and *C. burnetii*.

|  | CLEOA | *C. burnetii* RSA493 |
| --- | --- | --- |
| NCBI Accession | VFIV00000000 | AE016828.3 |
| Length (bp) | 1,561,173 | 1,995,488 |
| %GC | 40.6 | 42.7 |
| rRNA (5S, 16S, 23S) | 1,1,1 | 1,1,1 |
| tRNA | 42 | 42 |
| Functional genes | 889 | 1,798 |
| Pseudogenes | 660 | 197 |
| Single copy genes<sup>a</sup> | 106/111 | 106/111 |

<sup>a</sup> Albertson et al. 2013

**Figure 5.**
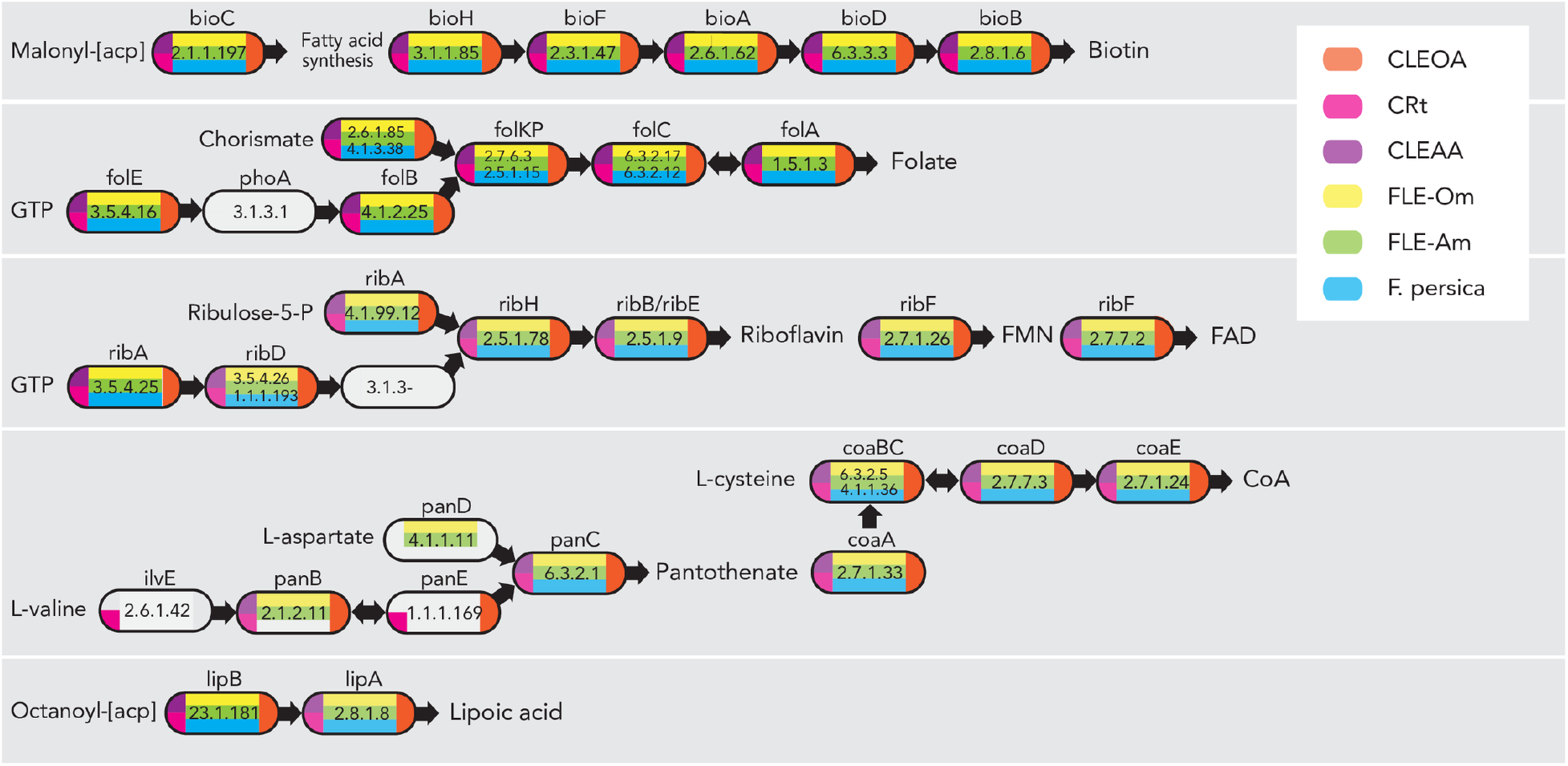
CLEs and FLEs encode B vitamin and cofactor biosynthetic pathways. CLEOA, CRt, CLEAA, FLE-Om (*Francisella* endosymbiont of *Ornithodoros moubata*; LVCE00000000), FLE-Am (*Francisella* endosymbiont of *Amblyomma maculatum*; LNCT00000000), and *F. persica* (FLE in *Argas arboreus*; NZ_CP013022) contain intact pathways for the synthesis of B vitamins and cofactors. Enzymes catalyzing each step are labeled with gene names and EC numbers. Blocks with no color indicate that a functional copy of a gene was not detected in that genome. Depictions of metabolic pathways modified from Gerhart et al. 2018.

In addition to the vitamin and cofactor biosynthesis genes, CLEOA contains 91 genes that have been deactivated or lost in *C. burnetii* (Supplemental Table S7). It is likely that many of these CLEOA-specific genes have functions in the tick ecosystem but are not useful during mammalian infections. Collectively, based on its genomic features (Table 1), recent loss of T4BSS (Fig. 2), and presence of intact pathways for synthesizing B vitamins and cofactors (Fig. 5), we conclude that a pathogenic *Coxiella* was recruited to function as a nutrient-provisioning endosymbiont in *O. amblus*.

#### CLEOA has not retained genes that allow *C. burnetii* to withstand the harsh environs of CCV Loss of T4BSS and associated genes

Because *C. burnetii* diverged from CLEOA recently, it provided us an opportunity to perform a thorough comparison of the human pathogen’s genome to that of a closely-related non-pathogen. Our analysis identified 980 functional genes in *C. burnetii* whose orthologs have been either pseudogenized or deleted in CLEOA (Supplemental Table S8). Among them are genes for T4BSS and for effectors secreted through T4BSS (Fig. 2; Supplemental Table S2) that enable *C. burnetii* to generate its intracellular replicative niche, termed *Coxiella*-containing vacuole (CCV) (Chen et al. 2010; Beare et al. 2011; Martinez et al. 2014, 2020; Newton et al. 2014). In addition, genes for PmrAB and EirA that control T4BSS activity, and for eight T4BSS effectors present on *C. burnetii*’s QpH1 plasmid have been inactivated in CLEOA (Maturana et al. 2013; Beare et al. 2014; Kuba et al. 2020). Therefore, the secretion system, which is critical to the intra-macrophage growth of *C. burnetii*, is clearly not required for CLEOA to grow within tick cells.

#### Loss of transporters of antibacterial molecules

To protect itself from noxious molecules produced by the host, *C. burnetii* likely depends on transport proteins that efflux harmful substances out of its cytoplasm. For instance, macrophages increase Cu^2+^ concentration within phagosomes to kill intracellular bacteria (Neyrolles et al. 2015), and *C. burnetii* probably utilizes a P-1B type ATPase to export copper from its cytoplasm to sustain intracellular growth (Rowland and Niederweis 2012). This ATPase, along with 18 others of unknown function, have been pseudogenized or deleted in CLEOA (Supplemental Table S9). In addition, *C. burnetii* encodes 25 putative transporters that could facilitate the pathogen’s growth within CCV, out of which, 19 have become non-functional in CLEOA (Supplemental Table S9), suggestive of the mild nature of the tick endosymbiont’s intracellular compartment where antibacterial molecules are probably not a major threat.

#### Diminished pH regulation

A defining feature of CCV is its acidity (pH ~4.75) (Vallejo Esquerra et al. 2017; Samanta et al. 2019). To compensate, *C. burnetii* utilizes several mechanisms to maintain its cytoplasmic pH close to neutral (Hackstadt 1983), many of which have been rendered non-functional in CLEOA. For example, the enzyme carbonic anhydrase that catalyzes the production of bicarbonate (HCO3-) buffer from CO2 (Vallejo Esquerra et al. 2017; Bury-Moné et al. 2008) has been pseudogenized in CLEOA (Supplemental Table S10). Another pH-regulating strategy used by *C. burnetii* is to remove excess protons in its cytoplasm via proton-antiporters such as the multi-protein Mrp antiporter, a pair of Na^+^/H^+^ antiporters, and a K^+^/H^+^ antiporter. *C. burnetii* also encodes a pair of glutamate/gamma-aminobutyrate (GABA) antiporters that export GABA in exchange for glutamate, thereby reducing the cytoplasmic proton content. The Mrp antiporter, one of the two Na^+^/H^+^ antiporters, and both glutamate/GABA antiporters have been pseudogenized in CLEOA (Supplemental Table S10). A hallmark of *C. burnetii* is its unusually high number of basic proteins (~46% of proteins have pI values ≥9; average pI 8.22) that could function as a “proton sink,” which allows the pathogen to maintain its cytoplasmic pH close to neutral (Seshadri et al. 2003). In contrast, only ~39% of proteins in CLEOA have pI values ≥9 (average pI 8.0), again illustrating a lack of acidic stress within CLEOA’s intracellular vacuole. Collectively, our data suggest that the endosymbiont does not face the constant threat of excess protons entering its cytoplasm, probably because its intracellular niche, unlike *C. burnetii*’s, has a pH closer to neutral.

#### Loss of cell membrane and cell wall genes

In gram-negative bacteria, inner and outer membranes along with peptidoglycan play important roles in stress response (Rowlett et al. 2017). In CLEOA, the gene that encodes PlsC, which converts lysophosphatidic acid into phosphatidic acid (PA), a universal intermediate in the biosynthesis of membrane phospholipids, has been pseudogenized. The *plsC* gene is essential in *Escherichia coli*, and a transposon insertion in this gene in *C. burnetii* caused severe intracellular growth defect (Coleman 1990; Martinez et al. 2014); hence, it is not clear how CLEOA is able to build its membranes without a functional *plsC*, but one possibility is that the endosymbiont utilizes PA obtained from its host. Another membrane-associated loss of function in CLEOA is the pseudogenization of the *pldA* gene that encodes phospholipase A (PldA), which is critical to *C. burnetii’*s outer membrane function and for optimal growth within macrophages (Stead et al. 2018). As for its peptidoglycan, CLEOA contains intact genes for D, D-transpeptidases (also known as penicillin-binding proteins) that catalyze 4-3 peptide cross-links between D-alanine and diaminopimelate; however, all L, D-transpeptidase genes (annotated as “enhanced entry proteins”) have been pseudogenized, indicating that the tick symbiont does not have the ability to generate 3–3 cross-links between diaminopimelate molecules in its peptidoglycan. These nonclassical cross-links contribute to *C. burnetii*’s environmental stability (Sandoz et al. 2016), and are probably not critical to CLEOA because the endosymbiont is passed vertically from one generation to next. Collectively, as observed in other endosymbionts (Nakabachi et al. 2006; McCutcheon and Moran 2010; Chong and Moran 2018), CLEOA lacks numerous proteins that are typically considered integral to the optimal functioning of bacterial cell membrane and cell wall.

#### Loss of antioxidant genes

An intricate network of antioxidants allows *C. burnetii* to thrives in a phagolysosome-derived intracellular vacuole (Mertens and Samuel 2012). In contrast to CCV, oxidative stress appears to be minimal in CLEOA’s intracellular vacuole because the endosymbiont contains only a streamlined version of *C. burnetii*’s antioxidant defense system. For instance, *C. burnetii* contains two superoxide dismutases (SODs), but CLEOA has retained the cytoplasmic Fe-containing SodB, but not SodC, the periplasmic Cu/Zn-SOD. OxyR, the master regulator of peroxide stress, along with a catalase, a peroxidase (AhpC2), a methionine sulfoxide reductase, a hemerythrin-like protein, and a glutathione transferase that together help mitigate oxidative stress have also been deactivated in CLEOA (Supplemental Table S11). In addition, *C. burnetii*, but not CLEOA, has the ability to synthesize queuine, a guanine analog, found in the first anticodon position of several post-transcriptionally modified tRNAs (Iwata-Reuyl 2003). The precise functions of queuine is not understood, but it is thought to promote the activity of antioxidant enzymes, including catalase, superoxide dismutase, and glutathione transferase, most of which, as mentioned above, have lost their functionality in CLEOA (Koh and Sarin 2018).

*C. burnetii* utilizes both cytochrome bd (encoded by genes *cydABX*) and cytochrome o (encoded by genes *cyoABCD*) as terminal oxidases, but CLEOA has only retained cytochrome o genes. Cytochrome bd, which also functions as a quinol peroxidase that prevents the buildup of oxidative free radicals (Endley et al. 2001; Omsland and Heinzen 2011), has become nonfunctional in the tick endosymbiont. In addition, CLEOA does not encode genes for an acid phosphatase and two sterol reductases that likely modify host proteins and cholesterol, respectively, to protect *C. burnetii* from host-induced oxidative stress (Seshadri et al. 2003; Gilk et al. 2010; Hill and Samuel 2011; Gilk 2012). Finally, *C. burnetii* is thought to compensate for the lack of the oxidative branch of Pentose Phosphate Pathway (PPP)— a major source of NADPH, by utilizing alternative NADPH-regenerating enzymes such as short chain dehydrogenases and sterol reductases, and by salvaging NAD^+^ from the host (Bitew et al. 2018, 2020). In CLEOA, all four short chain dehydrogenases, the two eukaryote-like sterol reductases, and the nicotinate-salvaging protein have become nonfunctional. In total, while the human pathogen contains several mechanisms to defend against oxidative stress, most of these antioxidant systems have been lost in CLEOA, most likely due to minimal oxidative stress experienced by the bacterium within tick cells. Collectively, the loss of T4BSS, transporters, pH regulation, cell wall modification, and antioxidant defense in CLEOA show that its intracellular vacuole is a less stressful place to live than the phagolysosome-derived CCV occupied by *C. burnetii*.

### Heme analog inhibits *C. burnetii* growth

Cytochromes require heme as a cofactor, but CLEOA does not contain a functional heme biosynthesis pathway, which is present in *C. burnetii* (Supplemental Table S12). The only intact heme biosynthesis gene in CLEOA is *ctaB*, which encodes an enzyme that converts heme b to heme o, a component of cytochrome o — the sole terminal cytochrome oxidase present in CLEOA (Saiki et al. 1992). Based on this evidence, the endosymbiont appears to import heme b from the tick hemocoel (vertebrate hemoglobin contains heme b) and converts it to heme o using the *ctaB*-encoded protoheme IX farnesyltransferase. Additionally, while *C. burnetii* can import ferrous iron released from iron-containing host molecules such as ferritin and transferrin (Sanchez and Omsland 2020), free Fe^2+^ does not seem to be important for CLEOA’s intracellular growth because the iron transporter FeoB has been pseudogenized, suggesting that host-derived heme b serves as the tick endosymbiont’s heme and iron source.

The heme biosynthesis pathway, while absent in CLEOA, is conserved in all strains of *C. burnetii*, probably because the iron-protoporphyrin molecule is critical to the pathogen’s ability to grow within human macrophages (Moses et al. 2017). We tested *C. burnetii*’s dependence on heme by treating both axenically grown and intracellular *C. burnetii* with gallium protoporphyrin IX (GaPPIX), which can replace heme in cytochromes and other heme-containing enzymes (Hijazi et al. 2017, 2018). As shown in Figure 6, ≥250 nM of GaPPIX caused significant inhibition of *C. burnetii* growth in ACCM-2, and treatment with ≥2μM of GaPPIX resulted in significant growth impairment of *C. burnetii* within THP-1 cells. Reassuringly, only GaPPIX concentrations of ≥512μM caused cytotoxicity in THP-1 cells (Supplemental Fig. S2), indicating that gallium compounds could potentially be used to treat *C. burnetii* infections.

**Figure 6.**
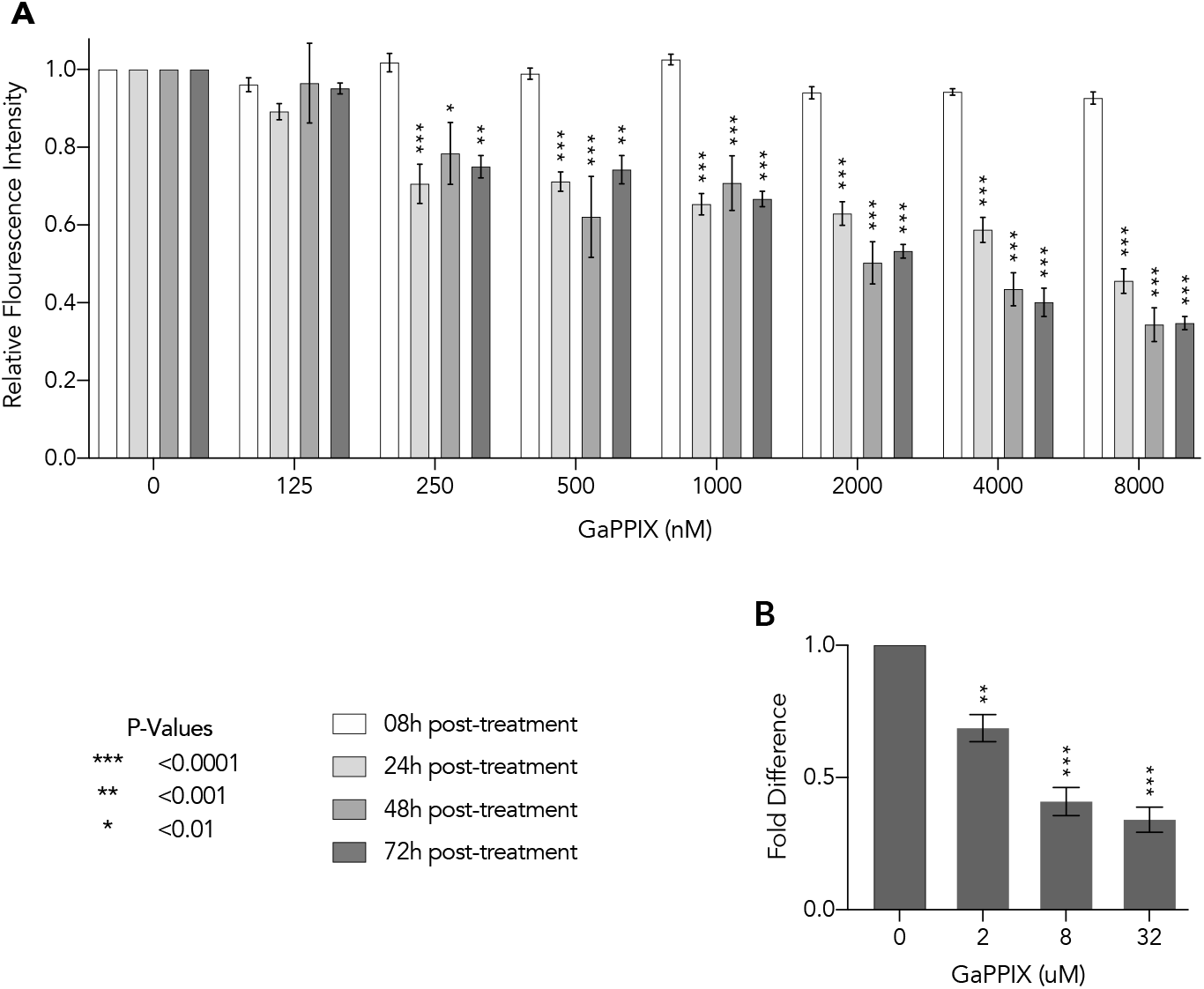
A heme analog reduces *C. burnetii* growth. (**A)** Bacteria growing in ACCM-2 were exposed to concentrations of GaPPIX shown in x-axis and were quantified using PicoGreen at 8h, 24h, 48h, and 72h post-treatment. Data shown are mean fluorescence intensity (± SE) compared to the vehicle control (0nM). Statistical significance was analyzed using two-way repeated measures ANOVA followed by Dunnett’s test (n=5). (**B)** At 72h post-treatment, bacterial growth within THP-1 cells was quantified using qPCR. Data shown as mean fold difference (± SE) compared to control. Statistical significance was analyzed using one-way ANOVA followed by Dunnett’s test (n=3).

## DISCUSSION

Although symbiotic and pathogenic coxiellae associated with ticks are found across the globe, it is not clear how pathogenesis and symbiosis evolved in this group of bacteria. Here we show that CLEOA, a soft-tick symbiont, and *C. burnetii*, a human pathogen, evolved recently from a common ancestor that contained genes necessary to infect macrophages. Additionally, while HGT contributed significantly to the evolution of *C. burnetii*, it occurred in ancestors prior to the divergence of CLEOA and *C. burnetii* lineages. These discoveries clarify the evolution of *C. burnetii*, which previously was thought to have evolved from an avirulent tick endosymbiont by gaining virulence factors via HGT. We further show that the pathogenic ancestry of *C. burnetii* and CLEOA fits into a general pattern of tick endosymbionts originating from erstwhile pathogens, thereby revealing the ability of ticks to convert pathogenic bacteria into beneficial endosymbionts. Lastly, by comparing the genomes of *C. burnetii* and CLEOA, we were able to gain new insights into the intracellular biology of both bacteria, and show that metabolic pathways retained only in the human pathogen are promising targets for the development of new treatments against Q fever.

### Emergence of tick-symbionts from virulent ancestors

*Coxiella* species related to CLEs infect a wide range of animals (Shivaprasad et al. 2008; Woc-Colburn et al. 2008; Angelakis et al. 2016; Seo et al. 2016; Guimard et al. 2017; Elliman and Owens 2020; Needle et al. 2020), but these infectious strains are not the closest relatives of *C. burnetii*; instead, the human pathogen’s closest relative is the soft-tick symbiont CLEOA. Akin to the CLEOA-*C. burnetii* relationship, CRt, the endosymbiont in *R*. *turanicus*, is closely related to pathogenic *Coxiella* (termed “*Candidatus* Coxiella massiliensis”) isolated from human skin infections, and a strain of *Coxiella* isolated from horse blood is closely related to CLEs present in *Haemaphysalis* ticks (Angelakis et al. 2016; Seo et al. 2016; Guimard et al. 2017). In addition to these pathogens, bacteria related to CLEs have repeatedly caused fatal bird and crayfish infections (Fig. 3; Shivaprasad et al. 2008; Woc-Colburn et al. 2008; Elliman and Owens 2020; Needle et al. 2020). Microscopic and histological data from avian infections demonstrated that the bacteria have the ability to generate CCV-like compartments within macrophages, and both avian and human skin infection strains have “small-cell” and “large-cell” morphologies — two distinct characteristics of *C. burnetii* — suggesting that the bacteria are genuine vertebrate pathogens (Shivaprasad et al. 2008; Woc-Colburn et al. 2008; Angelakis et al. 2016; Guimard et al. 2017; Needle et al. 2020). Further research, including sequencing their genomes, is required to elucidate the biology and pathogenicity of these infective strains and to understand why only one, i.e., *C. burnetii*, among several virulent lineages have evolved into a bona fide human pathogen.

### Tick endosymbionts are ephemeral

Phylogenies of only a few CLEs are congruent with those of their hosts (Duron et al. 2017; Binetruy et al. 2020), probably because older CLEs get replaced by newer CLEs derived from distantly-related coxiellae. In a similar fashion, FLEs seem to have replaced older CLEs in several tick lineages (Gerhart et al. 2016, 2018; Duron et al. 2017, 2018b). This ephemeral nature of CLEs is surprising because hematophagic arthropods typically need a reliable partner to gain nutrients that are scarce in vertebrate blood. Insects such as bedbugs and body lice that face similar nutrient scarcity have evolved stable long-term relationships with endosymbionts (Perotti et al. 2007; Hosokawa et al. 2010). It is not clear why that is not the case in ticks, but one possibility is that ticks do not need to establish long-term relationships because they frequently encounter bacteria that are predisposed to becoming nutrient-provisioning endosymbionts. For instance, both *Coxiella* and *Francisella* have the metabolic wherewithal to provide nutrients missing in vertebrate blood, thereby improving tick fitness and promoting the establishment of symbiotic relationships. Additionally, both bacteria are capable of infecting tick ovaries, which ensures their vertical transfer, and thus the maintenance of symbiotic relationships. Furthermore, tick behavior such as hyperparasitism, wherein an unfed tick steals blood by piercing the integument of a newly-fed tick, could promote the horizontal spread and eventual establishment of a new endosymbiont in a population (Helmy et al. 1983; Labruna et al. 2007; Llanos-Soto et al. 2019). The plentiful supply of potential endosymbionts in its environment could also obviate the need for ticks to evolve specialized symbiont-carrying organs (Perotti et al. 2007; Hosokawa et al. 2010, 2012; Rio et al. 2012; Simonet et al. 2018; Husnik et al. 2020), thereby explaining the absence bacteriomes in ticks. Alternatively, malpighian tubules, which extends into most parts of a tick’s body cavity, and to which CLEs and FLEs show strong tissue tropism, could be functioning as a stand-in for bacteriome (Klyachko et al. 2007; Machado-Ferreira et al. 2011; Lalzar et al. 2014; Duron et al. 2018b).

Another reason for the unstable nature of CLE-tick relationships could be that the constant turnover of endosymbionts protects ticks from being dependent on an endosymbiont with reduced nutrient-provisioning capability. For instance, when a free-living bacterium evolves into an obligate endosymbiont, its genome undergoes massive gene loss due to a combination of factors, including reduced selection to maintain superfluous genes, genetic drift and possibly adaptive evolution (Moran 1996; Clayton et al. 2016). This degenerative evolution, as observed in CLEOA, is a convergent feature of heritable symbionts and could result in the irreversible loss of fitness-improving genes, including genes that form the primary basis for symbiosis (Lambert and Moran 1998; Andersson and Andersson 1999; McCutcheon and Moran 2010). If no compensatory mechanisms are undertaken, this progressive gene loss could result in the extinction of both the tick and the endosymbiont. One strategy to avoid this fate is to periodically replace resident endosymbionts with new bacteria that have the needed metabolic capabilities (Russell et al. 2017; Bennett and Moran 2020).

Gaining new symbionts via horizontal transmission could also protect ticks from becoming dependent on a degraded endosymbiont. While the mechanistic details of this process are not understood, phylogenetic patterns of CLE and FLE distribution strongly indicate the occurrence of horizontal transmission of bacteria between ticks (Gerhart et al. 2016, 2018; Duron et al. 2017; Binetruy et al. 2020). It is not clear whether there is direct tick-to-tick transfer or whether unrelated ticks acquire similar bacteria from common environmental sources (Duron et al. 2018a). In either case, tick-endosymbiont relationships appear to be much more short-lived and dynamic than most insect-endosymbiont relationships. It should be noted however that not all tick-endosymbionts are short-lived. Ticks that carry CLEs belonging to Clade D (Fig. 3) appear to have established long-term relationships with their endosymbionts. For instance, CLEs in *A. americanum* and *A. sculptum* have highly reduced (~0.60 Mbp) genomes that are similar in size to *Buchnera*, which established its symbiosis with aphids more than 200 million years ago (Moran et al. 1993). Putting all this information together, it appears that a combination of vertical inheritance, horizontal transmission, and periodic replacement of old symbionts with new pathogen-derived symbionts underlies the complex distribution pattern of endosymbionts observed in ticks (Gottlieb et al. 2015; Smith et al. 2015; Gerhart et al. 2016, 2018; Duron et al. 2017; Tsementzi et al. 2018; Binetruy et al. 2020).

### Functions of CLEs and FLEs

While the exact functions of FLEs and CLEs have not been fully characterized, previous studies have shown that they are often the sole bacterium present in long-term laboratory tick colonies, an indication that the bacteria are essential to ticks’ wellbeing. In addition, removal of the resident endosymbiont via antibiotic treatment reduced tick fitness, which was reversed when ticks were provided with B vitamins (Zhong et al. 2007; Smith et al. 2015; Gerhart et al. 2016; Guizzo et al. 2017; Zhang et al. 2017; Duron et al. 2018b; Li et al. 2018; Ben-Yosef et al. 2020). Our genome analyses support a B vitamin-provisioning role for tick endosymbionts because the genes required to synthesize B7 (biotin), B9 (folic acid), and B2 (riboflavin), and three coenzymes (FAD, CoA, and lipoic acid) are conserved in all CLEs and FLEs. Future experiments should clarify whether any or all of these nutrients form the basis for the CLE-tick and FLE-tick symbioses.

### Pathogen-specific metabolic processes are potential targets to control Q fever

Genetic and physiological capabilities accumulated by a bacterium are critical to its ability to adapt to new environments, especially ones such as intra-macrophage vacuoles that do not facilitate the gain of new genes via HGT. In accordance with this idea, our analyses showed that virulence factors and metabolic genes utilized by *C. burnetii* to grow within CCV were present in the common ancestor of *C. burnetii* and CLEOA. Befitting its obligate endosymbiotic lifestyle, many of these genes have become non-functional in CLEOA, allowing us to identify metabolic processes that are likely critical to *C. burnetii*’s intracellular growth. One metabolite that is exclusively produced by the pathogen is heme, the iron-protoporphyrin required for oxidative phosphorylation, among other functions. To test the importance of heme to *C. burnetii*, we exposed the bacterium to GaPPIX, a Ga(III) complex of protoporphyrin IX. Ga(III) inhibits bacterial growth because it binds to biological complexes that normally binds to Fe(III), but under physiological conditions Ga(III) is not reduced to Ga(II), thereby disrupting essential redox-driven biological processes (Bernstein 1998). We chose GaPPIX over other gallium-based formulations because it could replace heme in cytochromes, is known to be bactericidal, and is not toxic to primary human fibroblasts and established cell lines (Stojiljkovic et al. 1999; Arivett et al. 2015; Hijazi et al. 2018). *C. burnetii* lacks homologs of known heme transporters (Moses et al. 2017), hence, it is not clear how GaPPIX enters into the pathogen, but our growth assays clearly demonstrated that the heme analog is very effective at inhibiting both axenic and intracellular growth of *C. burnetii* (Fig. 6). Encouragingly, a recent human trial showed that Ga could improve lung function in people with cystic fibrosis and chronic *Pseudomonas aeruginosa* lung infections, and that the molecule worked synergistically with other antibiotics to inhibit bacterial growth (Goss et al. 2018). Thus, Ga, which has been approved by FDA for intravenous administration (Bonchi et al. 2014), and its derivatives such as GaPPIX, hold great promise as new therapeutic tools. In conclusion, this study provide novel insights into the evolution and biology of pathogenesis and symbiosis in *Coxiella*, and demonstrate that the new information could be applied to developing innovative therapeutic strategies.

## METHODS

### Genome sequencing and assembly

An *O. amblus* female, collected from soil underneath rocks near a *Spheniscus humbolti* (Humboldt penguin) nesting area in Isla Grande de Atacama, Chile, was identified as described in Clifford *et al.* 1980. DNA was extracted from the tick using DNeasy Blood & Tissue kit (Qiagen) and was submitted to Yale Center for Genome Analysis for Illumina (NovaSeq) sequencing. The resulting 150 bp paired-end reads were trimmed using Trimmomatic resulting in approximately 220 million read pairs of suitable quality (Bolger et al. 2014). The reads were assembled into contigs using metaSPAdes (Nurk et al. 2017), and open reading frames (ORFs) were identified using Prodigal (Hyatt et al. 2010). Contigs containing *Coxiella* genes were tentatively identified using BLASTn and BLASTp by comparing to a database of all publicly available sequences from Coxiellacea members. CONCOCT (Alneberg et al. 2014) was used for binning contigs based on coverage and k-mer composition, and these finding were merged with BLAST-based binning results. Approximately 20 million paired reads that mapped to contigs identified as containing *Coxiella* genes were used for a final metaSPAdes assembly resulting in a total of 101 contigs. The final collection of contigs was verified using hmmsearch (Potter et al. 2018) to identify essential single-copy genes (Albertsen et al. 2013), as well as RNammer (Lagesen et al. 2007) and tRNAscan-SE (Chan and Lowe 2019) to identify ribosomal and transfer RNAs, respectively. We were unable to stitch the contigs together into a closed chromosome because the contigs were filled with repeat sequences; however, the presence of 106 out of 111 highly conserved single-copy genes in both CLEOA and *C. burnetii* indicate that most of the CLEOA genome is represented in the assembled contigs. The final set of 101 contigs were submitted to NCBI (accession VFIV00000000) and annotated using the Prokaryotic Genome Pipeline.

### Phylogenetic analysis

Orthofinder (Emms and Kelly 2015) was utilized to identify 205 single-copy genes present in 52 representative species from the order Legionellales (Supplemental Tables S1, S13) in order to build the comprehensive phylogenomic tree Supplemental Figure S3. A subset of 117 genes conserved in 30 species (Supplemental Tables S1, S13) were used to generate Figure 1. For both trees, nucleotide sequences were aligned individually using global MAFFT (Katoh and Standley 2013) and were then concatenated. GBlocks (Talavera and Castresana 2007) was used to cull ambiguously aligned regions and jModelTest2 (Darriba et al. 2012) was used to select the appropriate model (GTR+I+G). Maximum Likelihood trees were generated using RaxML and Bayesian trees were produced using MrBayes (Ronquist et al. 2012; Stamatakis 2014). The 16s rDNA trees were built using the same process as above, with the final tree based on 1203 nucleotide positions, and nodes with less than 70% support collapsed. To confirm the HGT origin of Node 5 genes, homologs were identified via BLASTp (NCBI nr database, e-value ≤10e-5, identity ≥30% identity, coverage ≥70%). The nucleotide sequences of the homologs were collected into a database, and the Phylomizer pipeline (https://github.com/Gabaldonlab/phylomizer) was used to generate individual Maximum Likelihood trees using the 75 most closely related homolog sequences. Each tree was then compared to an NCBI Taxonomy-based tree to validate HGT (Supplemental Fig. S1).

### Determination of nodes of gene origin

The presence of functional homologs of *C. burnetii* RSA493 (AE016828.3) genes in other members of the order Legionellales was determined using BLASTp (identity ≥30%, coverage ≥70%, e-value ≤10e-5), and pseudogenized homologs were detected using tBLASTn (identity ≥30%, coverage ≥50%, e-value ≤10e-5). The presence/absence profile was utilized in Gain and Loss Mapping Engine (GLOOME) (Cohen et al. 2010) to determine the posterior probability of each gene’s presence at nodes N1-N5. For each *C. burnetii* gene, the node of origin was marked as the oldest node at which posterior probability was ≥0.7, with all subsequent nodes also having posterior probability of ≥0.7, as described previously (Peer and Margalit 2014).

### Calculation of CAI and pI

We identified 22 highly conserved single-copy protein-coding genes in *C. burnetii* that were highly expressed in both ACCM-2 and within human macrophages based on previous RNA-seq data (Supplemental Table S5; Warrier et al. 2014; Wachter et al. 2019). CodonW (http://codonw.sourceforge.net) was used to generate CAI values for the 22 genes in order to generate a model for optimal codon usage in *C. burnetii*, which was then compared to CAI values of sets of genes acquired at each node. All 22 genes used to build the model belonged to Node 1, and were not included in this analysis. Potentially spurious genes (n=224) that did not have any detectable homologs outside of *C. burnetii*, as well as genes with undetermined nodes of origin (n=44) were excluded from this analysis (Supplemental Table S14). Isoelectric points (pI) for all proteins in CLEOA and *C. burnetii* RSA493 (AE016828.3) were calculated using IPC (Kozlowski 2016).

### GaPPIX susceptibility assay

A 10mM GaPPIX (Frontier Scientific) solution was prepared in dimethyl sulfoxide (DMSO) and was stored at 4°C under dark conditions until further use. *C. burnetii* was cultured in ACCM-2 for 2 days at 37°C, 5% CO_2_ and 2.5% O_2_, and ~2×10^7^ genome equivalents were resuspended in fresh ACCM-2 containing 125nM, 250nM, 500nM, 1mM, 2mM, 4mM, or 8mM GaPPIX in 96-well black-bottom microplates (Greiner Bio-One). Bacterial growth was measured using PicoGreen (Invitrogen) as described previously (Moses et al. 2017).

THP-1 human monocytes (ATCC, TIB-202) were cultured in sterile RPMI-1640 medium (Gibco) supplemented with 1mM sodium pyruvate, 0.05 mM beta-mercaptoethanol, 1% Pen-Strep, and 4500 mg/L glucose with 10% heat-inactivated fetal bovine serum (FBS) at 37°C, 5% CO_2_ in 6-well tissue culture plates. Prior to infection, cells were differentiated into macrophages by treating with 30 nM phorbol 12-myristate 13-acetate (PMA) for 24h, followed by resting in PMA-free RPMI for 24h. Infection of THP-1 cells with *C. burnetii* was carried out using a 7d bacterial culture at a multiplicity of infection of 25. After briefly washing the cells with PBS, bacteria-containing medium was added to each well and gently centrifuged for 10 minutes followed by incubation at 37°C, 5% CO_2_ for two hours. To remove extracellular bacteria, cells were washed three times with PBS, and replaced with antibiotic-free RPMI and were incubated for 48h before treating with GaPPIX- (2uM, 8uM, and 32uM) or DMSO- (as control) containing media. After 72h, cells were washed three times with PBS and intracellular bacterial load was measure using qPCR, as we described previously (Moses et al. 2017). Potential cytotoxicity of GaPPIX was determined by measuring the levels of released lactate dehydrogenase (LDH) in cell supernatants using an LDH Cytotoxicity Assay Kit (Invitrogen).

## DATA ACCESS

The CLEOA genome generated in this study has been submitted to the NCBI Assembly database (https://www.ncbi.nlm.nih.gov/assembly/) under Whole Genome Shotgun (WGS) accession prefix VFIV, BioSample SAMN12040594, and BioProject PRJNA548565.

## ACKNOWLEDGEMENTS

We thank Samantha Fancher, Andrew Ashford and Jim Archuleta for technical help, and Daniel González-Acuña for assisting in tick collection. This project was supported in part by NIH grants AI123464, AI126385, and AI133023 to R.R.

## SUPPLEMENTAL FIGURE LEGENDS

**Supplemental Figure S1. HGT contributed to *C. burnetii* evolution.** 88 Node 5 genes have phylogenies (left) inconsistent with established taxonomic relationships (right), indicating that they were acquired via HGT. Branches and species titles are colored by taxonomy as specified on Page 1. (**A**) Maximum likelihood gene trees with bootstrap values labeled at branchpoints. *C. burnetii* RSA493 genes are indicated by their locus tags and are highlighted in blue. Duplicate genes are shown together in single trees. (**B**) Taxonomy-based trees depict the established relationships among the taxa featured in A. Taxa sharing the same NCBI taxonomic identifier number are displayed only once even if multiple representatives are present in A.

**Supplemental Figure S2. GaPPIX is not toxic to macrophages at levels that significantly inhibit *C. burnetii* growth.** At 24h post-GaPPIX treatment of THP-1 derived macrophages, lactate dehydrogenase (LDH) activity was determined by measuring the level of resorufin formation using an LDH cytotoxicity assay. The cytotoxicity was reported as the percentage LDH released compared to the maximum LDH activity. Data are shown as mean ± SEM percentage LDH released, and statistical significance was analyzed using one-way ANOVA followed by Dunnett’s test (n=3).

**Supplemental Figure S3. Legionellales phylogenomic tree.** Maximum likelihood and Bayesian trees were generated using 205 single-copy protein-coding genes conserved in all represented taxa (Supplemental Tables S1, S13). Bootstrap support and posterior probabilities are labeled above and below each node respectively. This tree was used in GLOOME (Cohen et al. 2010) to determine the nodes of gene origin.

## REFERENCES

Albertsen M, Hugenholtz P, Skarshewski A, Nielsen KL, Tyson GW, Nielsen PH. 2013. Genome sequences of rare, uncultured bacteria obtained by differential coverage binning of multiple metagenomes. Nat Biotechnol 31: 533–538.

Almeida AP, Marcili A, Leite RC, Nieri-Bastos FA, Domingues LN, Martins JR, Labruna MB. 2012. *Coxiella* symbiont in the tick *Ornithodoros rostratus* (Acari: Argasidae). Ticks Tick-Borne Dis 3: 203–206.

Alneberg J, Bjarnason BS, de Bruijn I, Schirmer M, Quick J, Ijaz UZ, Lahti L, Loman NJ, Andersson AF, Quince C. 2014. Binning metagenomic contigs by coverage and composition. Nat Methods 11: 1144–1146.

Andersson JO, Andersson SG. 1999. Insights into the evolutionary process of genome degradation. Curr Opin Genet Dev 9: 664–671.

Angelakis E, Mediannikov O, Jos S-L, Berenger J-M, Parola P, Raoult D. 2016. *Candidatus Coxiella massiliensis* infection. Emerg Infect Dis 22: 285–288.

Arivett BA, Fiester SE, Ohneck EJ, Penwell WF, Kaufman CM, Relich RF, Actis LA. 2015. Antimicrobial activity of gallium protoporphyrin IX against *Acinetobacter baumannii* strains displaying different antibiotic resistance phenotypes. Antimicrob Agents Chemother 59: 7657–7665.

Beare PA, Gilk SD, Larson CL, Hill J, Stead CM, Omsland A, Cockrell DC, Howe D, Voth DE, Heinzen RA. 2011. Dot/Icm type IVB secretion system requirements for *Coxiella burnetii* growth in human macrophages. mBio 2: e00175–00111.

Beare PA, Sandoz KM, Larson CL, Howe D, Kronmiller B, Heinzen RA. 2014. Essential role for the response regulator PmrA in *Coxiella burnetii* type 4B secretion and colonization of mammalian host cells. J Bacteriol 196: 1925–1940.

Ben-Yosef M, Rot A, Mahagna M, Kapri E, Behar A, Gottlieb Y. 2020. *Coxiella*-like Endosymbiont of *Rhipicephalus sanguineus* is required for physiological processes during ontogeny. Front Microbiol 11: 493.

Bennett GM, Moran NA. 2015. Heritable symbiosis: The advantages and perils of an evolutionary rabbit hole. Proc Natl Acad Sci U S A 112: 10169–10176.

Bernstein LR. 1998. Mechanisms of therapeutic activity for gallium. Pharmacol Rev 50: 665–682.

Binetruy F, Buysse M, Lejarre Q, Barosi R, Villa M, Rahola N, Paupy C, Ayala D, Duron O. 2020. Microbial community structure reveals instability of nutritional symbiosis during the evolutionary radiation of *Amblyomma* ticks. Mol Ecol 29: 1016–1029.

Bitew MA, Hofmann J, De Souza DP, Wawegama NK, Newton HJ, Sansom FM. 2020. SdrA, an NADP(H)-regenerating enzyme, is crucial for *Coxiella burnetii* to resist oxidative stress and replicate intracellularly. Cell Microbiol 22: e13154.

Bitew MA, Khoo CA, Neha N, De Souza DP, Tull D, Wawegama NK, Newton HJ, Sansom FM. 2018. De novo NAD synthesis is required for intracellular replication of *Coxiella burnetii*, the causative agent of the neglected zoonotic disease Q fever. J Biol Chem 293: 18636–18645.

Bolger AM, Lohse M, Usadel B. 2014. Trimmomatic: a flexible trimmer for Illumina sequence data. Bioinformatics 30: 2114–2120.

Bonchi C, Imperi F, Minandri F, Visca P, Frangipani E. 2014. Repurposing of gallium-based drugs for antibacterial therapy: Gallium-based antibacterials. BioFactors 40: 303–312.

Burstein D, Amaro F, Zusman T, Lifshitz Z, Cohen O, Gilbert JA, Pupko T, Shuman HA, Segal G. 2016. Genomic analysis of 38 *Legionella* species identifies large and diverse effector repertoires. Nat Genet 48: 167–175.

Bury-Moné S, Mendz GL, Ball GE, Thibonnier M, Stingl K, Ecobichon C, Avé P, Huerre M, Labigne A, Thiberge J-M, et al. 2008. Roles of alpha and beta carbonic anhydrases of *Helicobacter pylori* in the urease-dependent response to acidity and in colonization of the murine gastric mucosa. Infect Immun 76: 497–509.

Chan PP, Lowe TM. 2019. TRNAscan-SE: Searching for tRNA genes in genomic sequences. Methods Mol Biol 1962: 1–14.

Chen C, Banga S, Mertens K, Weber MM, Gorbaslieva I, Tan Y, Luo Z-Q, Samuel JE. 2010. Large-scale identification and translocation of type IV secretion substrates by *Coxiella burnetii*. Proc Natl Acad Sci U S A 107: 21755–21760.

Chong RA, Moran NA. 2018. Evolutionary loss and replacement of *Buchnera*, the obligate endosymbiont of aphids. ISME J 12: 898–908.

Clayton AL, Jackson DG, Weiss RB, Dale C. 2016. Adaptation by deletogenic replication slippage in a nascent symbiont. Mol Biol Evol 33: 1957–1966.

Clifford CM, Hoogstraal H, Radovsky FJ, Stiller D, Keirans JE. 1980. *Ornithodoros* (*Alectorobius*) *amblus* (Acarina: Ixodoidea: Argasidae): identity, marine bird and human hosts, virus infections, and distribution in Peru. J Parasitol 66: 312–323.

Cohen O, Ashkenazy H, Belinky F, Huchon D, Pupko T. 2010. GLOOME: gain loss mapping engine. Bioinformatics 26: 2914–2915.

Coleman J. 1990. Characterization of *Escherichia coli* cells deficient in 1-acyl-sn-glycerol-3-phosphate acyltransferase activity. J Biol Chem 265: 17215–17221.

Cox HR. 1938. A filter-passing infectious agent isolated from ticks: III. Description of organism and cultivation experiments. Public Health Rep 53: 2270–2276.

Darriba D, Taboada GL, Doallo R, Posada D. 2012. jModelTest 2: more models, new heuristics and parallel computing. Nat Methods 9: 772–772.

Duron O, Binetruy F, Noël V, Cremaschi J, McCoy KD, Arnathau C, Plantard O, Goolsby J, Pérez de León AA, Heylen DJA, et al. 2017. Evolutionary changes in symbiont community structure in ticks. Mol Ecol 26: 2905–2921.

Duron O, Doublet P, Vavre F, Bouchon D. 2018a. The importance of revisiting Legionellales diversity. Trends Parasitol 34: 1027–1037.

Duron O, Gottlieb Y. 2020. Convergence of nutritional symbioses in obligate blood feeders. Trends Parasitol 36: 816–825.

Duron O, Morel O, Noël V, Buysse M, Binetruy F, Lancelot R, Loire E, Ménard C, Bouchez O, Vavre F, et al. 2018b. Tick-bacteria mutualism depends on B vitamin synthesis pathway. Curr Biol 28: 1896–1902.e5.

Duron O, Noël V, McCoy KD, Bonazzi M, Sidi-Boumedine K, Morel O, Vavre F, Zenner L, Jourdain E, Durand P, et al. 2015a. The recent evolution of a maternally-inherited endosymbiont of ticks led to the emergence of the Q fever pathogen, *Coxiella burnetii*. PLoS Pathog 11: e1004892.

Duron O, Sidi-Boumedine K, Rousset E, Moutailler S, Jourdain E. 2015b. The importance of ticks in Q fever transmission: What has (and has not) been demonstrated? Trends Parasitol 31: 536–552.

Eldin C, Mélenotte C, Mediannikov O, Ghigo E, Million M, Edouard S, Mege J-L, Maurin M, Raoult D. 2017. From Q fever to *Coxiella burnetii* infection: A paradigm change. Clin Microbiol Rev 30: 115–190.

Elliman JR, Owens L. 2020. Confirmation that *candidatus Coxiella cheraxi* from redclaw crayfish (*Cherax quadricarinatus*) is a close relative of *Coxiella burnetii*, the agent of Q-fever. Lett Appl Microbiol 71: 320–326.

Emms DM, Kelly S. 2015. OrthoFinder: solving fundamental biases in whole genome comparisons dramatically improves orthogroup inference accuracy. Genome Biol 16: 157.

Endley S, McMurray D, Ficht TA. 2001. Interruption of the cydB locus in *Brucella abortus* attenuates intracellular survival and virulence in the mouse model of infection. J Bacteriol 183: 2454–2462.

Gerhart JG, Dutcher HA, Brenner AE, Moses AS, Grubhoffer L, Raghavan R. 2018. Multiple acquisitions of pathogen-derived *Francisella* endosymbionts in soft ticks. Genome Biol Evol 10: 607–615.

Gerhart JG, Moses AS, Raghavan R. 2016. A *Francisella*-like endosymbiont in the Gulf Coast tick evolved from a mammalian pathogen. Sci Rep 6: 33670.

Gilk SD. 2012. Role of lipids in *Coxiella burnetii* infection. Adv Exp Med Biol 984: 199–213.

Gilk SD, Beare PA, Heinzen RA. 2010. *Coxiella burnetii* expresses a functional ∆24 sterol reductase. J Bacteriol 192: 6154–6159.

Goss CH, Kaneko Y, Khuu L, Anderson GD, Ravishankar S, Aitken ML, Lechtzin N, Zhou G, Czyz DM, McLean K, et al. 2018. Gallium disrupts bacterial iron metabolism and has therapeutic effects in mice and humans with lung infections. Sci Transl Med 10: eaat7520.

Gottlieb Y, Lalzar I, Klasson L. 2015. Distinctive genome reduction rates revealed by genomic analyses of two *Coxiella*-like endosymbionts in ticks. Genome Biol Evol 7: 1779–1796.

Guimard T, Amrane S, Prudent E, El Karkouri K, Raoult D, Angelakis E. 2017. Case report: Scalp eschar and neck lymphadenopathy associated with bacteremia due to *Coxiella*-like bacteria. Am J Trop Med Hyg 97: 1319–1322.

Guizzo MG, Parizi LF, Nunes RD, Schama R, Albano RM, Tirloni L, Oldiges DP, Vieira RP, Oliveira WHC, Leite M de S, et al. 2017. A *Coxiella* mutualist symbiont is essential to the development of *Rhipicephalus microplus*. Sci Rep 7: 17554.

Hackstadt T. 1983. Estimation of the cytoplasmic pH of *Coxiella burnetii* and effect of substrate oxidation on proton motive force. J Bacteriol 154: 591–597.

Helmy N, Khalil GM, Hoogstraal H. 1983. Hyperparasitism in *Ornithodoros erraticus*. J Parasitol 69: 229–233.

Hijazi S, Visaggio D, Pirolo M, Frangipani E, Bernstein L, Visca P. 2018. Antimicrobial activity of gallium compounds on ESKAPE pathogens. Front Cell Infect Microbiol 8: 316.

Hijazi S, Visca P, Frangipani E. 2017. Gallium-protoporphyrin IX inhibits *Pseudomonas aeruginosa* growth by targeting cytochromes. Front Cell Infect Microbiol 7: 12.

Hill J, Samuel JE. 2011. *Coxiella burnetii* acid phosphatase inhibits the release of reactive oxygen intermediates in polymorphonuclear leukocytes. Infect Immun 79: 414–420.

Hosokawa T, Koga R, Kikuchi Y, Meng X-Y, Fukatsu T. 2010. *Wolbachia* as a bacteriocyte-associated nutritional mutualist. Proc Natl Acad Sci 107: 769–774.

Hosokawa T, Nikoh N, Koga R, Satô M, Tanahashi M, Meng X-Y, Fukatsu T. 2012. Reductive genome evolution, host–symbiont co-speciation and uterine transmission of endosymbiotic bacteria in bat flies. ISME J 6: 577–587.

Husnik F, Hypsa V, Darby A. 2020. Insect-symbiont gene expression in the midgut bacteriocytes of a blood-sucking parasite. Genome Biol Evol 12: 429–442.

Hyatt D, Chen G-L, LoCascio PF, Land ML, Larimer FW, Hauser LJ. 2010. Prodigal: prokaryotic gene recognition and translation initiation site identification. BMC Bioinformatics 11: 119.

Iwata-Reuyl D. 2003. Biosynthesis of the 7-deazaguanosine hypermodified nucleosides of transfer RNA. Bioorganic Chem 31: 24–43.

Jansen R, Bussemaker HJ, Gerstein M. 2003. Revisiting the codon adaptation index from a whole-genome perspective: analyzing the relationship between gene expression and codon occurrence in yeast using a variety of models. Nucleic Acids Res 31: 2242–2251.

Katoh K, Standley DM. 2013. MAFFT multiple sequence alignment software version 7: improvements in performance and usability. Mol Biol Evol 30: 772–780.

Klyachko O, Stein BD, Grindle N, Clay K, Fuqua C. 2007. Localization and visualization of a *Coxiella*-type symbiont within the lone star tick, *Amblyomma americanum*. Appl Environ Microbiol 73: 6584–6594.

Koh CS, Sarin LP. 2018. Transfer RNA modification and infection – Implications for pathogenicity and host responses. Biochim Biophys Acta Gene Regul Mech 1861: 419–432.

Körner S, Makert GR, Mertens-Scholz K, Henning K, Pfeffer M, Starke A, Nijhof AM, Ulbert S. 2020. Uptake and fecal excretion of *Coxiella burnetii* by *Ixodes ricinus* and *Dermacentor marginatus* ticks. Parasit Vectors 13: 75.

Kozlowski LP. 2016. IPC – Isoelectric Point Calculator. Biol Direct 11: 55.

Kuba M, Neha N, Newton P, Lee YW, Bennett-Wood V, Hachani A, De Souza DP, Nijagal B, Dayalan S, Tull D, et al. 2020. EirA is a novel protein essential for intracellular replication of *Coxiella burnetii*. Infect Immun 88.

Labruna MB, Ahid SMM, Soares HS, Suassuna ACD. 2007. Hyperparasitism in *Amblyomma rotundatum* (Acari: Ixodidae). J Parasitol 93: 1531–1532.

Lagesen K, Hallin P, Rødland EA, Stærfeldt H-H, Rognes T, Ussery DW. 2007. RNAmmer: consistent and rapid annotation of ribosomal RNA genes. Nucleic Acids Res 35: 3100–3108.

Lalzar I, Friedmann Y, Gottlieb Y. 2014. Tissue tropism and vertical transmission of *Coxiella* in *Rhipicephalus sanguineus* and *Rhipicephalus turanicus* ticks: Tissue tropism of tick symbionts. Environ Microbiol 16: 3657–3668.

Lambert JD, Moran NA. 1998. Deleterious mutations destabilize ribosomal RNA in endosymbiotic bacteria. Proc Natl Acad Sci U S A 95: 4458–4462.

Lawrence JG, Ochman H. 1997. Amelioration of bacterial genomes: rates of change and exchange. J Mol Evol 44: 383–397.

Lerat E, Daubin V, Ochman H, Moran NA. 2005. Evolutionary origins of genomic repertoires in bacteria. PLoS Biol 3: e130.

Li L-H, Zhang Y, Zhu D. 2018. Effects of antibiotic treatment on the fecundity of *Rhipicephalus haemaphysaloides* ticks. Parasit Vectors 11.

Llanos-Soto S, Muñoz-Leal S, González-Acuña D. 2019. Hyperparasitism in the seabird tick *Ornithodoros amblus* (Acari: Argasidae). Syst Appl Acarol 24: 525.

Machado-Ferreira E, Dietrich G, Hojgaard A, Levin M, Piesman J, Zeidner NS, Soares CAG. 2011. *Coxiella* symbionts in the Cayenne tick *Amblyomma cajennense*. Microb Ecol 62: 134–142.

Martinez E, Cantet F, Fava L, Norville I, Bonazzi M. 2014. Identification of OmpA, a *Coxiella burnetii* protein involved in host cell invasion, by multi-phenotypic high-content screening. PLoS Pathog 10: e1004013.

Martinez E, Huc-Brandt S, Brelle S, Allombert J, Cantet F, Gannoun-Zaki L, Burette M, Martin M, Letourneur F, Bonazzi M, et al. 2020. The secreted protein kinase CstK from *Coxiella burnetii* influences vacuole development and interacts with the GTPase-activating host protein TBC1D5. J Biol Chem 295: 7391–7403.

Maturana P, Graham JG, Sharma UM, Voth DE. 2013. Refining the plasmid-encoded type IV secretion system substrate repertoire of Coxiella burnetii. J Bacteriol 195: 3269–3276.

Maurin M, Raoult D. 1999. Q Fever. Clin Microbiol Rev 12: 518–553.

McCutcheon JP, Moran NA. 2011. Extreme genome reduction in symbiotic bacteria. Nat Rev Microbiol 10: 13–26.

McCutcheon JP, Moran NA. 2010. Functional convergence in reduced genomes of bacterial symbionts spanning 200 My of evolution. Genome Biol Evol 2: 708–718.

Mertens K, Samuel JE. 2012. Defense mechanisms against oxidative stress in *Coxiella burnetii*: adaptation to a unique intracellular niche. Adv Exp Med Biol 984: 39–63.

Moran NA. 1996. Accelerated evolution and Muller’s rachet in endosymbiotic bacteria. Proc Natl Acad Sci U S A 93: 2873–2878.

Moran NA. 2002. Microbial minimalism: genome reduction in bacterial pathogens. Cell 108: 583–586.

Moran NA, Munson MA, Baumann P, Ishikawa H. 1993. A molecular clock in endosymbiotic bacteria is calibrated using the insect hosts. Proc Biol Sci 253: 167–171.

Moses AS, Millar JA, Bonazzi M, Beare PA, Raghavan R. 2017. Horizontally acquired biosynthesis genes boost *Coxiella burnetii*’s physiology. Front Cell Infect Microbiol 7: 174.

Nakabachi A, Yamashita A, Toh H, Ishikawa H, Dunbar HE, Moran NA, Hattori M. 2006. The 160-kilobase genome of the bacterial endosymbiont *Carsonella*. Science 314: 267.

Needle DB, Agnew DW, Bradway DS, Nordhausen RW, Garner MM. 2020. Avian coxiellosis in nine psittacine birds, one black-browed barbet, and one paradise tanager. Avian Pathol J WVPA 49: 268–274.

Newton HJ, Kohler LJ, McDonough JA, Temoche-Diaz M, Crabill E, Hartland EL, Roy CR. 2014. A screen of *Coxiella burnetii* mutants reveals important roles for Dot/Icm effectors and host autophagy in vacuole biogenesis. PLOS Pathog 10: e1004286.

Neyrolles O, Wolschendorf F, Mitra A, Niederweis M. 2015. *Mycobacteria*, metals, and the macrophage. Immunol Rev 264: 249–263.

Nurk S, Meleshko D, Korobeynikov A, Pevzner PA. 2017. metaSPAdes: a new versatile metagenomic assembler. Genome Res 27: 824–834.

Ochman H, Lawrence JG, Groisman EA. 2000. Lateral gene transfer and the nature of bacterial innovation. Nature 405: 299–304.

Omsland A, Heinzen RA. 2011. Life on the outside: the rescue of *Coxiella burnetii* from its host cell. Annu Rev Microbiol 65: 111–128.

Peer A, Margalit H. 2014. Evolutionary patterns of *Escherichia coli* small RNAs and their regulatory interactions. RNA 20: 994–1003.

Potter SC, Luciani A, Eddy SR, Park Y, Lopez R, Finn RD. 2018. HMMER web server: 2018 update. Nucleic Acids Res 46: W200–W204.

Perotti MA, Allen JM, Reed DL, Braig HR. 2007. Host-symbiont interactions of the primary endosymbiont of human head and body lice. FASEB J 21: 1058–1066.

Raghavan R, Kelkar YD, Ochman H. 2012. A selective force favoring increased G+C content in bacterial genes. Proc Natl Acad Sci U S A 109: 14504–14507.

Rio RVM, Symula RE, Wang J, Lohs C, Wu Y-N, Snyder AK, Bjornson RD, Oshima K, Biehl BS, Perna NT, et al. 2012. Insight into the transmission biology and species-specific functional capabilities of tsetse (Diptera: glossinidae) obligate symbiont *Wigglesworthia*. MBio 3: 00240–11.

Ronquist F, Teslenko M, van der Mark P, Ayres DL, Darling A, Höhna S, Larget B, Liu L, Suchard MA, Huelsenbeck JP. 2012. MrBayes 3.2: efficient Bayesian phylogenetic inference and model choice across a large model space. Syst Biol 61: 539–542.

Rowland JL, Niederweis M. 2012. Resistance mechanisms of *Mycobacterium tuberculosis* against phagosomal copper overload. Tuberc Edinb Scotl 92: 202–210.

Rowlett VW, Mallampalli VKPS, Karlstaedt A, Dowhan W, Taegtmeyer H, Margolin W, Vitrac H. 2017. Impact of membrane phospholipid alterations in *Escherichia coli* on cellular function and bacterial stress adaptation. J Bacteriol 199: e00849–16.

Russell JA, Oliver KM, Hansen AK. 2017. Band-aids for *Buchnera* and B vitamins for all. Mol Ecol 26: 2199–2203.

Saiki K, Mogi T, Anraku Y. 1992. Heme O biosynthesis in *Escherichia coli*: the cyoE gene in the cytochrome bo operon encodes a protoheme IX farnesyltransferase. Biochem Biophys Res Commun 189: 1491–1497.

Samanta D, Clemente TM, Schuler BE, Gilk SD. 2019. *Coxiella burnetii* type 4B secretion system-dependent manipulation of endolysosomal maturation is required for bacterial growth. PLoS Pathog 15: e1007855.

Sanchez SE, Omsland A. 2020. Critical role for molecular iron in *Coxiella burnetii* replication and viability. mSphere 5: e00458–20.

Sandoz KM, Popham DL, Beare PA, Sturdevant DE, Hansen B, Nair V, Heinzen RA. 2016. Transcriptional profiling of *Coxiella burnetii* reveals extensive cell wall remodeling in the small cell variant developmental form. PloS One 11: e0149957.

Segal G, Russo JJ, Shuman HA. 1999. Relationships between a new type IV secretion system and the icm/dot virulence system of *Legionella pneumophila*. Mol Microbiol 34: 799–809.

Seo M-G, Lee S-H, VanBik D, Ouh I-O, Yun S-H, Choi E, Park Y-S, Lee S-E, Kim JW, Cho G-J, et al. 2016. Detection and genotyping of *Coxiella burnetii* and *Coxiella*-like bacteria in horses in South Korea. PloS One 11: e0156710.

Seshadri R, Paulsen IT, Eisen JA, Read TD, Nelson KE, Nelson WC, Ward NL, Tettelin H, Davidsen TM, Beanan MJ, et al. 2003. Complete genome sequence of the Q-fever pathogen *Coxiella burnetii*. Proc Natl Acad Sci U S A 100: 5455–5460.

Sharp PM, Li WH. 1987. The codon adaptation index--a measure of directional synonymous codon usage bias, and its potential applications. Nucleic Acids Res 15: 1281–1295.

Shivaprasad HL, Cadenas MB, Diab SS, Nordhausen R, Bradway D, Crespo R, Breitschwerdt EB. 2008. *Coxiella*-like infection in psittacines and a toucan. Avian Dis 52: 426–432.

Simonet P, Gaget K, Balmand S, Ribeiro Lopes M, Parisot N, Buhler K, Duport G, Vulsteke V, Febvay G, Heddi A, et al. 2018. Bacteriocyte cell death in the pea aphid/*Buchnera* symbiotic system. Proc Natl Acad Sci 115: E1819–E1828.

Smith DJW, Derrick EH. 1940. Studies in the epidemiology of Q fever: 1. The isolation of six strains of *Rickettsia burneti* from the tick *Haemaphysalis humerosa*. Aust J Exp Biol Med Sci 18: 1–8.

Smith TA, Driscoll T, Gillespie JJ, Raghavan R. 2015. A *Coxiella*-like endosymbiont is a potential vitamin source for the Lone Star tick. Genome Biol Evol 7: 831–838.

Stamatakis A. 2014. RAxML version 8: a tool for phylogenetic analysis and post-analysis of large phylogenies. Bioinformatics 30: 1312–1313.

Stead CM, Cockrell DC, Beare PA, Miller HE, Heinzen RA. 2018. A *Coxiella burnetii* phospholipase A homolog pldA is required for optimal growth in macrophages and developmental form lipid remodeling. BMC Microbiol 18: 33.

Stojiljkovic I, Kumar V, Srinivasan N. 1999. Non-iron metalloporphyrins: potent antibacterial compounds that exploit haem/Hb uptake systems of pathogenic bacteria. Mol Microbiol 31: 429–442.

Talavera G, Castresana J. 2007. Improvement of phylogenies after removing divergent and ambiguously aligned blocks from protein sequence alignments. Syst Biol 56: 564–577.

Tsementzi D, Castro Gordillo J, Mahagna M, Gottlieb Y, Konstantinidis KT. 2018. Comparison of closely related, uncultivated *Coxiella* tick endosymbiont population genomes reveals clues about the mechanisms of symbiosis. Environ Microbiol 20: 1751–1764.

Vallejo Esquerra E, Yang H, Sanchez SE, Omsland A. 2017. Physicochemical and nutritional requirements for axenic replication suggest physiological basis for *Coxiella burnetii* niche restriction. Front Cell Infect Microbiol 7: 190.

Wachter S, Bonazzi M, Shifflett K, Moses AS, Raghavan R, Minnick MF. 2019. A CsrA-binding, *trans*-acting sRNA of *Coxiella burnetii* is necessary for optimal intracellular growth and vacuole formation during early infection of host cells. J Bacteriol 201: e00524–19.

Warrier I, Hicks LD, Battisti JM, Raghavan R, Minnick MF. 2014. Identification of novel small RNAs and characterization of the 6S RNA of *Coxiella burnetii*. PLoS One 9: e100147.

Wernegreen JJ, Ochman H, Jones IB, Moran NA. 2000. Decoupling of genome size and sequence divergence in a symbiotic bacterium. J Bacteriol 182: 3867–3869.

Woc-Colburn AM, Garner MM, Bradway D, West G, D’Agostino J, Trupkiewicz J, Barr B, Anderson SE, Rurangirwa FR, Nordhausen RW. 2008. Fatal coxiellosis in Swainson’s Blue Mountain Rainbow Lorikeets (*Trichoglossus haematodus moluccanus*). Vet Pathol 45: 247–254.

Zhang C-M, Li N-X, Zhang T-T, Qiu Z-X, Li Y, Li L-W, Liu J-Z. 2017. Endosymbiont CLS-HI plays a role in reproduction and development of *Haemaphysalis longicornis*. Exp Appl Acarol 73: 429–438.

Zhong J, Jasinskas A, Barbour AG. 2007. Antibiotic treatment of the tick vector *Amblyomma americanum* reduced reproductive fitness. PLoS One 2: e405.

