## Supplemental Figure 1 for "Ticks convert pathogenic *Coxiella* into endosymbionts"

Acidobacteria

Aquificae

Deferribacteres

Elusimicrobia

FCB Group

Fusobacteria

Nitrospinae Tectomicrobia Group

Nitrospirae

PVC Group

Spirochaetes

Synergistetes

Terrabacteria Group

Thermodesulfobacteria

Thermotogae

Proteobacteria: Acidithiobacillia

Proteobacteria: Alphaproteobacteria

Proteobacteria: Betaproteobacteria

Proteobacteria: Candidatus Muproteobacteria

Proteobacteria: Delta Epsilon Subdivisions

Proteobacteria: Gammaproteobacteria

Proteobacteria: Hydrogenophilalia

Proteobacteria: Oligoflexia

Proteobacteria: Zetaproteobacteria

Proteobacteria: Unclassified & Environmental Samples

Bacteria: Unclassified & Environmental Samples

Viruses & Constructs

Eukaryota

Archaea

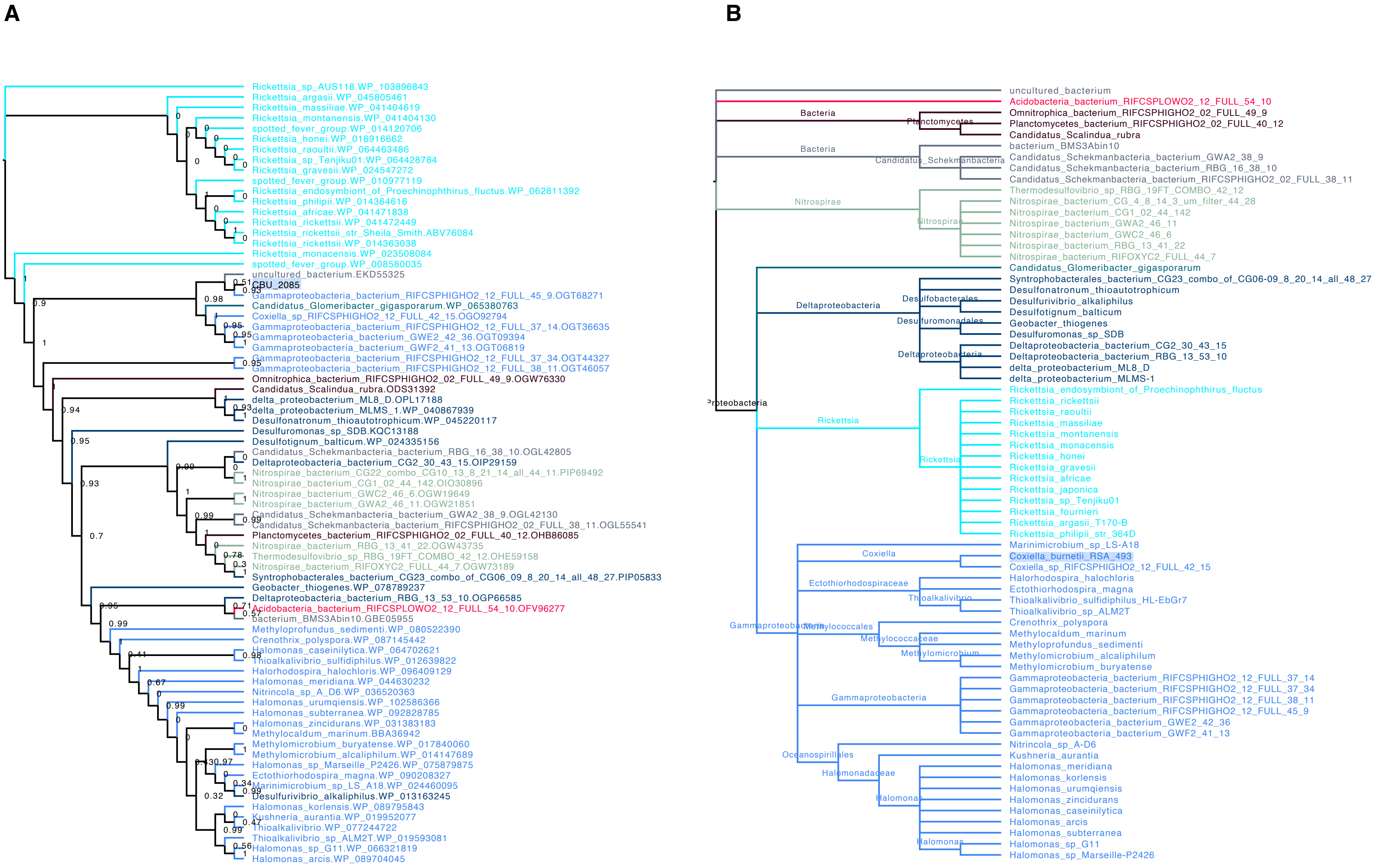
A
Rickettsia_sp_AUS118.WP_103896843
Rickettsia_argasii.WP_045805461
Rickettsia_massiliae.WP_041404619
Rickettsia_montanensis.WP_041404130
spotted_fever_group.WP_014120706
Rickettsia_honei.WP_016916662
Rickettsia_raoultii.WP_064463486
Rickettsia_sp_Tenjiku01.WP_064428784
Rickettsia_gravesii.WP_024547272
spotted_fever_group.WP_010977119
Rickettsia_endosymbiont_of_Proechinophthirus_fluctus.WP_062811392
Rickettsia_philipii.WP_014364616
Rickettsia_africae.WP_041471838
Rickettsia_rickettsii.WP_041472449
Rickettsia_rickettsii_str_Sheila_Smith.ABV76084
Rickettsia_rickettsii.WP_014363038
Rickettsia_monacensis.WP_023508084
spotted_fever_group.WP_008580035
uncultured_bacterium.EKD55325
CBU_2085
Gammaproteobacteria_bacterium_RIFCSPHIGHO2_12_FULL_45_9.OGT68271
Candidatus_Glomeribacter_gigasporarum.WP_065380763
Coxiella_sp_RIFCSPHIGHO2_12_FULL_42_15.OGO92794
Gammaproteobacteria_bacterium_RIFCSPHIGHO2_12_FULL_37_14.OGT36635
Gammaproteobacteria_bacterium_GWE2_42_36.OGT09394
Gammaproteobacteria_bacterium_GWF2_41_13.OGT06819
Gammaproteobacteria_bacterium_RIFCSPHIGHO2_12_FULL_37_34.OGT44327
Gammaproteobacteria_bacterium_RIFCSPHIGHO2_12_FULL_38_11.OGT46057
Omnitrophica_bacterium_RIFCSPHIGHO2_02_FULL_49_9.OGW76330
Candidatus_Scalindua_rubra.ODS31392
delta_proteobacterium_ML8_D.OPL17188
delta_proteobacterium_MLMS_1.WP_040867939
Desulfonatronum_thioautotrophicum.WP_045220117
Desulfuromonas_sp_SDB.KQC13188
Desulfotignum_balticum.WP_024335156
Candidatus_Schekmanbacteria_bacterium_RBG_16_38_10.OGL42805
Deltaproteobacteria_bacterium_CG2_30_43_15.OIP29159
Nitrospirae_bacterium_CG22_combo_CG10_13_8_21_14_all_44_11.PIP69492
Nitrospirae_bacterium_CG1_02_44_142.OIO30896
Nitrospirae_bacterium_GWC2_46_6.OGW19649
Nitrospirae_bacterium_GWA2_46_11.OGW21851
Candidatus_Schekmanbacteria_bacterium_GWA2_38_9.OGL42130
Candidatus_Schekmanbacteria_bacterium_RIFCSPHIGHO2_02_FULL_38_11.OGL55541
Planctomycetes_bacterium_RIFCSPHIGHO2_02_FULL_40_12.OHB86085
Nitrospirae_bacterium_RBG_13_41_22.OGW43735
Thermodesulfovibrio_sp_RBG_19FT_COMBO_42_12.OHE59158
Nitrospirae_bacterium_RIFOXYC2_FULL_44_7.OGW73189
Syntrophobacterales_bacterium_CG23_combo_of_CG06_09_8_20_14_all_48_27.PIP05833
Geobacter_thiogenes.WP_078789237
Deltaproteobacteria_bacterium_RBG_13_53_10.OGP66585
Acidobacteria_bacterium_RIFCSPLOWO2_12_FULL_54_10.OFV96277
bacterium_BMS3Abin10.GBE05955
Methyloprofundus_sedimenti.WP_080522390
Crenothrix_polyspora.WP_087145442
Halomonas_caseinilytica.WP_064702621
Thioalkalivibrio_sulfidiphilus.WP_012639822
Halorhodospira_halochloris.WP_096409129
Halomonas_meridiana.WP_044630232
Nitrincola_sp_A_D6.WP_036520363
Halomonas_urumqiensis.WP_102586366
Halomonas_subterranea.WP_092828785
Halomonas_zincidurans.WP_031383183
Methylocaldum_marinum.BBA36942
Methylomicrobium_buryatense.WP_017840060
Methylomicrobium_alcaliphilum.WP_014147689
Halomonas_sp_Marseille_P2426.WP_075879875
Ectothiorhodospira_magna.WP_090208327
Marinimicrobium_sp_LS_A18.WP_024460095
Desulfurivibrio_alkaliphilus.WP_013163245
Halomonas_korlensis.WP_089795843
Kushneria_aurantia.WP_019952077
Thioalkalivibrio.WP_077244722
Thioalkalivibrio_sp_ALM2T.WP_019593081
Halomonas_sp_G11.WP_066321819
Halomonas_arcis.WP_089704045
B
uncultured_bacterium
Acidobacteria_bacterium_RIFCSPLOWO2_12_FULL_54_10
Bacteria
Planctomycetes
Omnitrophica_bacterium_RIFCSPHIGHO2_02_FULL_49_9
Planctomycetes_bacterium_RIFCSPHIGHO2_02_FULL_40_12
Candidatus_Scalindua_rubra
bacterium_BMS3Abin10
Bacteria
Candidatus_Schekmanbacteria
Candidatus_Schekmanbacteria_bacterium_GWA2_38_9
Candidatus_Schekmanbacteria_bacterium_RBG_16_38_10
Candidatus_Schekmanbacteria_bacterium_RIFCSPHIGHO2_02_FULL_38_11
Nitrospirae
Thermodesulfovibrio_sp_RBG_19FT_COMBO_42_12
Nitrospirae_bacterium_CG_4_8_14_3_um_filter_44_28
Nitrospirae
Nitrospirae_bacterium_CG1_02_44_142
Nitrospirae_bacterium_GWA2_46_11
Nitrospirae_bacterium_GWC2_46_6
Nitrospirae_bacterium_RBG_13_41_22
Nitrospirae_bacterium_RIFOXYC2_FULL_44_7
Candidatus_Glomeribacter_gigasporarum
Syntrophobacterales_bacterium_CG23_combo_of_CG06-09_8_20_14_all_48_27
Desulfonatronum_thioautotrophicum
Deltaproteobacteria
Desulfobacterales
Desulfurivibrio_alkaliphilus
Desulfotignum_balticum
Desulfuromonadales
Geobacter_thiogenes
Desulfuromonas_sp_SDB
Deltaproteobacteria
Deltaproteobacteria_bacterium_CG2_30_43_15
Deltaproteobacteria_bacterium_RBG_13_53_10
delta_proteobacterium_ML8_D
delta_proteobacterium_MLMS-1
Proteobacteria
Rickettsia_endosymbiont_of_Proechinophthirus_fluctus
Rickettsia_rickettsii
Rickettsia_raoultii
Rickettsia
Rickettsia_massiliae
Rickettsia_montanensis
Rickettsia_monacensis
Rickettsia_honei
Rickettsia
Rickettsia_gravesii
Rickettsia_africae
Rickettsia_japonica
Rickettsia_sp_Tenjiku01
Rickettsia_fournieri
Rickettsia_argasii_T170-B
Rickettsia_philipii_str_364D
Marinimicrobium_sp_LS-A18
Coxiella
Coxiella_burnetii_RSA_493
Coxiella_sp_RIFCSPHIGHO2_12_FULL_42_15
Ectothiorhodospiraceae
Halorhodospira_halochloris
Ectothiorhodospira_magna
Thioalkalivibrio
Thioalkalivibrio_sulfidiphilus_HL-EbGr7
Thioalkalivibrio_sp_ALM2T
Gammaproteobacteria
Methylococcales
Crenothrix_polyspora
Methylocaldum_marinum
Methylococcaceae
Methyloprofundus_sedimenti
Methylomicrobium
Methylomicrobium_alcaliphilum
Methylomicrobium_buryatense
Gammaproteobacteria
Gammaproteobacteria_bacterium_RIFCSPHIGHO2_12_FULL_37_14
Gammaproteobacteria_bacterium_RIFCSPHIGHO2_12_FULL_37_34
Gammaproteobacteria_bacterium_RIFCSPHIGHO2_12_FULL_38_11
Gammaproteobacteria_bacterium_RIFCSPHIGHO2_12_FULL_45_9
Gammaproteobacteria_bacterium_GWE2_42_36
Gammaproteobacteria_bacterium_GWF2_41_13
Oceanospirillales
Nitrincola_sp_A-D6
Kushneria_aurantia
Halomonadaceae
Halomonas_meridiana
Halomonas_korlensis
Halomonas_urumqiensis
Halomonas
Halomonas_zincidurans
Halomonas_caseinilytica
Halomonas_arcis
Halomonas_subterranea
Halomonas
Halomonas_sp_G11
Halomonas_sp_Marseille-P2426

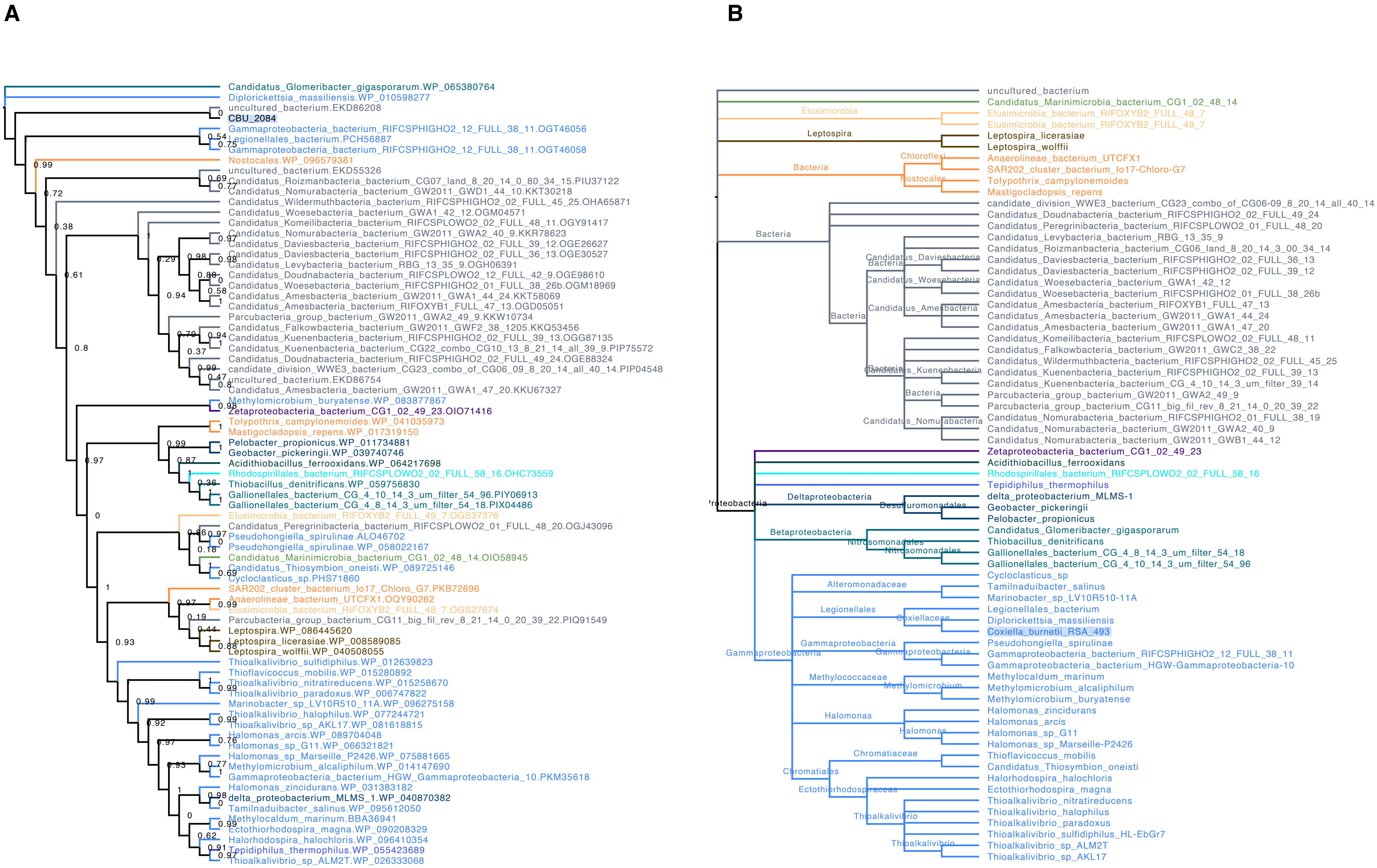
A
Candidatus_Glomeribacter_gigasporarum.WP_065380764
Diplorickettsia_massiliensis.WP_010598277
uncultured_bacterium.EKD86208
CBU_2084
Gammaproteobacteria_bacterium_RIFCSPHIGHO2_12_FULL_38_11.OGT46056
Legionellales_bacterium.PCH56887
Gammaproteobacteria_bacterium_RIFCSPHIGHO2_12_FULL_38_11.OGT46058
Nostocales.WP_096579381
uncultured_bacterium.EKD55326
Candidatus_Roizmanbacteria_bacterium_CG07_land_8_20_14_0_80_34_15.PIU37122
Candidatus_Nomurabacteria_bacterium_GW2011_GWD1_44_10.KKT30218
Candidatus_Wildermuthbacteria_bacterium_RIFCSPHIGHO2_02_FULL_45_25.OHA65871
Candidatus_Woesebacteria_bacterium_GWA1_42_12.OGM04571
Candidatus_Komeilibacteria_bacterium_RIFCSPLOWO2_02_FULL_48_11.OGY91417
Candidatus_Nomurabacteria_bacterium_GW2011_GWA2_40_9.KKR78623
Candidatus_Daviesbacteria_bacterium_RIFCSPHIGHO2_02_FULL_39_12.OGE26627
Candidatus_Daviesbacteria_bacterium_RIFCSPHIGHO2_02_FULL_36_13.OGE30527
Candidatus_Levybacteria_bacterium_RBG_13_35_9.OGH06391
Candidatus_Doudnabacteria_bacterium_RIFCSPLOWO2_12_FULL_42_9.OGE98610
Candidatus_Woesebacteria_bacterium_RIFCSPHIGHO2_01_FULL_38_26b.OGM18969
Candidatus_Amesbacteria_bacterium_GW2011_GWA1_44_24.KKT58069
Candidatus_Amesbacteria_bacterium_RIFOXYB1_FULL_47_13.OGD05051
Parcubacteria_group_bacterium_GW2011_GWA2_49_9.KKW10734
Candidatus_Falkowbacteria_bacterium_GW2011_GWF2_38_1205.KKQ53456
Candidatus_Kuenenbacteria_bacterium_RIFCSPHIGHO2_02_FULL_39_13.OGG87135
Candidatus_Kuenenbacteria_bacterium_CG22_combo_CG10_13_8_21_14_all_39_9.PIP75572
Candidatus_Doudnabacteria_bacterium_RIFCSPHIGHO2_02_FULL_49_24.OGE88324
candidate_division_WWE3_bacterium_CG23_combo_of_CG06_09_8_20_14_all_40_14.PIP04548
uncultured_bacterium.EKD86754
Candidatus_Amesbacteria_bacterium_GW2011_GWA1_47_20.KKU67327
Methylomicrobium_buryatense.WP_083877867
Zetaproteobacteria_bacterium_CG1_02_49_23.OIO71416
Tolypothrix_campylonemoides.WP_041035973
Mastigocladopsis_repens.WP_017319150
Pelobacter_propionicus.WP_011734881
Geobacter_pickeringii.WP_039740746
Acidithiobacillus_ferrooxidans.WP_064217698
Rhodospirillales_bacterium_RIFCSPLOWO2_02_FULL_58_16.OHC73559
Thiobacillus_denitrificans.WP_059756830
Gallionellales_bacterium_CG_4_10_14_3_um_filter_54_96.PIY06913
Gallionellales_bacterium_CG_4_8_14_3_um_filter_54_18.PIX04486
Elusimicrobia_bacterium_RIFOXYB2_FULL_49_7.OGS37376
Candidatus_Peregrinibacteria_bacterium_RIFCSPLOWO2_01_FULL_48_20.OGJ43096
Pseudohongiella_spirulinae.ALO46702
Pseudohongiella_spirulinae.WP_058022167
Candidatus_Marinimicrobia_bacterium_CG1_02_48_14.OIO58945
Candidatus_Thiosymbion_oneisti.WP_089725146
Cycloclasticus_sp.PHS71860
SAR202_cluster_bacterium_Io17_Chloro_G7.PKB72696
Anaerolineae_bacterium_UTCFX1.OQY90262
Elusimicrobia_bacterium_RIFOXYB2_FULL_48_7.OGS27674
Parcubacteria_group_bacterium_CG11_big_fil_rev_8_21_14_0_20_39_22.PIQ91549
Leptospira.WP_086445620
Leptospira_licerasiae.WP_008589085
Leptospira_wolffii.WP_040508055
Thioalkalivibrio_sulfidiphilus.WP_012639823
Thioflavicoccus_mobilis.WP_015280892
Thioalkalivibrio_nitratireducens.WP_015258670
Thioalkalivibrio_paradoxus.WP_006747822
Marinobacter_sp_LV10R510_11A.WP_096275158
Thioalkalivibrio_halophilus.WP_077244721
Thioalkalivibrio_sp_AKL17.WP_081618815
Halomonas_arcis.WP_089704048
Halomonas_sp_G11.WP_066321821
Halomonas_sp_Marseille_P2426.WP_075881665
Methylomicrobium_alcaliphilum.WP_014147690
Gammaproteobacteria_bacterium_HGW_Gammaproteobacteria_10.PKM35618
Halomonas_zincidurans.WP_031383182
delta_proteobacterium_MLMS_1.WP_040870382
Tamilnaduibacter_salinus.WP_095612050
Methylocaldum_marinum.BBA36941
Ectothiorhodospira_magna.WP_090208329
Halorhodospira_halochloris.WP_096410354
Tepidiphilus_thermophilus.WP_055423689
Thioalkalivibrio_sp_ALM2T.WP_026333068
B
uncultured_bacterium
Candidatus_Marinimicrobia_bacterium_CG1_02_48_14
Elusimicrobia
Elusimicrobia_bacterium_RIFOXYB2_FULL_48_7
Elusimicrobia_bacterium_RIFOXYB2_FULL_49_7
Leptospira
Leptospira_licerasiae
Leptospira_wolffii
Chloroflexi
Bacteria
Anaerolineae_bacterium_UTCFX1
SAR202_cluster_bacterium_Io17-Chloro-G7
Nostocales
Tolypothrix_campylonemoides
Mastigocladopsis_repens
candidate_division_WWE3_bacterium_CG23_combo_of_CG06-09_8_20_14_all_40_14
Candidatus_Doudnabacteria_bacterium_RIFCSPHIGHO2_02_FULL_49_24
Candidatus_Peregrinibacteria_bacterium_RIFCSPLOWO2_01_FULL_48_20
Candidatus_Levybacteria_bacterium_RBG_13_35_9
Candidatus_Roizmanbacteria_bacterium_CG06_land_8_20_14_3_00_34_14
Candidatus_Daviesbacteria
Candidatus_Daviesbacteria_bacterium_RIFCSPHIGHO2_02_FULL_36_13
Candidatus_Daviesbacteria_bacterium_RIFCSPHIGHO2_02_FULL_39_12
Candidatus_Woesebacteria
Candidatus_Woesebacteria_bacterium_GWA1_42_12
Candidatus_Woesebacteria_bacterium_RIFCSPHIGHO2_01_FULL_38_26b
Candidatus_Amesbacteria
Candidatus_Amesbacteria_bacterium_RIFOXYB1_FULL_47_13
Candidatus_Amesbacteria_bacterium_GW2011_GWA1_44_24
Candidatus_Amesbacteria_bacterium_GW2011_GWA1_47_20
Candidatus_Komeilibacteria_bacterium_RIFCSPLOWO2_02_FULL_48_11
Candidatus_Falkowbacteria_bacterium_GW2011_GWC2_38_22
Candidatus_Wildermuthbacteria_bacterium_RIFCSPHIGHO2_02_FULL_45_25
Candidatus_Kuenenbacteria
Candidatus_Kuenenbacteria_bacterium_RIFCSPHIGHO2_02_FULL_39_13
Candidatus_Kuenenbacteria_bacterium_CG_4_10_14_3_um_filter_39_14
Parcubacteria_group_bacterium_GW2011_GWA2_49_9
Parcubacteria_group_bacterium_CG11_big_fil_rev_8_21_14_0_20_39_22
Candidatus_Nomurabacteria
Candidatus_Nomurabacteria_bacterium_RIFCSPHIGHO2_01_FULL_38_19
Candidatus_Nomurabacteria_bacterium_GW2011_GWA2_40_9
Candidatus_Nomurabacteria_bacterium_GW2011_GWB1_44_12
Proteobacteria
Zetaproteobacteria_bacterium_CG1_02_49_23
Acidithiobacillus_ferrooxidans
Rhodospirillales_bacterium_RIFCSPLOWO2_02_FULL_58_16
Tepidiphilus_thermophilus
Deltaproteobacteria
Desulfuromonadales
delta_proteobacterium_MLMS-1
Geobacter_pickeringii
Pelobacter_propionicus
Betaproteobacteria
Candidatus_Glomeribacter_gigasporarum
Thiobacillus_denitrificans
Nitrosomonadales
Gallionellales_bacterium_CG_4_8_14_3_um_filter_54_18
Gallionellales_bacterium_CG_4_10_14_3_um_filter_54_96
Gammaproteobacteria
Cycloclasticus_sp
Alteromonadaceae
Tamilnaduibacter_salinus
Marinobacter_sp_LV10R510-11A
Legionellales
Coxiellaceae
Legionellales_bacterium
Diplorickettsia_massiliensis
Coxiella_burnetii_RSA_493
Pseudohongiella_spirulinae
Gammaproteobacteria_bacterium_RIFCSPHIGHO2_12_FULL_38_11
Gammaproteobacteria_bacterium_HGW-Gammaproteobacteria-10
Methylococcaceae
Methylomicrobium
Methylocaldum_marinum
Methylomicrobium_alcaliphilum
Methylomicrobium_buryatense
Halomonas
Halomonas_zincidurans
Halomonas_arcis
Halomonas_sp_G11
Halomonas_sp_Marseille-P2426
Chromatiales
Chromatiaceae
Thioflavicoccus_mobilis
Candidatus_Thiosymbion_oneisti
Ectothiorhodospiraceae
Halorhodospira_halochloris
Ectothiorhodospira_magna
Thioalkalivibrio
Thioalkalivibrio_nitratireducens
Thioalkalivibrio_halophilus
Thioalkalivibrio_paradoxus
Thioalkalivibrio_sulfidiphilus_HL-EbGr7
Thioalkalivibrio_sp_ALM2T
Thioalkalivibrio_sp_AKL17

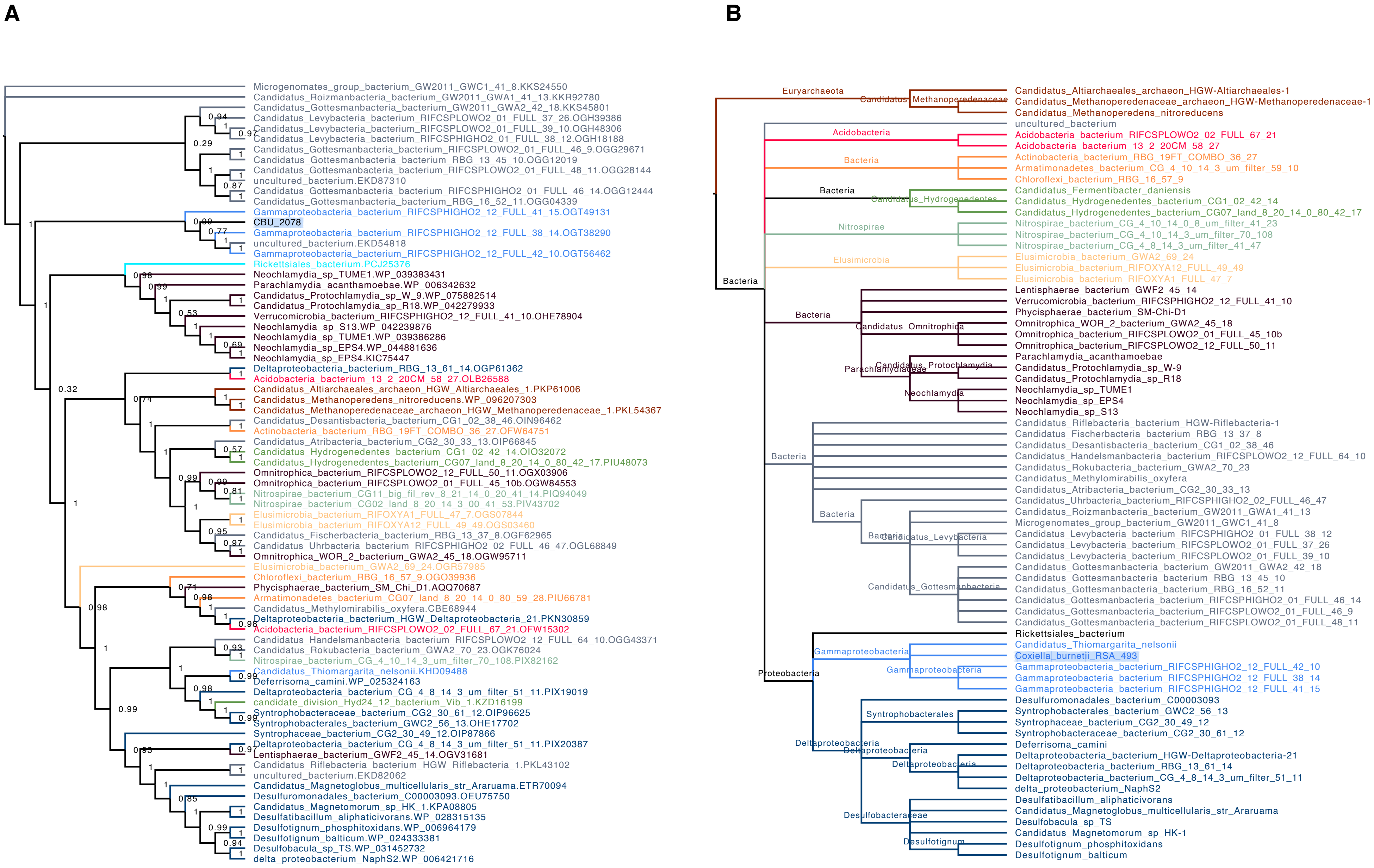
A
B
Microgenomates_group_bacterium_GW2011_GWC1_41_8.KKS24550
Candidatus_Roizmanbacteria_bacterium_GW2011_GWA1_41_13.KKR92780
Candidatus_Gottesmanbacteria_bacterium_GW2011_GWA2_42_18.KKS45801
Candidatus_Levybacteria_bacterium_RIFCSPLOWO2_01_FULL_37_26.OGH39386
Candidatus_Levybacteria_bacterium_RIFCSPLOWO2_01_FULL_39_10.OGH48306
Candidatus_Levybacteria_bacterium_RIFCSPHIGHO2_01_FULL_38_12.OGH18188
Candidatus_Gottesmanbacteria_bacterium_RIFCSPLOWO2_01_FULL_46_9.OGG29671
Candidatus_Gottesmanbacteria_bacterium_RBG_13_45_10.OGG12019
Candidatus_Gottesmanbacteria_bacterium_RIFCSPLOWO2_01_FULL_48_11.OGG28144
uncultured_bacterium.EKD87310
Candidatus_Gottesmanbacteria_bacterium_RIFCSPHIGHO2_01_FULL_46_14.OGG12444
Candidatus_Gottesmanbacteria_bacterium_RBG_16_52_11.OGG04339
Gammaproteobacteria_bacterium_RIFCSPHIGHO2_12_FULL_41_15.OGT49131
CBU_2078
Gammaproteobacteria_bacterium_RIFCSPHIGHO2_12_FULL_38_14.OGT38290
uncultured_bacterium.EKD54818
Gammaproteobacteria_bacterium_RIFCSPHIGHO2_12_FULL_42_10.OGT56462
Rickettsiales_bacterium.PCJ25376
Neochlamydia_sp_TUME1.WP_039383431
Parachlamydia_acanthamoebae.WP_006342632
Candidatus_Protochlamydia_sp_W_9.WP_075882514
Candidatus_Protochlamydia_sp_R18.WP_042279933
Verrucomicrobia_bacterium_RIFCSPHIGHO2_12_FULL_41_10.OHE78904
Neochlamydia_sp_S13.WP_042239876
Neochlamydia_sp_TUME1.WP_039386286
Neochlamydia_sp_EPS4.WP_044881636
Neochlamydia_sp_EPS4.KIC75447
Deltaproteobacteria_bacterium_RBG_13_61_14.OGP61362
Acidobacteria_bacterium_13_2_20CM_58_27.OLB26588
Candidatus_Altiarchaeales_archaeon_HGW_Altiarchaeales_1.PKP61006
Candidatus_Methanoperedens_nitroreducens.WP_096207303
Candidatus_Methanoperedenaceae_archaeon_HGW_Methanoperedenaceae_1.PKL54367
Candidatus_Desantisbacteria_bacterium_CG1_02_38_46.OIN96462
Actinobacteria_bacterium_RBG_19FT_COMBO_36_27.OFW64751
Candidatus_Atribacteria_bacterium_CG2_30_33_13.OIP66845
Candidatus_Hydrogenedentes_bacterium_CG1_02_42_14.OIO32072
Candidatus_Hydrogenedentes_bacterium_CG07_land_8_20_14_0_80_42_17.PIU48073
Omnitrophica_bacterium_RIFCSPLOWO2_12_FULL_50_11.OGX03906
Omnitrophica_bacterium_RIFCSPLOWO2_01_FULL_45_10b.OGW84553
Nitrospirae_bacterium_CG11_big_fil_rev_8_21_14_0_20_41_14.PIQ94049
Nitrospirae_bacterium_CG02_land_8_20_14_3_00_41_53.PIV43702
Elusimicrobia_bacterium_RIFOXYA1_FULL_47_7.OGS07844
Elusimicrobia_bacterium_RIFOXYA12_FULL_49_49.OGS03460
Candidatus_Fischerbacteria_bacterium_RBG_13_37_8.OGF62965
Candidatus_Uhrbacteria_bacterium_RIFCSPHIGHO2_02_FULL_46_47.OGL68849
Omnitrophica_WOR_2_bacterium_GWA2_45_18.OGW95711
Elusimicrobia_bacterium_GWA2_69_24.OGR57985
Chloroflexi_bacterium_RBG_16_57_9.OGO39936
Phycisphaerae_bacterium_SM_Chi_D1.AQQ70687
Armatimonadetes_bacterium_CG07_land_8_20_14_0_80_59_28.PIU66781
Candidatus_Methylomirabilis_oxyfera.CBE68944
Deltaproteobacteria_bacterium_HGW_Deltaproteobacteria_21.PKN30859
Acidobacteria_bacterium_RIFCSPLOWO2_02_FULL_67_21.OFW15302
Candidatus_Handelsmanbacteria_bacterium_RIFCSPLOWO2_12_FULL_64_10.OGG43371
Candidatus_Rokubacteria_bacterium_GWA2_70_23.OGK76024
Nitrospirae_bacterium_CG_4_10_14_3_um_filter_70_108.PIX82162
Candidatus_Thiomargarita_nelsonii.KHD09488
Deferrisoma_camini.WP_025324163
Deltaproteobacteria_bacterium_CG_4_8_14_3_um_filter_51_11.PIX19019
candidate_division_Hyd24_12_bacterium_Vib_1.KZD16199
Syntrophobacteraceae_bacterium_CG2_30_61_12.OIP96625
Syntrophobacterales_bacterium_GWC2_56_13.OHE17702
Syntrophaceae_bacterium_CG2_30_49_12.OIP87866
Deltaproteobacteria_bacterium_CG_4_8_14_3_um_filter_51_11.PIX20387
Lentisphaerae_bacterium_GWF2_45_14.OGV31681
Candidatus_Riflebacteria_bacterium_HGW_Riflebacteria_1.PKL43102
uncultured_bacterium.EKD82062
Candidatus_Magnetoglobus_multicellularis_str_Araruama.ETR70094
Desulfuromonadales_bacterium_C00003093.OEU75750
Candidatus_Magnetomorum_sp_HK_1.KPA08805
Desulfatibacillum_aliphaticivorans.WP_028315135
Desulfotignum_phosphitoxidans.WP_006964179
Desulfotignum_balticum.WP_024333381
Desulfobacula_sp_TS.WP_031452732
delta_proteobacterium_NaphS2.WP_006421716
Euryarchaeota
Candidatus_Methanoperedenaceae
Candidatus_Altiarchaeales_archaeon_HGW-Altiarchaeales-1
Candidatus_Methanoperedenaceae_archaeon_HGW-Methanoperedenaceae-1
Candidatus_Methanoperedens_nitroreducens
uncultured_bacterium
Acidobacteria
Acidobacteria_bacterium_RIFCSPLOWO2_02_FULL_67_21
Acidobacteria_bacterium_13_2_20CM_58_27
Bacteria
Actinobacteria_bacterium_RBG_19FT_COMBO_36_27
Armatimonadetes_bacterium_CG_4_10_14_3_um_filter_59_10
Chloroflexi_bacterium_RBG_16_57_9
Bacteria
Candidatus_Hydrogenedentes
Candidatus_Fermentibacter_daniensis
Candidatus_Hydrogenedentes_bacterium_CG1_02_42_14
Candidatus_Hydrogenedentes_bacterium_CG07_land_8_20_14_0_80_42_17
Nitrospirae
Nitrospirae_bacterium_CG_4_10_14_0_8_um_filter_41_23
Nitrospirae_bacterium_CG_4_10_14_3_um_filter_70_108
Nitrospirae_bacterium_CG_4_8_14_3_um_filter_41_47
Elusimicrobia
Elusimicrobia_bacterium_GWA2_69_24
Elusimicrobia_bacterium_RIFOXYA12_FULL_49_49
Elusimicrobia_bacterium_RIFOXYA1_FULL_47_7
Bacteria
Lentisphaerae_bacterium_GWF2_45_14
Verrucomicrobia_bacterium_RIFCSPHIGHO2_12_FULL_41_10
Phycisphaerae_bacterium_SM-Chi-D1
Bacteria
Candidatus_Omnitrophica
Omnitrophica_WOR_2_bacterium_GWA2_45_18
Omnitrophica_bacterium_RIFCSPLOWO2_01_FULL_45_10b
Omnitrophica_bacterium_RIFCSPLOWO2_12_FULL_50_11
Parachlamydiaceae
Candidatus_Protochlamydia
Neochlamydia
Parachlamydia_acanthamoebae
Candidatus_Protochlamydia_sp_W-9
Candidatus_Protochlamydia_sp_R18
Neochlamydia_sp_TUME1
Neochlamydia_sp_EPS4
Neochlamydia_sp_S13
Bacteria
Candidatus_Riflebacteria_bacterium_HGW-Riflebacteria-1
Candidatus_Fischerbacteria_bacterium_RBG_13_37_8
Candidatus_Desantisbacteria_bacterium_CG1_02_38_46
Candidatus_Handelsmanbacteria_bacterium_RIFCSPLOWO2_12_FULL_64_10
Candidatus_Rokubacteria_bacterium_GWA2_70_23
Candidatus_Methylomirabilis_oxyfera
Candidatus_Atribacteria_bacterium_CG2_30_33_13
Bacteria
Bacteria
Candidatus_Levybacteria
Candidatus_Gottesmanbacteria
Candidatus_Uhrbacteria_bacterium_RIFCSPHIGHO2_02_FULL_46_47
Candidatus_Roizmanbacteria_bacterium_GW2011_GWA1_41_13
Microgenomates_group_bacterium_GW2011_GWC1_41_8
Candidatus_Levybacteria_bacterium_RIFCSPHIGHO2_01_FULL_38_12
Candidatus_Levybacteria_bacterium_RIFCSPLOWO2_01_FULL_37_26
Candidatus_Levybacteria_bacterium_RIFCSPLOWO2_01_FULL_39_10
Candidatus_Gottesmanbacteria_bacterium_GW2011_GWA2_42_18
Candidatus_Gottesmanbacteria_bacterium_RBG_13_45_10
Candidatus_Gottesmanbacteria_bacterium_RBG_16_52_11
Candidatus_Gottesmanbacteria_bacterium_RIFCSPHIGHO2_01_FULL_46_14
Candidatus_Gottesmanbacteria_bacterium_RIFCSPLOWO2_01_FULL_46_9
Candidatus_Gottesmanbacteria_bacterium_RIFCSPLOWO2_01_FULL_48_11
Proteobacteria
Rickettsiales_bacterium
Gammaproteobacteria
Gammaproteobacteria
Candidatus_Thiomargarita_nelsonii
Coxiella_burnetii_RSA_493
Gammaproteobacteria_bacterium_RIFCSPHIGHO2_12_FULL_42_10
Gammaproteobacteria_bacterium_RIFCSPHIGHO2_12_FULL_38_14
Gammaproteobacteria_bacterium_RIFCSPHIGHO2_12_FULL_41_15
Desulfuromonadales_bacterium_C00003093
Syntrophobacterales
Syntrophobacterales_bacterium_GWC2_56_13
Syntrophaceae_bacterium_CG2_30_49_12
Syntrophobacteraceae_bacterium_CG2_30_61_12
Deltaproteobacteria
Deltaproteobacteria
Deltaproteobacteria
Deferrisoma_camini
Deltaproteobacteria_bacterium_HGW-Deltaproteobacteria-21
Deltaproteobacteria_bacterium_RBG_13_61_14
Deltaproteobacteria_bacterium_CG_4_8_14_3_um_filter_51_11
delta_proteobacterium_NaphS2
Desulfobacteraceae
Desulfotignum
Desulfatibacillum_aliphaticivorans
Candidatus_Magnetoglobus_multicellularis_str_Araruama
Desulfobacula_sp_TS
Candidatus_Magnetomorum_sp_HK-1
Desulfotignum_phosphitoxidans
Desulfotignum_balticum

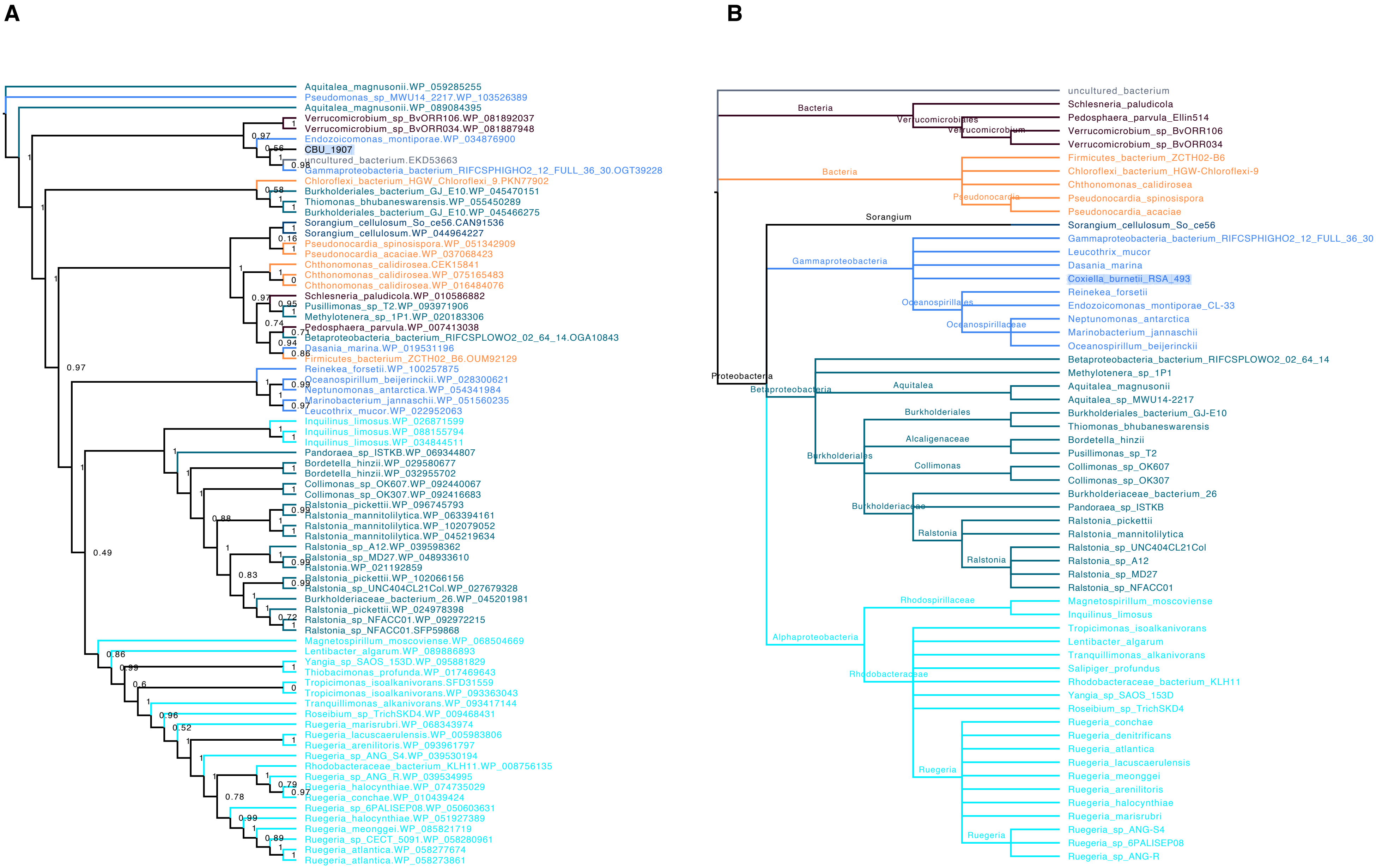
A
Aquitalea_magnusonii.WP_059285255
Pseudomonas_sp_MWU14_2217.WP_103526389
Aquitalea_magnusonii.WP_089084395
Verrucomicrobium_sp_BvORR106.WP_081892037
Verrucomicrobium_sp_BvORR034.WP_081887948
Endozoicomonas_montiporae.WP_034876900
CBU_1907
uncultured_bacterium.EKD53663
Gammaproteobacteria_bacterium_RIFCSPHIGHO2_12_FULL_36_30.OGT39228
Chloroflexi_bacterium_HGW_Chloroflexi_9.PKN77902
Burkholderiales_bacterium_GJ_E10.WP_045470151
Thiomonas_bhubaneswarensis.WP_055450289
Burkholderiales_bacterium_GJ_E10.WP_045466275
Sorangium_cellulosum_So_ce56.CAN91536
Sorangium_cellulosum.WP_044964227
Pseudonocardia_spinosispora.WP_051342909
Pseudonocardia_acaciae.WP_037068423
Chthonomonas_calidirosea.CEK15841
Chthonomonas_calidirosea.WP_075165483
Chthonomonas_calidirosea.WP_016484076
Schlesneria_paludicola.WP_010586882
Pusillimonas_sp_T2.WP_093971906
Methylotenera_sp_1P1.WP_020183306
Pedosphaera_parvula.WP_007413038
Betaproteobacteria_bacterium_RIFCSPLOWO2_02_64_14.OGA10843
Dasania_marina.WP_019531196
Firmicutes_bacterium_ZCTH02_B6.OUM92129
Reinekea_forsetii.WP_100257875
Oceanospirillum_beijerinckii.WP_028300621
Neptunomonas_antarctica.WP_054341984
Marinobacterium_jannaschii.WP_051560235
Leucothrix_mucor.WP_022952063
Inquilinus_limosus.WP_026871599
Inquilinus_limosus.WP_088155794
Inquilinus_limosus.WP_034844511
Pandoraea_sp_ISTKB.WP_069344807
Bordetella_hinzii.WP_029580677
Bordetella_hinzii.WP_032955702
Collimonas_sp_OK607.WP_092440067
Collimonas_sp_OK307.WP_092416683
Ralstonia_pickettii.WP_096745793
Ralstonia_mannitolilytica.WP_063394161
Ralstonia_mannitolilytica.WP_102079052
Ralstonia_mannitolilytica.WP_045219634
Ralstonia_sp_A12.WP_039598362
Ralstonia_sp_MD27.WP_048933610
Ralstonia.WP_021192859
Ralstonia_pickettii.WP_102066156
Ralstonia_sp_UNC404CL21Col.WP_027679328
Burkholderiaceae_bacterium_26.WP_045201981
Ralstonia_pickettii.WP_024978398
Ralstonia_sp_NFACC01.WP_092972215
Ralstonia_sp_NFACC01.SFP59868
Magnetospirillum_moscoviense.WP_068504669
Lentibacter_algarum.WP_089886893
Yangia_sp_SAOS_153D.WP_095881829
Thiobacimonas_profunda.WP_017469643
Tropicimonas_isoalkanivorans.SFD31559
Tropicimonas_isoalkanivorans.WP_093363043
Tranquillimonas_alkanivorans.WP_093417144
Roseibium_sp_TrichSKD4.WP_009468431
Ruegeria_marisrubri.WP_068343974
Ruegeria_lacuscaerulensis.WP_005983806
Ruegeria_arenilitoris.WP_093961797
Ruegeria_sp_ANG_S4.WP_039530194
Rhodobacteraceae_bacterium_KLH11.WP_008756135
Ruegeria_sp_ANG_R.WP_039534995
Ruegeria_halocynthiae.WP_074735029
Ruegeria_conchae.WP_010439424
Ruegeria_sp_6PALISEP08.WP_050603631
Ruegeria_halocynthiae.WP_051927389
Ruegeria_meonggei.WP_085821719
Ruegeria_sp_CECT_5091.WP_058280961
Ruegeria_atlantica.WP_058277674
Ruegeria_atlantica.WP_058273861
B
uncultured_bacterium
Bacteria
Verrucomicrobiales
Verrucomicrobium
Schlesneria_paludicola
Pedosphaera_parvula_Ellin514
Verrucomicrobium_sp_BvORR106
Verrucomicrobium_sp_BvORR034
Firmicutes_bacterium_ZCTH02-B6
Chloroflexi_bacterium_HGW-Chloroflexi-9
Chthonomonas_calidirosea
Pseudonocardia
Pseudonocardia_spinosispora
Pseudonocardia_acaciae
Sorangium
Sorangium_cellulosum_So_ce56
Gammaproteobacteria
Gammaproteobacteria_bacterium_RIFCSPHIGHO2_12_FULL_36_30
Leucothrix_mucor
Dasania_marina
Coxiella_burnetii_RSA_493
Oceanospirillales
Reinekea_forsetii
Endozoicomonas_montiporae_CL-33
Oceanospirillaceae
Neptunomonas_antarctica
Marinobacterium_jannaschii
Oceanospirillum_beijerinckii
Proteobacteria
Betaproteobacteria
Betaproteobacteria_bacterium_RIFCSPLOWO2_02_64_14
Methylotenera_sp_1P1
Aquitalea
Aquitalea_magnusonii
Aquitalea_sp_MWU14-2217
Burkholderiales
Burkholderiales_bacterium_GJ-E10
Thiomonas_bhubaneswarensis
Alcaligenaceae
Bordetella_hinzii
Pusillimonas_sp_T2
Collimonas
Collimonas_sp_OK607
Collimonas_sp_OK307
Burkholderiaceae
Burkholderiaceae_bacterium_26
Pandoraea_sp_ISTKB
Ralstonia
Ralstonia_pickettii
Ralstonia_mannitolilytica
Ralstonia_sp_UNC404CL21Col
Ralstonia_sp_A12
Ralstonia_sp_MD27
Ralstonia_sp_NFACC01
Alphaproteobacteria
Rhodospirillaceae
Magnetospirillum_moscoviense
Inquilinus_limosus
Rhodobacteraceae
Tropicimonas_isoalkanivorans
Lentibacter_algarum
Tranquillimonas_alkanivorans
Salipiger_profundus
Rhodobacteraceae_bacterium_KLH11
Yangia_sp_SAOS_153D
Roseibium_sp_TrichSKD4
Ruegeria
Ruegeria_conchae
Ruegeria_denitrificans
Ruegeria_atlantica
Ruegeria_lacuscaerulensis
Ruegeria_meonggei
Ruegeria_arenilitoris
Ruegeria_halocynthiae
Ruegeria_marisrubri
Ruegeria_sp_ANG-S4
Ruegeria_sp_6PALISEP08
Ruegeria_sp_ANG-R

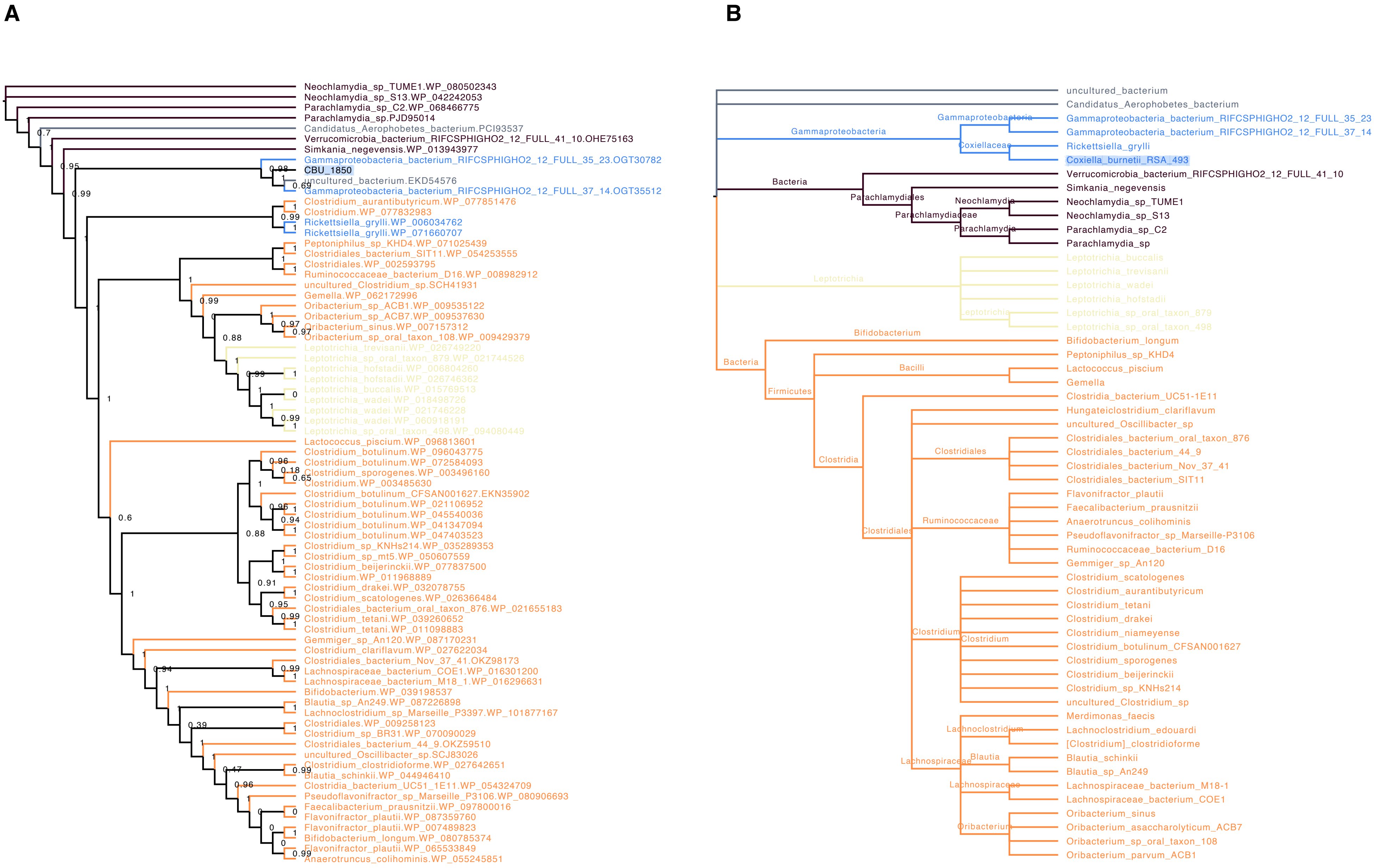
A
Neochlamydia_sp_TUME1.WP_080502343
Neochlamydia_sp_S13.WP_042242053
Parachlamydia_sp_C2.WP_068466775
Parachlamydia_sp.PJD95014
Candidatus_Aerophobetes_bacterium.PCI93537
Verrucomicrobia_bacterium_RIFCSPHIGHO2_12_FULL_41_10.OHE75163
Simkania_negevensis.WP_013943977
Gammaproteobacteria_bacterium_RIFCSPHIGHO2_12_FULL_35_23.OGT30782
CBU_1850
uncultured_bacterium.EKD54576
Gammaproteobacteria_bacterium_RIFCSPHIGHO2_12_FULL_37_14.OGT35512
Clostridium_aurantibutyricum.WP_077851476
Clostridium.WP_077832983
Rickettsiella_grylli.WP_006034762
Rickettsiella_grylli.WP_071660707
Peptoniphilus_sp_KHD4.WP_071025439
Clostridiales_bacterium_SIT11.WP_054253555
Clostridiales.WP_002593795
Ruminococcaceae_bacterium_D16.WP_008982912
uncultured_Clostridium_sp.SCH41931
Gemella.WP_062172996
Oribacterium_sp_ACB1.WP_009535122
Oribacterium_sp_ACB7.WP_009537630
Oribacterium_sinus.WP_007157312
Oribacterium_sp_oral_taxon_108.WP_009429379
Leptotrichia_trevisanii.WP_026749220
Leptotrichia_sp_oral_taxon_879.WP_021744526
Leptotrichia_hofstadii.WP_006804260
Leptotrichia_hofstadii.WP_026746362
Leptotrichia_buccalis.WP_015769513
Leptotrichia_wadei.WP_018498726
Leptotrichia_wadei.WP_021746228
Leptotrichia_wadei.WP_060918191
Leptotrichia_sp_oral_taxon_498.WP_094080449
Lactococcus_piscium.WP_096813601
Clostridium_botulinum.WP_096043775
Clostridium_botulinum.WP_072584093
Clostridium_sporogenes.WP_003496160
Clostridium.WP_003485630
Clostridium_botulinum_CFSAN001627.EKN35902
Clostridium_botulinum.WP_021106952
Clostridium_botulinum.WP_045540036
Clostridium_botulinum.WP_041347094
Clostridium_botulinum.WP_047403523
Clostridium_sp_KNHs214.WP_035289353
Clostridium_sp_mt5.WP_050607559
Clostridium_beijerinckii.WP_077837500
Clostridium.WP_011968889
Clostridium_drakei.WP_032078755
Clostridium_scatologenes.WP_026366484
Clostridiales_bacterium_oral_taxon_876.WP_021655183
Clostridium_tetani.WP_039260652
Clostridium_tetani.WP_011098883
Gemmiger_sp_An120.WP_087170231
Clostridium_clariflavum.WP_027622034
Clostridiales_bacterium_Nov_37_41.OKZ98173
Lachnospiraceae_bacterium_COE1.WP_016301200
Lachnospiraceae_bacterium_M18_1.WP_016296631
Bifidobacterium.WP_039198537
Blautia_sp_An249.WP_087226898
Lachnoclostridium_sp_Marseille_P3397.WP_101877167
Clostridiales.WP_009258123
Clostridium_sp_BR31.WP_070090029
Clostridiales_bacterium_44_9.OKZ59510
uncultured_Oscillibacter_sp.SCJ83026
Clostridium_clostridioforme.WP_027642651
Blautia_schinkii.WP_044946410
Clostridia_bacterium_UC51_1E11.WP_054324709
Pseudoflavonifractor_sp_Marseille_P3106.WP_080906693
Faecalibacterium_prausnitzii.WP_097800016
Flavonifractor_plautii.WP_087359760
Flavonifractor_plautii.WP_007489823
Bifidobacterium_longum.WP_080785374
Flavonifractor_plautii.WP_065533849
Anaerotruncus_colihominis.WP_055245851
B
uncultured_bacterium
Candidatus_Aerophobetes_bacterium
Gammaproteobacteria
Gammaproteobacteria
Coxiellaceae
Gammaproteobacteria_bacterium_RIFCSPHIGHO2_12_FULL_35_23
Gammaproteobacteria_bacterium_RIFCSPHIGHO2_12_FULL_37_14
Rickettsiella_grylli
Coxiella_burnetii_RSA_493
Bacteria
Verrucomicrobia_bacterium_RIFCSPHIGHO2_12_FULL_41_10
Parachlamydiales
Simkania_negevensis
Neochlamydia
Neochlamydia_sp_TUME1
Parachlamydiaceae
Neochlamydia_sp_S13
Parachlamydia
Parachlamydia_sp_C2
Parachlamydia_sp
Leptotrichia
Leptotrichia_buccalis
Leptotrichia_trevisanii
Leptotrichia_wadei
Leptotrichia_hofstadii
Leptotrichia
Leptotrichia_sp_oral_taxon_879
Leptotrichia_sp_oral_taxon_498
Bifidobacterium
Bifidobacterium_longum
Bacteria
Peptoniphilus_sp_KHD4
Bacilli
Lactococcus_piscium
Gemella
Firmicutes
Clostridia_bacterium_UC51-1E11
Hungateiclostridium_clariflavum
uncultured_Oscillibacter_sp
Clostridiales
Clostridiales_bacterium_oral_taxon_876
Clostridiales_bacterium_44_9
Clostridia
Clostridiales_bacterium_Nov_37_41
Clostridiales_bacterium_SIT11
Flavonifractor_plautii
Faecalibacterium_prausnitzii
Ruminococcaceae
Anaerotruncus_colihominis
Clostridiales
Pseudoflavonifractor_sp_Marseille-P3106
Ruminococcaceae_bacterium_D16
Gemmiger_sp_An120
Clostridium_scatologenes
Clostridium_aurantibutyricum
Clostridium_tetani
Clostridium_drakei
Clostridium
Clostridium_niameyense
Clostridium
Clostridium_botulinum_CFSAN001627
Clostridium_sporogenes
Clostridium_beijerinckii
Clostridium_sp_KNHs214
uncultured_Clostridium_sp
Merdimonas_faecis
Lachnoclostridium
Lachnoclostridium_edouardi
[Clostridium]_clostridioforme
Blautia
Blautia_schinkii
Lachnospiraceae
Blautia_sp_An249
Lachnospiraceae
Lachnospiraceae_bacterium_M18-1
Lachnospiraceae_bacterium_COE1
Oribacterium_sinus
Oribacterium
Oribacterium_asaccharolyticum_ACB7
Oribacterium_sp_oral_taxon_108
Oribacterium_parvum_ACB1

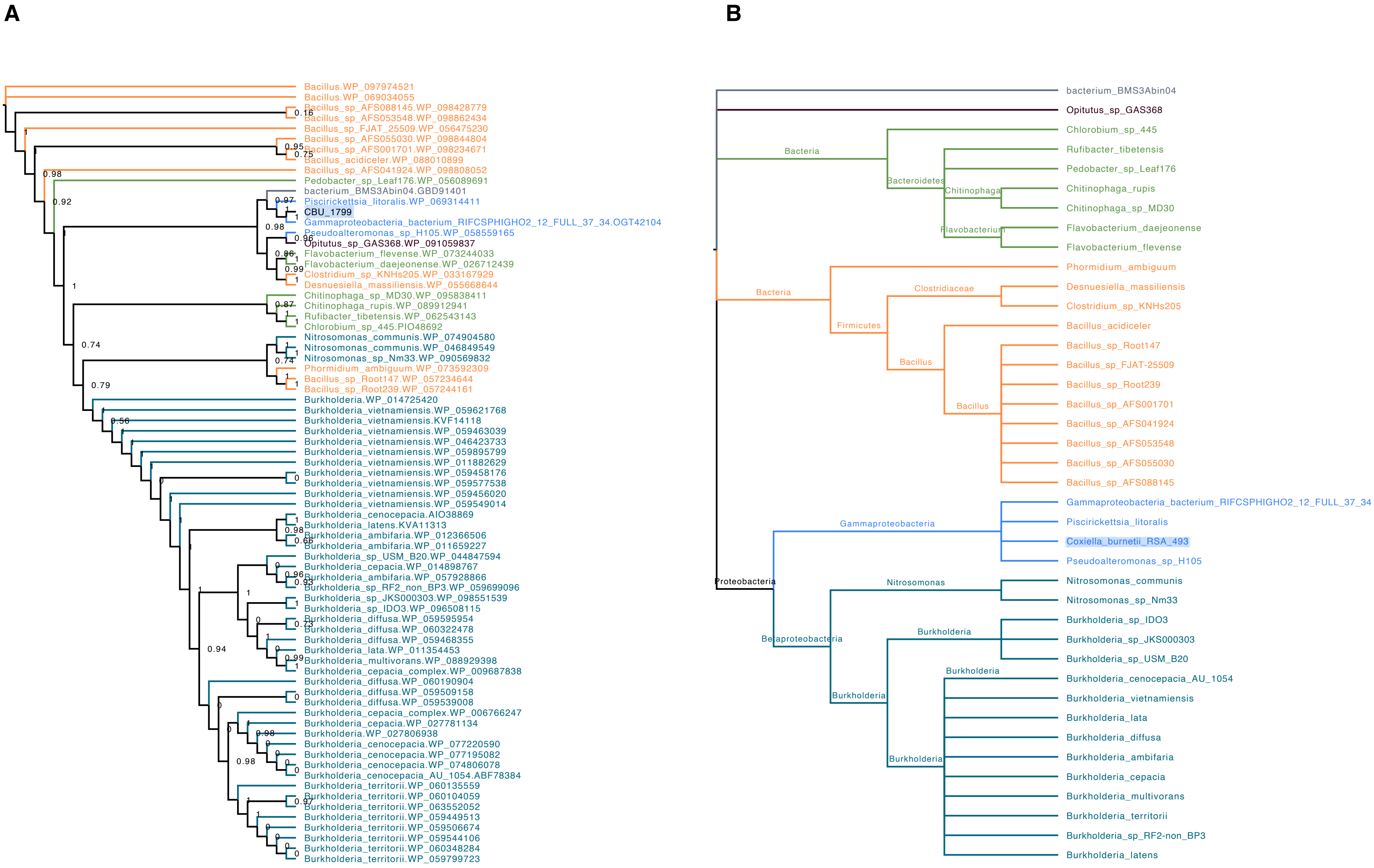
A
Bacillus.WP_097974521
Bacillus.WP_069034055
Bacillus_sp_AFS088145.WP_098428779
Bacillus_sp_AFS053548.WP_098862434
Bacillus_sp_FJAT_25509.WP_056475230
Bacillus_sp_AFS055030.WP_098844804
Bacillus_sp_AFS001701.WP_098234671
Bacillus_acidiceler.WP_088010899
Bacillus_sp_AFS041924.WP_098808052
Pedobacter_sp_Leaf176.WP_056089691
bacterium_BMS3Abin04.GBD91401
Piscirickettsia_litoralis.WP_069314411
CBU_1799
Gammaproteobacteria_bacterium_RIFCSPHIGHO2_12_FULL_37_34.OGT42104
Pseudoalteromonas_sp_H105.WP_058559165
Opitutus_sp_GAS368.WP_091059837
Flavobacterium_flevense.WP_073244033
Flavobacterium_daejeonense.WP_026712439
Clostridium_sp_KNHs205.WP_033167929
Desnuesiella_massiliensis.WP_055668644
Chitinophaga_sp_MD30.WP_095838411
Chitinophaga_rupis.WP_089912941
Rufibacter_tibetensis.WP_062543143
Chlorobium_sp_445.PIO48692
Nitrosomonas_communis.WP_074904580
Nitrosomonas_communis.WP_046849549
Nitrosomonas_sp_Nm33.WP_090569832
Phormidium_ambiguum.WP_073592309
Bacillus_sp_Root147.WP_057234644
Bacillus_sp_Root239.WP_057244161
Burkholderia.WP_014725420
Burkholderia_vietnamiensis.WP_059621768
Burkholderia_vietnamiensis.KVF14118
Burkholderia_vietnamiensis.WP_059463039
Burkholderia_vietnamiensis.WP_046423733
Burkholderia_vietnamiensis.WP_059895799
Burkholderia_vietnamiensis.WP_011882629
Burkholderia_vietnamiensis.WP_059458176
Burkholderia_vietnamiensis.WP_059577538
Burkholderia_vietnamiensis.WP_059456020
Burkholderia_vietnamiensis.WP_059549014
Burkholderia_cenocepacia.AIO38869
Burkholderia_latens.KVA11313
Burkholderia_ambifaria.WP_012366506
Burkholderia_ambifaria.WP_011659227
Burkholderia_sp_USM_B20.WP_044847594
Burkholderia_cepacia.WP_014898767
Burkholderia_ambifaria.WP_057928866
Burkholderia_sp_RF2_non_BP3.WP_059699096
Burkholderia_sp_JKS000303.WP_098551539
Burkholderia_sp_IDO3.WP_096508115
Burkholderia_diffusa.WP_059595954
Burkholderia_diffusa.WP_060322478
Burkholderia_diffusa.WP_059468355
Burkholderia_lata.WP_011354453
Burkholderia_multivorans.WP_088929398
Burkholderia_cepacia_complex.WP_009687838
Burkholderia_diffusa.WP_060190904
Burkholderia_diffusa.WP_059509158
Burkholderia_diffusa.WP_059539008
Burkholderia_cepacia_complex.WP_006766247
Burkholderia_cepacia.WP_027781134
Burkholderia.WP_027806938
Burkholderia_cenocepacia.WP_077220590
Burkholderia_cenocepacia.WP_077195082
Burkholderia_cenocepacia.WP_074806078
Burkholderia_cenocepacia_AU_1054.ABF78384
Burkholderia_territorii.WP_060135559
Burkholderia_territorii.WP_060104059
Burkholderia_territorii.WP_063552052
Burkholderia_territorii.WP_059449513
Burkholderia_territorii.WP_059506674
Burkholderia_territorii.WP_059544106
Burkholderia_territorii.WP_060348284
Burkholderia_territorii.WP_059799723
B
bacterium_BMS3Abin04
Opitutus_sp_GAS368
Bacteria
Bacteroidetes
Chitinophaga
Flavobacterium
Chlorobium_sp_445
Rufibacter_tibetensis
Pedobacter_sp_Leaf176
Chitinophaga_rupis
Chitinophaga_sp_MD30
Flavobacterium_daejeonense
Flavobacterium_flevense
Bacteria
Firmicutes
Clostridiaceae
Bacillus
Bacillus
Phormidium_ambiguum
Desnuesiella_massiliensis
Clostridium_sp_KNHs205
Bacillus_acidiceler
Bacillus_sp_Root147
Bacillus_sp_FJAT-25509
Bacillus_sp_Root239
Bacillus_sp_AFS001701
Bacillus_sp_AFS041924
Bacillus_sp_AFS053548
Bacillus_sp_AFS055030
Bacillus_sp_AFS088145
Proteobacteria
Gammaproteobacteria
Gammaproteobacteria_bacterium_RIFCSPHIGHO2_12_FULL_37_34
Piscirickettsia_litoralis
Coxiella_burnetii_RSA_493
Pseudoalteromonas_sp_H105
Betaproteobacteria
Nitrosomonas
Nitrosomonas_communis
Nitrosomonas_sp_Nm33
Burkholderia
Burkholderia
Burkholderia
Burkholderia
Burkholderia_sp_IDO3
Burkholderia_sp_JKS000303
Burkholderia_sp_USM_B20
Burkholderia_cenocepacia_AU_1054
Burkholderia_vietnamiensis
Burkholderia_lata
Burkholderia_diffusa
Burkholderia_ambifaria
Burkholderia_cepacia
Burkholderia_multivorans
Burkholderia_territorii
Burkholderia_sp_RF2-non_BP3
Burkholderia_latens

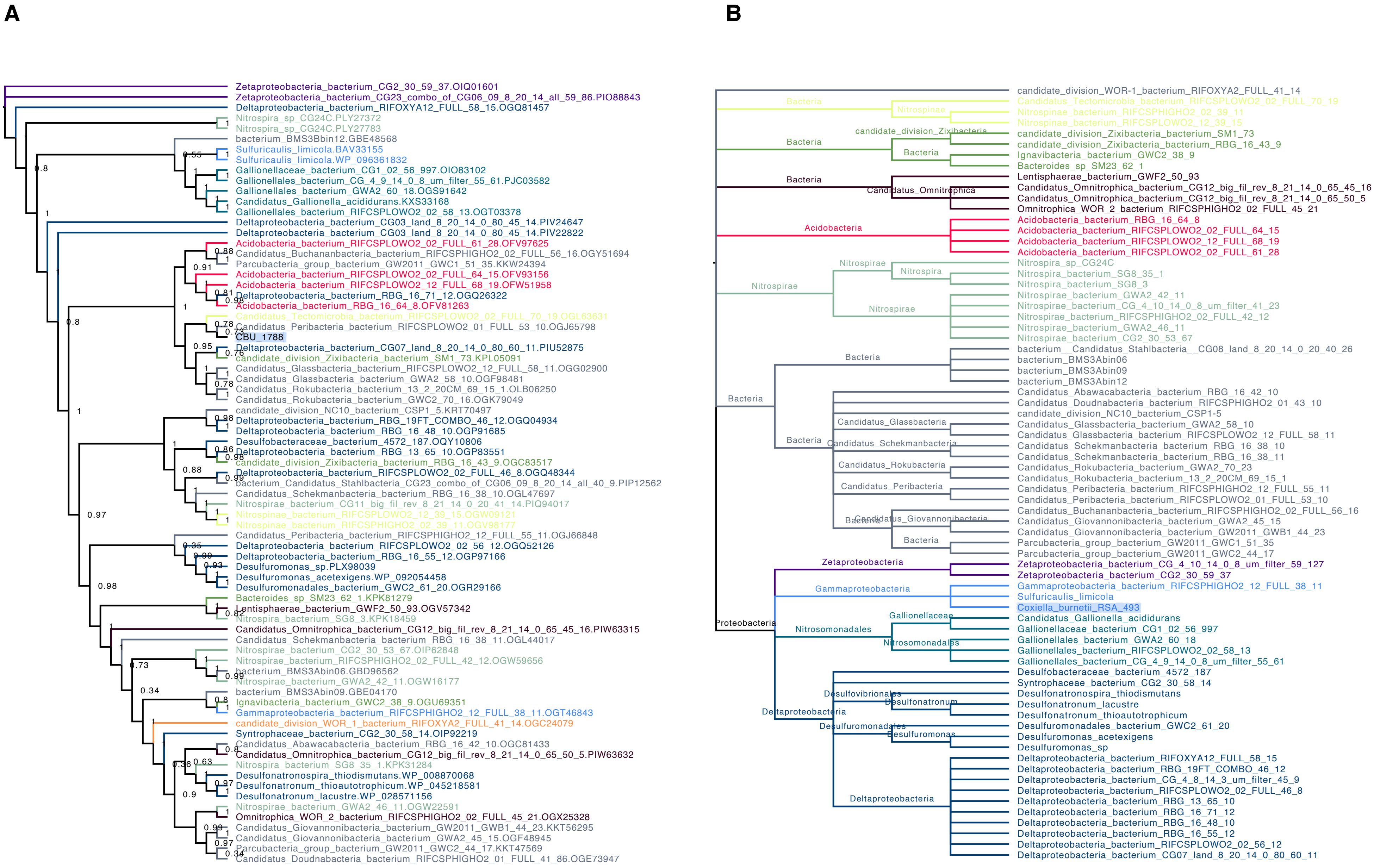
A
Zetaproteobacteria_bacterium_CG2_30_59_37.OIQ01601
Zetaproteobacteria_bacterium_CG23_combo_of_CG06_09_8_20_14_all_59_86.PIO88843
Deltaproteobacteria_bacterium_RIFOXYA12_FULL_58_15.OGQ81457
Nitrospira_sp_CG24C.PLY27372
Nitrospira_sp_CG24C.PLY27783
bacterium_BMS3Bbin12.GBE48568
Sulfuricaulis_limicola.BAV33155
Sulfuricaulis_limicola.WP_096361832
Gallionellaceae_bacterium_CG1_02_56_997.OIO83102
Gallionellales_bacterium_CG_4_9_14_0_8_um_filter_55_61.PJC03582
Gallionellales_bacterium_GWA2_60_18.OGS91642
Candidatus_Gallionella_acididurans.KXS33168
Gallionellales_bacterium_RIFCSPLOWO2_02_58_13.OGT03378
Deltaproteobacteria_bacterium_CG03_land_8_20_14_0_80_45_14.PIV24647
Deltaproteobacteria_bacterium_CG03_land_8_20_14_0_80_45_14.PIV22822
Acidobacteria_bacterium_RIFCSPLOWO2_02_FULL_61_28.OFV97625
Candidatus_Buchananbacteria_bacterium_RIFCSPHIGHO2_02_FULL_56_16.OGY51694
Parcubacteria_group_bacterium_GW2011_GWC1_51_35.KKW24394
Acidobacteria_bacterium_RIFCSPLOWO2_02_FULL_64_15.OFV93156
Acidobacteria_bacterium_RIFCSPLOWO2_12_FULL_68_19.OFW51958
Deltaproteobacteria_bacterium_RBG_16_71_12.OGQ26322
Acidobacteria_bacterium_RBG_16_64_8.OFV81263
Candidatus_Tectomicrobia_bacterium_RIFCSPLOWO2_02_FULL_70_19.OGL63631
Candidatus_Peribacteria_bacterium_RIFCSPLOWO2_01_FULL_53_10.OGJ65798
CBU_1788
Deltaproteobacteria_bacterium_CG07_land_8_20_14_0_80_60_11.PIU52875
candidate_division_Zixibacteria_bacterium_SM1_73.KPL05091
Candidatus_Glassbacteria_bacterium_RIFCSPLOWO2_12_FULL_58_11.OGG02900
Candidatus_Glassbacteria_bacterium_GWA2_58_10.OGF98481
Candidatus_Rokubacteria_bacterium_13_2_20CM_69_15_1.OLB06250
Candidatus_Rokubacteria_bacterium_GWC2_70_16.OGK79049
candidate_division_NC10_bacterium_CSP1_5.KRT70497
Deltaproteobacteria_bacterium_RBG_19FT_COMBO_46_12.OGQ04934
Deltaproteobacteria_bacterium_RBG_16_48_10.OGP91685
Desulfobacteraceae_bacterium_4572_187.OQY10806
Deltaproteobacteria_bacterium_RBG_13_65_10.OGP83551
candidate_division_Zixibacteria_bacterium_RBG_16_43_9.OGC83517
Deltaproteobacteria_bacterium_RIFCSPLOWO2_02_FULL_46_8.OGQ48344
bacterium_Candidatus_Stahlbacteria_CG23_combo_of_CG06_09_8_20_14_all_40_9.PIP12562
Candidatus_Schekmanbacteria_bacterium_RBG_16_38_10.OGL47697
Nitrospirae_bacterium_CG11_big_fil_rev_8_21_14_0_20_41_14.PIQ94017
Nitrospinae_bacterium_RIFCSPLOWO2_12_39_15.OGW09121
Nitrospinae_bacterium_RIFCSPHIGHO2_02_39_11.OGV98177
Candidatus_Peribacteria_bacterium_RIFCSPHIGHO2_12_FULL_55_11.OGJ66848
Deltaproteobacteria_bacterium_RIFCSPLOWO2_02_56_12.OGQ52126
Deltaproteobacteria_bacterium_RBG_16_55_12.OGP97166
Desulfuromonas_sp.PLX98039
Desulfuromonas_acetexigens.WP_092054458
Desulfuromonadales_bacterium_GWC2_61_20.OGR29166
Bacteroides_sp_SM23_62_1.KPK81279
Lentisphaerae_bacterium_GWF2_50_93.OGV57342
Nitrospira_bacterium_SG8_3.KPK18459
Candidatus_Omnitrophica_bacterium_CG12_big_fil_rev_8_21_14_0_65_45_16.PIW63315
Candidatus_Schekmanbacteria_bacterium_RBG_16_38_11.OGL44017
Nitrospirae_bacterium_CG2_30_53_67.OIP62848
Nitrospirae_bacterium_RIFCSPHIGHO2_02_FULL_42_12.OGW59656
bacterium_BMS3Abin06.GBD96562
Nitrospirae_bacterium_GWA2_42_11.OGW16177
bacterium_BMS3Abin09.GBE04170
Ignavibacteria_bacterium_GWC2_38_9.OGU69351
Gammaproteobacteria_bacterium_RIFCSPHIGHO2_12_FULL_38_11.OGT46843
candidate_division_WOR_1_bacterium_RIFOXYA2_FULL_41_14.OGC24079
Syntrophaceae_bacterium_CG2_30_58_14.OIP92219
Candidatus_Abawacabacteria_bacterium_RBG_16_42_10.OGC81433
Candidatus_Omnitrophica_bacterium_CG12_big_fil_rev_8_21_14_0_65_50_5.PIW63632
Nitrospira_bacterium_SG8_35_1.KPK31284
Desulfonatronospira_thiodismutans.WP_008870068
Desulfonatronum_thioautotrophicum.WP_045218581
Desulfonatronum_lacustre.WP_028571156
Nitrospirae_bacterium_GWA2_46_11.OGW22591
Omnitrophica_WOR_2_bacterium_RIFCSPHIGHO2_02_FULL_45_21.OGX25328
Candidatus_Giovannonibacteria_bacterium_GW2011_GWB1_44_23.KKT56295
Candidatus_Giovannonibacteria_bacterium_GWA2_45_15.OGF48945
Parcubacteria_group_bacterium_GW2011_GWC2_44_17.KKT47569
Candidatus_Doudnabacteria_bacterium_RIFCSPHIGHO2_01_FULL_41_86.OGE73947
B
candidate_division_WOR-1_bacterium_RIFOXYA2_FULL_41_14
Bacteria
Nitrospinae
Candidatus_Tectomicrobia_bacterium_RIFCSPLOWO2_02_FULL_70_19
Nitrospinae_bacterium_RIFCSPHIGHO2_02_39_11
Nitrospinae_bacterium_RIFCSPLOWO2_12_39_15
candidate_division_Zixibacteria
candidate_division_Zixibacteria_bacterium_SM1_73
candidate_division_Zixibacteria_bacterium_RBG_16_43_9
Ignavibacteria_bacterium_GWC2_38_9
Bacteroides_sp_SM23_62_1
Lentisphaerae_bacterium_GWF2_50_93
Candidatus_Omnitrophica
Candidatus_Omnitrophica_bacterium_CG12_big_fil_rev_8_21_14_0_65_45_16
Candidatus_Omnitrophica_bacterium_CG12_big_fil_rev_8_21_14_0_65_50_5
Omnitrophica_WOR_2_bacterium_RIFCSPHIGHO2_02_FULL_45_21
Acidobacteria
Acidobacteria_bacterium_RBG_16_64_8
Acidobacteria_bacterium_RIFCSPLOWO2_02_FULL_64_15
Acidobacteria_bacterium_RIFCSPLOWO2_12_FULL_68_19
Acidobacteria_bacterium_RIFCSPLOWO2_02_FULL_61_28
Nitrospirae
Nitrospira
Nitrospira_sp_CG24C
Nitrospira_bacterium_SG8_35_1
Nitrospira_bacterium_SG8_3
Nitrospirae_bacterium_GWA2_42_11
Nitrospirae_bacterium_CG_4_10_14_0_8_um_filter_41_23
Nitrospirae_bacterium_RIFCSPHIGHO2_02_FULL_42_12
Nitrospirae_bacterium_GWA2_46_11
Nitrospirae_bacterium_CG2_30_53_67
bacterium__Candidatus_Stahlbacteria__CG08_land_8_20_14_0_20_40_26
bacterium_BMS3Abin06
bacterium_BMS3Abin09
bacterium_BMS3Abin12
Candidatus_Abawacabacteria_bacterium_RBG_16_42_10
Candidatus_Doudnabacteria_bacterium_RIFCSPHIGHO2_01_43_10
candidate_division_NC10_bacterium_CSP1-5
Candidatus_Glassbacteria
Candidatus_Glassbacteria_bacterium_GWA2_58_10
Candidatus_Glassbacteria_bacterium_RIFCSPLOWO2_12_FULL_58_11
Candidatus_Schekmanbacteria
Candidatus_Schekmanbacteria_bacterium_RBG_16_38_10
Candidatus_Schekmanbacteria_bacterium_RBG_16_38_11
Candidatus_Rokubacteria
Candidatus_Rokubacteria_bacterium_GWA2_70_23
Candidatus_Rokubacteria_bacterium_13_2_20CM_69_15_1
Candidatus_Peribacteria
Candidatus_Peribacteria_bacterium_RIFCSPHIGHO2_12_FULL_55_11
Candidatus_Peribacteria_bacterium_RIFCSPLOWO2_01_FULL_53_10
Candidatus_Buchananbacteria_bacterium_RIFCSPHIGHO2_02_FULL_56_16
Candidatus_Giovannonibacteria
Candidatus_Giovannonibacteria_bacterium_GWA2_45_15
Candidatus_Giovannonibacteria_bacterium_GW2011_GWB1_44_23
Parcubacteria_group_bacterium_GW2011_GWC1_51_35
Parcubacteria_group_bacterium_GW2011_GWC2_44_17
Proteobacteria
Zetaproteobacteria
Zetaproteobacteria_bacterium_CG_4_10_14_0_8_um_filter_59_127
Zetaproteobacteria_bacterium_CG2_30_59_37
Gammaproteobacteria
Gammaproteobacteria_bacterium_RIFCSPHIGHO2_12_FULL_38_11
Sulfuricaulis_limicola
Coxiella_burnetii_RSA_493
Nitrosomonadales
Gallionellaceae
Candidatus_Gallionella_acididurans
Gallionellaceae_bacterium_CG1_02_56_997
Gallionellales_bacterium_GWA2_60_18
Gallionellales_bacterium_RIFCSPLOWO2_02_58_13
Gallionellales_bacterium_CG_4_9_14_0_8_um_filter_55_61
Deltaproteobacteria
Desulfovibrionales
Desulfonatronum
Desulfuromonadales
Desulfuromonas
Desulfobacteraceae_bacterium_4572_187
Syntrophaceae_bacterium_CG2_30_58_14
Desulfonatronospira_thiodismutans
Desulfonatronum_lacustre
Desulfonatronum_thioautotrophicum
Desulfuromonadales_bacterium_GWC2_61_20
Desulfuromonas_acetexigens
Desulfuromonas_sp
Deltaproteobacteria_bacterium_RIFOXYA12_FULL_58_15
Deltaproteobacteria_bacterium_RBG_19FT_COMBO_46_12
Deltaproteobacteria_bacterium_CG_4_8_14_3_um_filter_45_9
Deltaproteobacteria_bacterium_RIFCSPLOWO2_02_FULL_46_8
Deltaproteobacteria_bacterium_RBG_13_65_10
Deltaproteobacteria_bacterium_RBG_16_71_12
Deltaproteobacteria_bacterium_RBG_16_48_10
Deltaproteobacteria_bacterium_RBG_16_55_12
Deltaproteobacteria_bacterium_RIFCSPLOWO2_02_56_12
Deltaproteobacteria_bacterium_CG07_land_8_20_14_0_80_60_11

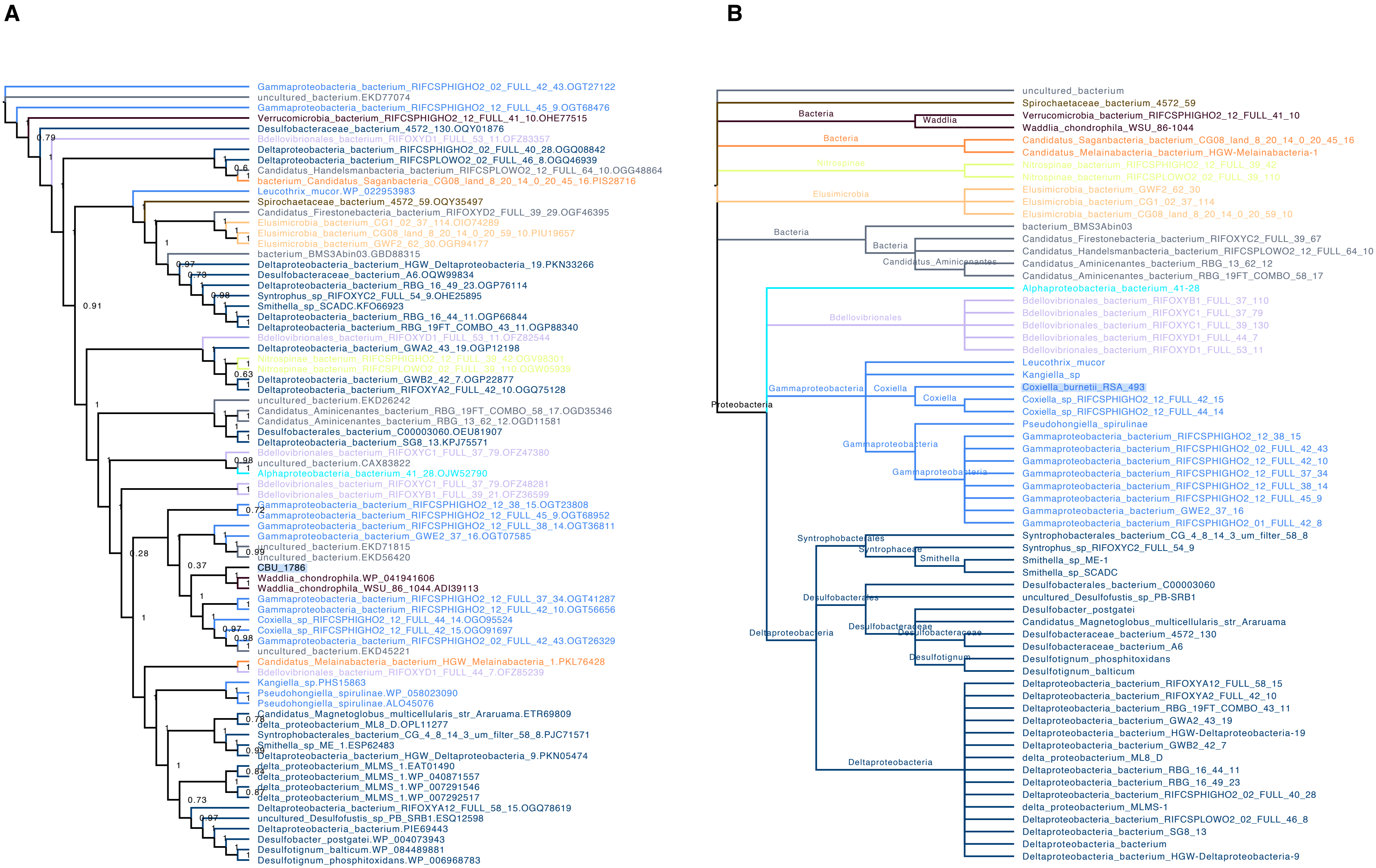
A
Gammaproteobacteria_bacterium_RIFCSPHIGHO2_02_FULL_42_43.OGT27122
uncultured_bacterium.EKD77074
Gammaproteobacteria_bacterium_RIFCSPHIGHO2_12_FULL_45_9.OGT68476
Verrucomicrobia_bacterium_RIFCSPHIGHO2_12_FULL_41_10.OHE77515
Desulfobacteraceae_bacterium_4572_130.OQY01876
Bdellovibrionales_bacterium_RIFOXYD1_FULL_53_11.OFZ83357
Deltaproteobacteria_bacterium_RIFCSPHIGHO2_02_FULL_40_28.OGQ08842
Deltaproteobacteria_bacterium_RIFCSPLOWO2_02_FULL_46_8.OGQ46939
Candidatus_Handelsmanbacteria_bacterium_RIFCSPLOWO2_12_FULL_64_10.OGG48864
bacterium_Candidatus_Saganbacteria_CG08_land_8_20_14_0_20_45_16.PIS28716
Leucothrix_mucor.WP_022953983
Spirochaetaceae_bacterium_4572_59.OQY35497
Candidatus_Firestonebacteria_bacterium_RIFOXYD2_FULL_39_29.OGF46395
Elusimicrobia_bacterium_CG1_02_37_114.OIO74289
Elusimicrobia_bacterium_CG08_land_8_20_14_0_20_59_10.PIU19657
Elusimicrobia_bacterium_GWF2_62_30.OGR94177
bacterium_BMS3Abin03.GBD88315
Deltaproteobacteria_bacterium_HGW_Deltaproteobacteria_19.PKN33266
Desulfobacteraceae_bacterium_A6.OQW99834
Deltaproteobacteria_bacterium_RBG_16_49_23.OGP76114
Syntrophus_sp_RIFOXYC2_FULL_54_9.OHE25895
Smithella_sp_SCADC.KFO66923
Deltaproteobacteria_bacterium_RBG_16_44_11.OGP66844
Deltaproteobacteria_bacterium_RBG_19FT_COMBO_43_11.OGP88340
Bdellovibrionales_bacterium_RIFOXYD1_FULL_53_11.OFZ82544
Deltaproteobacteria_bacterium_GWA2_43_19.OGP12198
Nitrospinae_bacterium_RIFCSPHIGHO2_12_FULL_39_42.OGV98301
Nitrospinae_bacterium_RIFCSPLOWO2_02_FULL_39_110.OGW05939
Deltaproteobacteria_bacterium_GWB2_42_7.OGP22877
Deltaproteobacteria_bacterium_RIFOXYA2_FULL_42_10.OGQ75128
uncultured_bacterium.EKD26242
Candidatus_Aminicenantes_bacterium_RBG_19FT_COMBO_58_17.OGD35346
Candidatus_Aminicenantes_bacterium_RBG_13_62_12.OGD11581
Desulfobacterales_bacterium_C00003060.OEU81907
Deltaproteobacteria_bacterium_SG8_13.KPJ75571
Bdellovibrionales_bacterium_RIFOXYC1_FULL_37_79.OFZ47380
uncultured_bacterium.CAX83822
Alphaproteobacteria_bacterium_41_28.OJW52790
Bdellovibrionales_bacterium_RIFOXYC1_FULL_37_79.OFZ48281
Bdellovibrionales_bacterium_RIFOXYB1_FULL_39_21.OFZ36599
Gammaproteobacteria_bacterium_RIFCSPHIGHO2_12_38_15.OGT23808
Gammaproteobacteria_bacterium_RIFCSPHIGHO2_12_FULL_45_9.OGT68952
Gammaproteobacteria_bacterium_RIFCSPHIGHO2_12_FULL_38_14.OGT36811
Gammaproteobacteria_bacterium_GWE2_37_16.OGT07585
uncultured_bacterium.EKD71815
uncultured_bacterium.EKD56420
CBU_1786
Waddlia_chondrophila.WP_041941606
Waddlia_chondrophila_WSU_86_1044.ADI39113
Gammaproteobacteria_bacterium_RIFCSPHIGHO2_12_FULL_37_34.OGT41287
Gammaproteobacteria_bacterium_RIFCSPHIGHO2_12_FULL_42_10.OGT56656
Coxiella_sp_RIFCSPHIGHO2_12_FULL_44_14.OGO95524
Coxiella_sp_RIFCSPHIGHO2_12_FULL_42_15.OGO91697
Gammaproteobacteria_bacterium_RIFCSPHIGHO2_02_FULL_42_43.OGT26329
uncultured_bacterium.EKD45221
Candidatus_Melainabacteria_bacterium_HGW_Melainabacteria_1.PKL76428
Bdellovibrionales_bacterium_RIFOXYD1_FULL_44_7.OFZ85239
Kangiella_sp.PHS15863
Pseudohongiella_spirulinae.WP_058023090
Pseudohongiella_spirulinae.ALO45076
Candidatus_Magnetoglobus_multicellularis_str_Araruama.ETR69809
delta_proteobacterium_ML8_D.OPL11277
Syntrophobacterales_bacterium_CG_4_8_14_3_um_filter_58_8.PJC71571
Smithella_sp_ME_1.ESP62483
Deltaproteobacteria_bacterium_HGW_Deltaproteobacteria_9.PKN05474
delta_proteobacterium_MLMS_1.EAT01490
delta_proteobacterium_MLMS_1.WP_040871557
delta_proteobacterium_MLMS_1.WP_007291546
delta_proteobacterium_MLMS_1.WP_007292517
Deltaproteobacteria_bacterium_RIFOXYA12_FULL_58_15.OGQ78619
uncultured_Desulfofustis_sp_PB_SRB1.ESQ12598
Deltaproteobacteria_bacterium.PIE69443
Desulfobacter_postgatei.WP_004073943
Desulfotignum_balticum.WP_084489881
Desulfotignum_phosphitoxidans.WP_006968783
B
uncultured_bacterium
Spirochaetaceae_bacterium_4572_59
Bacteria
Waddlia
Verrucomicrobia_bacterium_RIFCSPHIGHO2_12_FULL_41_10
Waddlia_chondrophila_WSU_86-1044
Bacteria
Candidatus_Saganbacteria_bacterium_CG08_land_8_20_14_0_20_45_16
Candidatus_Melainabacteria_bacterium_HGW-Melainabacteria-1
Nitrospinae
Nitrospinae_bacterium_RIFCSPHIGHO2_12_FULL_39_42
Nitrospinae_bacterium_RIFCSPLOWO2_02_FULL_39_110
Elusimicrobia
Elusimicrobia_bacterium_GWF2_62_30
Elusimicrobia_bacterium_CG1_02_37_114
Elusimicrobia_bacterium_CG08_land_8_20_14_0_20_59_10
Bacteria
bacterium_BMS3Abin03
Bacteria
Candidatus_Firestonebacteria_bacterium_RIFOXYC2_FULL_39_67
Candidatus_Handelsmanbacteria_bacterium_RIFCSPLOWO2_12_FULL_64_10
Candidatus_Aminicenantes
Candidatus_Aminicenantes_bacterium_RBG_13_62_12
Candidatus_Aminicenantes_bacterium_RBG_19FT_COMBO_58_17
Alphaproteobacteria_bacterium_41-28
Bdellovibrionales
Bdellovibrionales_bacterium_RIFOXYB1_FULL_37_110
Bdellovibrionales_bacterium_RIFOXYC1_FULL_37_79
Bdellovibrionales_bacterium_RIFOXYC1_FULL_39_130
Bdellovibrionales_bacterium_RIFOXYD1_FULL_44_7
Bdellovibrionales_bacterium_RIFOXYD1_FULL_53_11
Leucothrix_mucor
Kangiella_sp
Gammaproteobacteria
Coxiella
Coxiella_burnetii_RSA_493
Coxiella
Coxiella_sp_RIFCSPHIGHO2_12_FULL_42_15
Coxiella_sp_RIFCSPHIGHO2_12_FULL_44_14
Proteobacteria
Pseudohongiella_spirulinae
Gammaproteobacteria
Gammaproteobacteria_bacterium_RIFCSPHIGHO2_12_38_15
Gammaproteobacteria_bacterium_RIFCSPHIGHO2_02_FULL_42_43
Gammaproteobacteria
Gammaproteobacteria_bacterium_RIFCSPHIGHO2_12_FULL_42_10
Gammaproteobacteria_bacterium_RIFCSPHIGHO2_12_FULL_37_34
Gammaproteobacteria_bacterium_RIFCSPHIGHO2_12_FULL_38_14
Gammaproteobacteria_bacterium_RIFCSPHIGHO2_12_FULL_45_9
Gammaproteobacteria_bacterium_GWE2_37_16
Gammaproteobacteria_bacterium_RIFCSPHIGHO2_01_FULL_42_8
Syntrophobacterales
Syntrophobacterales_bacterium_CG_4_8_14_3_um_filter_58_8
Syntrophaceae
Syntrophus_sp_RIFOXYC2_FULL_54_9
Smithella
Smithella_sp_ME-1
Smithella_sp_SCADC
Desulfobacterales
Desulfobacterales_bacterium_C00003060
uncultured_Desulfofustis_sp_PB-SRB1
Desulfobacter_postgatei
Deltaproteobacteria
Desulfobacteraceae
Desulfobacteraceae
Candidatus_Magnetoglobus_multicellularis_str_Araruama
Desulfobacteraceae_bacterium_4572_130
Desulfobacteraceae_bacterium_A6
Desulfotignum
Desulfotignum_phosphitoxidans
Desulfotignum_balticum
Deltaproteobacteria_bacterium_RIFOXYA12_FULL_58_15
Deltaproteobacteria_bacterium_RIFOXYA2_FULL_42_10
Deltaproteobacteria_bacterium_RBG_19FT_COMBO_43_11
Deltaproteobacteria_bacterium_GWA2_43_19
Deltaproteobacteria_bacterium_HGW-Deltaproteobacteria-19
Deltaproteobacteria_bacterium_GWB2_42_7
delta_proteobacterium_ML8_D
Deltaproteobacteria
Deltaproteobacteria_bacterium_RBG_16_44_11
Deltaproteobacteria_bacterium_RBG_16_49_23
Deltaproteobacteria_bacterium_RIFCSPHIGHO2_02_FULL_40_28
delta_proteobacterium_MLMS-1
Deltaproteobacteria_bacterium_RIFCSPLOWO2_02_FULL_46_8
Deltaproteobacteria_bacterium_SG8_13
Deltaproteobacteria_bacterium
Deltaproteobacteria_bacterium_HGW-Deltaproteobacteria-9

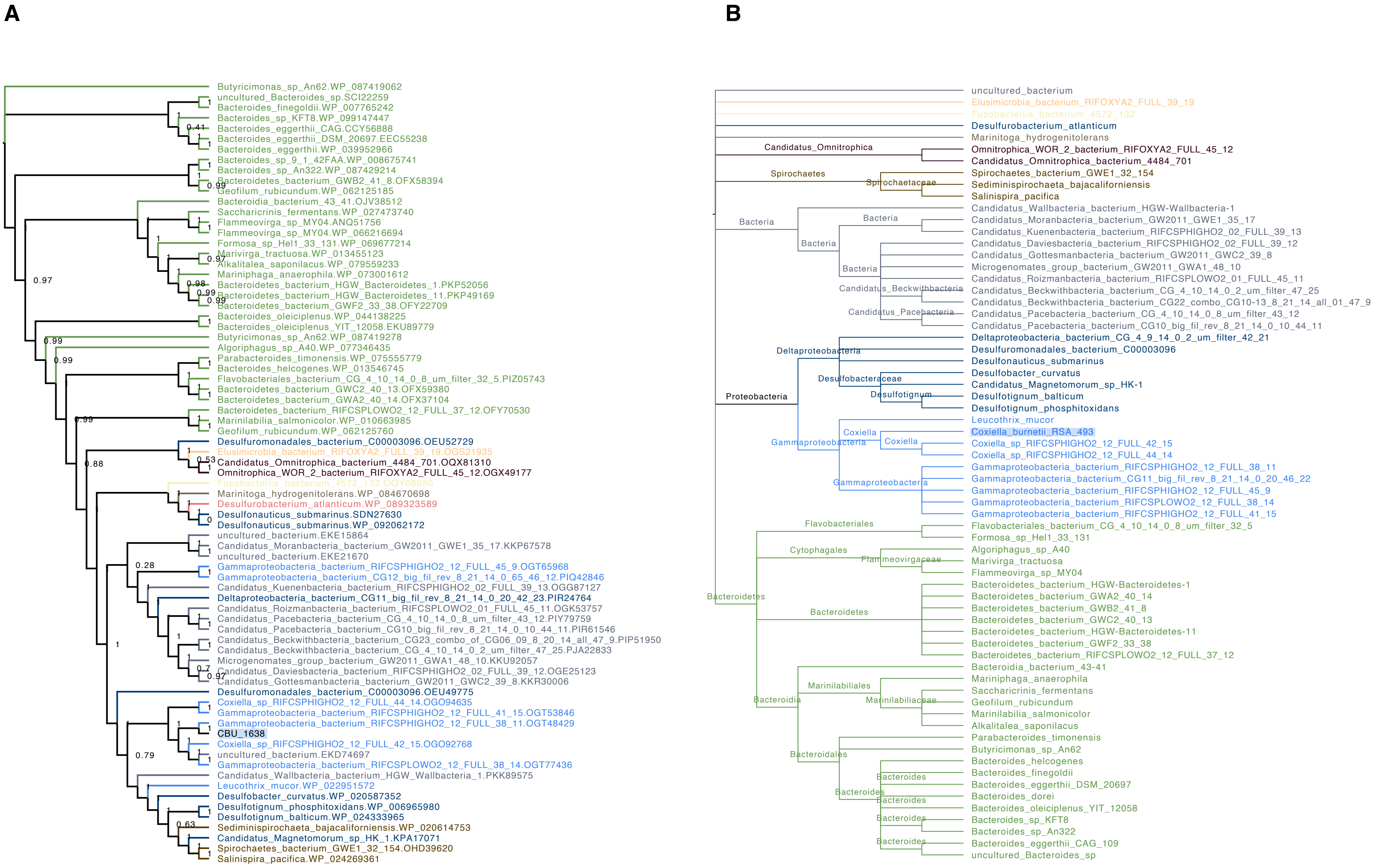
A
Butyricimonas_sp_An62.WP_087419062
uncultured_Bacteroides_sp.SCI22259
Bacteroides_finegoldii.WP_007765242
Bacteroides_sp_KFT8.WP_099147447
Bacteroides_eggerthii_CAG.CCY56888
Bacteroides_eggerthii_DSM_20697.EEC55238
Bacteroides_eggerthii.WP_039952966
Bacteroides_sp_9_1_42FAA.WP_008675741
Bacteroides_sp_An322.WP_087429214
Bacteroidetes_bacterium_GWB2_41_8.OFX58394
Geofilum_rubicundum.WP_062125185
Bacteroidia_bacterium_43_41.OJV38512
Saccharicrinis_fermentans.WP_027473740
Flammeovirga_sp_MY04.ANQ51756
Flammeovirga_sp_MY04.WP_066216694
Formosa_sp_Hel1_33_131.WP_069677214
Marivirga_tractuosa.WP_013455123
Alkalitalea_saponilacus.WP_079559233
Mariniphaga_anaerophila.WP_073001612
Bacteroidetes_bacterium_HGW_Bacteroidetes_1.PKP52056
Bacteroidetes_bacterium_HGW_Bacteroidetes_11.PKP49169
Bacteroidetes_bacterium_GWF2_33_38.OFY22709
Bacteroides_oleiciplenus.WP_044138225
Bacteroides_oleiciplenus_YIT_12058.EKU89779
Butyricimonas_sp_An62.WP_087419278
Algoriphagus_sp_A40.WP_077346435
Parabacteroides_timonensis.WP_075555779
Bacteroides_helcogenes.WP_013546745
Flavobacteriales_bacterium_CG_4_10_14_0_8_um_filter_32_5.PIZ05743
Bacteroidetes_bacterium_GWC2_40_13.OFX59380
Bacteroidetes_bacterium_GWA2_40_14.OFX37104
Bacteroidetes_bacterium_RIFCSPLOWO2_12_FULL_37_12.OFY70530
Marinilabilia_salmonicolor.WP_010663985
Geofilum_rubicundum.WP_062125760
Desulfuromonadales_bacterium_C00003096.OEU52729
Elusimicrobia_bacterium_RIFOXYA2_FULL_39_19.OGS21935
Candidatus_Omnitrophica_bacterium_4484_701.OQX81310
Omnitrophica_WOR_2_bacterium_RIFOXYA2_FULL_45_12.OGX49177
Fusobacteria_bacterium_4572_132.OQY08680
Marinitoga_hydrogenitolerans.WP_084670698
Desulfurobacterium_atlanticum.WP_089323589
Desulfonauticus_submarinus.SDN27630
Desulfonauticus_submarinus.WP_092062172
uncultured_bacterium.EKE15864
Candidatus_Moranbacteria_bacterium_GW2011_GWE1_35_17.KKP67578
uncultured_bacterium.EKE21670
Gammaproteobacteria_bacterium_RIFCSPHIGHO2_12_FULL_45_9.OGT65968
Gammaproteobacteria_bacterium_CG12_big_fil_rev_8_21_14_0_65_46_12.PIQ42846
Candidatus_Kuenenbacteria_bacterium_RIFCSPHIGHO2_02_FULL_39_13.OGG87127
Deltaproteobacteria_bacterium_CG11_big_fil_rev_8_21_14_0_20_42_23.PIR24764
Candidatus_Roizmanbacteria_bacterium_RIFCSPLOWO2_01_FULL_45_11.OGK53757
Candidatus_Pacebacteria_bacterium_CG_4_10_14_0_8_um_filter_43_12.PIY79759
Candidatus_Pacebacteria_bacterium_CG10_big_fil_rev_8_21_14_0_10_44_11.PIR61546
Candidatus_Beckwithbacteria_bacterium_CG23_combo_of_CG06_09_8_20_14_all_47_9.PIP51950
Candidatus_Beckwithbacteria_bacterium_CG_4_10_14_0_2_um_filter_47_25.PJA22833
Microgenomates_group_bacterium_GW2011_GWA1_48_10.KKU92057
Candidatus_Daviesbacteria_bacterium_RIFCSPHIGHO2_02_FULL_39_12.OGE25123
Candidatus_Gottesmanbacteria_bacterium_GW2011_GWC2_39_8.KKR30006
Desulfuromonadales_bacterium_C00003096.OEU49775
Coxiella_sp_RIFCSPHIGHO2_12_FULL_44_14.OGO94635
Gammaproteobacteria_bacterium_RIFCSPHIGHO2_12_FULL_41_15.OGT53846
Gammaproteobacteria_bacterium_RIFCSPHIGHO2_12_FULL_38_11.OGT48429
CBU_1638
Coxiella_sp_RIFCSPHIGHO2_12_FULL_42_15.OGO92768
uncultured_bacterium.EKD74697
Gammaproteobacteria_bacterium_RIFCSPLOWO2_12_FULL_38_14.OGT77436
Candidatus_Wallbacteria_bacterium_HGW_Wallbacteria_1.PKK89575
Leucothrix_mucor.WP_022951572
Desulfobacter_curvatus.WP_020587352
Desulfotignum_phosphitoxidans.WP_006965980
Desulfotignum_balticum.WP_024333965
Sediminispirochaeta_bajacaliforniensis.WP_020614753
Candidatus_Magnetomorum_sp_HK_1.KPA17071
Spirochaetes_bacterium_GWE1_32_154.OHD39620
Salinispira_pacifica.WP_024269361
B
uncultured_bacterium
Elusimicrobia_bacterium_RIFOXYA2_FULL_39_19
Fusobacteria_bacterium_4572_132
Desulfurobacterium_atlanticum
Marinitoga_hydrogenitolerans
Candidatus_Omnitrophica
Omnitrophica_WOR_2_bacterium_RIFOXYA2_FULL_45_12
Candidatus_Omnitrophica_bacterium_4484_701
Spirochaetes
Spirochaetaceae
Spirochaetes_bacterium_GWE1_32_154
Sediminispirochaeta_bajacaliforniensis
Salinispira_pacifica
Bacteria
Candidatus_Wallbacteria_bacterium_HGW-Wallbacteria-1
Candidatus_Moranbacteria_bacterium_GW2011_GWE1_35_17
Candidatus_Kuenenbacteria_bacterium_RIFCSPHIGHO2_02_FULL_39_13
Candidatus_Daviesbacteria_bacterium_RIFCSPHIGHO2_02_FULL_39_12
Candidatus_Gottesmanbacteria_bacterium_GW2011_GWC2_39_8
Microgenomates_group_bacterium_GW2011_GWA1_48_10
Candidatus_Roizmanbacteria_bacterium_RIFCSPLOWO2_01_FULL_45_11
Candidatus_Beckwithbacteria
Candidatus_Beckwithbacteria_bacterium_CG_4_10_14_0_2_um_filter_47_25
Candidatus_Beckwithbacteria_bacterium_CG22_combo_CG10-13_8_21_14_all_01_47_9
Candidatus_Pacebacteria
Candidatus_Pacebacteria_bacterium_CG_4_10_14_0_8_um_filter_43_12
Candidatus_Pacebacteria_bacterium_CG10_big_fil_rev_8_21_14_0_10_44_11
Proteobacteria
Deltaproteobacteria
Deltaproteobacteria_bacterium_CG_4_9_14_0_2_um_filter_42_21
Desulfuromonadales_bacterium_C00003096
Desulfonauticus_submarinus
Desulfobacteraceae
Desulfobacter_curvatus
Candidatus_Magnetomorum_sp_HK-1
Desulfotignum
Desulfotignum_balticum
Desulfotignum_phosphitoxidans
Gammaproteobacteria
Leucothrix_mucor
Coxiella
Coxiella_burnetii_RSA_493
Coxiella_sp_RIFCSPHIGHO2_12_FULL_42_15
Coxiella_sp_RIFCSPHIGHO2_12_FULL_44_14
Gammaproteobacteria_bacterium_RIFCSPHIGHO2_12_FULL_38_11
Gammaproteobacteria_bacterium_CG11_big_fil_rev_8_21_14_0_20_46_22
Gammaproteobacteria_bacterium_RIFCSPHIGHO2_12_FULL_45_9
Gammaproteobacteria_bacterium_RIFCSPLOWO2_12_FULL_38_14
Gammaproteobacteria_bacterium_RIFCSPHIGHO2_12_FULL_41_15
Bacteroidetes
Flavobacteriales
Flavobacteriales_bacterium_CG_4_10_14_0_8_um_filter_32_5
Formosa_sp_Hel1_33_131
Cytophagales
Flammeovirgaceae
Algoriphagus_sp_A40
Marivirga_tractuosa
Flammeovirga_sp_MY04
Bacteroidetes_bacterium_HGW-Bacteroidetes-1
Bacteroidetes_bacterium_GWA2_40_14
Bacteroidetes_bacterium_GWB2_41_8
Bacteroidetes_bacterium_GWC2_40_13
Bacteroidetes_bacterium_HGW-Bacteroidetes-11
Bacteroidetes_bacterium_GWF2_33_38
Bacteroidetes_bacterium_RIFCSPLOWO2_12_FULL_37_12
Bacteroidia
Bacteroidia_bacterium_43-41
Marinilabiliales
Marinilabiliaceae
Mariniphaga_anaerophila
Saccharicrinis_fermentans
Geofilum_rubicundum
Marinilabilia_salmonicolor
Alkalitalea_saponilacus
Bacteroidales
Parabacteroides_timonensis
Butyricimonas_sp_An62
Bacteroides
Bacteroides_helcogenes
Bacteroides_finegoldii
Bacteroides_eggerthii_DSM_20697
Bacteroides_dorei
Bacteroides_oleiciplenus_YIT_12058
Bacteroides_sp_KFT8
Bacteroides_sp_An322
Bacteroides_eggerthii_CAG_109
uncultured_Bacteroides_sp

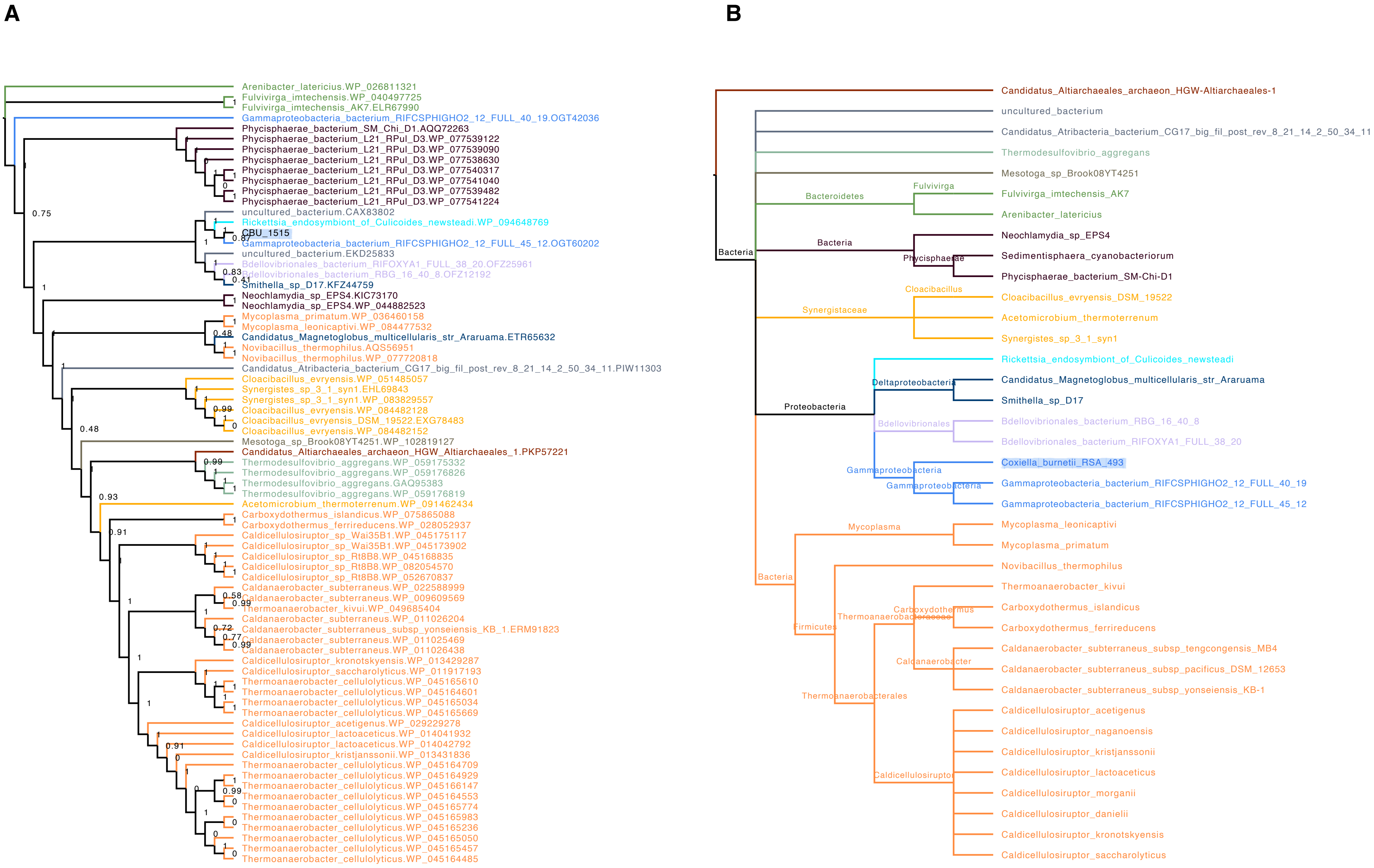
A
Arenibacter_latericius.WP_026811321
Fulvivirga_imtechensis.WP_040497725
Fulvivirga_imtechensis_AK7.ELR67990
Gammaproteobacteria_bacterium_RIFCSPHIGHO2_12_FULL_40_19.OGT42036
Phycisphaerae_bacterium_SM_Chi_D1.AQQ72263
Phycisphaerae_bacterium_L21_RPul_D3.WP_077539122
Phycisphaerae_bacterium_L21_RPul_D3.WP_077539090
Phycisphaerae_bacterium_L21_RPul_D3.WP_077538630
Phycisphaerae_bacterium_L21_RPul_D3.WP_077540317
Phycisphaerae_bacterium_L21_RPul_D3.WP_077541040
Phycisphaerae_bacterium_L21_RPul_D3.WP_077539482
Phycisphaerae_bacterium_L21_RPul_D3.WP_077541224
uncultured_bacterium.CAX83802
Rickettsia_endosymbiont_of_Culicoides_newsteadi.WP_094648769
CBU_1515
Gammaproteobacteria_bacterium_RIFCSPHIGHO2_12_FULL_45_12.OGT60202
uncultured_bacterium.EKD25833
Bdellovibrionales_bacterium_RIFOXYA1_FULL_38_20.OFZ25961
Bdellovibrionales_bacterium_RBG_16_40_8.OFZ12192
Smithella_sp_D17.KFZ44759
Neochlamydia_sp_EPS4.KIC73170
Neochlamydia_sp_EPS4.WP_044882523
Mycoplasma_primatum.WP_036460158
Mycoplasma_leonicaptivi.WP_084477532
Candidatus_Magnetoglobus_multicellularis_str_Araruama.ETR65632
Novibacillus_thermophilus.AQS56951
Novibacillus_thermophilus.WP_077720818
Candidatus_Atribacteria_bacterium_CG17_big_fil_post_rev_8_21_14_2_50_34_11.PIW11303
Cloacibacillus_evryensis.WP_051485057
Synergistes_sp_3_1_syn1.EHL69843
Synergistes_sp_3_1_syn1.WP_083829557
Cloacibacillus_evryensis.WP_084482128
Cloacibacillus_evryensis_DSM_19522.EXG78483
Cloacibacillus_evryensis.WP_084482152
Mesotoga_sp_Brook08YT4251.WP_102819127
Candidatus_Altiarchaeales_archaeon_HGW_Altiarchaeales_1.PKP57221
Thermodesulfovibrio_aggregans.WP_059175332
Thermodesulfovibrio_aggregans.WP_059176826
Thermodesulfovibrio_aggregans.GAQ95383
Thermodesulfovibrio_aggregans.WP_059176819
Acetomicrobium_thermoterrenum.WP_091462434
Carboxydothermus_islandicus.WP_075865088
Carboxydothermus_ferrireducens.WP_028052937
Caldicellulosiruptor_sp_Wai35B1.WP_045175117
Caldicellulosiruptor_sp_Wai35B1.WP_045173902
Caldicellulosiruptor_sp_Rt8B8.WP_045168835
Caldicellulosiruptor_sp_Rt8B8.WP_082054570
Caldicellulosiruptor_sp_Rt8B8.WP_052670837
Caldanaerobacter_subterraneus.WP_022588999
Caldanaerobacter_subterraneus.WP_009609569
Thermoanaerobacter_kivui.WP_049685404
Caldanaerobacter_subterraneus.WP_011026204
Caldanaerobacter_subterraneus_subsp_yonseiensis_KB_1.ERM91823
Caldanaerobacter_subterraneus.WP_011025469
Caldanaerobacter_subterraneus.WP_011026438
Caldicellulosiruptor_kronotskyensis.WP_013429287
Caldicellulosiruptor_saccharolyticus.WP_011917193
Thermoanaerobacter_cellulolyticus.WP_045165610
Thermoanaerobacter_cellulolyticus.WP_045164601
Thermoanaerobacter_cellulolyticus.WP_045165034
Thermoanaerobacter_cellulolyticus.WP_045165669
Caldicellulosiruptor_acetigenus.WP_029229278
Caldicellulosiruptor_lactoaceticus.WP_014041932
Caldicellulosiruptor_lactoaceticus.WP_014042792
Caldicellulosiruptor_kristjanssonii.WP_013431836
Thermoanaerobacter_cellulolyticus.WP_045164709
Thermoanaerobacter_cellulolyticus.WP_045164929
Thermoanaerobacter_cellulolyticus.WP_045166147
Thermoanaerobacter_cellulolyticus.WP_045164553
Thermoanaerobacter_cellulolyticus.WP_045165774
Thermoanaerobacter_cellulolyticus.WP_045165983
Thermoanaerobacter_cellulolyticus.WP_045165236
Thermoanaerobacter_cellulolyticus.WP_045165050
Thermoanaerobacter_cellulolyticus.WP_045165457
Thermoanaerobacter_cellulolyticus.WP_045164485
B
Candidatus_Altiarchaeales_archaeon_HGW-Altiarchaeales-1
uncultured_bacterium
Candidatus_Atribacteria_bacterium_CG17_big_fil_post_rev_8_21_14_2_50_34_11
Thermodesulfovibrio_aggregans
Mesotoga_sp_Brook08YT4251
Bacteroidetes
Fulvivirga
Fulvivirga_imtechensis_AK7
Arenibacter_latericius
Bacteria
Phycisphaerae
Neochlamydia_sp_EPS4
Sedimentisphaera_cyanobacteriorum
Phycisphaerae_bacterium_SM-Chi-D1
Synergistaceae
Cloacibacillus
Cloacibacillus_evryensis_DSM_19522
Acetomicrobium_thermoterrenum
Synergistes_sp_3_1_syn1
Rickettsia_endosymbiont_of_Culicoides_newsteadi
Deltaproteobacteria
Candidatus_Magnetoglobus_multicellularis_str_Araruama
Smithella_sp_D17
Proteobacteria
Bdellovibrionales
Bdellovibrionales_bacterium_RBG_16_40_8
Bdellovibrionales_bacterium_RIFOXYA1_FULL_38_20
Gammaproteobacteria
Coxiella_burnetii_RSA_493
Gammaproteobacteria_bacterium_RIFCSPHIGHO2_12_FULL_40_19
Gammaproteobacteria_bacterium_RIFCSPHIGHO2_12_FULL_45_12
Mycoplasma
Mycoplasma_leonicaptivi
Mycoplasma_primatum
Novibacillus_thermophilus
Firmicutes
Thermoanaerobacteraceae
Carboxydothermus
Thermoanaerobacter_kivui
Carboxydothermus_islandicus
Carboxydothermus_ferrireducens
Caldanaerobacter
Caldanaerobacter_subterraneus_subsp_tengcongensis_MB4
Caldanaerobacter_subterraneus_subsp_pacificus_DSM_12653
Caldanaerobacter_subterraneus_subsp_yonseiensis_KB-1
Thermoanaerobacterales
Caldicellulosiruptor
Caldicellulosiruptor_acetigenus
Caldicellulosiruptor_naganoensis
Caldicellulosiruptor_kristjanssonii
Caldicellulosiruptor_lactoaceticus
Caldicellulosiruptor_morganii
Caldicellulosiruptor_danielii
Caldicellulosiruptor_kronotskyensis
Caldicellulosiruptor_saccharolyticus

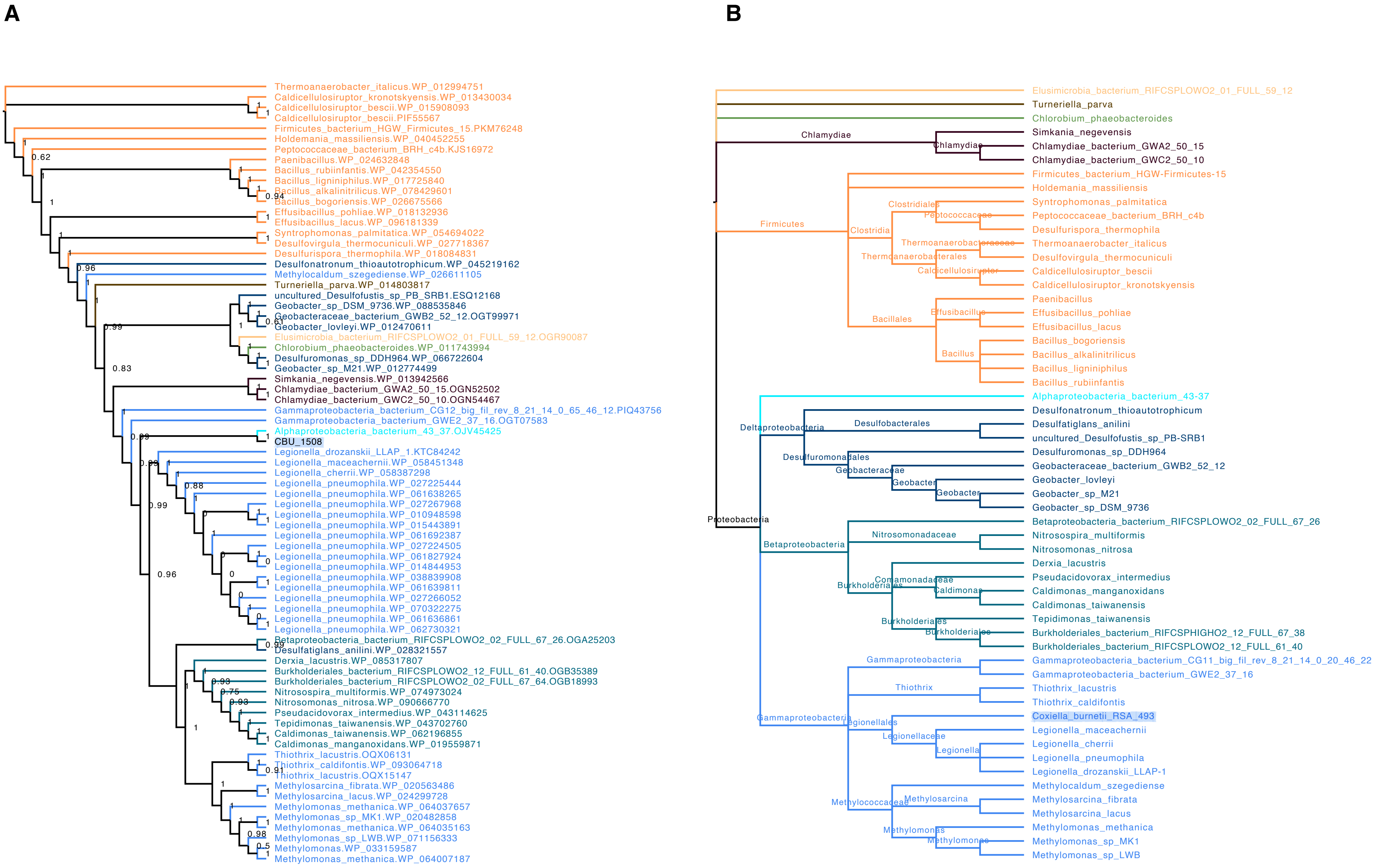
A
Thermoanaerobacter_italicus.WP_012994751
Caldicellulosiruptor_kronotskyensis.WP_013430034
Caldicellulosiruptor_bescii.WP_015908093
Caldicellulosiruptor_bescii.PIF55567
Firmicutes_bacterium_HGW_Firmicutes_15.PKM76248
Holdemania_massiliensis.WP_040452265
Peptococcaceae_bacterium_BRH_c4b.KJS16972
Paenibacillus.WP_024632848
Bacillus_rubiinfantis.WP_042354550
Bacillus_ligniniphilus.WP_017725840
Bacillus_alkalinitrilicus.WP_078429601
Bacillus_bogoriensis.WP_026675566
Effusibacillus_pohliae.WP_018132936
Effusibacillus_lacus.WP_096181339
Syntrophomonas_palmitatica.WP_054694022
Desulfovirgula_thermocuniculi.WP_027718367
Desulfurispora_thermophila.WP_018084831
Desulfonatronum_thioautotrophicum.WP_045219162
Methylocaldum_szegediense.WP_026611105
Turneriella_parva.WP_014803817
uncultured_Desulfofustis_sp_PB_SRB1.ESQ12168
Geobacter_sp_DSM_9736.WP_088535846
Geobacteraceae_bacterium_GWB2_52_12.OGT99971
Geobacter_lovleyi.WP_012470611
Elusimicrobia_bacterium_RIFCSPLOWO2_01_FULL_59_12.OGR90087
Chlorobium_phaeobacteroides.WP_011743994
Desulfuromonas_sp_DDH964.WP_066722604
Geobacter_sp_M21.WP_012774499
Simkania_negevensis.WP_013942566
Chlamydiae_bacterium_GWA2_50_15.OGN52502
Chlamydiae_bacterium_GWC2_50_10.OGN54467
Gammaproteobacteria_bacterium_CG12_big_fil_rev_8_21_14_0_65_46_12.PIQ43756
Gammaproteobacteria_bacterium_GWE2_37_16.OGT07583
Alphaproteobacteria_bacterium_43_37.OJV45425
CBU_1508
Legionella_drozanskii_LLAP_1.KTC84242
Legionella_maceachernii.WP_058451348
Legionella_cherrii.WP_058387298
Legionella_pneumophila.WP_027225444
Legionella_pneumophila.WP_061638265
Legionella_pneumophila.WP_027267968
Legionella_pneumophila.WP_010948598
Legionella_pneumophila.WP_015443891
Legionella_pneumophila.WP_061692387
Legionella_pneumophila.WP_027224505
Legionella_pneumophila.WP_061827924
Legionella_pneumophila.WP_014844953
Legionella_pneumophila.WP_038839908
Legionella_pneumophila.WP_061639811
Legionella_pneumophila.WP_027266052
Legionella_pneumophila.WP_070322275
Legionella_pneumophila.WP_061636861
Legionella_pneumophila.WP_062730321
Betaproteobacteria_bacterium_RIFCSPLOWO2_02_FULL_67_26.OGA25203
Desulfatiglans_anilini.WP_028321557
Derxia_lacustris.WP_085317807
Burkholderiales_bacterium_RIFCSPLOWO2_12_FULL_61_40.OGB35389
Burkholderiales_bacterium_RIFCSPLOWO2_02_FULL_67_64.OGB18993
Nitrosospira_multiformis.WP_074973024
Nitrosomonas_nitrosa.WP_090666770
Pseudacidovorax_intermedius.WP_043114625
Tepidimonas_taiwanensis.WP_043702760
Caldimonas_taiwanensis.WP_062196855
Caldimonas_manganoxidans.WP_019559871
Thiothrix_lacustris.OQX06131
Thiothrix_caldifontis.WP_093064718
Thiothrix_lacustris.OQX15147
Methylosarcina_fibrata.WP_020563486
Methylosarcina_lacus.WP_024299728
Methylomonas_methanica.WP_064037657
Methylomonas_sp_MK1.WP_020482858
Methylomonas_methanica.WP_064035163
Methylomonas_sp_LWB.WP_071156333
Methylomonas.WP_033159587
Methylomonas_methanica.WP_064007187
B
Elusimicrobia_bacterium_RIFCSPLOWO2_01_FULL_59_12
Turneriella_parva
Chlorobium_phaeobacteroides
Chlamydiae
Chlamydiae
Simkania_negevensis
Chlamydiae_bacterium_GWA2_50_15
Chlamydiae_bacterium_GWC2_50_10
Firmicutes
Clostridia
Clostridiales
Peptococcaceae
Thermoanaerobacterales
Thermoanaerobacteraceae
Caldicellulosiruptor
Bacillales
Effusibacillus
Bacillus
Firmicutes_bacterium_HGW-Firmicutes-15
Holdemania_massiliensis
Syntrophomonas_palmitatica
Peptococcaceae_bacterium_BRH_c4b
Desulfurispora_thermophila
Thermoanaerobacter_italicus
Desulfovirgula_thermocuniculi
Caldicellulosiruptor_bescii
Caldicellulosiruptor_kronotskyensis
Paenibacillus
Effusibacillus_pohliae
Effusibacillus_lacus
Bacillus_bogoriensis
Bacillus_alkalinitrilicus
Bacillus_ligniniphilus
Bacillus_rubiinfantis
Proteobacteria
Deltaproteobacteria
Desulfobacterales
Desulfuromonadales
Geobacteraceae
Geobacter
Geobacter
Alphaproteobacteria_bacterium_43-37
Desulfonatronum_thioautotrophicum
Desulfatiglans_anilini
uncultured_Desulfofustis_sp_PB-SRB1
Desulfuromonas_sp_DDH964
Geobacteraceae_bacterium_GWB2_52_12
Geobacter_lovleyi
Geobacter_sp_M21
Geobacter_sp_DSM_9736
Betaproteobacteria
Nitrosomonadaceae
Burkholderiales
Comamonadaceae
Caldimonas
Burkholderiales
Burkholderiales
Betaproteobacteria_bacterium_RIFCSPLOWO2_02_FULL_67_26
Nitrosospira_multiformis
Nitrosomonas_nitrosa
Derxia_lacustris
Pseudacidovorax_intermedius
Caldimonas_manganoxidans
Caldimonas_taiwanensis
Tepidimonas_taiwanensis
Burkholderiales_bacterium_RIFCSPHIGHO2_12_FULL_67_38
Burkholderiales_bacterium_RIFCSPLOWO2_12_FULL_61_40
Gammaproteobacteria
Gammaproteobacteria
Thiothrix
Legionellales
Legionellaceae
Legionella
Methylococcaceae
Methylosarcina
Methylomonas
Methylomonas
Gammaproteobacteria_bacterium_CG11_big_fil_rev_8_21_14_0_20_46_22
Gammaproteobacteria_bacterium_GWE2_37_16
Thiothrix_lacustris
Thiothrix_caldifontis
Coxiella_burnetii_RSA_493
Legionella_maceachernii
Legionella_cherrii
Legionella_pneumophila
Legionella_drozanskii_LLAP-1
Methylocaldum_szegediense
Methylosarcina_fibrata
Methylosarcina_lacus
Methylomonas_methanica
Methylomonas_sp_MK1
Methylomonas_sp_LWB

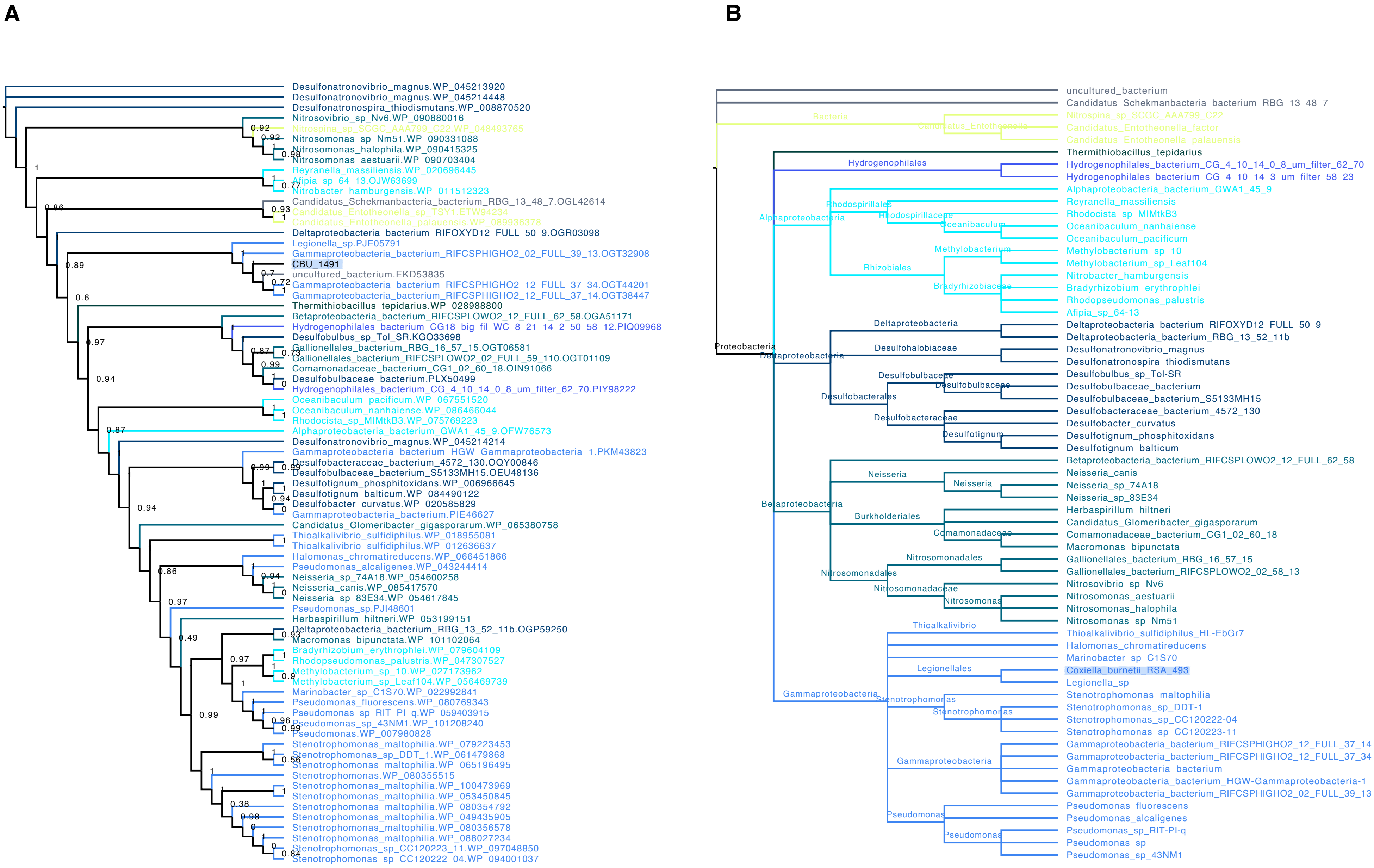
A
Desulfonatronovibrio_magnus.WP_045213920
Desulfonatronovibrio_magnus.WP_045214448
Desulfonatronospira_thiodismutans.WP_008870520
Nitrosovibrio_sp_Nv6.WP_090880016
Nitrospina_sp_SCGC_AAA799_C22.WP_048493765
Nitrosomonas_sp_Nm51.WP_090331088
Nitrosomonas_halophila.WP_090415325
Nitrosomonas_aestuarii.WP_090703404
Reyranella_massiliensis.WP_020696445
Afipia_sp_64_13.OJW63699
Nitrobacter_hamburgensis.WP_011512323
Candidatus_Schekmanbacteria_bacterium_RBG_13_48_7.OGL42614
Candidatus_Entotheonella_sp_TSY1.ETW94234
Candidatus_Entotheonella_palauensis.WP_089936378
Deltaproteobacteria_bacterium_RIFOXYD12_FULL_50_9.OGR03098
Legionella_sp.PJE05791
Gammaproteobacteria_bacterium_RIFCSPHIGHO2_02_FULL_39_13.OGT32908
CBU_1491
uncultured_bacterium.EKD53835
Gammaproteobacteria_bacterium_RIFCSPHIGHO2_12_FULL_37_34.OGT44201
Gammaproteobacteria_bacterium_RIFCSPHIGHO2_12_FULL_37_14.OGT38447
Thermithiobacillus_tepidarius.WP_028988800
Betaproteobacteria_bacterium_RIFCSPLOWO2_12_FULL_62_58.OGA51171
Hydrogenophilales_bacterium_CG18_big_fil_WC_8_21_14_2_50_58_12.PIQ09968
Desulfobulbus_sp_Tol_SR.KGO33698
Gallionellales_bacterium_RBG_16_57_15.OGT06581
Gallionellales_bacterium_RIFCSPLOWO2_02_FULL_59_110.OGT01109
Comamonadaceae_bacterium_CG1_02_60_18.OIN91066
Desulfobulbaceae_bacterium.PLX50499
Hydrogenophilales_bacterium_CG_4_10_14_0_8_um_filter_62_70.PIY98222
Oceanibaculum_pacificum.WP_067551520
Oceanibaculum_nanhaiense.WP_086466044
Rhodocista_sp_MIMtkB3.WP_075769223
Alphaproteobacteria_bacterium_GWA1_45_9.OFW76573
Desulfonatronovibrio_magnus.WP_045214214
Gammaproteobacteria_bacterium_HGW_Gammaproteobacteria_1.PKM43823
Desulfobacteraceae_bacterium_4572_130.OQY00846
Desulfobulbaceae_bacterium_S5133MH15.OEU48136
Desulfotignum_phosphitoxidans.WP_006966645
Desulfotignum_balticum.WP_084490122
Desulfobacter_curvatus.WP_020585829
Gammaproteobacteria_bacterium.PIE46627
Candidatus_Glomeribacter_gigasporarum.WP_065380758
Thioalkalivibrio_sulfidiphilus.WP_018955081
Thioalkalivibrio_sulfidiphilus.WP_012636637
Halomonas_chromatireducens.WP_066451866
Pseudomonas_alcaligenes.WP_043244414
Neisseria_sp_74A18.WP_054600258
Neisseria_canis.WP_085417570
Neisseria_sp_83E34.WP_054617845
Pseudomonas_sp.PJI48601
Herbaspirillum_hiltneri.WP_053199151
Deltaproteobacteria_bacterium_RBG_13_52_11b.OGP59250
Macromonas_bipunctata.WP_101102064
Bradyrhizobium_erythrophlei.WP_079604109
Rhodopseudomonas_palustris.WP_047307527
Methylobacterium_sp_10.WP_027173962
Methylobacterium_sp_Leaf104.WP_056469739
Marinobacter_sp_C1S70.WP_022992841
Pseudomonas_fluorescens.WP_080769343
Pseudomonas_sp_RIT_PI_q.WP_059403915
Pseudomonas_sp_43NM1.WP_101208240
Pseudomonas.WP_007980828
Stenotrophomonas_maltophilia.WP_079223453
Stenotrophomonas_sp_DDT_1.WP_061479868
Stenotrophomonas_maltophilia.WP_065196495
Stenotrophomonas.WP_080355515
Stenotrophomonas_maltophilia.WP_100473969
Stenotrophomonas_maltophilia.WP_053450845
Stenotrophomonas_maltophilia.WP_080354792
Stenotrophomonas_maltophilia.WP_049435905
Stenotrophomonas_maltophilia.WP_080356578
Stenotrophomonas_maltophilia.WP_088027234
Stenotrophomonas_sp_CC120223_11.WP_097048850
Stenotrophomonas_sp_CC120222_04.WP_094001037
B
uncultured_bacterium
Candidatus_Schekmanbacteria_bacterium_RBG_13_48_7
Bacteria
Candidatus_Entotheonella
Nitrospina_sp_SCGC_AAA799_C22
Candidatus_Entotheonella_factor
Candidatus_Entotheonella_palauensis
Thermithiobacillus_tepidarius
Hydrogenophilales
Hydrogenophilales_bacterium_CG_4_10_14_0_8_um_filter_62_70
Hydrogenophilales_bacterium_CG_4_10_14_3_um_filter_58_23
Alphaproteobacteria
Alphaproteobacteria_bacterium_GWA1_45_9
Rhodospirillales
Rhodospirillaceae
Reyranella_massiliensis
Rhodocista_sp_MIMtkB3
Oceanibaculum
Oceanibaculum_nanhaiense
Oceanibaculum_pacificum
Methylobacterium
Methylobacterium_sp_10
Methylobacterium_sp_Leaf104
Rhizobiales
Bradyrhizobiaceae
Nitrobacter_hamburgensis
Bradyrhizobium_erythrophlei
Rhodopseudomonas_palustris
Afipia_sp_64-13
Proteobacteria
Deltaproteobacteria
Deltaproteobacteria_bacterium_RIFOXYD12_FULL_50_9
Deltaproteobacteria_bacterium_RBG_13_52_11b
Desulfohalobiaceae
Desulfonatronovibrio_magnus
Desulfonatronospira_thiodismutans
Desulfobulbaceae
Desulfobulbus_sp_Tol-SR
Desulfobulbaceae_bacterium
Desulfobulbaceae_bacterium_S5133MH15
Desulfobacterales
Desulfobacteraceae
Desulfobacteraceae_bacterium_4572_130
Desulfobacter_curvatus
Desulfotignum
Desulfotignum_phosphitoxidans
Desulfotignum_balticum
Betaproteobacteria
Betaproteobacteria_bacterium_RIFCSPLOWO2_12_FULL_62_58
Neisseria
Neisseria_canis
Neisseria_sp_74A18
Neisseria_sp_83E34
Burkholderiales
Herbaspirillum_hiltneri
Candidatus_Glomeribacter_gigasporarum
Comamonadaceae
Comamonadaceae_bacterium_CG1_02_60_18
Macromonas_bipunctata
Nitrosomonadales
Gallionellales_bacterium_RBG_16_57_15
Gallionellales_bacterium_RIFCSPLOWO2_02_58_13
Nitrosomonadaceae
Nitrosovibrio_sp_Nv6
Nitrosomonas
Nitrosomonas_aestuarii
Nitrosomonas_halophila
Nitrosomonas_sp_Nm51
Thioalkalivibrio
Thioalkalivibrio_sulfidiphilus_HL-EbGr7
Halomonas_chromatireducens
Marinobacter_sp_C1S70
Legionellales
Coxiella_burnetii_RSA_493
Legionella_sp
Gammaproteobacteria
Stenotrophomonas
Stenotrophomonas_maltophilia
Stenotrophomonas_sp_DDT-1
Stenotrophomonas_sp_CC120222-04
Stenotrophomonas_sp_CC120223-11
Gammaproteobacteria_bacterium_RIFCSPHIGHO2_12_FULL_37_14
Gammaproteobacteria_bacterium_RIFCSPHIGHO2_12_FULL_37_34
Gammaproteobacteria_bacterium
Gammaproteobacteria_bacterium_HGW-Gammaproteobacteria-1
Gammaproteobacteria_bacterium_RIFCSPHIGHO2_02_FULL_39_13
Pseudomonas
Pseudomonas_fluorescens
Pseudomonas_alcaligenes
Pseudomonas_sp_RIT-PI-q
Pseudomonas_sp
Pseudomonas_sp_43NM1

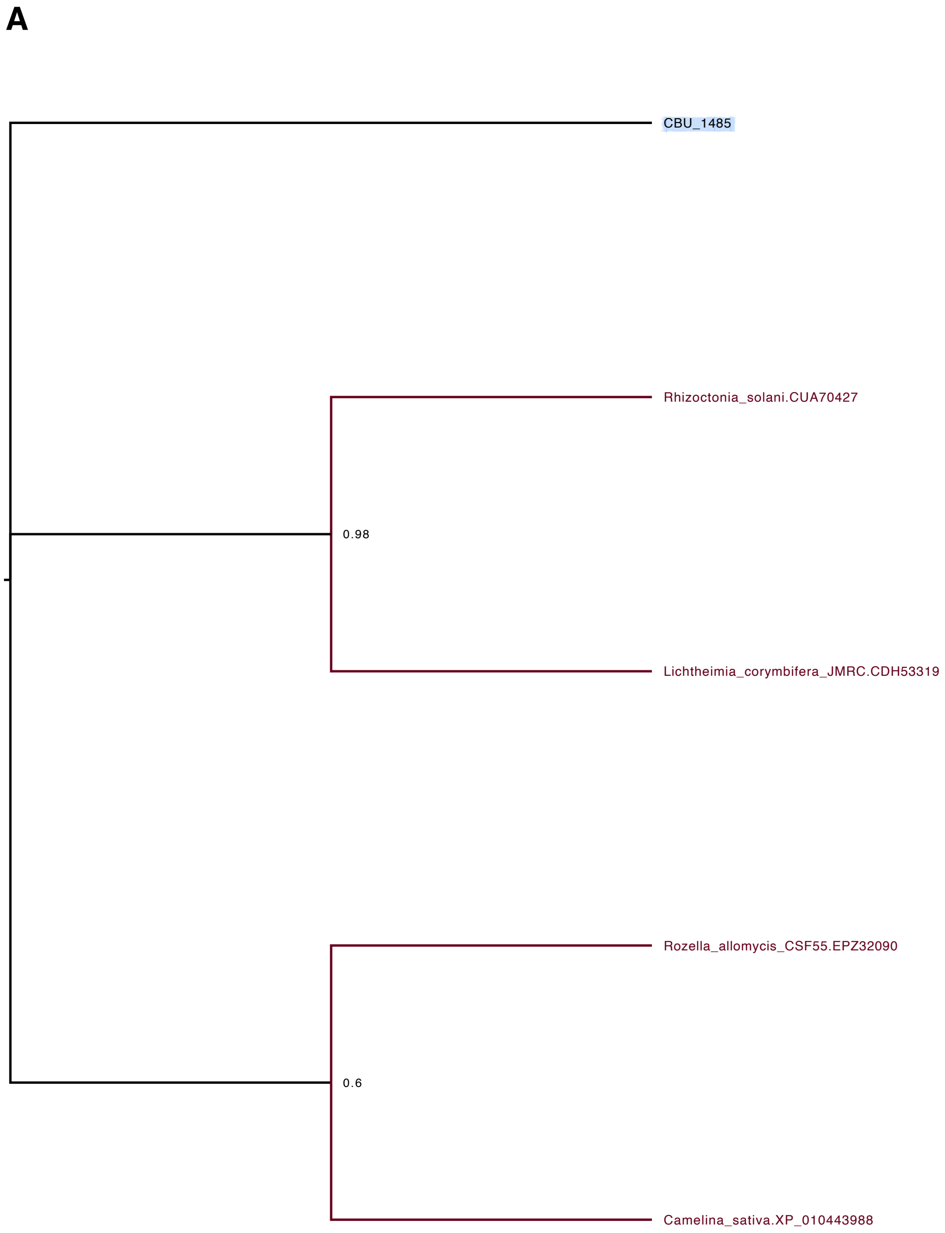
A
CBU_1485
Rhizoctonia_solani.CUA70427
0.98
Lichtheimia_corymbifera_JMRC.CDH53319
Rozella_allomycis_CSF55.EPZ32090
0.6
Camelina_sativa.XP_010443988

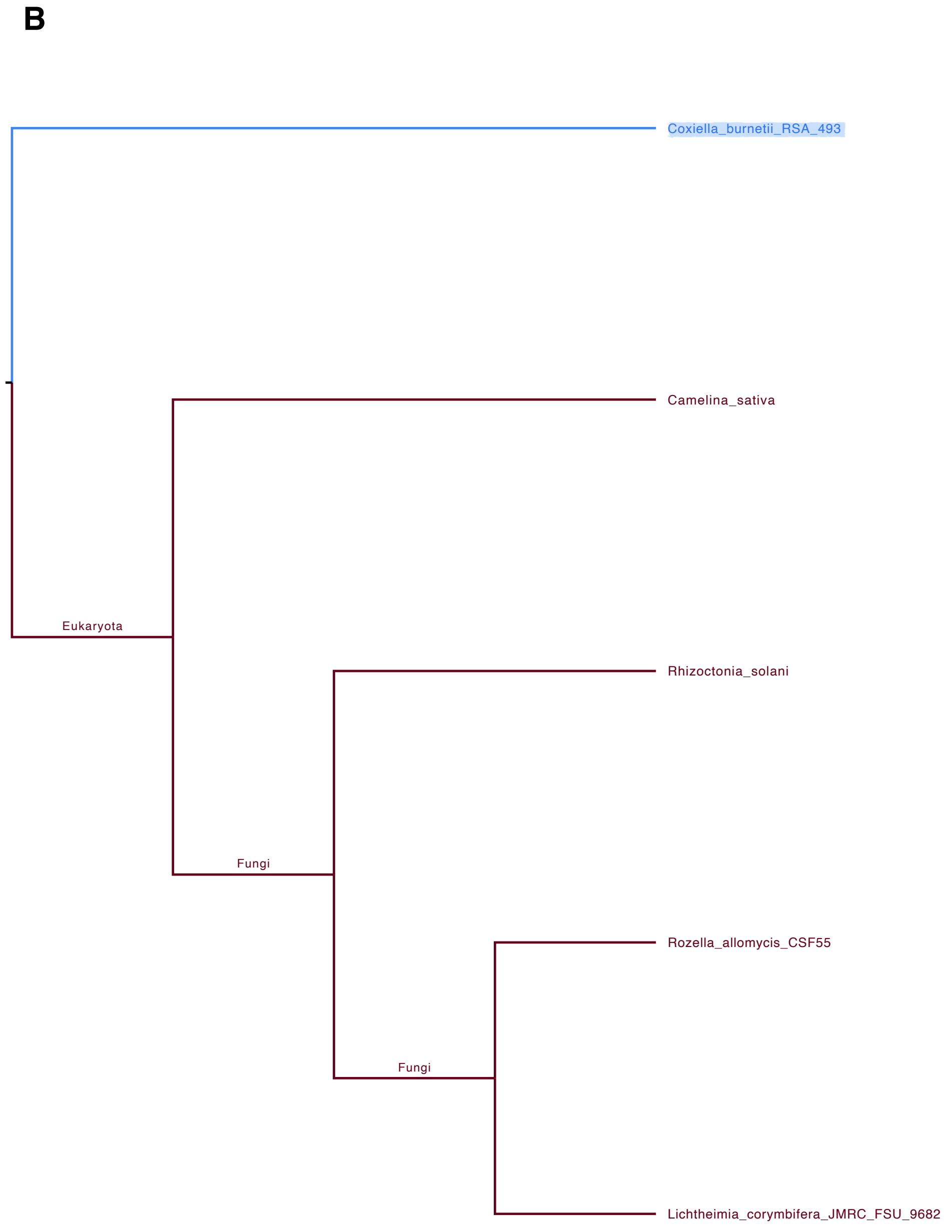
B
Coxiella_burnetii_RSA_493
Camelina_sativa
Eukaryota
Rhizoctonia_solani
Fungi
Rozella_allomycis_CSF55
Fungi
Lichtheimia_corymbifera_JMRC_FSU_9682

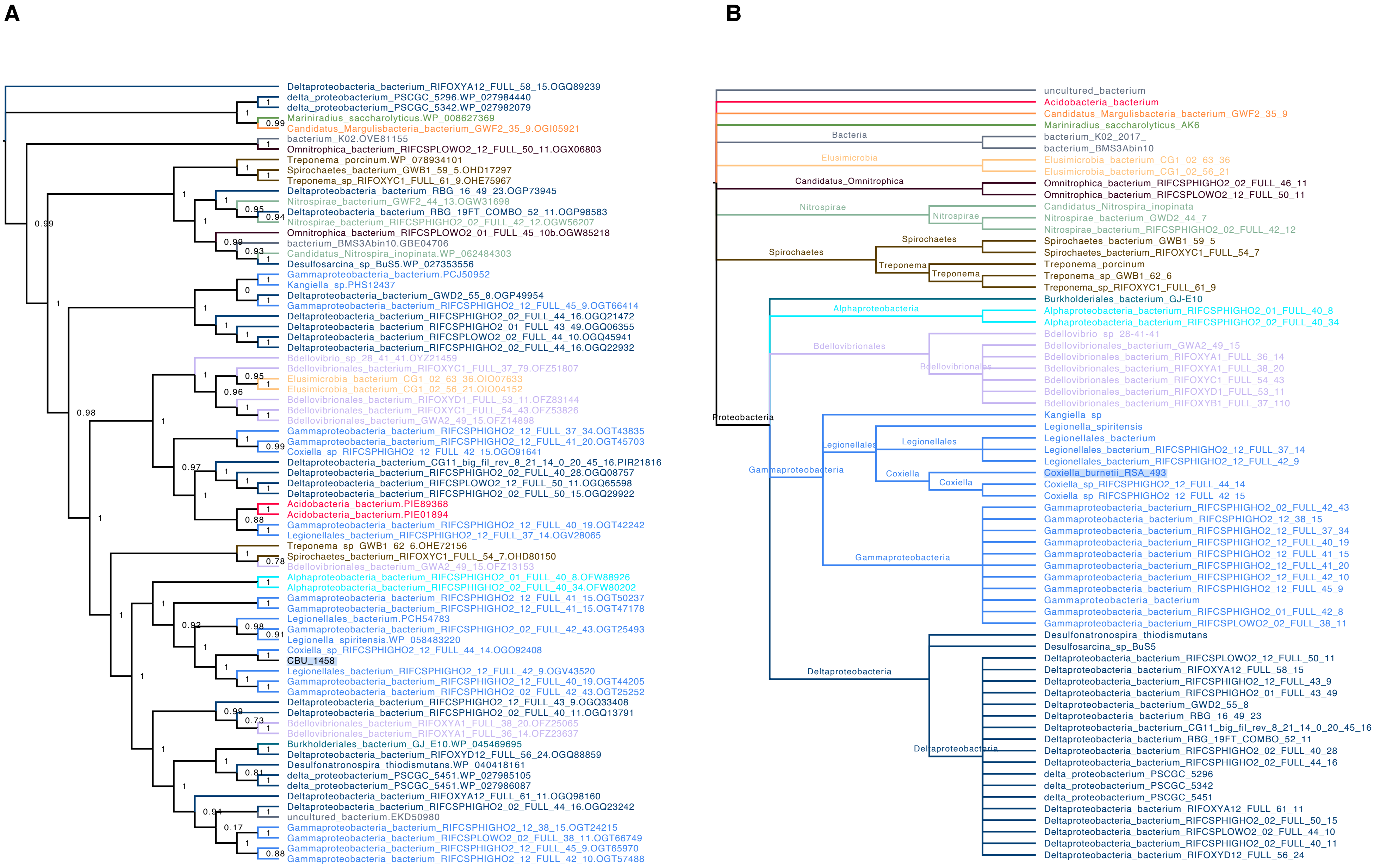
A
Deltaproteobacteria_bacterium_RIFOXYA12_FULL_58_15.OGQ89239
delta_proteobacterium_PSCGC_5296.WP_027984440
delta_proteobacterium_PSCGC_5342.WP_027982079
Mariniradius_saccharolyticus.WP_008627369
Candidatus_Margulisbacteria_bacterium_GWF2_35_9.OGI05921
bacterium_K02.OVE81155
Omnitrophica_bacterium_RIFCSPLOWO2_12_FULL_50_11.OGX06803
Treponema_porcinum.WP_078934101
Spirochaetes_bacterium_GWB1_59_5.OHD17297
Treponema_sp_RIFOXYC1_FULL_61_9.OHE75967
Deltaproteobacteria_bacterium_RBG_16_49_23.OGP73945
Nitrospirae_bacterium_GWF2_44_13.OGW31698
Deltaproteobacteria_bacterium_RBG_19FT_COMBO_52_11.OGP98583
Nitrospirae_bacterium_RIFCSPHIGHO2_02_FULL_42_12.OGW56207
Omnitrophica_bacterium_RIFCSPLOWO2_01_FULL_45_10b.OGW85218
bacterium_BMS3Abin10.GBE04706
Candidatus_Nitrospira_inopinata.WP_062484303
Desulfosarcina_sp_BuS5.WP_027353556
Gammaproteobacteria_bacterium.PCJ50952
Kangiella_sp.PHS12437
Deltaproteobacteria_bacterium_GWD2_55_8.OGP49954
Gammaproteobacteria_bacterium_RIFCSPHIGHO2_12_FULL_45_9.OGT66414
Deltaproteobacteria_bacterium_RIFCSPHIGHO2_02_FULL_44_16.OGQ21472
Deltaproteobacteria_bacterium_RIFCSPHIGHO2_01_FULL_43_49.OGQ06355
Deltaproteobacteria_bacterium_RIFCSPLOWO2_02_FULL_44_10.OGQ45941
Deltaproteobacteria_bacterium_RIFCSPHIGHO2_02_FULL_44_16.OGQ22932
Bdellovibrio_sp_28_41_41.OYZ21459
Bdellovibrionales_bacterium_RIFOXYC1_FULL_37_79.OFZ51807
Elusimicrobia_bacterium_CG1_02_63_36.OIO07633
Elusimicrobia_bacterium_CG1_02_56_21.OIO04152
Bdellovibrionales_bacterium_RIFOXYD1_FULL_53_11.OFZ83144
Bdellovibrionales_bacterium_RIFOXYC1_FULL_54_43.OFZ53826
Bdellovibrionales_bacterium_GWA2_49_15.OFZ14898
Gammaproteobacteria_bacterium_RIFCSPHIGHO2_12_FULL_37_34.OGT43835
Gammaproteobacteria_bacterium_RIFCSPHIGHO2_12_FULL_41_20.OGT45703
Coxiella_sp_RIFCSPHIGHO2_12_FULL_42_15.OGO91641
Deltaproteobacteria_bacterium_CG11_big_fil_rev_8_21_14_0_20_45_16.PIR21816
Deltaproteobacteria_bacterium_RIFCSPHIGHO2_02_FULL_40_28.OGQ08757
Deltaproteobacteria_bacterium_RIFCSPLOWO2_12_FULL_50_11.OGQ65598
Deltaproteobacteria_bacterium_RIFCSPHIGHO2_02_FULL_50_15.OGQ29922
Acidobacteria_bacterium.PIE89368
Acidobacteria_bacterium.PIE01894
Gammaproteobacteria_bacterium_RIFCSPHIGHO2_12_FULL_40_19.OGT42242
Legionellales_bacterium_RIFCSPHIGHO2_12_FULL_37_14.OGV28065
Treponema_sp_GWB1_62_6.OHE72156
Spirochaetes_bacterium_RIFOXYC1_FULL_54_7.OHD80150
Bdellovibrionales_bacterium_GWA2_49_15.OFZ13153
Alphaproteobacteria_bacterium_RIFCSPHIGHO2_01_FULL_40_8.OFW88926
Alphaproteobacteria_bacterium_RIFCSPHIGHO2_02_FULL_40_34.OFW80202
Gammaproteobacteria_bacterium_RIFCSPHIGHO2_12_FULL_41_15.OGT50237
Gammaproteobacteria_bacterium_RIFCSPHIGHO2_12_FULL_41_15.OGT47178
Legionellales_bacterium.PCH54783
Gammaproteobacteria_bacterium_RIFCSPHIGHO2_02_FULL_42_43.OGT25493
Legionella_spiritensis.WP_058483220
Coxiella_sp_RIFCSPHIGHO2_12_FULL_44_14.OGO92408
CBU_1458
Legionellales_bacterium_RIFCSPHIGHO2_12_FULL_42_9.OGV43520
Gammaproteobacteria_bacterium_RIFCSPHIGHO2_12_FULL_40_19.OGT44205
Gammaproteobacteria_bacterium_RIFCSPHIGHO2_02_FULL_42_43.OGT25252
Deltaproteobacteria_bacterium_RIFCSPHIGHO2_12_FULL_43_9.OGQ33408
Deltaproteobacteria_bacterium_RIFCSPHIGHO2_02_FULL_40_11.OGQ13791
Bdellovibrionales_bacterium_RIFOXYA1_FULL_38_20.OFZ25065
Bdellovibrionales_bacterium_RIFOXYA1_FULL_36_14.OFZ23637
Burkholderiales_bacterium_GJ_E10.WP_045469695
Deltaproteobacteria_bacterium_RIFOXYD12_FULL_56_24.OGQ88859
Desulfonatronospira_thiodismutans.WP_040418161
delta_proteobacterium_PSCGC_5451.WP_027985105
delta_proteobacterium_PSCGC_5451.WP_027986087
Deltaproteobacteria_bacterium_RIFOXYA12_FULL_61_11.OGQ98160
Deltaproteobacteria_bacterium_RIFCSPHIGHO2_02_FULL_44_16.OGQ23242
uncultured_bacterium.EKD50980
Gammaproteobacteria_bacterium_RIFCSPHIGHO2_12_38_15.OGT24215
Gammaproteobacteria_bacterium_RIFCSPLOWO2_02_FULL_38_11.OGT66749
Gammaproteobacteria_bacterium_RIFCSPHIGHO2_12_FULL_45_9.OGT65970
Gammaproteobacteria_bacterium_RIFCSPHIGHO2_12_FULL_42_10.OGT57488
B
uncultured_bacterium
Acidobacteria_bacterium
Candidatus_Margulisbacteria_bacterium_GWF2_35_9
Mariniradius_saccharolyticus_AK6
Bacteria
bacterium_K02_2017_
bacterium_BMS3Abin10
Elusimicrobia
Elusimicrobia_bacterium_CG1_02_63_36
Elusimicrobia_bacterium_CG1_02_56_21
Candidatus_Omnitrophica
Omnitrophica_bacterium_RIFCSPHIGHO2_02_FULL_46_11
Omnitrophica_bacterium_RIFCSPLOWO2_12_FULL_50_11
Nitrospirae
Candidatus_Nitrospira_inopinata
Nitrospirae_bacterium_GWD2_44_7
Nitrospirae_bacterium_RIFCSPHIGHO2_02_FULL_42_12
Spirochaetes
Spirochaetes_bacterium_GWB1_59_5
Spirochaetes_bacterium_RIFOXYC1_FULL_54_7
Treponema
Treponema_porcinum
Treponema_sp_GWB1_62_6
Treponema_sp_RIFOXYC1_FULL_61_9
Burkholderiales_bacterium_GJ-E10
Alphaproteobacteria
Alphaproteobacteria_bacterium_RIFCSPHIGHO2_01_FULL_40_8
Alphaproteobacteria_bacterium_RIFCSPHIGHO2_02_FULL_40_34
Bdellovibrionales
Bdellovibrio_sp_28-41-41
Bdellovibrionales_bacterium_GWA2_49_15
Bdellovibrionales_bacterium_RIFOXYA1_FULL_36_14
Bdellovibrionales_bacterium_RIFOXYA1_FULL_38_20
Bdellovibrionales_bacterium_RIFOXYC1_FULL_54_43
Bdellovibrionales_bacterium_RIFOXYD1_FULL_53_11
Bdellovibrionales_bacterium_RIFOXYB1_FULL_37_110
Proteobacteria
Kangiella_sp
Legionella_spiritensis
Legionellales
Legionellales_bacterium
Legionellales_bacterium_RIFCSPHIGHO2_12_FULL_37_14
Legionellales_bacterium_RIFCSPHIGHO2_12_FULL_42_9
Gammaproteobacteria
Coxiella
Coxiella_burnetii_RSA_493
Coxiella_sp_RIFCSPHIGHO2_12_FULL_44_14
Coxiella_sp_RIFCSPHIGHO2_12_FULL_42_15
Gammaproteobacteria_bacterium_RIFCSPHIGHO2_02_FULL_42_43
Gammaproteobacteria_bacterium_RIFCSPHIGHO2_12_38_15
Gammaproteobacteria_bacterium_RIFCSPHIGHO2_12_FULL_37_34
Gammaproteobacteria_bacterium_RIFCSPHIGHO2_12_FULL_40_19
Gammaproteobacteria_bacterium_RIFCSPHIGHO2_12_FULL_41_15
Gammaproteobacteria_bacterium_RIFCSPHIGHO2_12_FULL_41_20
Gammaproteobacteria_bacterium_RIFCSPHIGHO2_12_FULL_42_10
Gammaproteobacteria_bacterium_RIFCSPHIGHO2_12_FULL_45_9
Gammaproteobacteria_bacterium
Gammaproteobacteria_bacterium_RIFCSPHIGHO2_01_FULL_42_8
Gammaproteobacteria_bacterium_RIFCSPLOWO2_02_FULL_38_11
Desulfonatronospira_thiodismutans
Desulfosarcina_sp_BuS5
Deltaproteobacteria
Deltaproteobacteria_bacterium_RIFCSPLOWO2_12_FULL_50_11
Deltaproteobacteria_bacterium_RIFOXYA12_FULL_58_15
Deltaproteobacteria_bacterium_RIFCSPHIGHO2_12_FULL_43_9
Deltaproteobacteria_bacterium_RIFCSPHIGHO2_01_FULL_43_49
Deltaproteobacteria_bacterium_GWD2_55_8
Deltaproteobacteria_bacterium_RBG_16_49_23
Deltaproteobacteria_bacterium_CG11_big_fil_rev_8_21_14_0_20_45_16
Deltaproteobacteria_bacterium_RBG_19FT_COMBO_52_11
Deltaproteobacteria_bacterium_RIFCSPHIGHO2_02_FULL_40_28
Deltaproteobacteria_bacterium_RIFCSPHIGHO2_02_FULL_44_16
delta_proteobacterium_PSCGC_5296
delta_proteobacterium_PSCGC_5342
delta_proteobacterium_PSCGC_5451
Deltaproteobacteria_bacterium_RIFOXYA12_FULL_61_11
Deltaproteobacteria_bacterium_RIFCSPHIGHO2_02_FULL_50_15
Deltaproteobacteria_bacterium_RIFCSPLOWO2_02_FULL_44_10
Deltaproteobacteria_bacterium_RIFCSPHIGHO2_02_FULL_40_11
Deltaproteobacteria_bacterium_RIFOXYD12_FULL_56_24

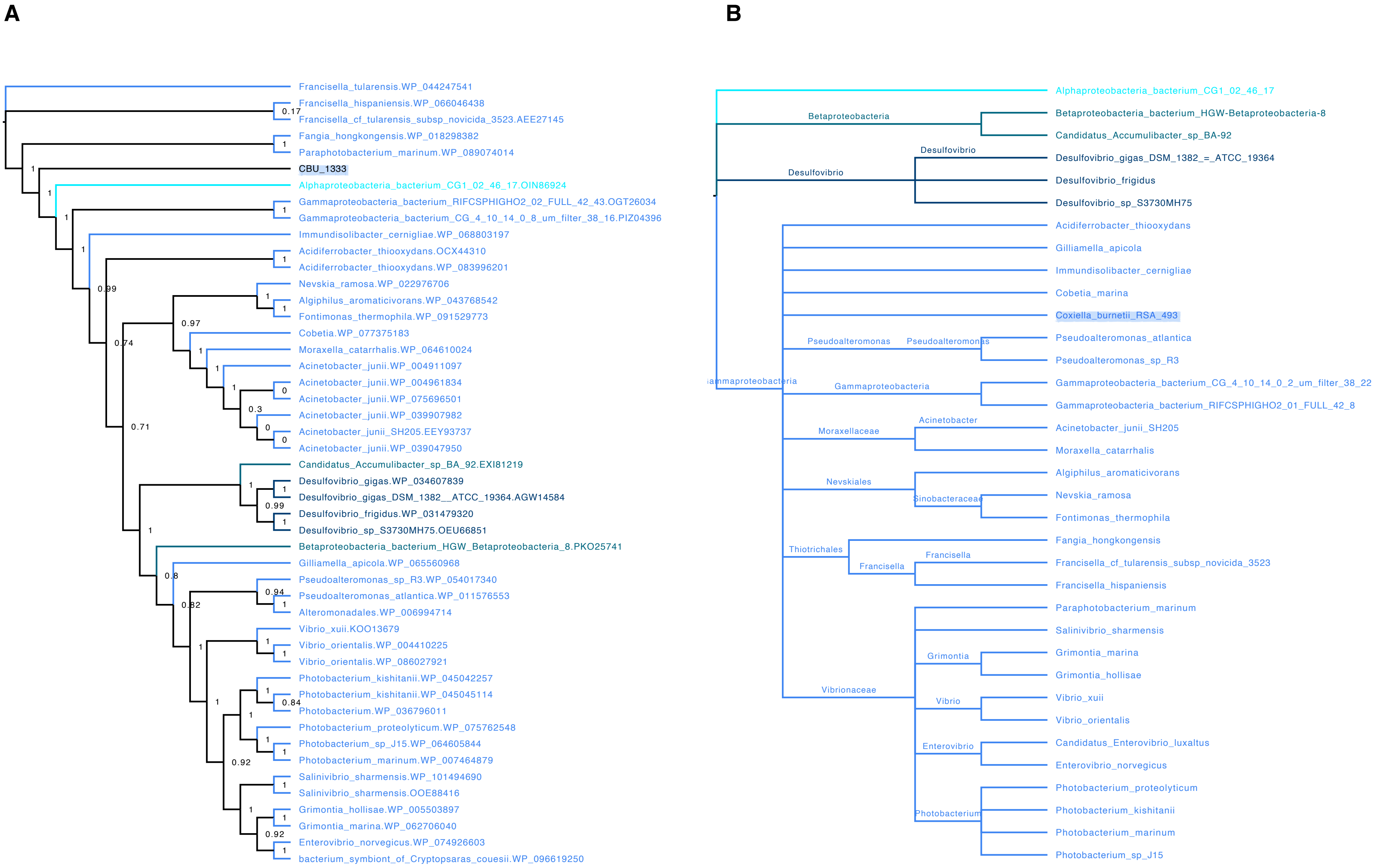
A
Francisella_tularensis.WP_044247541
Francisella_hispaniensis.WP_066046438
Francisella_cf_tularensis_subsp_novicida_3523.AEE27145
Fangia_hongkongensis.WP_018298382
Paraphotobacterium_marinum.WP_089074014
CBU_1333
Alphaproteobacteria_bacterium_CG1_02_46_17.OIN86924
Gammaproteobacteria_bacterium_RIFCSPHIGHO2_02_FULL_42_43.OGT26034
Gammaproteobacteria_bacterium_CG_4_10_14_0_8_um_filter_38_16.PIZ04396
Immundisolibacter_cernigliae.WP_068803197
Acidiferrobacter_thiooxydans.OCX44310
Acidiferrobacter_thiooxydans.WP_083996201
Nevskia_ramosa.WP_022976706
Algiphilus_aromaticivorans.WP_043768542
Fontimonas_thermophila.WP_091529773
Cobetia.WP_077375183
Moraxella_catarrhalis.WP_064610024
Acinetobacter_junii.WP_004911097
Acinetobacter_junii.WP_004961834
Acinetobacter_junii.WP_075696501
Acinetobacter_junii.WP_039907982
Acinetobacter_junii_SH205.EEY93737
Acinetobacter_junii.WP_039047950
Candidatus_Accumulibacter_sp_BA_92.EXI81219
Desulfovibrio_gigas.WP_034607839
Desulfovibrio_gigas_DSM_1382__ATCC_19364.AGW14584
Desulfovibrio_frigidus.WP_031479320
Desulfovibrio_sp_S3730MH75.OEU66851
Betaproteobacteria_bacterium_HGW_Betaproteobacteria_8.PKO25741
Gilliamella_apicola.WP_065560968
Pseudoalteromonas_sp_R3.WP_054017340
Pseudoalteromonas_atlantica.WP_011576553
Alteromonadales.WP_006994714
Vibrio_xuii.KOO13679
Vibrio_orientalis.WP_004410225
Vibrio_orientalis.WP_086027921
Photobacterium_kishitanii.WP_045042257
Photobacterium_kishitanii.WP_045045114
Photobacterium.WP_036796011
Photobacterium_proteolyticum.WP_075762548
Photobacterium_sp_J15.WP_064605844
Photobacterium_marinum.WP_007464879
Salinivibrio_sharmensis.WP_101494690
Salinivibrio_sharmensis.OOE88416
Grimontia_hollisae.WP_005503897
Grimontia_marina.WP_062706040
Enterovibrio_norvegicus.WP_074926603
bacterium_symbiont_of_Cryptopsaras_couesii.WP_096619250
B
Alphaproteobacteria_bacterium_CG1_02_46_17
Betaproteobacteria
Betaproteobacteria_bacterium_HGW-Betaproteobacteria-8
Candidatus_Accumulibacter_sp_BA-92
Desulfovibrio
Desulfovibrio_gigas_DSM_1382_=_ATCC_19364
Desulfovibrio_frigidus
Desulfovibrio_sp_S3730MH75
Acidiferrobacter_thiooxydans
Gilliamella_apicola
Immundisolibacter_cernigliae
Cobetia_marina
Coxiella_burnetii_RSA_493
Pseudoalteromonas
Pseudoalteromonas_atlantica
Pseudoalteromonas_sp_R3
Gammaproteobacteria
Gammaproteobacteria_bacterium_CG_4_10_14_0_2_um_filter_38_22
Gammaproteobacteria_bacterium_RIFCSPHIGHO2_01_FULL_42_8
Moraxellaceae
Acinetobacter
Acinetobacter_junii_SH205
Moraxella_catarrhalis
Nevskiales
Sinobacteraceae
Algiphilus_aromaticivorans
Nevskia_ramosa
Fontimonas_thermophila
Thiotrichales
Francisella
Fangia_hongkongensis
Francisella_cf_tularensis_subsp_novicida_3523
Francisella_hispaniensis
Vibrionaceae
Paraphotobacterium_marinum
Salinivibrio_sharmensis
Grimontia
Grimontia_marina
Grimontia_hollisae
Vibrio
Vibrio_xuii
Vibrio_orientalis
Enterovibrio
Candidatus_Enterovibrio_luxaltus
Enterovibrio_norvegicus
Photobacterium
Photobacterium_proteolyticum
Photobacterium_kishitanii
Photobacterium_marinum
Photobacterium_sp_J15

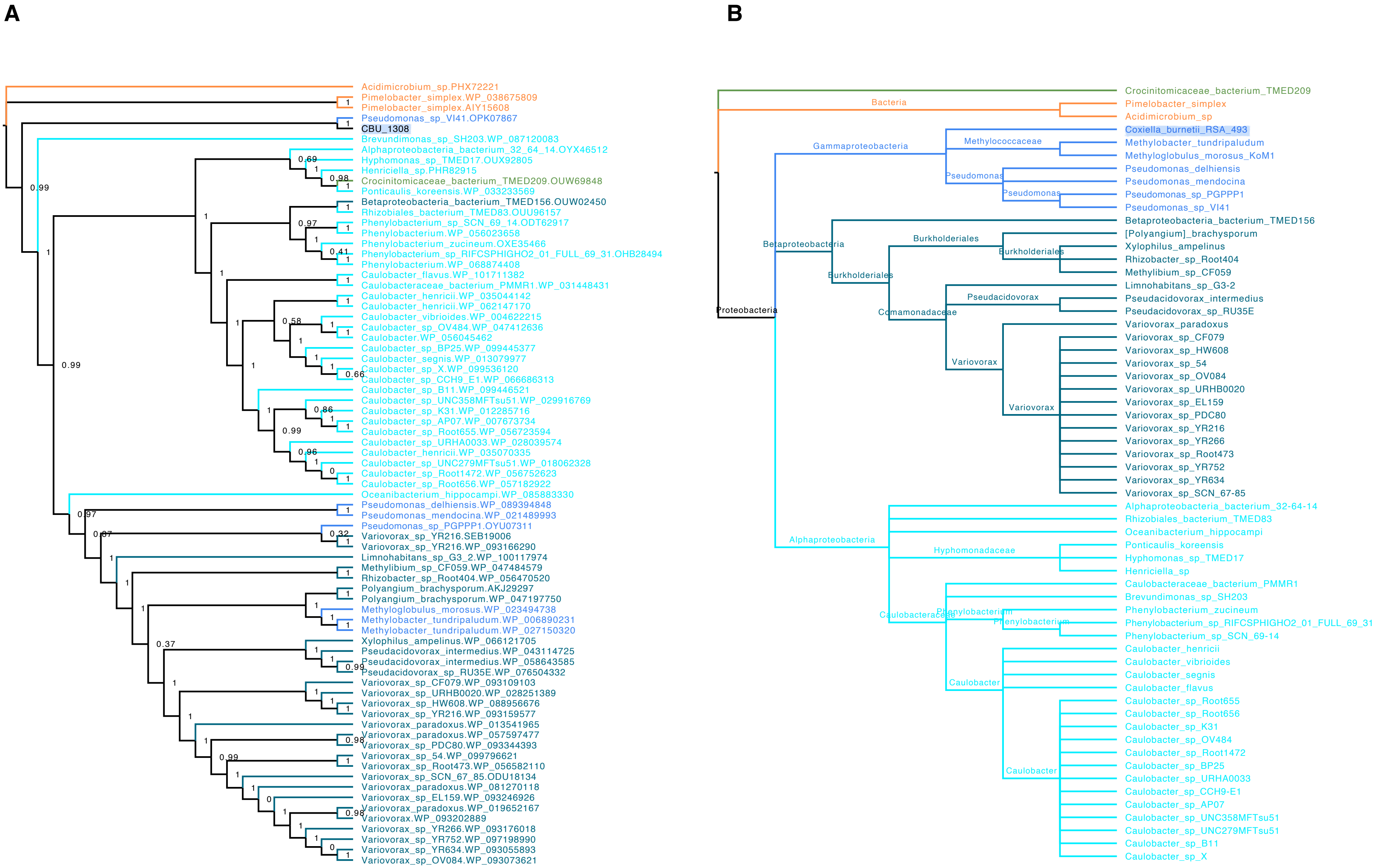
A
Acidimicrobium_sp.PHX72221
Pimelobacter_simplex.WP_038675809
Pimelobacter_simplex.AIY15608
Pseudomonas_sp_VI41.OPK07867
CBU_1308
Brevundimonas_sp_SH203.WP_087120083
Alphaproteobacteria_bacterium_32_64_14.OYX46512
Hyphomonas_sp_TMED17.OUX92805
Henriciella_sp.PHR82915
Crocinitomicaceae_bacterium_TMED209.OUW69848
Ponticaulis_koreensis.WP_033233569
Betaproteobacteria_bacterium_TMED156.OUW02450
Rhizobiales_bacterium_TMED83.OUU96157
Phenylobacterium_sp_SCN_69_14.ODT62917
Phenylobacterium.WP_056023658
Phenylobacterium_zucineum.OXE35466
Phenylobacterium_sp_RIFCSPHIGHO2_01_FULL_69_31.OHB28494
Phenylobacterium.WP_068874408
Caulobacter_flavus.WP_101711382
Caulobacteraceae_bacterium_PMMR1.WP_031448431
Caulobacter_henricii.WP_035044142
Caulobacter_henricii.WP_062147170
Caulobacter_vibrioides.WP_004622215
Caulobacter_sp_OV484.WP_047412636
Caulobacter.WP_056045462
Caulobacter_sp_BP25.WP_099445377
Caulobacter_segnis.WP_013079977
Caulobacter_sp_X.WP_099536120
Caulobacter_sp_CCH9_E1.WP_066686313
Caulobacter_sp_B11.WP_099446521
Caulobacter_sp_UNC358MFTsu51.WP_029916769
Caulobacter_sp_K31.WP_012285716
Caulobacter_sp_AP07.WP_007673734
Caulobacter_sp_Root655.WP_056723594
Caulobacter_sp_URHA0033.WP_028039574
Caulobacter_henricii.WP_035070335
Caulobacter_sp_UNC279MFTsu51.WP_018062328
Caulobacter_sp_Root1472.WP_056752623
Caulobacter_sp_Root656.WP_057182922
Oceanibacterium_hippocampi.WP_085883330
Pseudomonas_delhiensis.WP_089394848
Pseudomonas_mendocina.WP_021489993
Pseudomonas_sp_PGPPP1.OYU07311
Variovorax_sp_YR216.SEB19006
Variovorax_sp_YR216.WP_093166290
Limnohabitans_sp_G3_2.WP_100117974
Methylibium_sp_CF059.WP_047484579
Rhizobacter_sp_Root404.WP_056470520
Polyangium_brachysporum.AKJ29297
Polyangium_brachysporum.WP_047197750
Methyloglobulus_morosus.WP_023494738
Methylobacter_tundripaludum.WP_006890231
Methylobacter_tundripaludum.WP_027150320
Xylophilus_ampelinus.WP_066121705
Pseudacidovorax_intermedius.WP_043114725
Pseudacidovorax_intermedius.WP_058643585
Pseudacidovorax_sp_RU35E.WP_076504332
Variovorax_sp_CF079.WP_093109103
Variovorax_sp_URHB0020.WP_028251389
Variovorax_sp_HW608.WP_088956676
Variovorax_sp_YR216.WP_093159577
Variovorax_paradoxus.WP_013541965
Variovorax_paradoxus.WP_057597477
Variovorax_sp_PDC80.WP_093344393
Variovorax_sp_54.WP_099796621
Variovorax_sp_Root473.WP_056582110
Variovorax_sp_SCN_67_85.ODU18134
Variovorax_paradoxus.WP_081270118
Variovorax_sp_EL159.WP_093246926
Variovorax_paradoxus.WP_019652167
Variovorax.WP_093202889
Variovorax_sp_YR266.WP_093176018
Variovorax_sp_YR752.WP_097198990
Variovorax_sp_YR634.WP_093055893
Variovorax_sp_OV084.WP_093073621
B
Bacteria
Proteobacteria
Gammaproteobacteria
Methylococcaceae
Pseudomonas
Betaproteobacteria
Burkholderiales
Comamonadaceae
Pseudacidovorax
Variovorax
Alphaproteobacteria
Hyphomonadaceae
Caulobacteraceae
Phenylobacterium
Caulobacter
Crocinitomicaceae_bacterium_TMED209
Pimelobacter_simplex
Acidimicrobium_sp
Coxiella_burnetii_RSA_493
Methylobacter_tundripaludum
Methyloglobulus_morosus_KoM1
Pseudomonas_delhiensis
Pseudomonas_mendocina
Pseudomonas_sp_PGPPP1
Pseudomonas_sp_VI41
Betaproteobacteria_bacterium_TMED156
[Polyangium]_brachysporum
Xylophilus_ampelinus
Rhizobacter_sp_Root404
Methylibium_sp_CF059
Limnohabitans_sp_G3-2
Pseudacidovorax_intermedius
Pseudacidovorax_sp_RU35E
Variovorax_paradoxus
Variovorax_sp_CF079
Variovorax_sp_HW608
Variovorax_sp_54
Variovorax_sp_OV084
Variovorax_sp_URHB0020
Variovorax_sp_EL159
Variovorax_sp_PDC80
Variovorax_sp_YR216
Variovorax_sp_YR266
Variovorax_sp_Root473
Variovorax_sp_YR752
Variovorax_sp_YR634
Variovorax_sp_SCN_67-85
Alphaproteobacteria_bacterium_32-64-14
Rhizobiales_bacterium_TMED83
Oceanibacterium_hippocampi
Ponticaulis_koreensis
Hyphomonas_sp_TMED17
Henriciella_sp
Caulobacteraceae_bacterium_PMMR1
Brevundimonas_sp_SH203
Phenylobacterium_zucineum
Phenylobacterium_sp_RIFCSPHIGHO2_01_FULL_69_31
Phenylobacterium_sp_SCN_69-14
Caulobacter_henricii
Caulobacter_vibrioides
Caulobacter_segnis
Caulobacter_flavus
Caulobacter_sp_Root655
Caulobacter_sp_Root656
Caulobacter_sp_K31
Caulobacter_sp_OV484
Caulobacter_sp_Root1472
Caulobacter_sp_BP25
Caulobacter_sp_URHA0033
Caulobacter_sp_CCH9-E1
Caulobacter_sp_AP07
Caulobacter_sp_UNC358MFTsu51
Caulobacter_sp_UNC279MFTsu51
Caulobacter_sp_B11
Caulobacter_sp_X

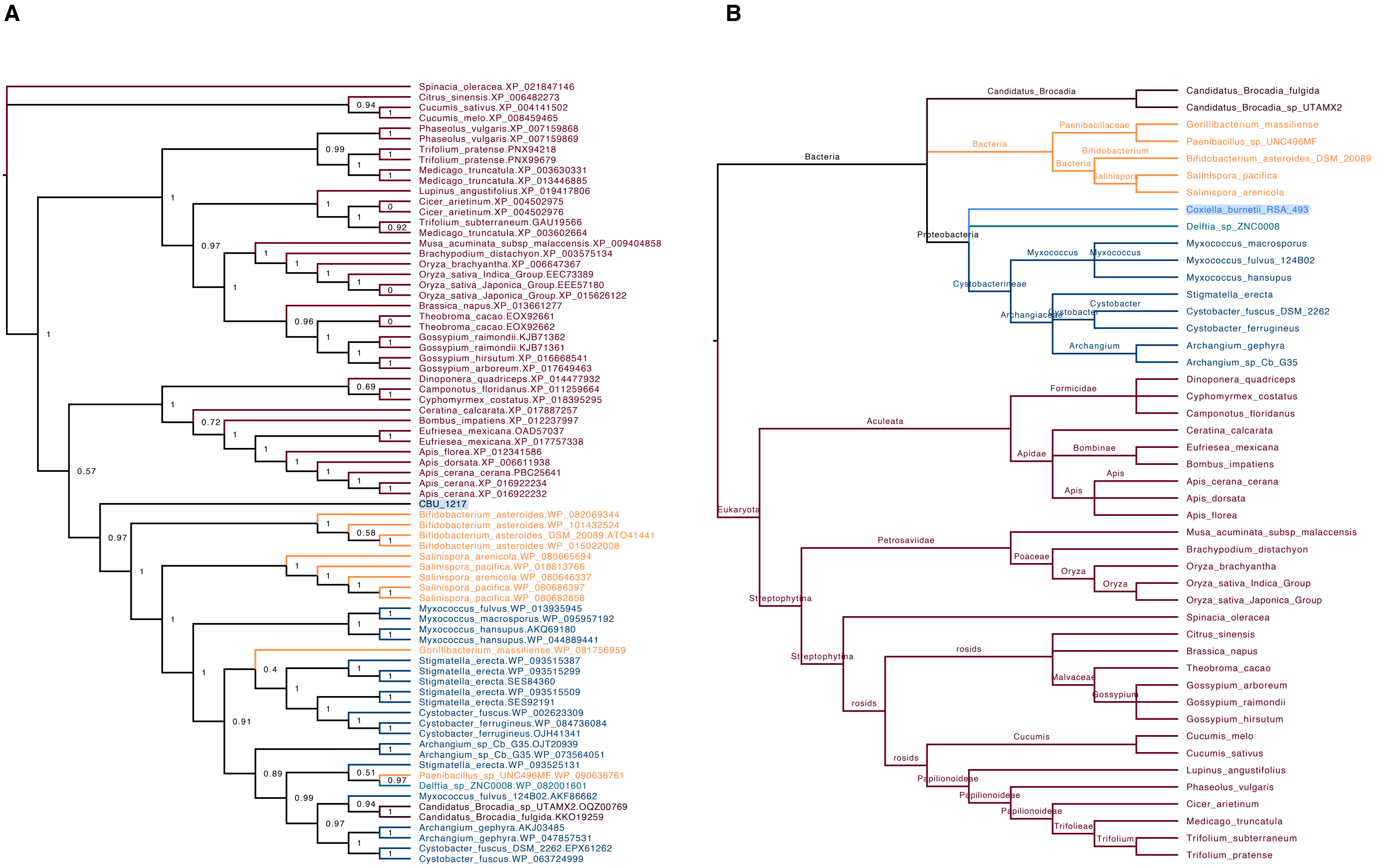
A
Spinacia_oleracea.XP_021847146
Citrus_sinensis.XP_006482273
Cucumis_sativus.XP_004141502
Cucumis_melo.XP_008459465
Phaseolus_vulgaris.XP_007159868
Phaseolus_vulgaris.XP_007159869
Trifolium_pratense.PNX94218
Trifolium_pratense.PNX99679
Medicago_truncatula.XP_003630331
Medicago_truncatula.XP_013446885
Lupinus_angustifolius.XP_019417806
Cicer_arietinum.XP_004502975
Cicer_arietinum.XP_004502976
Trifolium_subterraneum.GAU19566
Medicago_truncatula.XP_003602664
Musa_acuminata_subsp_malaccensis.XP_009404858
Brachypodium_distachyon.XP_003575134
Oryza_brachyantha.XP_006647367
Oryza_sativa_Indica_Group.EEC73389
Oryza_sativa_Japonica_Group.EEE57180
Oryza_sativa_Japonica_Group.XP_015626122
Brassica_napus.XP_013661277
Theobroma_cacao.EOX92661
Theobroma_cacao.EOX92662
Gossypium_raimondii.KJB71362
Gossypium_raimondii.KJB71361
Gossypium_hirsutum.XP_016668541
Gossypium_arboreum.XP_017649463
Dinoponera_quadriceps.XP_014477932
Camponotus_floridanus.XP_011259664
Cyphomyrmex_costatus.XP_018395295
Ceratina_calcarata.XP_017887257
Bombus_impatiens.XP_012237997
Eufriesea_mexicana.OAD57037
Eufriesea_mexicana.XP_017757338
Apis_florea.XP_012341586
Apis_dorsata.XP_006611938
Apis_cerana_cerana.PBC25641
Apis_cerana.XP_016922234
Apis_cerana.XP_016922232
CBU_1217
Bifidobacterium_asteroides.WP_082069344
Bifidobacterium_asteroides.WP_101432524
Bifidobacterium_asteroides_DSM_20089.ATO41441
Bifidobacterium_asteroides.WP_015022008
Salinispora_arenicola.WP_080665694
Salinispora_pacifica.WP_018813766
Salinispora_arenicola.WP_080646337
Salinispora_pacifica.WP_080686397
Salinispora_pacifica.WP_080682858
Myxococcus_fulvus.WP_013935945
Myxococcus_macrosporus.WP_095957192
Myxococcus_hansupus.AKQ69180
Myxococcus_hansupus.WP_044889441
Gorillibacterium_massiliense.WP_081756959
Stigmatella_erecta.WP_093515387
Stigmatella_erecta.WP_093515299
Stigmatella_erecta.SES84360
Stigmatella_erecta.WP_093515509
Stigmatella_erecta.SES92191
Cystobacter_fuscus.WP_002623309
Cystobacter_ferrugineus.WP_084736084
Cystobacter_ferrugineus.OJH41341
Archangium_sp_Cb_G35.OJT20939
Archangium_sp_Cb_G35.WP_073564051
Stigmatella_erecta.WP_093525131
Paenibacillus_sp_UNC496MF.WP_090636761
Delftia_sp_ZNC0008.WP_082001601
Myxococcus_fulvus_124B02.AKF86662
Candidatus_Brocadia_sp_UTAMX2.OQZ00769
Candidatus_Brocadia_fulgida.KKO19259
Archangium_gephyra.AKJ03485
Archangium_gephyra.WP_047857531
Cystobacter_fuscus_DSM_2262.EPX61262
Cystobacter_fuscus.WP_063724999
B
Bacteria
Candidatus_Brocadia
Candidatus_Brocadia_fulgida
Candidatus_Brocadia_sp_UTAMX2
Paenibacillaceae
Gorillibacterium_massiliense
Paenibacillus_sp_UNC496MF
Bifidobacterium
Bifidobacterium_asteroides_DSM_20089
Salinispora
Salinispora_pacifica
Salinispora_arenicola
Proteobacteria
Coxiella_burnetii_RSA_493
Delftia_sp_ZNC0008
Cystobacterineae
Myxococcus
Myxococcus_macrosporus
Myxococcus_fulvus_124B02
Myxococcus_hansupus
Archangiaceae
Cystobacter
Stigmatella_erecta
Cystobacter_fuscus_DSM_2262
Cystobacter_ferrugineus
Archangium
Archangium_gephyra
Archangium_sp_Cb_G35
Eukaryota
Aculeata
Formicidae
Dinoponera_quadriceps
Cyphomyrmex_costatus
Camponotus_floridanus
Apidae
Bombinae
Apis
Ceratina_calcarata
Eufriesea_mexicana
Bombus_impatiens
Apis_cerana_cerana
Apis_dorsata
Apis_florea
Streptophytina
Petrosaviidae
Poaceae
Oryza
Musa_acuminata_subsp_malaccensis
Brachypodium_distachyon
Oryza_brachyantha
Oryza_sativa_Indica_Group
Oryza_sativa_Japonica_Group
rosids
Malvaceae
Gossypium
Spinacia_oleracea
Citrus_sinensis
Brassica_napus
Theobroma_cacao
Gossypium_arboreum
Gossypium_raimondii
Gossypium_hirsutum
Cucumis
Cucumis_melo
Cucumis_sativus
Papilionoideae
Trifolieae
Trifolium
Lupinus_angustifolius
Phaseolus_vulgaris
Cicer_arietinum
Medicago_truncatula
Trifolium_subterraneum
Trifolium_pratense

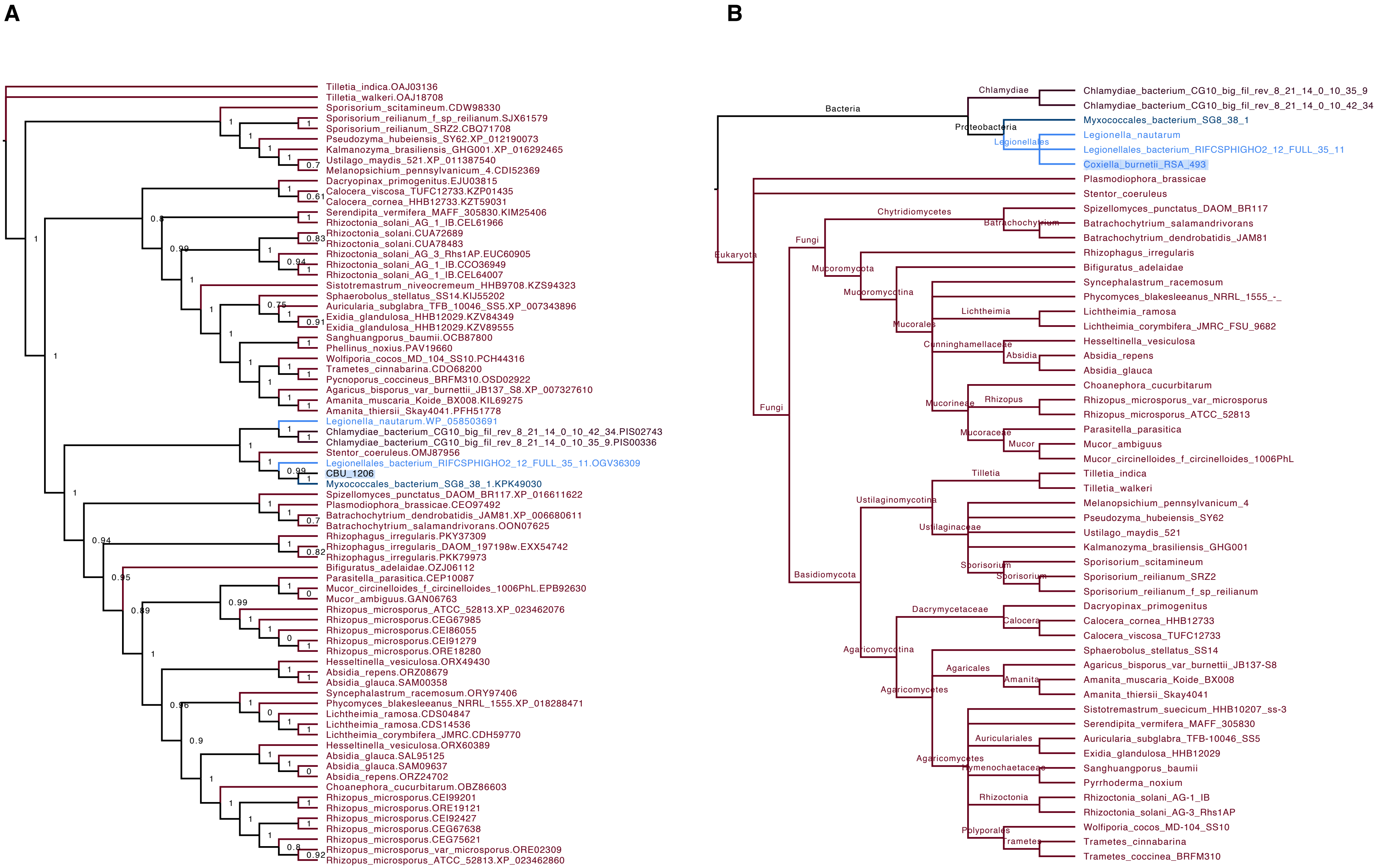
A
Tilletia_indica.OAJ03136
Tilletia_walkeri.OAJ18708
Sporisorium_scitamineum.CDW98330
Sporisorium_reilianum_f_sp_reilianum.SJX61579
Sporisorium_reilianum_SRZ2.CBQ71708
Pseudozyma_hubeiensis_SY62.XP_012190073
Kalmanozyma_brasiliensis_GHG001.XP_016292465
Ustilago_maydis_521.XP_011387540
Melanopsichium_pennsylvanicum_4.CDI52369
Dacryopinax_primogenitus.EJU03815
Calocera_viscosa_TUFC12733.KZP01435
Calocera_cornea_HHB12733.KZT59031
Serendipita_vermifera_MAFF_305830.KIM25406
Rhizoctonia_solani_AG_1_IB.CEL61966
Rhizoctonia_solani.CUA72689
Rhizoctonia_solani.CUA78483
Rhizoctonia_solani_AG_3_Rhs1AP.EUC60905
Rhizoctonia_solani_AG_1_IB.CCO36949
Rhizoctonia_solani_AG_1_IB.CEL64007
Sistotremastrum_niveocremeum_HHB9708.KZS94323
Sphaerobolus_stellatus_SS14.KIJ55202
Auricularia_subglabra_TFB_10046_SS5.XP_007343896
Exidia_glandulosa_HHB12029.KZV84349
Exidia_glandulosa_HHB12029.KZV89555
Sanghuangporus_baumii.OCB87800
Phellinus_noxius.PAV19660
Wolfiporia_cocos_MD_104_SS10.PCH44316
Trametes_cinnabarina.CDO68200
Pycnoporus_coccineus_BRFM310.OSD02922
Agaricus_bisporus_var_burnettii_JB137_S8.XP_007327610
Amanita_muscaria_Koide_BX008.KIL69275
Amanita_thiersii_Skay4041.PFH51778
Legionella_nautarum.WP_058503691
Chlamydiae_bacterium_CG10_big_fil_rev_8_21_14_0_10_42_34.PIS02743
Chlamydiae_bacterium_CG10_big_fil_rev_8_21_14_0_10_35_9.PIS00336
Stentor_coeruleus.OMJ87956
Legionellales_bacterium_RIFCSPHIGHO2_12_FULL_35_11.OGV36309
CBU_1206
Myxococcales_bacterium_SG8_38_1.KPK49030
Spizellomyces_punctatus_DAOM_BR117.XP_016611622
Plasmodiophora_brassicae.CEO97492
Batrachochytrium_dendrobatidis_JAM81.XP_006680611
Batrachochytrium_salamandrivorans.OON07625
Rhizophagus_irregularis.PKY37309
Rhizophagus_irregularis_DAOM_197198w.EXX54742
Rhizophagus_irregularis.PKK79973
Bifiguratus_adelaidae.OZJ06112
Parasitella_parasitica.CEP10087
Mucor_circinelloides_f_circinelloides_1006PhL.EPB92630
Mucor_ambiguus.GAN06763
Rhizopus_microsporus_ATCC_52813.XP_023462076
Rhizopus_microsporus.CEG67985
Rhizopus_microsporus.CEI86055
Rhizopus_microsporus.CEI91279
Rhizopus_microsporus.ORE18280
Hesseltinella_vesiculosa.ORX49430
Absidia_repens.ORZ08679
Absidia_glauca.SAM00358
Syncephalastrum_racemosum.ORY97406
Phycomyces_blakesleeanus_NRRL_1555.XP_018288471
Lichtheimia_ramosa.CDS04847
Lichtheimia_ramosa.CDS14536
Lichtheimia_corymbifera_JMRC.CDH59770
Hesseltinella_vesiculosa.ORX60389
Absidia_glauca.SAL95125
Absidia_glauca.SAM09637
Absidia_repens.ORZ24702
Choanephora_cucurbitarum.OBZ86603
Rhizopus_microsporus.CEI99201
Rhizopus_microsporus.ORE19121
Rhizopus_microsporus.CEI92427
Rhizopus_microsporus.CEG67638
Rhizopus_microsporus.CEG75621
Rhizopus_microsporus_var_microsporus.ORE02309
Rhizopus_microsporus_ATCC_52813.XP_023462860
B
Bacteria
Chlamydiae
Chlamydiae_bacterium_CG10_big_fil_rev_8_21_14_0_10_35_9
Chlamydiae_bacterium_CG10_big_fil_rev_8_21_14_0_10_42_34
Proteobacteria
Myxococcales_bacterium_SG8_38_1
Legionellales
Legionella_nautarum
Legionellales_bacterium_RIFCSPHIGHO2_12_FULL_35_11
Coxiella_burnetii_RSA_493
Eukaryota
Plasmodiophora_brassicae
Stentor_coeruleus
Fungi
Chytridiomycetes
Batrachochytrium
Spizellomyces_punctatus_DAOM_BR117
Batrachochytrium_salamandrivorans
Batrachochytrium_dendrobatidis_JAM81
Mucoromycota
Rhizophagus_irregularis
Mucoromycotina
Bifiguratus_adelaidae
Mucorales
Syncephalastrum_racemosum
Phycomyces_blakesleeanus_NRRL_1555_-_
Lichtheimia
Lichtheimia_ramosa
Lichtheimia_corymbifera_JMRC_FSU_9682
Cunninghamellaceae
Absidia
Hesseltinella_vesiculosa
Absidia_repens
Absidia_glauca
Mucorineae
Rhizopus
Choanephora_cucurbitarum
Rhizopus_microsporus_var_microsporus
Rhizopus_microsporus_ATCC_52813
Mucoraceae
Mucor
Parasitella_parasitica
Mucor_ambiguus
Mucor_circinelloides_f_circinelloides_1006PhL
Basidiomycota
Ustilaginomycotina
Tilletia
Tilletia_indica
Tilletia_walkeri
Ustilaginaceae
Melanopsichium_pennsylvanicum_4
Pseudozyma_hubeiensis_SY62
Ustilago_maydis_521
Kalmanozyma_brasiliensis_GHG001
Sporisorium
Sporisorium_scitamineum
Sporisorium_reilianum_SRZ2
Sporisorium_reilianum_f_sp_reilianum
Agaricomycotina
Dacrymycetaceae
Calocera
Dacryopinax_primogenitus
Calocera_cornea_HHB12733
Calocera_viscosa_TUFC12733
Agaricomycetes
Sphaerobolus_stellatus_SS14
Agaricales
Amanita
Agaricus_bisporus_var_burnettii_JB137-S8
Amanita_muscaria_Koide_BX008
Amanita_thiersii_Skay4041
Sistotremastrum_suecicum_HHB10207_ss-3
Serendipita_vermifera_MAFF_305830
Auriculariales
Auricularia_subglabra_TFB-10046_SS5
Exidia_glandulosa_HHB12029
Hymenochaetaceae
Sanghuangporus_baumii
Pyrrhoderma_noxium
Rhizoctonia
Rhizoctonia_solani_AG-1_IB
Rhizoctonia_solani_AG-3_Rhs1AP
Polyporales
Trametes
Wolfiporia_cocos_MD-104_SS10
Trametes_cinnabarina
Trametes_coccinea_BRFM310

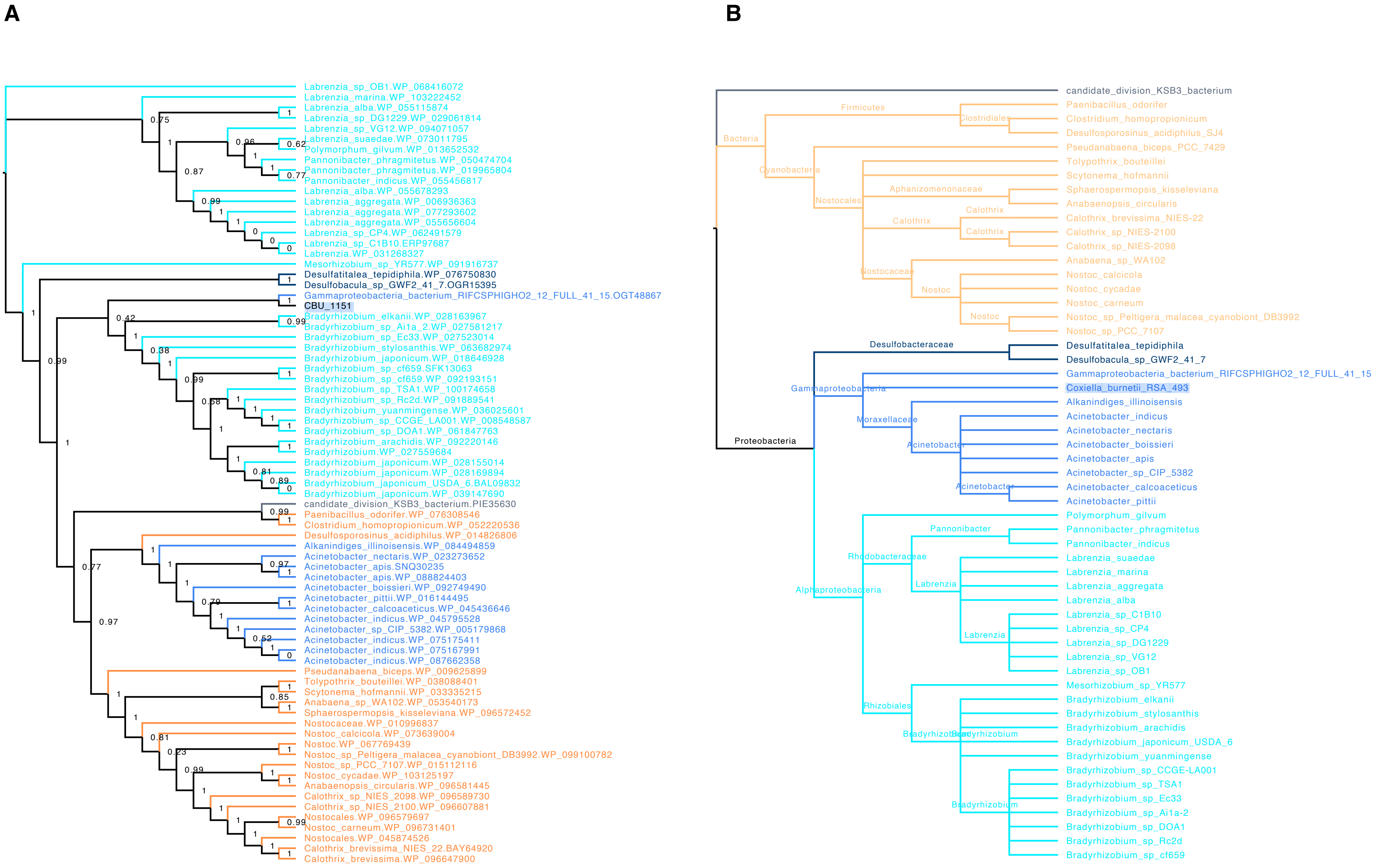

A
Labrenzia_sp_OB1.WP_068416072
Labrenzia_marina.WP_103222452
Labrenzia_alba.WP_055115874
Labrenzia_sp_DG1229.WP_029061814
Labrenzia_sp_VG12.WP_094071057
Labrenzia_suaedae.WP_073011795
Polymorphum_gilvum.WP_013652532
Pannonibacter_phragmitetus.WP_050474704
Pannonibacter_phragmitetus.WP_019965804
Pannonibacter_indicus.WP_055456817
Labrenzia_alba.WP_055678293
Labrenzia_aggregata.WP_008936363
Labrenzia_aggregata.WP_077293602
Labrenzia_aggregata.WP_055656604
Labrenzia_sp_CP4.WP_062491579
Labrenzia_sp_C1B10.ERP97687
Labrenzia.WP_031268327
Mesorhizobium_sp_YR577.WP_091916737
Desulfatitalea_tepidiphila.WP_076750830
Desulfobacula_sp_GWF2_41_7.OGR15395
Gammaproteobacteria_bacterium_RIFCSPHIGHO2_12_FULL_41_15.OGT48867
CBU_1151
Bradyrhizobium_elkanii.WP_028163967
Bradyrhizobium_sp_Ai1a_2.WP_027581217
Bradyrhizobium_sp_Ec33.WP_027523014
Bradyrhizobium_stylosanthis.WP_063682974
Bradyrhizobium_japonicum.WP_018646928
Bradyrhizobium_sp_cf659.SFK13063
Bradyrhizobium_sp_cf659.WP_092193151
Bradyrhizobium_sp_TSA1.WP_100174658
Bradyrhizobium_sp_Rc2d.WP_091889541
Bradyrhizobium_yuanmingense.WP_036025601
Bradyrhizobium_sp_CCGE_LA001.WP_008548587
Bradyrhizobium_sp_DOA1.WP_061847763
Bradyrhizobium_arachidis.WP_092220146
Bradyrhizobium.WP_027559684
Bradyrhizobium_japonicum.WP_028155014
Bradyrhizobium_japonicum.WP_028169894
Bradyrhizobium_japonicum_USDA_6.BAL09832
Bradyrhizobium_japonicum.WP_039147690
candidate_division_KSB3_bacterium.PIE35630
Paenibacillus_odorifer.WP_076308546
Clostridium_homopropionicum.WP_052220536
Desulfosporosinus_acidiphilus.WP_014826806
Alkanindiges_illinoisensis.WP_084494859
Acinetobacter_nectaris.WP_023273652
Acinetobacter_apis.SNQ30235
Acinetobacter_apis.WP_088824403
Acinetobacter_boissieri.WP_092749490
Acinetobacter_pittii.WP_016144495
Acinetobacter_calcoaceticus.WP_045436646
Acinetobacter_indicus.WP_045795528
Acinetobacter_sp_CIP_5382.WP_005179868
Acinetobacter_indicus.WP_075175411
Acinetobacter_indicus.WP_075167991
Acinetobacter_indicus.WP_087662358
Pseudanabaena_biceps.WP_009625899
Tolypothrix_bouteillei.WP_038088401
Scytonema_hofmannii.WP_033335215
Anabaena_sp_WA102.WP_053540173
Sphaerospermopsis_kisseleviana.WP_096572452
Nostocaceae.WP_010996837
Nostoc_calcicola.WP_073639004
Nostoc.WP_067769439
Nostoc_sp_Peltigera_malacea_cyanobiont_DB3992.WP_099100782
Nostoc_sp_PCC_7107.WP_015112116
Nostoc_cycadae.WP_103125197
Anabaenopsis_circularis.WP_096581445
Calothrix_sp_NIES_2098.WP_096589730
Calothrix_sp_NIES_2100.WP_096607881
Nostocales.WP_096579697
Nostoc_carneum.WP_096731401
Nostocales.WP_045874526
Calothrix_brevissima_NIES_22.BAY64920
Calothrix_brevissima.WP_096647900
B
candidate_division_KSB3_bacterium
Bacteria
Firmicutes
Clostridiales
Paenibacillus_odorifer
Clostridium_homopropionicum
Desulfosporosinus_acidiphilus_SJ4
Cyanobacteria
Pseudanabaena_biceps_PCC_7429
Nostocales
Tolypothrix_bouteillei
Scytonema_hofmannii
Aphanizomenonaceae
Sphaerospermopsis_kisseleviana
Anabaenopsis_circularis
Calothrix
Calothrix_brevissima_NIES-22
Calothrix_sp_NIES-2100
Calothrix_sp_NIES-2098
Nostocaceae
Nostoc
Anabaena_sp_WA102
Nostoc_calcicola
Nostoc_cycadae
Nostoc_carneum
Nostoc_sp_Peltigera_malacea_cyanobiont_DB3992
Nostoc_sp_PCC_7107
Proteobacteria
Desulfobacteraceae
Desulfatitalea_tepidiphila
Desulfobacula_sp_GWF2_41_7
Gammaproteobacteria
Gammaproteobacteria_bacterium_RIFCSPHIGHO2_12_FULL_41_15
Coxiella_burnetii_RSA_493
Moraxellaceae
Alkanindiges_illinoisensis
Acinetobacter
Acinetobacter_indicus
Acinetobacter_nectaris
Acinetobacter_boissieri
Acinetobacter_apis
Acinetobacter_sp_CIP_5382
Acinetobacter_calcoaceticus
Acinetobacter_pittii
Alphaproteobacteria
Rhodobacteraceae
Pannonibacter
Polymorphum_gilvum
Pannonibacter_phragmitetus
Pannonibacter_indicus
Labrenzia
Labrenzia_suaedae
Labrenzia_marina
Labrenzia_aggregata
Labrenzia_alba
Labrenzia_sp_C1B10
Labrenzia_sp_CP4
Labrenzia_sp_DG1229
Labrenzia_sp_VG12
Labrenzia_sp_OB1
Rhizobiales
Bradyrhizobium
Mesorhizobium_sp_YR577
Bradyrhizobium_elkanii
Bradyrhizobium_stylosanthis
Bradyrhizobium_arachidis
Bradyrhizobium_japonicum_USDA_6
Bradyrhizobium_yuanmingense
Bradyrhizobium_sp_CCGE-LA001
Bradyrhizobium_sp_TSA1
Bradyrhizobium_sp_Ec33
Bradyrhizobium_sp_Ai1a-2
Bradyrhizobium_sp_DOA1
Bradyrhizobium_sp_Rc2d
Bradyrhizobium_sp_cf659

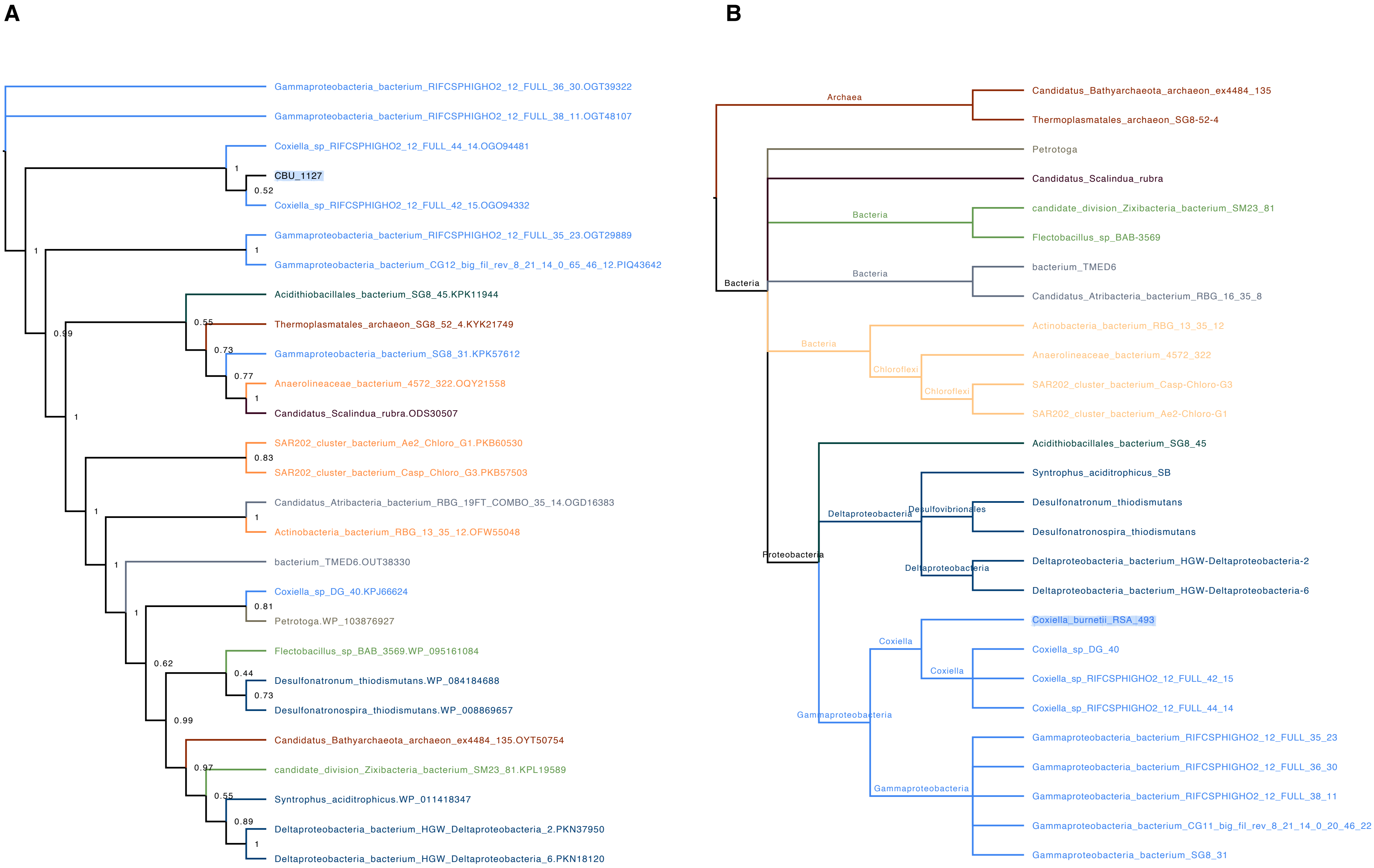
A
Gammaproteobacteria_bacterium_RIFCSPHIGHO2_12_FULL_36_30.OGT39322
Gammaproteobacteria_bacterium_RIFCSPHIGHO2_12_FULL_38_11.OGT48107
Coxiella_sp_RIFCSPHIGHO2_12_FULL_44_14.OGO94481
1
CBU_1127
0.52
Coxiella_sp_RIFCSPHIGHO2_12_FULL_42_15.OGO94332
Gammaproteobacteria_bacterium_RIFCSPHIGHO2_12_FULL_35_23.OGT29889
1
1
Gammaproteobacteria_bacterium_CG12_big_fil_rev_8_21_14_0_65_46_12.PIQ43642
Acidithiobacillales_bacterium_SG8_45.KPK11944
0.55
Thermoplasmatales_archaeon_SG8_52_4.KYK21749
0.99
0.73
Gammaproteobacteria_bacterium_SG8_31.KPK57612
0.77
Anaerolineaceae_bacterium_4572_322.OQY21558
1
1
Candidatus_Scalindua_rubra.ODS30507
SAR202_cluster_bacterium_Ae2_Chloro_G1.PKB60530
0.83
SAR202_cluster_bacterium_Casp_Chloro_G3.PKB57503
Candidatus_Atribacteria_bacterium_RBG_19FT_COMBO_35_14.OGD16383
1
1
Actinobacteria_bacterium_RBG_13_35_12.OFW55048
1
bacterium_TMED6.OUT38330
Coxiella_sp_DG_40.KPJ66624
1
0.81
Petrotoga.WP_103876927
Flectobacillus_sp_BAB_3569.WP_095161084
0.62
0.44
Desulfonatronum_thiodismutans.WP_084184688
0.73
Desulfonatronospira_thiodismutans.WP_008869657
0.99
Candidatus_Bathyarchaeota_archaeon_ex4484_135.OYT50754
0.97
candidate_division_Zixibacteria_bacterium_SM23_81.KPL19589
0.55
Syntrophus_aciditrophicus.WP_011418347
0.89
Deltaproteobacteria_bacterium_HGW_Deltaproteobacteria_2.PKN37950
1
Deltaproteobacteria_bacterium_HGW_Deltaproteobacteria_6.PKN18120
B
Archaea
Candidatus_Bathyarchaeota_archaeon_ex4484_135
Thermoplasmatales_archaeon_SG8-52-4
Petrotoga
Candidatus_Scalindua_rubra
Bacteria
candidate_division_Zixibacteria_bacterium_SM23_81
Flectobacillus_sp_BAB-3569
Bacteria
bacterium_TMED6
Bacteria
Candidatus_Atribacteria_bacterium_RBG_16_35_8
Actinobacteria_bacterium_RBG_13_35_12
Bacteria
Anaerolineaceae_bacterium_4572_322
Chloroflexi
Chloroflexi
SAR202_cluster_bacterium_Casp-Chloro-G3
SAR202_cluster_bacterium_Ae2-Chloro-G1
Acidithiobacillales_bacterium_SG8_45
Syntrophus_aciditrophicus_SB
Desulfonatronum_thiodismutans
Deltaproteobacteria
Desulfovibrionales
Desulfonatronospira_thiodismutans
Proteobacteria
Deltaproteobacteria_bacterium_HGW-Deltaproteobacteria-2
Deltaproteobacteria
Deltaproteobacteria_bacterium_HGW-Deltaproteobacteria-6
Coxiella_burnetii_RSA_493
Coxiella
Coxiella_sp_DG_40
Coxiella
Coxiella_sp_RIFCSPHIGHO2_12_FULL_42_15
Coxiella_sp_RIFCSPHIGHO2_12_FULL_44_14
Gammaproteobacteria
Gammaproteobacteria_bacterium_RIFCSPHIGHO2_12_FULL_35_23
Gammaproteobacteria_bacterium_RIFCSPHIGHO2_12_FULL_36_30
Gammaproteobacteria
Gammaproteobacteria_bacterium_RIFCSPHIGHO2_12_FULL_38_11
Gammaproteobacteria_bacterium_CG11_big_fil_rev_8_21_14_0_20_46_22
Gammaproteobacteria_bacterium_SG8_31

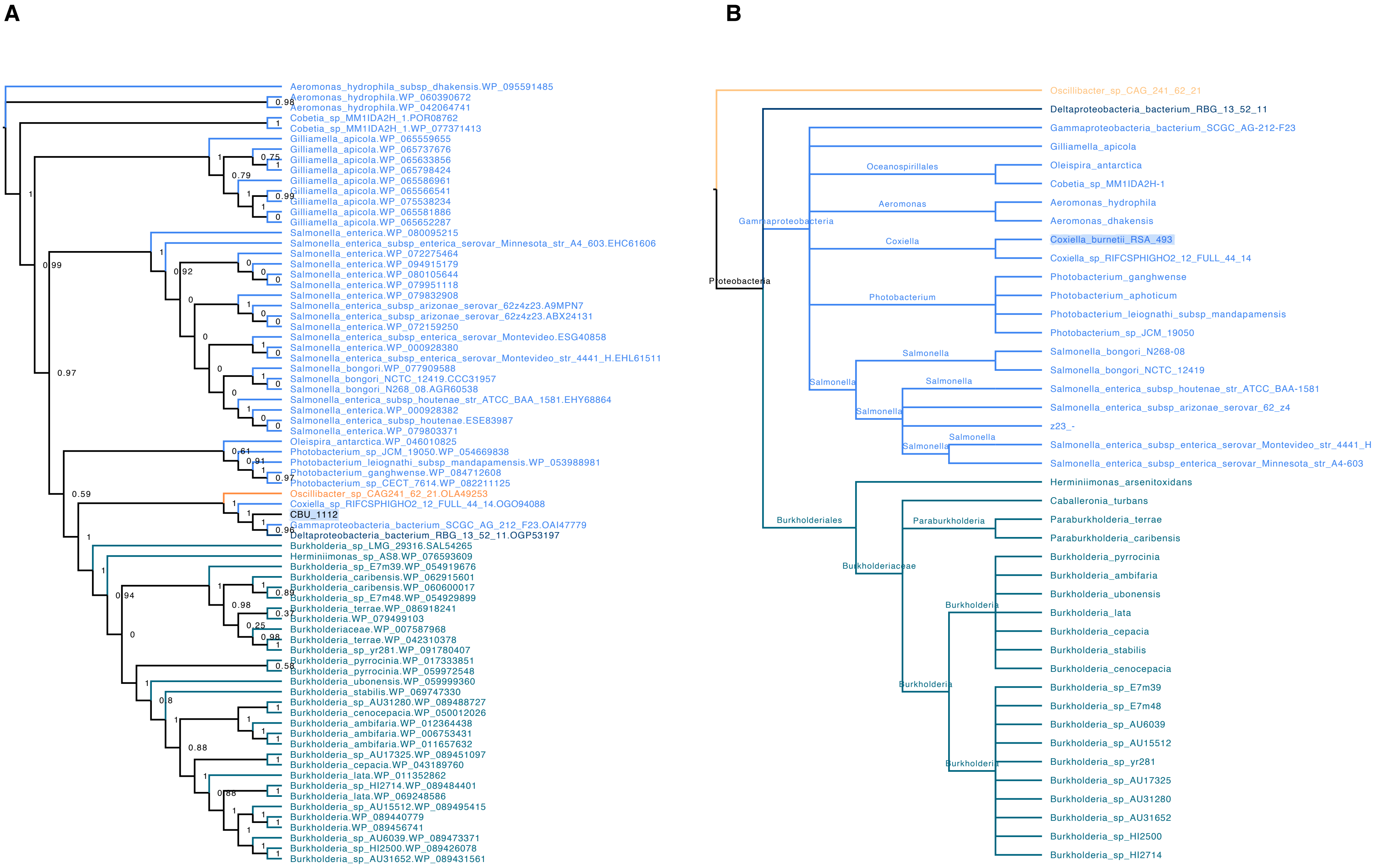
A
Aeromonas_hydrophila_subsp_dhakensis.WP_095591485
Aeromonas_hydrophila.WP_060390672
Aeromonas_hydrophila.WP_042064741
Cobetia_sp_MM1IDA2H_1.POR08762
Cobetia_sp_MM1IDA2H_1.WP_077371413
Gilliamella_apicola.WP_065559655
Gilliamella_apicola.WP_065737676
Gilliamella_apicola.WP_065633856
Gilliamella_apicola.WP_065798424
Gilliamella_apicola.WP_065586961
Gilliamella_apicola.WP_065566541
Gilliamella_apicola.WP_075538234
Gilliamella_apicola.WP_065581886
Gilliamella_apicola.WP_065652287
Salmonella_enterica.WP_080095215
Salmonella_enterica_subsp_enterica_serovar_Minnesota_str_A4_603.EHC61606
Salmonella_enterica.WP_072275464
Salmonella_enterica.WP_094915179
Salmonella_enterica.WP_080105644
Salmonella_enterica.WP_079951118
Salmonella_enterica.WP_079832908
Salmonella_enterica_subsp_arizonae_serovar_62z4z23.A9MPN7
Salmonella_enterica_subsp_arizonae_serovar_62z4z23.ABX24131
Salmonella_enterica.WP_072159250
Salmonella_enterica_subsp_enterica_serovar_Montevideo.ESG40858
Salmonella_enterica.WP_000928380
Salmonella_enterica_subsp_enterica_serovar_Montevideo_str_4441_H.EHL61511
Salmonella_bongori.WP_077909588
Salmonella_bongori_NCTC_12419.CCC31957
Salmonella_bongori_N268_08.AGR60538
Salmonella_enterica_subsp_houtenae_str_ATCC_BAA_1581.EHY68864
Salmonella_enterica.WP_000928382
Salmonella_enterica_subsp_houtenae.ESE83987
Salmonella_enterica.WP_079803371
Oleispira_antarctica.WP_046010825
Photobacterium_sp_JCM_19050.WP_054669838
Photobacterium_leiognathi_subsp_mandapamensis.WP_053988981
Photobacterium_ganghwense.WP_084712608
Photobacterium_sp_CECT_7614.WP_082211125
Oscillibacter_sp_CAG241_62_21.OLA49253
Coxiella_sp_RIFCSPHIGHO2_12_FULL_44_14.OGO94088
CBU_1112
Gammaproteobacteria_bacterium_SCGC_AG_212_F23.OAI47779
Deltaproteobacteria_bacterium_RBG_13_52_11.OGP53197
Burkholderia_sp_LMG_29316.SAL54265
Herminiimonas_sp_AS8.WP_076593609
Burkholderia_sp_E7m39.WP_054919676
Burkholderia_caribensis.WP_062915601
Burkholderia_caribensis.WP_060600017
Burkholderia_sp_E7m48.WP_054929899
Burkholderia_terrae.WP_086918241
Burkholderia.WP_079499103
Burkholderiaceae.WP_007587968
Burkholderia_terrae.WP_042310378
Burkholderia_sp_yr281.WP_091780407
Burkholderia_pyrrocinia.WP_017333851
Burkholderia_pyrrocinia.WP_059972548
Burkholderia_ubonensis.WP_059999360
Burkholderia_stabilis.WP_069747330
Burkholderia_sp_AU31280.WP_089488727
Burkholderia_cenocepacia.WP_050012026
Burkholderia_ambifaria.WP_012364438
Burkholderia_ambifaria.WP_006753431
Burkholderia_ambifaria.WP_011657632
Burkholderia_sp_AU17325.WP_089451097
Burkholderia_cepacia.WP_043189760
Burkholderia_lata.WP_011352862
Burkholderia_sp_HI2714.WP_089484401
Burkholderia_lata.WP_069248586
Burkholderia_sp_AU15512.WP_089495415
Burkholderia.WP_089440779
Burkholderia.WP_089456741
Burkholderia_sp_AU6039.WP_089473371
Burkholderia_sp_HI2500.WP_089426078
Burkholderia_sp_AU31652.WP_089431561
B
Oscillibacter_sp_CAG_241_62_21
Deltaproteobacteria_bacterium_RBG_13_52_11
Gammaproteobacteria_bacterium_SCGC_AG-212-F23
Gilliamella_apicola
Oceanospirillales
Oleispira_antarctica
Cobetia_sp_MM1IDA2H-1
Aeromonas
Aeromonas_hydrophila
Aeromonas_dhakensis
Gammaproteobacteria
Coxiella
Coxiella_burnetii_RSA_493
Coxiella_sp_RIFCSPHIGHO2_12_FULL_44_14
Proteobacteria
Photobacterium
Photobacterium_ganghwense
Photobacterium_aphoticum
Photobacterium_leiognathi_subsp_mandapamensis
Photobacterium_sp_JCM_19050
Salmonella
Salmonella_bongori_N268-08
Salmonella_bongori_NCTC_12419
Salmonella_enterica_subsp_houtenae_str_ATCC_BAA-1581
Salmonella_enterica_subsp_arizonae_serovar_62_z4
z23_-
Salmonella_enterica_subsp_enterica_serovar_Montevideo_str_4441_H
Salmonella_enterica_subsp_enterica_serovar_Minnesota_str_A4-603
Herminiimonas_arsenitoxidans
Caballeronia_turbans
Burkholderiales
Paraburkholderia
Paraburkholderia_terrae
Paraburkholderia_caribensis
Burkholderiaceae
Burkholderia_pyrrocinia
Burkholderia_ambifaria
Burkholderia_ubonensis
Burkholderia
Burkholderia_lata
Burkholderia_cepacia
Burkholderia_stabilis
Burkholderia_cenocepacia
Burkholderia_sp_E7m39
Burkholderia_sp_E7m48
Burkholderia_sp_AU6039
Burkholderia_sp_AU15512
Burkholderia_sp_yr281
Burkholderia_sp_AU17325
Burkholderia_sp_AU31280
Burkholderia_sp_AU31652
Burkholderia_sp_HI2500
Burkholderia_sp_HI2714

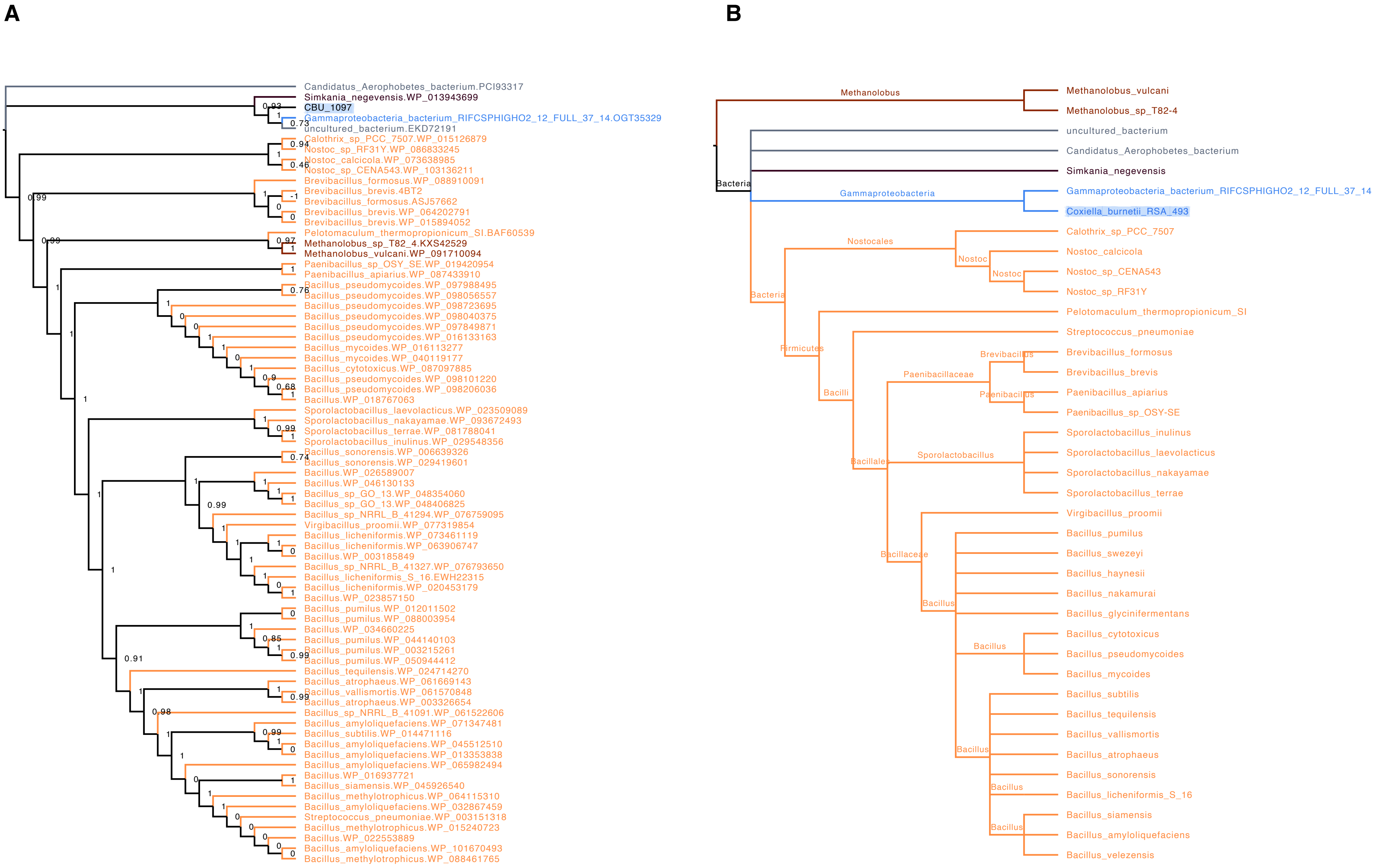
A
Candidatus_Aerophobetes_bacterium.PCI93317
Simkania_negevensis.WP_013943699
CBU_1097
Gammaproteobacteria_bacterium_RIFCSPHIGHO2_12_FULL_37_14.OGT35329
uncultured_bacterium.EKD72191
Calothrix_sp_PCC_7507.WP_015126879
Nostoc_sp_RF31Y.WP_086833245
Nostoc_calcicola.WP_073638985
Nostoc_sp_CENA543.WP_103136211
Brevibacillus_formosus.WP_088910091
Brevibacillus_brevis.4BT2
Brevibacillus_formosus.ASJ57662
Brevibacillus_brevis.WP_064202791
Brevibacillus_brevis.WP_015894052
Pelotomaculum_thermopropionicum_SI.BAF60539
Methanolobus_sp_T82_4.KXS42529
Methanolobus_vulcani.WP_091710094
Paenibacillus_sp_OSY_SE.WP_019420954
Paenibacillus_apiarius.WP_087433910
Bacillus_pseudomycoides.WP_097988495
Bacillus_pseudomycoides.WP_098056557
Bacillus_pseudomycoides.WP_098723695
Bacillus_pseudomycoides.WP_098040375
Bacillus_pseudomycoides.WP_097849871
Bacillus_pseudomycoides.WP_016133163
Bacillus_mycoides.WP_016113277
Bacillus_mycoides.WP_040119177
Bacillus_cytotoxicus.WP_087097885
Bacillus_pseudomycoides.WP_098101220
Bacillus_pseudomycoides.WP_098206036
Bacillus.WP_018767063
Sporolactobacillus_laevolacticus.WP_023509089
Sporolactobacillus_nakayamae.WP_093672493
Sporolactobacillus_terrae.WP_081788041
Sporolactobacillus_inulinus.WP_029548356
Bacillus_sonorensis.WP_006639326
Bacillus_sonorensis.WP_029419601
Bacillus.WP_026589007
Bacillus.WP_046130133
Bacillus_sp_GO_13.WP_048354060
Bacillus_sp_GO_13.WP_048406825
Bacillus_sp_NRRL_B_41294.WP_076759095
Virgibacillus_proomii.WP_077319854
Bacillus_licheniformis.WP_073461119
Bacillus_licheniformis.WP_063906747
Bacillus.WP_003185849
Bacillus_sp_NRRL_B_41327.WP_076793650
Bacillus_licheniformis_S_16.EWH22315
Bacillus_licheniformis.WP_020453179
Bacillus.WP_023857150
Bacillus_pumilus.WP_012011502
Bacillus_pumilus.WP_088003954
Bacillus.WP_034660225
Bacillus_pumilus.WP_044140103
Bacillus_pumilus.WP_003215261
Bacillus_pumilus.WP_050944412
Bacillus_tequilensis.WP_024714270
Bacillus_atrophaeus.WP_061669143
Bacillus_vallismortis.WP_061570848
Bacillus_atrophaeus.WP_003326654
Bacillus_sp_NRRL_B_41091.WP_061522606
Bacillus_amyloliquefaciens.WP_071347481
Bacillus_subtilis.WP_014471116
Bacillus_amyloliquefaciens.WP_045512510
Bacillus_amyloliquefaciens.WP_013353838
Bacillus_amyloliquefaciens.WP_065982494
Bacillus.WP_016937721
Bacillus_siamensis.WP_045926540
Bacillus_methylotrophicus.WP_064115310
Bacillus_amyloliquefaciens.WP_032867459
Streptococcus_pneumoniae.WP_003151318
Bacillus_methylotrophicus.WP_015240723
Bacillus.WP_022553889
Bacillus_amyloliquefaciens.WP_101670493
Bacillus_methylotrophicus.WP_088461765
B
Methanolobus
Methanolobus_vulcani
Methanolobus_sp_T82-4
uncultured_bacterium
Candidatus_Aerophobetes_bacterium
Simkania_negevensis
Bacteria
Gammaproteobacteria
Gammaproteobacteria_bacterium_RIFCSPHIGHO2_12_FULL_37_14
Coxiella_burnetii_RSA_493
Nostocales
Calothrix_sp_PCC_7507
Nostoc
Nostoc_calcicola
Nostoc
Nostoc_sp_CENA543
Nostoc_sp_RF31Y
Bacteria
Pelotomaculum_thermopropionicum_SI
Streptococcus_pneumoniae
Firmicutes
Brevibacillus
Brevibacillus_formosus
Brevibacillus_brevis
Paenibacillaceae
Paenibacillus
Paenibacillus_apiarius
Paenibacillus_sp_OSY-SE
Bacilli
Sporolactobacillus_inulinus
Sporolactobacillus_laevolacticus
Bacillales
Sporolactobacillus
Sporolactobacillus_nakayamae
Sporolactobacillus_terrae
Virgibacillus_proomii
Bacillus_pumilus
Bacillaceae
Bacillus_swezeyi
Bacillus_haynesii
Bacillus_nakamurai
Bacillus
Bacillus_glycinifermentans
Bacillus_cytotoxicus
Bacillus
Bacillus_pseudomycoides
Bacillus_mycoides
Bacillus_subtilis
Bacillus_tequilensis
Bacillus_vallismortis
Bacillus
Bacillus_atrophaeus
Bacillus_sonorensis
Bacillus
Bacillus_licheniformis_S_16
Bacillus_siamensis
Bacillus
Bacillus_amyloliquefaciens
Bacillus_velezensis

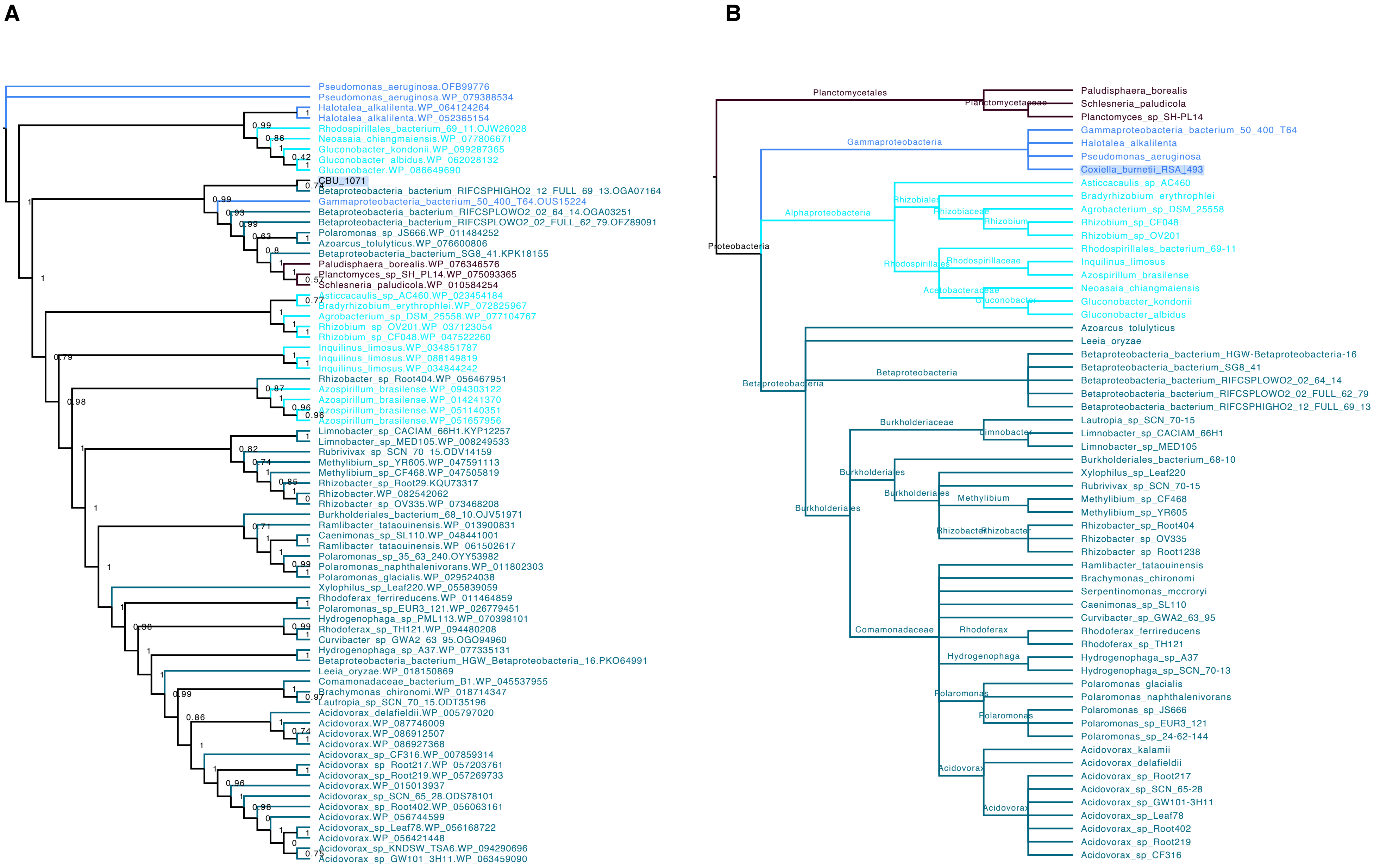
A
Pseudomonas_aeruginosa.OFB99776
Pseudomonas_aeruginosa.WP_079388534
Halotalea_alkalilenta.WP_064124264
Halotalea_alkalilenta.WP_052365154
Rhodospirillales_bacterium_69_11.OJW26028
Neoasaia_chiangmaiensis.WP_077806671
Gluconobacter_kondonii.WP_099287365
Gluconobacter_albidus.WP_062028132
Gluconobacter.WP_086649690
CBU_1071
Betaproteobacteria_bacterium_RIFCSPHIGHO2_12_FULL_69_13.OGA07164
Gammaproteobacteria_bacterium_50_400_T64.OUS15224
Betaproteobacteria_bacterium_RIFCSPLOWO2_02_64_14.OGA03251
Betaproteobacteria_bacterium_RIFCSPLOWO2_02_FULL_62_79.OFZ89091
Polaromonas_sp_JS666.WP_011484252
Azoarcus_tolulyticus.WP_076600806
Betaproteobacteria_bacterium_SG8_41.KPK18155
Paludisphaera_borealis.WP_076346576
Planctomyces_sp_SH_PL14.WP_075093365
Schlesneria_paludicola.WP_010584254
Asticcacaulis_sp_AC460.WP_023454184
Bradyrhizobium_erythrophlei.WP_072825967
Agrobacterium_sp_DSM_25558.WP_077104767
Rhizobium_sp_OV201.WP_037123054
Rhizobium_sp_CF048.WP_047522260
Inquilinus_limosus.WP_034851787
Inquilinus_limosus.WP_088149819
Inquilinus_limosus.WP_034844242
Rhizobacter_sp_Root404.WP_056467951
Azospirillum_brasilense.WP_094303122
Azospirillum_brasilense.WP_014241370
Azospirillum_brasilense.WP_051140351
Azospirillum_brasilense.WP_051657956
Limnobacter_sp_CACIAM_66H1.KYP12257
Limnobacter_sp_MED105.WP_008249533
Rubrivivax_sp_SCN_70_15.ODV14159
Methylibium_sp_YR605.WP_047591113
Methylibium_sp_CF468.WP_047505819
Rhizobacter_sp_Root29.KQU73317
Rhizobacter.WP_082542062
Rhizobacter_sp_OV335.WP_073468208
Burkholderiales_bacterium_68_10.OJV51971
Ramlibacter_tataouinensis.WP_013900831
Caenimonas_sp_SL110.WP_048441001
Ramlibacter_tataouinensis.WP_061502617
Polaromonas_sp_35_63_240.OYY53982
Polaromonas_naphthalenivorans.WP_011802303
Polaromonas_glacialis.WP_029524038
Xylophilus_sp_Leaf220.WP_055839059
Rhodoferax_ferrireducens.WP_011464859
Polaromonas_sp_EUR3_121.WP_026779451
Hydrogenophaga_sp_PML113.WP_070398101
Rhodoferax_sp_TH121.WP_094480208
Curvibacter_sp_GWA2_63_95.OGO94960
Hydrogenophaga_sp_A37.WP_077335131
Betaproteobacteria_bacterium_HGW_Betaproteobacteria_16.PKO64991
Leeia_oryzae.WP_018150869
Comamonadaceae_bacterium_B1.WP_045537955
Brachymonas_chironomi.WP_018714347
Lautropia_sp_SCN_70_15.ODT35196
Acidovorax_delafieldii.WP_005797020
Acidovorax.WP_087746009
Acidovorax.WP_086912507
Acidovorax.WP_086927368
Acidovorax_sp_CF316.WP_007859314
Acidovorax_sp_Root217.WP_057203761
Acidovorax_sp_Root219.WP_057269733
Acidovorax.WP_015013937
Acidovorax_sp_SCN_65_28.ODS78101
Acidovorax_sp_Root402.WP_056063161
Acidovorax.WP_056744599
Acidovorax_sp_Leaf78.WP_056168722
Acidovorax.WP_056421448
Acidovorax_sp_KNDSW_TSA6.WP_094290696
Acidovorax_sp_GW101_3H11.WP_063459090
B
Planctomycetales
Planctomycetaceae
Paludisphaera_borealis
Schlesneria_paludicola
Planctomyces_sp_SH-PL14
Proteobacteria
Gammaproteobacteria
Gammaproteobacteria_bacterium_50_400_T64
Halotalea_alkalilenta
Pseudomonas_aeruginosa
Coxiella_burnetii_RSA_493
Alphaproteobacteria
Rhizobiales
Rhizobiaceae
Rhizobium
Asticcacaulis_sp_AC460
Bradyrhizobium_erythrophlei
Agrobacterium_sp_DSM_25558
Rhizobium_sp_CF048
Rhizobium_sp_OV201
Rhodospirillales
Rhodospirillaceae
Acetobacteraceae
Gluconobacter
Rhodospirillales_bacterium_69-11
Inquilinus_limosus
Azospirillum_brasilense
Neoasaia_chiangmaiensis
Gluconobacter_kondonii
Gluconobacter_albidus
Betaproteobacteria
Azoarcus_tolulyticus
Leeia_oryzae
Betaproteobacteria_bacterium_HGW-Betaproteobacteria-16
Betaproteobacteria_bacterium_SG8_41
Betaproteobacteria_bacterium_RIFCSPLOWO2_02_64_14
Betaproteobacteria_bacterium_RIFCSPLOWO2_02_FULL_62_79
Betaproteobacteria_bacterium_RIFCSPHIGHO2_12_FULL_69_13
Burkholderiales
Burkholderiaceae
Limnobacter
Lautropia_sp_SCN_70-15
Limnobacter_sp_CACIAM_66H1
Limnobacter_sp_MED105
Burkholderiales_bacterium_68-10
Methylibium
Rhizobacter
Xylophilus_sp_Leaf220
Rubrivivax_sp_SCN_70-15
Methylibium_sp_CF468
Methylibium_sp_YR605
Rhizobacter_sp_Root404
Rhizobacter_sp_OV335
Rhizobacter_sp_Root1238
Comamonadaceae
Rhodoferax
Hydrogenophaga
Polaromonas
Acidovorax
Ramlibacter_tataouinensis
Brachymonas_chironomi
Serpentinomonas_mccroryi
Caenimonas_sp_SL110
Curvibacter_sp_GWA2_63_95
Rhodoferax_ferrireducens
Rhodoferax_sp_TH121
Hydrogenophaga_sp_A37
Hydrogenophaga_sp_SCN_70-13
Polaromonas_glacialis
Polaromonas_naphthalenivorans
Polaromonas_sp_JS666
Polaromonas_sp_EUR3_121
Polaromonas_sp_24-62-144
Acidovorax_kalamii
Acidovorax_delafieldii
Acidovorax_sp_Root217
Acidovorax_sp_SCN_65-28
Acidovorax_sp_GW101-3H11
Acidovorax_sp_Leaf78
Acidovorax_sp_Root402
Acidovorax_sp_Root219
Acidovorax_sp_CF316

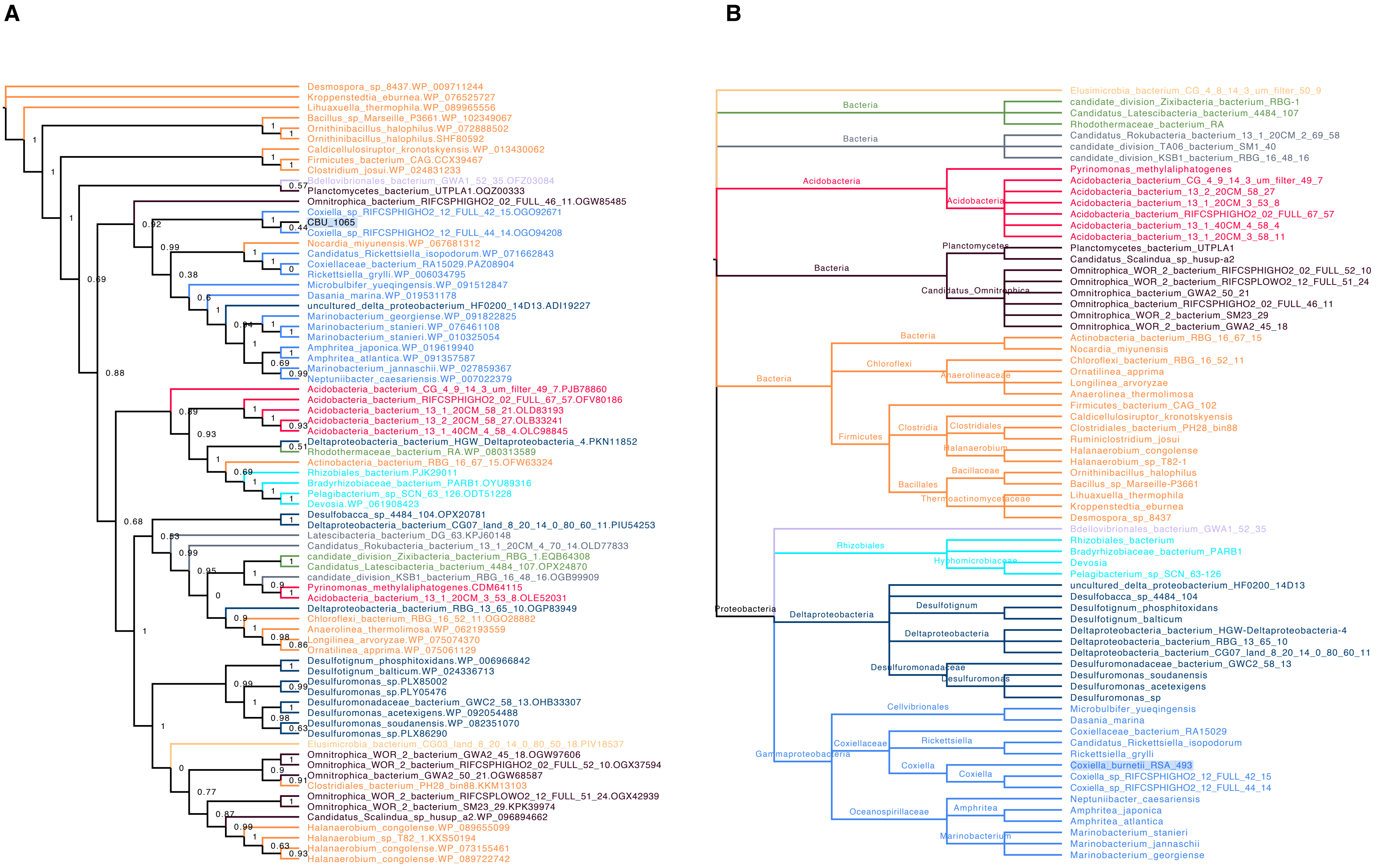
A
Desmospora_sp_8437.WP_009711244
Kroppenstedtia_eburnea.WP_076525727
Lihuaxuella_thermophila.WP_089965556
Bacillus_sp_Marseille_P3661.WP_102349067
Ornithinibacillus_halophilus.WP_072888502
Ornithinibacillus_halophilus.SHF80592
Caldicellulosiruptor_kronotskyensis.WP_013430062
Firmicutes_bacterium_CAG.CCX39467
Clostridium_josui.WP_024831233
Bdellovibrionales_bacterium_GWA1_52_35.OFZ03084
Planctomycetes_bacterium_UTPLA1.OQZ00333
Omnitrophica_bacterium_RIFCSPHIGHO2_02_FULL_46_11.OGW85485
Coxiella_sp_RIFCSPHIGHO2_12_FULL_42_15.OGO92671
CBU_1065
Coxiella_sp_RIFCSPHIGHO2_12_FULL_44_14.OGO94208
Nocardia_miyunensis.WP_067681312
Candidatus_Rickettsiella_isopodorum.WP_071662843
Coxiellaceae_bacterium_RA15029.PAZ08904
Rickettsiella_grylli.WP_006034795
Microbulbifer_yueqingensis.WP_091512847
Dasania_marina.WP_019531178
uncultured_delta_proteobacterium_HF0200_14D13.ADI19227
Marinobacterium_georgiense.WP_091822825
Marinobacterium_stanieri.WP_076461108
Marinobacterium_stanieri.WP_010325054
Amphritea_japonica.WP_019619940
Amphritea_atlantica.WP_091357587
Marinobacterium_jannaschii.WP_027859367
Neptuniibacter_caesariensis.WP_007022379
Acidobacteria_bacterium_CG_4_9_14_3_um_filter_49_7.PJB78860
Acidobacteria_bacterium_RIFCSPHIGHO2_02_FULL_67_57.OFV80186
Acidobacteria_bacterium_13_1_20CM_58_21.OLD83193
Acidobacteria_bacterium_13_2_20CM_58_27.OLB33241
Acidobacteria_bacterium_13_1_40CM_4_58_4.OLC98845
Deltaproteobacteria_bacterium_HGW_Deltaproteobacteria_4.PKN11852
Rhodothermaceae_bacterium_RA.WP_080313589
Actinobacteria_bacterium_RBG_16_67_15.OFW63324
Rhizobiales_bacterium.PJK29011
Bradyrhizobiaceae_bacterium_PARB1.OYU89316
Pelagibacterium_sp_SCN_63_126.ODT51228
Devosia.WP_061908423
Desulfobacca_sp_4484_104.OPX20781
Deltaproteobacteria_bacterium_CG07_land_8_20_14_0_80_60_11.PIU54253
Latescibacteria_bacterium_DG_63.KPJ60148
Candidatus_Rokubacteria_bacterium_13_1_20CM_4_70_14.OLD77833
candidate_division_Zixibacteria_bacterium_RBG_1.EQB64308
Candidatus_Latescibacteria_bacterium_4484_107.OPX24870
candidate_division_KSB1_bacterium_RBG_16_48_16.OGB99909
Pyrinomonas_methylaliphatogenes.CDM64115
Acidobacteria_bacterium_13_1_20CM_3_53_8.OLE52031
Deltaproteobacteria_bacterium_RBG_13_65_10.OGP83949
Chloroflexi_bacterium_RBG_16_52_11.OGO28882
Anaerolinea_thermolimosa.WP_062193559
Longilinea_arvoryzae.WP_075074370
Ornatilinea_apprima.WP_075061129
Desulfotignum_phosphitoxidans.WP_006966842
Desulfotignum_balticum.WP_024336713
Desulfuromonas_sp.PLX85002
Desulfuromonas_sp.PLY05476
Desulfuromonadaceae_bacterium_GWC2_58_13.OHB33307
Desulfuromonas_acetexigens.WP_092054488
Desulfuromonas_soudanensis.WP_082351070
Desulfuromonas_sp.PLX86290
Elusimicrobia_bacterium_CG03_land_8_20_14_0_80_50_18.PIV18537
Omnitrophica_WOR_2_bacterium_GWA2_45_18.OGW97606
Omnitrophica_WOR_2_bacterium_RIFCSPHIGHO2_02_FULL_52_10.OGX37594
Omnitrophica_bacterium_GWA2_50_21.OGW68587
Clostridiales_bacterium_PH28_bin88.KKM13103
Omnitrophica_WOR_2_bacterium_RIFCSPLOWO2_12_FULL_51_24.OGX42939
Omnitrophica_WOR_2_bacterium_SM23_29.KPK39974
Candidatus_Scalindua_sp_husup_a2.WP_096894662
Halanaerobium_congolense.WP_089655099
Halanaerobium_sp_T82_1.KXS50194
Halanaerobium_congolense.WP_073155461
Halanaerobium_congolense.WP_089722742
B
Elusimicrobia_bacterium_CG_4_8_14_3_um_filter_50_9
Bacteria
candidate_division_Zixibacteria_bacterium_RBG-1
Candidatus_Latescibacteria_bacterium_4484_107
Rhodothermaceae_bacterium_RA
Bacteria
Candidatus_Rokubacteria_bacterium_13_1_20CM_2_69_58
candidate_division_TA06_bacterium_SM1_40
candidate_division_KSB1_bacterium_RBG_16_48_16
Acidobacteria
Pyrinomonas_methylaliphatogenes
Acidobacteria
Acidobacteria_bacterium_CG_4_9_14_3_um_filter_49_7
Acidobacteria_bacterium_13_2_20CM_58_27
Acidobacteria_bacterium_13_1_20CM_3_53_8
Acidobacteria_bacterium_RIFCSPHIGHO2_02_FULL_67_57
Acidobacteria_bacterium_13_1_40CM_4_58_4
Acidobacteria_bacterium_13_1_20CM_3_58_11
Planctomycetes
Planctomycetes_bacterium_UTPLA1
Candidatus_Scalindua_sp_husup-a2
Bacteria
Candidatus_Omnitrophica
Omnitrophica_WOR_2_bacterium_RIFCSPHIGHO2_02_FULL_52_10
Omnitrophica_WOR_2_bacterium_RIFCSPLOWO2_12_FULL_51_24
Omnitrophica_bacterium_GWA2_50_21
Omnitrophica_bacterium_RIFCSPHIGHO2_02_FULL_46_11
Omnitrophica_WOR_2_bacterium_SM23_29
Omnitrophica_WOR_2_bacterium_GWA2_45_18
Bacteria
Actinobacteria_bacterium_RBG_16_67_15
Nocardia_miyunensis
Bacteria
Chloroflexi
Chloroflexi_bacterium_RBG_16_52_11
Anaerolineaceae
Ornatilinea_apprima
Longilinea_arvoryzae
Anaerolinea_thermolimosa
Firmicutes_bacterium_CAG_102
Clostridia
Clostridiales
Caldicellulosiruptor_kronotskyensis
Clostridiales_bacterium_PH28_bin88
Ruminiclostridium_josui
Firmicutes
Halanaerobium
Halanaerobium_congolense
Halanaerobium_sp_T82-1
Bacillaceae
Ornithinibacillus_halophilus
Bacillus_sp_Marseille-P3661
Bacillales
Thermoactinomycetaceae
Lihuaxuella_thermophila
Kroppenstedtia_eburnea
Desmospora_sp_8437
Bdellovibrionales_bacterium_GWA1_52_35
Rhizobiales
Rhizobiales_bacterium
Bradyrhizobiaceae_bacterium_PARB1
Hyphomicrobiaceae
Devosia
Pelagibacterium_sp_SCN_63-126
uncultured_delta_proteobacterium_HF0200_14D13
Desulfobacca_sp_4484_104
Desulfotignum
Desulfotignum_phosphitoxidans
Desulfotignum_balticum
Proteobacteria
Deltaproteobacteria
Deltaproteobacteria
Deltaproteobacteria_bacterium_HGW-Deltaproteobacteria-4
Deltaproteobacteria_bacterium_RBG_13_65_10
Deltaproteobacteria_bacterium_CG07_land_8_20_14_0_80_60_11
Desulfuromonadaceae
Desulfuromonadaceae_bacterium_GWC2_58_13
Desulfuromonas
Desulfuromonas_soudanensis
Desulfuromonas_acetexigens
Desulfuromonas_sp
Cellvibrionales
Microbulbifer_yueqingensis
Dasania_marina
Rickettsiella
Coxiellaceae_bacterium_RA15029
Coxiellaceae
Candidatus_Rickettsiella_isopodorum
Rickettsiella_grylli
Gammaproteobacteria
Coxiella
Coxiella_burnetii_RSA_493
Coxiella
Coxiella_sp_RIFCSPHIGHO2_12_FULL_42_15
Coxiella_sp_RIFCSPHIGHO2_12_FULL_44_14
Neptuniibacter_caesariensis
Amphritea
Amphritea_japonica
Oceanospirillaceae
Amphritea_atlantica
Marinobacterium
Marinobacterium_stanieri
Marinobacterium_jannaschii
Marinobacterium_georgiense

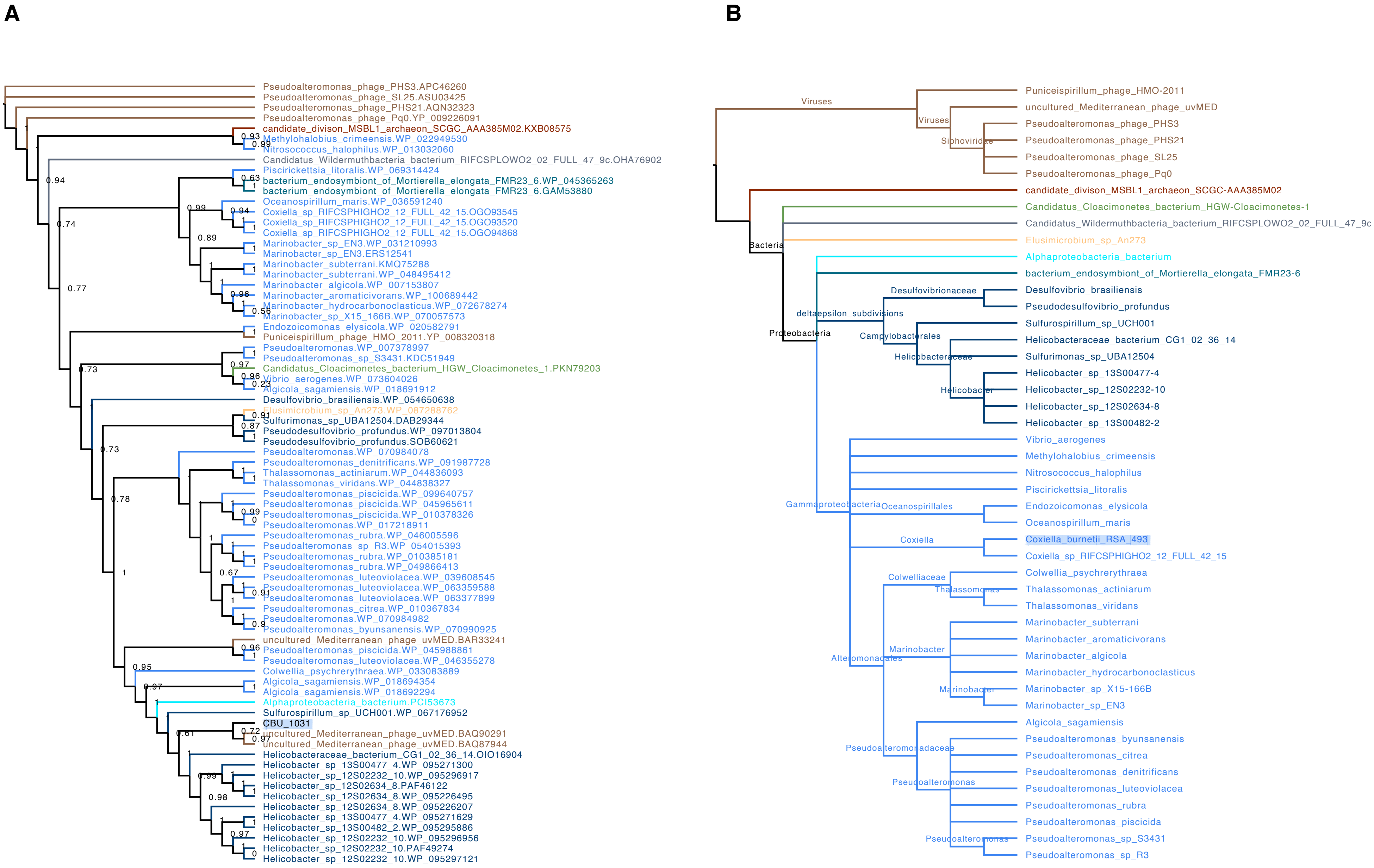
A
Pseudoalteromonas_phage_PHS3.APC46260
Pseudoalteromonas_phage_SL25.ASU03425
Pseudoalteromonas_phage_PHS21.AQN32323
Pseudoalteromonas_phage_Pq0.YP_009226091
candidate_divison_MSBL1_archaeon_SCGC_AAA385M02.KXB08575
Methylohalobius_crimeensis.WP_022949530
Nitrosococcus_halophilus.WP_013032060
Candidatus_Wildermuthbacteria_bacterium_RIFCSPLOWO2_02_FULL_47_9c.OHA76902
Piscirickettsia_litoralis.WP_069314424
bacterium_endosymbiont_of_Mortierella_elongata_FMR23_6.WP_045365263
bacterium_endosymbiont_of_Mortierella_elongata_FMR23_6.GAM53880
Oceanospirillum_maris.WP_036591240
Coxiella_sp_RIFCSPHIGHO2_12_FULL_42_15.OGO93545
Coxiella_sp_RIFCSPHIGHO2_12_FULL_42_15.OGO93520
Coxiella_sp_RIFCSPHIGHO2_12_FULL_42_15.OGO94868
Marinobacter_sp_EN3.WP_031210993
Marinobacter_sp_EN3.ERS12541
Marinobacter_subterrani.KMQ75288
Marinobacter_subterrani.WP_048495412
Marinobacter_algicola.WP_007153807
Marinobacter_aromaticivorans.WP_100689442
Marinobacter_hydrocarbonoclasticus.WP_072678274
Marinobacter_sp_X15_166B.WP_070057573
Endozoicomonas_elysicola.WP_020582791
Puniceispirillum_phage_HMO_2011.YP_008320318
Pseudoalteromonas.WP_007378997
Pseudoalteromonas_sp_S3431.KDC51949
Candidatus_Cloacimonetes_bacterium_HGW_Cloacimonetes_1.PKN79203
Vibrio_aerogenes.WP_073604026
Algicola_sagamiensis.WP_018691912
Desulfovibrio_brasiliensis.WP_054650638
Elusimicrobium_sp_An273.WP_087288762
Sulfurimonas_sp_UBA12504.DAB29344
Pseudodesulfovibrio_profundus.WP_097013804
Pseudodesulfovibrio_profundus.SOB60621
Pseudoalteromonas.WP_070984078
Pseudoalteromonas_denitrificans.WP_091987728
Thalassomonas_actiniarum.WP_044836093
Thalassomonas_viridans.WP_044838327
Pseudoalteromonas_piscicida.WP_099640757
Pseudoalteromonas_piscicida.WP_045965611
Pseudoalteromonas_piscicida.WP_010378326
Pseudoalteromonas.WP_017218911
Pseudoalteromonas_rubra.WP_046005596
Pseudoalteromonas_sp_R3.WP_054015393
Pseudoalteromonas_rubra.WP_010385181
Pseudoalteromonas_rubra.WP_049866413
Pseudoalteromonas_luteoviolacea.WP_039608545
Pseudoalteromonas_luteoviolacea.WP_063359588
Pseudoalteromonas_luteoviolacea.WP_063377899
Pseudoalteromonas_citrea.WP_010367834
Pseudoalteromonas.WP_070984982
Pseudoalteromonas_byunsanensis.WP_070990925
uncultured_Mediterranean_phage_uvMED.BAR33241
Pseudoalteromonas_piscicida.WP_045988861
Pseudoalteromonas_luteoviolacea.WP_046355278
Colwellia_psychrerythraea.WP_033083889
Algicola_sagamiensis.WP_018694354
Algicola_sagamiensis.WP_018692294
Alphaproteobacteria_bacterium.PCI53673
Sulfurospirillum_sp_UCH001.WP_067176952
CBU_1031
uncultured_Mediterranean_phage_uvMED.BAQ90291
uncultured_Mediterranean_phage_uvMED.BAQ87944
Helicobacteraceae_bacterium_CG1_02_36_14.OIO16904
Helicobacter_sp_13S00477_4.WP_095271300
Helicobacter_sp_12S02232_10.WP_095296917
Helicobacter_sp_12S02634_8.PAF46122
Helicobacter_sp_12S02634_8.WP_095226495
Helicobacter_sp_12S02634_8.WP_095226207
Helicobacter_sp_13S00477_4.WP_095271629
Helicobacter_sp_13S00482_2.WP_095295886
Helicobacter_sp_12S02232_10.WP_095296956
Helicobacter_sp_12S02232_10.PAF49274
Helicobacter_sp_12S02232_10.WP_095297121
B
Viruses
Siphoviridae
Bacteria
Proteobacteria
deltaepsilon_subdivisions
Desulfovibrionaceae
Campylobacterales
Helicobacteraceae
Helicobacter
Gammaproteobacteria
Oceanospirillales
Coxiella
Colwelliaceae
Thalassomonas
Alteromonadales
Marinobacter
Pseudoalteromonadaceae
Pseudoalteromonas
Puniceispirillum_phage_HMO-2011
uncultured_Mediterranean_phage_uvMED
Pseudoalteromonas_phage_PHS3
Pseudoalteromonas_phage_PHS21
Pseudoalteromonas_phage_SL25
Pseudoalteromonas_phage_Pq0
candidate_divison_MSBL1_archaeon_SCGC-AAA385M02
Candidatus_Cloacimonetes_bacterium_HGW-Cloacimonetes-1
Candidatus_Wildermuthbacteria_bacterium_RIFCSPLOWO2_02_FULL_47_9c
Elusimicrobium_sp_An273
Alphaproteobacteria_bacterium
bacterium_endosymbiont_of_Mortierella_elongata_FMR23-6
Desulfovibrio_brasiliensis
Pseudodesulfovibrio_profundus
Sulfurospirillum_sp_UCH001
Helicobacteraceae_bacterium_CG1_02_36_14
Sulfurimonas_sp_UBA12504
Helicobacter_sp_13S00477-4
Helicobacter_sp_12S02232-10
Helicobacter_sp_12S02634-8
Helicobacter_sp_13S00482-2
Vibrio_aerogenes
Methylohalobius_crimeensis
Nitrosococcus_halophilus
Piscirickettsia_litoralis
Endozoicomonas_elysicola
Oceanospirillum_maris
Coxiella_burnetii_RSA_493
Coxiella_sp_RIFCSPHIGHO2_12_FULL_42_15
Colwellia_psychrerythraea
Thalassomonas_actiniarum
Thalassomonas_viridans
Marinobacter_subterrani
Marinobacter_aromaticivorans
Marinobacter_algicola
Marinobacter_hydrocarbonoclasticus
Marinobacter_sp_X15-166B
Marinobacter_sp_EN3
Algicola_sagamiensis
Pseudoalteromonas_byunsanensis
Pseudoalteromonas_citrea
Pseudoalteromonas_denitrificans
Pseudoalteromonas_luteoviolacea
Pseudoalteromonas_rubra
Pseudoalteromonas_piscicida
Pseudoalteromonas_sp_S3431
Pseudoalteromonas_sp_R3

A

B

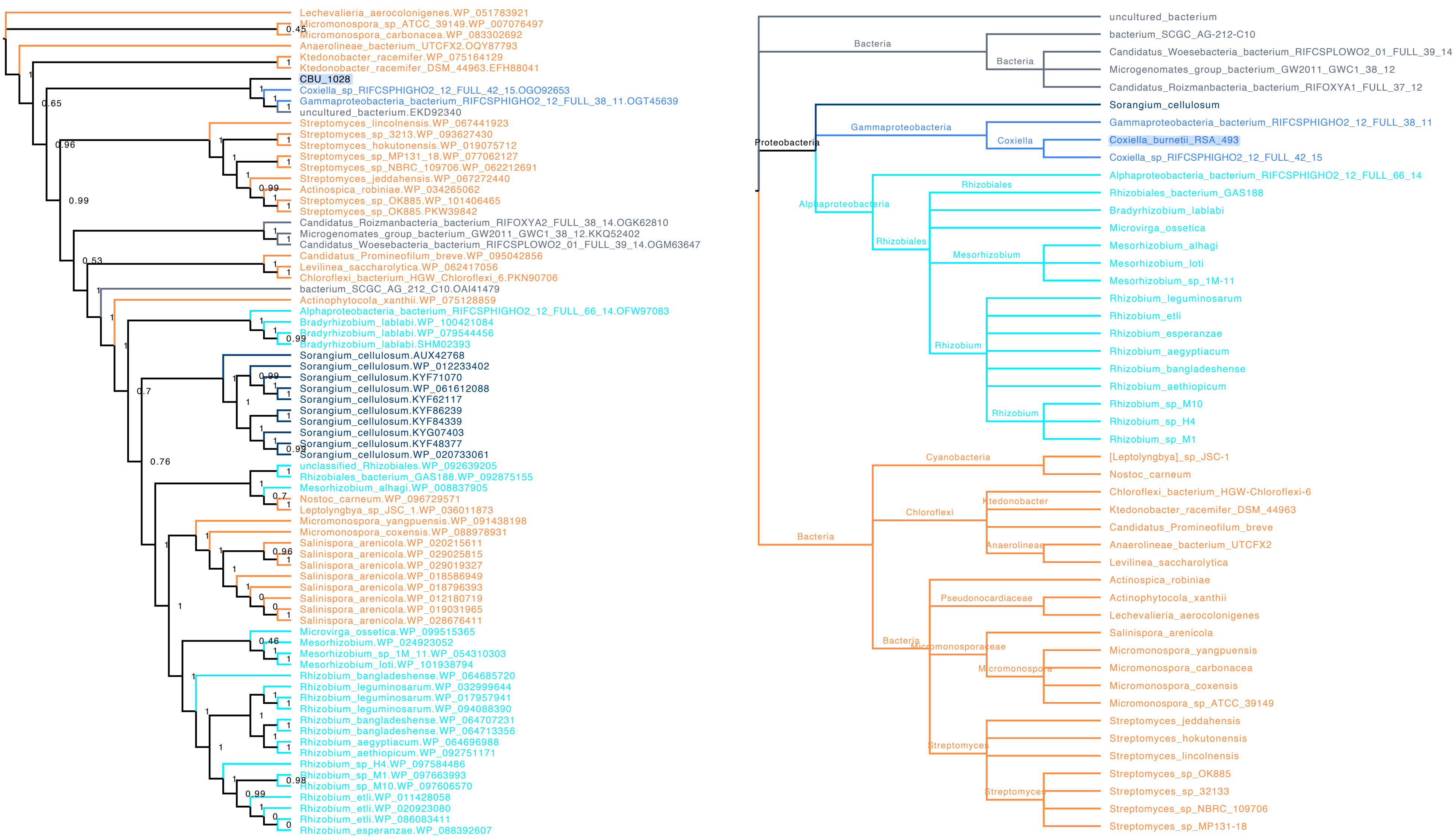

Lechevalieria_aerocolonigenes.WP_051783921
Micromonospora_sp_ATCC_39149.WP_007076497
Micromonospora_carbonacea.WP_083302692
Anaerolineae_bacterium_UTCFX2.OQY87793
Ktedonobacter_racemifer.WP_075164129
Ktedonobacter_racemifer_DSM_44963.EFH88041
CBU_1028
Coxiella_sp_RIFCSPHIGHO2_12_FULL_42_15.OGO92653
Gammaproteobacteria_bacterium_RIFCSPHIGHO2_12_FULL_38_11.OGT45639
uncultured_bacterium.EKD92340
Streptomyces_lincolnensis.WP_067441923
Streptomyces_sp_3213.WP_093627430
Streptomyces_hokutonensis.WP_019075712
Streptomyces_sp_MP131_18.WP_077062127
Streptomyces_sp_NBRC_109706.WP_062212691
Streptomyces_jeddahensis.WP_067272440
Actinospica_robiniae.WP_034265062
Streptomyces_sp_OK885.WP_101406465
Streptomyces_sp_OK885.PKW39842
Candidatus_Roizmanbacteria_bacterium_RIFOXYA2_FULL_38_14.OGK62810
Microgenomates_group_bacterium_GW2011_GWC1_38_12.KKQ52402
Candidatus_Woesebacteria_bacterium_RIFCSPLOWO2_01_FULL_39_14.OGM63647
Candidatus_Promineofilum_breve.WP_095042856
Levilinea_saccharolytica.WP_062417056
Chloroflexi_bacterium_HGW_Chloroflexi_6.PKN90706
bacterium_SCGC_AG_212_C10.OAI41479
Actinophytocola_xanthii.WP_075128859
Alphaproteobacteria_bacterium_RIFCSPHIGHO2_12_FULL_66_14.OFW97083
Bradyrhizobium_lablabi.WP_100421084
Bradyrhizobium_lablabi.WP_079544456
Bradyrhizobium_lablabi.SHM02393
Sorangium_cellulosum.AUX42768
Sorangium_cellulosum.WP_012233402
Sorangium_cellulosum.KYF71070
Sorangium_cellulosum.WP_061612088
Sorangium_cellulosum.KYF62117
Sorangium_cellulosum.KYF86239
Sorangium_cellulosum.KYF84339
Sorangium_cellulosum.KYG07403
Sorangium_cellulosum.KYF48377
Sorangium_cellulosum.WP_020733061
unclassified_Rhizobiales.WP_092639205
Rhizobiales_bacterium_GAS188.WP_092875155
Mesorhizobium_alhagi.WP_008837905
Nostoc_carneum.WP_096729571
Leptolyngbya_sp_JSC_1.WP_036011873
Micromonospora_yangpuensis.WP_091438198
Micromonospora_coxensis.WP_088978931
Salinispora_arenicola.WP_020215611
Salinispora_arenicola.WP_029025815
Salinispora_arenicola.WP_029019327
Salinispora_arenicola.WP_018586949
Salinispora_arenicola.WP_018796393
Salinispora_arenicola.WP_012180719
Salinispora_arenicola.WP_019031965
Salinispora_arenicola.WP_028676411
Microvirga_ossetica.WP_099515365
Mesorhizobium.WP_024923052
Mesorhizobium_sp_1M_11.WP_054310303
Mesorhizobium_loti.WP_101938794
Rhizobium_bangladeshense.WP_064685720
Rhizobium_leguminosarum.WP_032999644
Rhizobium_leguminosarum.WP_017957941
Rhizobium_leguminosarum.WP_094088390
Rhizobium_bangladeshense.WP_064707231
Rhizobium_bangladeshense.WP_064713356
Rhizobium_aegyptiacum.WP_064696988
Rhizobium_aethiopicum.WP_092751171
Rhizobium_sp_H4.WP_097584486
Rhizobium_sp_M1.WP_097663993
Rhizobium_sp_M10.WP_097606570
Rhizobium_etli.WP_011428058
Rhizobium_etli.WP_020923080
Rhizobium_etli.WP_086083411
Rhizobium_esperanzae.WP_088392607
uncultured_bacterium
Bacteria
bacterium_SCGC_AG-212-C10
Bacteria
Candidatus_Woesebacteria_bacterium_RIFCSPLOWO2_01_FULL_39_14
Microgenomates_group_bacterium_GW2011_GWC1_38_12
Candidatus_Roizmanbacteria_bacterium_RIFOXYA1_FULL_37_12
Sorangium_cellulosum
Gammaproteobacteria
Gammaproteobacteria_bacterium_RIFCSPHIGHO2_12_FULL_38_11
Proteobacteria
Coxiella
Coxiella_burnetii_RSA_493
Coxiella_sp_RIFCSPHIGHO2_12_FULL_42_15
Alphaproteobacteria_bacterium_RIFCSPHIGHO2_12_FULL_66_14
Rhizobiales
Rhizobiales_bacterium_GAS188
Alphaproteobacteria
Bradyrhizobium_lablabi
Microvirga_ossetica
Rhizobiales
Mesorhizobium_alhagi
Mesorhizobium
Mesorhizobium_loti
Mesorhizobium_sp_1M-11
Rhizobium_leguminosarum
Rhizobium_etli
Rhizobium_esperanzae
Rhizobium
Rhizobium_aegyptiacum
Rhizobium_bangladeshense
Rhizobium_aethiopicum
Rhizobium_sp_M10
Rhizobium
Rhizobium_sp_H4
Rhizobium_sp_M1
Cyanobacteria
[Leptolyngbya]_sp_JSC-1
Nostoc_carneum
Chloroflexi_bacterium_HGW-Chloroflexi-6
Ktedonobacter
Chloroflexi
Ktedonobacter_racemifer_DSM_44963
Candidatus_Promineofilum_breve
Bacteria
Anaerolineae
Anaerolineae_bacterium_UTCFX2
Levilinea_saccharolytica
Actinospica_robiniae
Pseudonocardiaceae
Actinophytocola_xanthii
Lechevalieria_aerocolonigenes
Salinispora_arenicola
Bacteria
Micromonosporaceae
Micromonospora_yangpuensis
Micromonospora
Micromonospora_carbonacea
Micromonospora_coxensis
Micromonospora_sp_ATCC_39149
Streptomyces_jeddahensis
Streptomyces_hokutonensis
Streptomyces
Streptomyces_lincolnensis
Streptomyces_sp_OK885
Streptomyces
Streptomyces_sp_32133
Streptomyces_sp_NBRC_109706
Streptomyces_sp_MP131-18

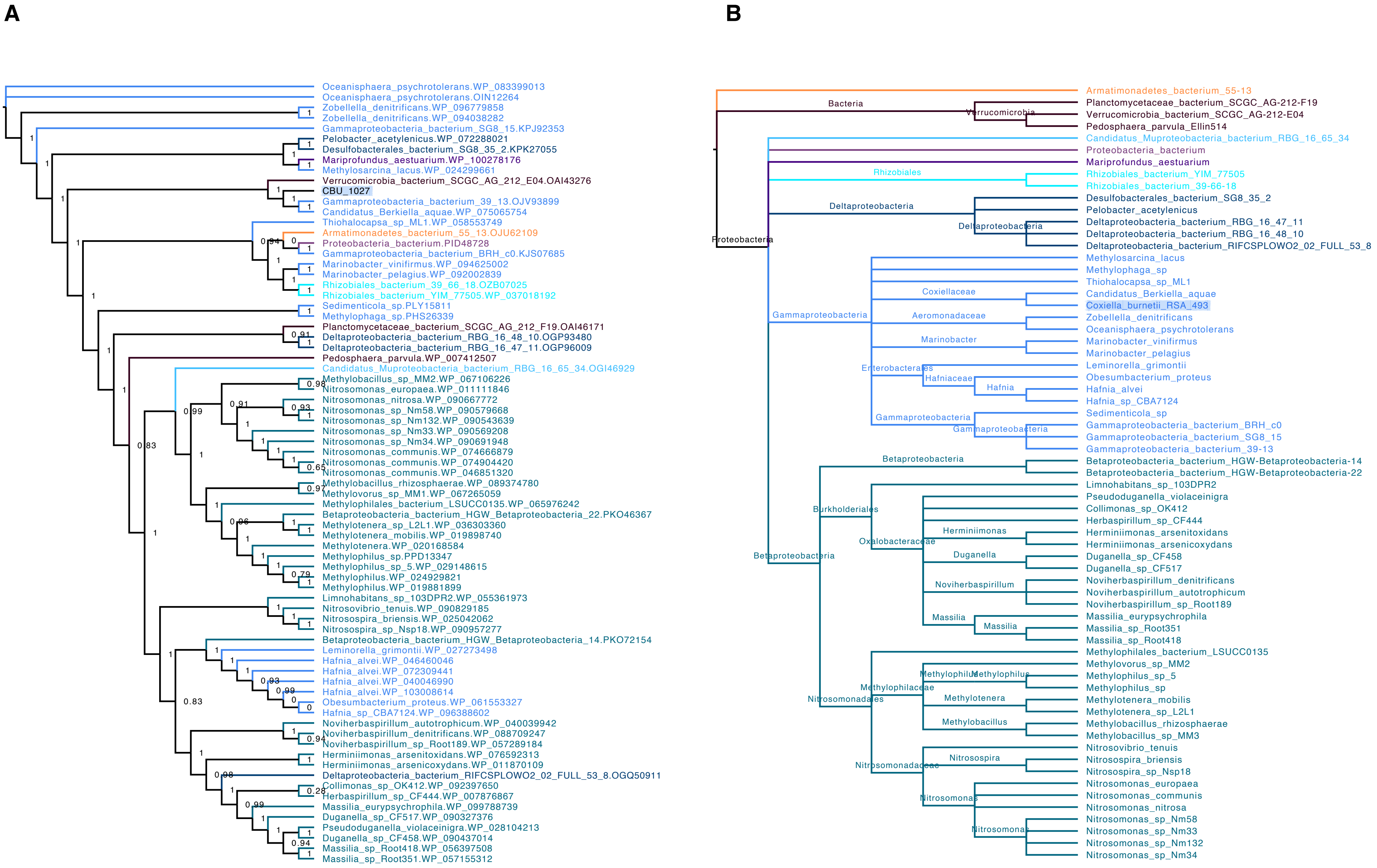
A
Oceanisphaera_psychrotolerans.WP_083399013
Oceanisphaera_psychrotolerans.OIN12264
Zobellella_denitrificans.WP_096779858
Zobellella_denitrificans.WP_094038282
Gammaproteobacteria_bacterium_SG8_15.KPJ92353
Pelobacter_acetylenicus.WP_072288021
Desulfobacterales_bacterium_SG8_35_2.KPK27055
Mariprofundus_aestuarium.WP_100278176
Methylosarcina_lacus.WP_024299661
Verrucomicrobia_bacterium_SCGC_AG_212_E04.OAI43276
CBU_1027
Gammaproteobacteria_bacterium_39_13.OJV93899
Candidatus_Berkiella_aquae.WP_075065754
Thiohalocapsa_sp_ML1.WP_058553749
Armatimonadetes_bacterium_55_13.OJU62109
Proteobacteria_bacterium.PID48728
Gammaproteobacteria_bacterium_BRH_c0.KJS07685
Marinobacter_vinifirmus.WP_094625002
Marinobacter_pelagius.WP_092002839
Rhizobiales_bacterium_39_66_18.OZB07025
Rhizobiales_bacterium_YIM_77505.WP_037018192
Sedimenticola_sp.PLY15811
Methylophaga_sp.PHS26339
Planctomycetaceae_bacterium_SCGC_AG_212_F19.OAI46171
Deltaproteobacteria_bacterium_RBG_16_48_10.OGP93480
Deltaproteobacteria_bacterium_RBG_16_47_11.OGP96009
Pedosphaera_parvula.WP_007412507
Candidatus_Muproteobacteria_bacterium_RBG_16_65_34.OGI46929
Methylobacillus_sp_MM2.WP_067106226
Nitrosomonas_europaea.WP_011111846
Nitrosomonas_nitrosa.WP_090667772
Nitrosomonas_sp_Nm58.WP_090579668
Nitrosomonas_sp_Nm132.WP_090543639
Nitrosomonas_sp_Nm33.WP_090569208
Nitrosomonas_sp_Nm34.WP_090691948
Nitrosomonas_communis.WP_074666879
Nitrosomonas_communis.WP_074904420
Nitrosomonas_communis.WP_046851320
Methylobacillus_rhizosphaerae.WP_089374780
Methylovorus_sp_MM1.WP_067265059
Methylophilales_bacterium_LSUCC0135.WP_065976242
Betaproteobacteria_bacterium_HGW_Betaproteobacteria_22.PKO46367
Methylotenera_sp_L2L1.WP_036303360
Methylotenera_mobilis.WP_019898740
Methylotenera.WP_020168584
Methylophilus_sp.PPD13347
Methylophilus_sp_5.WP_029148615
Methylophilus.WP_024929821
Methylophilus.WP_019881899
Limnohabitans_sp_103DPR2.WP_055361973
Nitrosovibrio_tenuis.WP_090829185
Nitrosospira_briensis.WP_025042062
Nitrosospira_sp_Nsp18.WP_090957277
Betaproteobacteria_bacterium_HGW_Betaproteobacteria_14.PKO72154
Leminorella_grimontii.WP_027273498
Hafnia_alvei.WP_046460046
Hafnia_alvei.WP_072309441
Hafnia_alvei.WP_040046990
Hafnia_alvei.WP_103008614
Obesumbacterium_proteus.WP_061553327
Hafnia_sp_CBA7124.WP_096388602
Noviherbaspirillum_autotrophicum.WP_040039942
Noviherbaspirillum_denitrificans.WP_088709247
Noviherbaspirillum_sp_Root189.WP_057289184
Herminiimonas_arsenitoxidans.WP_076592313
Herminiimonas_arsenicoxydans.WP_011870109
Deltaproteobacteria_bacterium_RIFCSPLOWO2_02_FULL_53_8.OGQ50911
Collimonas_sp_OK412.WP_092397650
Herbaspirillum_sp_CF444.WP_007876867
Massilia_eurypsychrophila.WP_099788739
Duganella_sp_CF517.WP_090327376
Pseudoduganella_violaceinigra.WP_028104213
Duganella_sp_CF458.WP_090437014
Massilia_sp_Root418.WP_056397508
Massilia_sp_Root351.WP_057155312
B
Bacteria
Verrucomicrobia
Proteobacteria
Rhizobiales
Deltaproteobacteria
Gammaproteobacteria
Coxiellaceae
Aeromonadaceae
Marinobacter
Enterobacterales
Hafniaceae
Hafnia
Betaproteobacteria
Burkholderiales
Oxalobacteraceae
Herminiimonas
Duganella
Noviherbaspirillum
Massilia
Nitrosomonadales
Methylophilaceae
Methylophilus
Methylotenera
Methylobacillus
Nitrosomonadaceae
Nitrosospira
Nitrosomonas
Armatimonadetes_bacterium_55-13
Planctomycetaceae_bacterium_SCGC_AG-212-F19
Verrucomicrobia_bacterium_SCGC_AG-212-E04
Pedosphaera_parvula_Ellin514
Candidatus_Muproteobacteria_bacterium_RBG_16_65_34
Proteobacteria_bacterium
Mariprofundus_aestuarium
Rhizobiales_bacterium_YIM_77505
Rhizobiales_bacterium_39-66-18
Desulfobacterales_bacterium_SG8_35_2
Pelobacter_acetylenicus
Deltaproteobacteria_bacterium_RBG_16_47_11
Deltaproteobacteria_bacterium_RBG_16_48_10
Deltaproteobacteria_bacterium_RIFCSPLOWO2_02_FULL_53_8
Methylosarcina_lacus
Methylophaga_sp
Thiohalocapsa_sp_ML1
Candidatus_Berkiella_aquae
Coxiella_burnetii_RSA_493
Zobellella_denitrificans
Oceanisphaera_psychrotolerans
Marinobacter_vinifirmus
Marinobacter_pelagius
Leminorella_grimontii
Obesumbacterium_proteus
Hafnia_alvei
Hafnia_sp_CBA7124
Sedimenticola_sp
Gammaproteobacteria_bacterium_BRH_c0
Gammaproteobacteria_bacterium_SG8_15
Gammaproteobacteria_bacterium_39-13
Betaproteobacteria_bacterium_HGW-Betaproteobacteria-14
Betaproteobacteria_bacterium_HGW-Betaproteobacteria-22
Limnohabitans_sp_103DPR2
Pseudoduganella_violaceinigra
Collimonas_sp_OK412
Herbaspirillum_sp_CF444
Herminiimonas_arsenitoxidans
Herminiimonas_arsenicoxydans
Duganella_sp_CF458
Duganella_sp_CF517
Noviherbaspirillum_denitrificans
Noviherbaspirillum_autotrophicum
Noviherbaspirillum_sp_Root189
Massilia_eurypsychrophila
Massilia_sp_Root351
Massilia_sp_Root418
Methylophilales_bacterium_LSUCC0135
Methylovorus_sp_MM2
Methylophilus_sp_5
Methylophilus_sp
Methylotenera_mobilis
Methylotenera_sp_L2L1
Methylobacillus_rhizosphaerae
Methylobacillus_sp_MM3
Nitrosovibrio_tenuis
Nitrosospira_briensis
Nitrosospira_sp_Nsp18
Nitrosomonas_europaea
Nitrosomonas_communis
Nitrosomonas_nitrosa
Nitrosomonas_sp_Nm58
Nitrosomonas_sp_Nm33
Nitrosomonas_sp_Nm132
Nitrosomonas_sp_Nm34

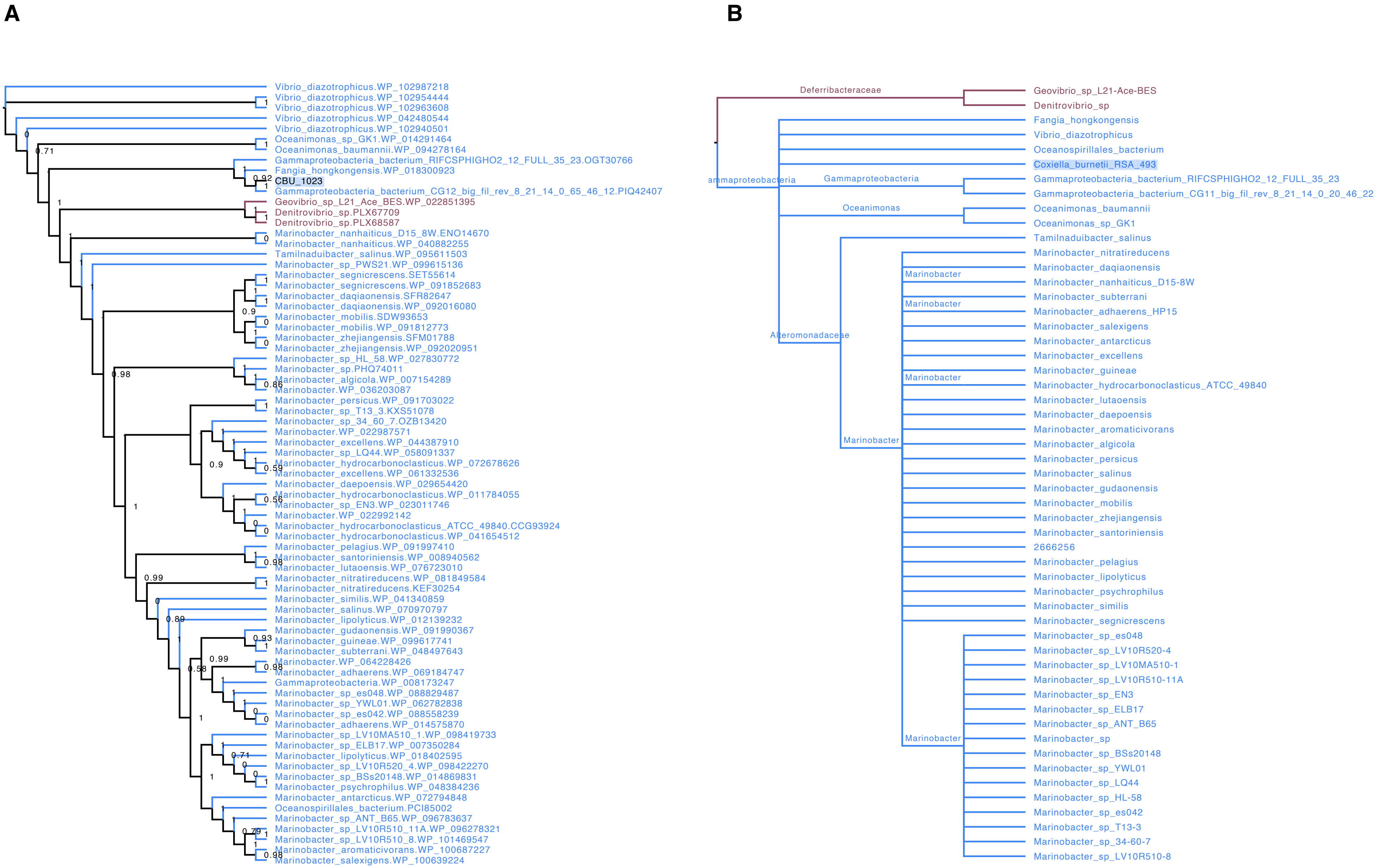
A
Vibrio_diazotrophicus.WP_102987218
Vibrio_diazotrophicus.WP_102954444
Vibrio_diazotrophicus.WP_102963608
Vibrio_diazotrophicus.WP_042480544
Vibrio_diazotrophicus.WP_102940501
Oceanimonas_sp_GK1.WP_014291464
Oceanimonas_baumannii.WP_094278164
Gammaproteobacteria_bacterium_RIFCSPHIGHO2_12_FULL_35_23.OGT30766
Fangia_hongkongensis.WP_018300923
CBU_1023
Gammaproteobacteria_bacterium_CG12_big_fil_rev_8_21_14_0_65_46_12.PIQ42407
Geovibrio_sp_L21_Ace_BES.WP_022851395
Denitrovibrio_sp.PLX67709
Denitrovibrio_sp.PLX68587
Marinobacter_nanhaiticus_D15_8W.ENO14670
Marinobacter_nanhaiticus.WP_040882255
Tamilnaduibacter_salinus.WP_095611503
Marinobacter_sp_PWS21.WP_099615136
Marinobacter_segnicrescens.SET55614
Marinobacter_segnicrescens.WP_091852683
Marinobacter_daqiaonensis.SFR82647
Marinobacter_daqiaonensis.WP_092016080
Marinobacter_mobilis.SDW93653
Marinobacter_mobilis.WP_091812773
Marinobacter_zhejiangensis.SFM01788
Marinobacter_zhejiangensis.WP_092020951
Marinobacter_sp_HL_58.WP_027830772
Marinobacter_sp.PHQ74011
Marinobacter_algicola.WP_007154289
Marinobacter.WP_036203087
Marinobacter_persicus.WP_091703022
Marinobacter_sp_T13_3.KXS51078
Marinobacter_sp_34_60_7.OZB13420
Marinobacter.WP_022987571
Marinobacter_excellens.WP_044387910
Marinobacter_sp_LQ44.WP_058091337
Marinobacter_hydrocarbonoclasticus.WP_072678626
Marinobacter_excellens.WP_061332536
Marinobacter_daepoensis.WP_029654420
Marinobacter_hydrocarbonoclasticus.WP_011784055
Marinobacter_sp_EN3.WP_023011746
Marinobacter.WP_022992142
Marinobacter_hydrocarbonoclasticus_ATCC_49840.CCG93924
Marinobacter_hydrocarbonoclasticus.WP_041654512
Marinobacter_pelagius.WP_091997410
Marinobacter_santoriniensis.WP_008940562
Marinobacter_lutaoensis.WP_076723010
Marinobacter_nitratireducens.WP_081849584
Marinobacter_nitratireducens.KEF30254
Marinobacter_similis.WP_041340859
Marinobacter_salinus.WP_070970797
Marinobacter_lipolyticus.WP_012139232
Marinobacter_gudaonensis.WP_091990367
Marinobacter_guineae.WP_099617741
Marinobacter_subterrani.WP_048497643
Marinobacter.WP_064228426
Marinobacter_adhaerens.WP_069184747
Gammaproteobacteria.WP_008173247
Marinobacter_sp_es048.WP_088829487
Marinobacter_sp_YWL01.WP_062782838
Marinobacter_sp_es042.WP_088558239
Marinobacter_adhaerens.WP_014575870
Marinobacter_sp_LV10MA510_1.WP_098419733
Marinobacter_sp_ELB17.WP_007350284
Marinobacter_lipolyticus.WP_018402595
Marinobacter_sp_LV10R520_4.WP_098422270
Marinobacter_sp_BSs20148.WP_014869831
Marinobacter_psychrophilus.WP_048384236
Marinobacter_antarcticus.WP_072794848
Oceanospirillales_bacterium.PCI85002
Marinobacter_sp_ANT_B65.WP_096783637
Marinobacter_sp_LV10R510_11A.WP_096278321
Marinobacter_sp_LV10R510_8.WP_101469547
Marinobacter_aromaticivorans.WP_100687227
Marinobacter_salexigens.WP_100639224
B
Deferribacteraceae
Geovibrio_sp_L21-Ace-BES
Denitrovibrio_sp
Fangia_hongkongensis
Vibrio_diazotrophicus
Oceanospirillales_bacterium
Coxiella_burnetii_RSA_493
Gammaproteobacteria
Gammaproteobacteria
Gammaproteobacteria_bacterium_RIFCSPHIGHO2_12_FULL_35_23
Gammaproteobacteria_bacterium_CG11_big_fil_rev_8_21_14_0_20_46_22
Oceanimonas
Oceanimonas_baumannii
Oceanimonas_sp_GK1
Tamilnaduibacter_salinus
Alteromonadaceae
Marinobacter
Marinobacter_nitratireducens
Marinobacter_daqiaonensis
Marinobacter_nanhaiticus_D15-8W
Marinobacter_subterrani
Marinobacter_adhaerens_HP15
Marinobacter_salexigens
Marinobacter_antarcticus
Marinobacter_excellens
Marinobacter_guineae
Marinobacter_hydrocarbonoclasticus_ATCC_49840
Marinobacter_lutaoensis
Marinobacter_daepoensis
Marinobacter_aromaticivorans
Marinobacter_algicola
Marinobacter_persicus
Marinobacter_salinus
Marinobacter_gudaonensis
Marinobacter_mobilis
Marinobacter_zhejiangensis
Marinobacter_santoriniensis
2666256
Marinobacter_pelagius
Marinobacter_lipolyticus
Marinobacter_psychrophilus
Marinobacter_similis
Marinobacter_segnicrescens
Marinobacter_sp_es048
Marinobacter_sp_LV10R520-4
Marinobacter_sp_LV10MA510-1
Marinobacter_sp_LV10R510-11A
Marinobacter_sp_EN3
Marinobacter_sp_ELB17
Marinobacter_sp_ANT_B65
Marinobacter_sp
Marinobacter_sp_BSs20148
Marinobacter_sp_YWL01
Marinobacter_sp_LQ44
Marinobacter_sp_HL-58
Marinobacter_sp_es042
Marinobacter_sp_T13-3
Marinobacter_sp_34-60-7
Marinobacter_sp_LV10R510-8

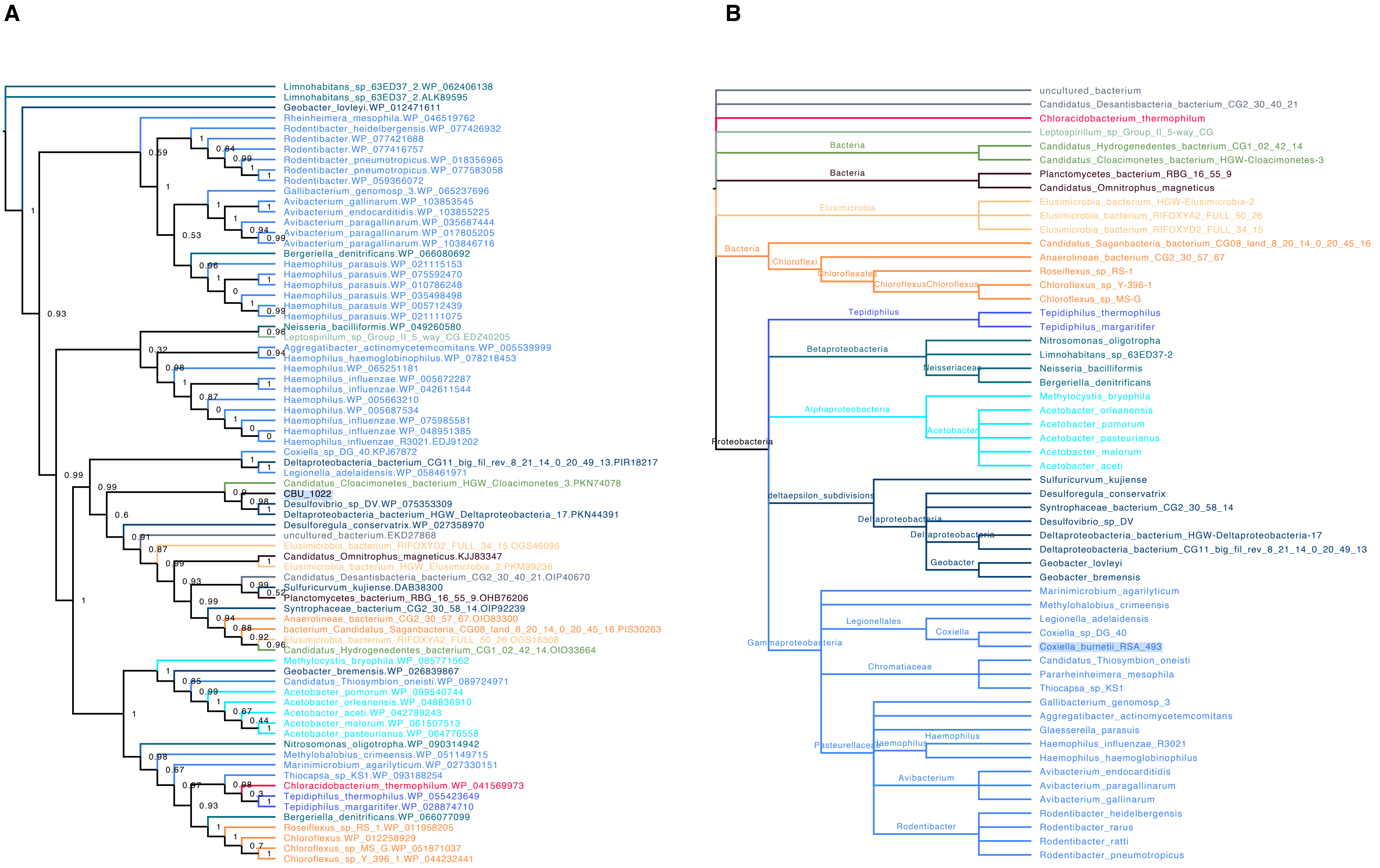
A
Limnohabitans_sp_63ED37_2.WP_062406138
Limnohabitans_sp_63ED37_2.ALK89595
Geobacter_lovleyi.WP_012471611
Rheinheimera_mesophila.WP_046519762
Rodentibacter_heidelbergensis.WP_077426932
Rodentibacter.WP_077421688
Rodentibacter.WP_077416757
Rodentibacter_pneumotropicus.WP_018356965
Rodentibacter_pneumotropicus.WP_077583058
Rodentibacter.WP_059366072
Gallibacterium_genomosp_3.WP_065237696
Avibacterium_gallinarum.WP_103853545
Avibacterium_endocarditidis.WP_103855225
Avibacterium_paragallinarum.WP_035687444
Avibacterium_paragallinarum.WP_017805205
Avibacterium_paragallinarum.WP_103846716
Bergeriella_denitrificans.WP_066080692
Haemophilus_parasuis.WP_021115153
Haemophilus_parasuis.WP_075592470
Haemophilus_parasuis.WP_010786248
Haemophilus_parasuis.WP_035498498
Haemophilus_parasuis.WP_005712439
Haemophilus_parasuis.WP_021111075
Neisseria_bacilliformis.WP_049260580
Leptospirillum_sp_Group_II_5_way_CG.EDZ40205
Aggregatibacter_actinomycetemcomitans.WP_005539999
Haemophilus_haemoglobinophilus.WP_078218453
Haemophilus.WP_065251181
Haemophilus_influenzae.WP_005672287
Haemophilus_influenzae.WP_042611544
Haemophilus.WP_005663210
Haemophilus.WP_005687534
Haemophilus_influenzae.WP_075985581
Haemophilus_influenzae.WP_048951385
Haemophilus_influenzae_R3021.EDJ91202
Coxiella_sp_DG_40.KPJ67872
Deltaproteobacteria_bacterium_CG11_big_fil_rev_8_21_14_0_20_49_13.PIR18217
Legionella_adelaidensis.WP_058461971
Candidatus_Cloacimonetes_bacterium_HGW_Cloacimonetes_3.PKN74078
CBU_1022
Desulfovibrio_sp_DV.WP_075353309
Deltaproteobacteria_bacterium_HGW_Deltaproteobacteria_17.PKN44391
Desulforegula_conservatrix.WP_027358970
uncultured_bacterium.EKD27868
Elusimicrobia_bacterium_RIFOXYD2_FULL_34_15.OGS46096
Candidatus_Omnitrophus_magneticus.KJJ83347
Elusimicrobia_bacterium_HGW_Elusimicrobia_2.PKM99236
Candidatus_Desantisbacteria_bacterium_CG2_30_40_21.OIP40670
Sulfuricurvum_kujiense.DAB38300
Planctomycetes_bacterium_RBG_16_55_9.OHB76206
Syntrophaceae_bacterium_CG2_30_58_14.OIP92239
Anaerolineae_bacterium_CG2_30_57_67.OIO83300
bacterium_Candidatus_Saganbacteria_CG08_land_8_20_14_0_20_45_16.PIS30263
Elusimicrobia_bacterium_RIFOXYA2_FULL_50_26.OGS18308
Candidatus_Hydrogenedentes_bacterium_CG1_02_42_14.OIO33664
Methylocystis_bryophila.WP_085771562
Geobacter_bremensis.WP_026839867
Candidatus_Thiosymbion_oneisti.WP_089724971
Acetobacter_pomorum.WP_099540744
Acetobacter_orleanensis.WP_048836910
Acetobacter_aceti.WP_042789243
Acetobacter_malorum.WP_061507513
Acetobacter_pasteurianus.WP_064776558
Nitrosomonas_oligotropha.WP_090314942
Methylohalobius_crimeensis.WP_051149715
Marinimicrobium_agarilyticum.WP_027330151
Thiocapsa_sp_KS1.WP_093188254
Chloracidobacterium_thermophilum.WP_041569973
Tepidiphilus_thermophilus.WP_055423649
Tepidiphilus_margaritifer.WP_028874710
Bergeriella_denitrificans.WP_066077099
Roseiflexus_sp_RS_1.WP_011958205
Chloroflexus.WP_012258929
Chloroflexus_sp_MS_G.WP_051871037
Chloroflexus_sp_Y_396_1.WP_044232441
B
uncultured_bacterium
Candidatus_Desantisbacteria_bacterium_CG2_30_40_21
Chloracidobacterium_thermophilum
Leptospirillum_sp_Group_II_5-way_CG
Bacteria
Candidatus_Hydrogenedentes_bacterium_CG1_02_42_14
Candidatus_Cloacimonetes_bacterium_HGW-Cloacimonetes-3
Bacteria
Planctomycetes_bacterium_RBG_16_55_9
Candidatus_Omnitrophus_magneticus
Elusimicrobia
Elusimicrobia_bacterium_HGW-Elusimicrobia-2
Elusimicrobia_bacterium_RIFOXYA2_FULL_50_26
Elusimicrobia_bacterium_RIFOXYD2_FULL_34_15
Bacteria
Candidatus_Saganbacteria_bacterium_CG08_land_8_20_14_0_20_45_16
Chloroflexi
Anaerolineae_bacterium_CG2_30_57_67
Chloroflexales
Roseiflexus_sp_RS-1
ChloroflexusChloroflexus
Chloroflexus_sp_Y-396-1
Chloroflexus_sp_MS-G
Tepidiphilus
Tepidiphilus_thermophilus
Tepidiphilus_margaritifer
Betaproteobacteria
Nitrosomonas_oligotropha
Limnohabitans_sp_63ED37-2
Neisseriaceae
Neisseria_bacilliformis
Bergeriella_denitrificans
Alphaproteobacteria
Methylocystis_bryophila
Acetobacter
Acetobacter_orleanensis
Acetobacter_pomorum
Acetobacter_pasteurianus
Acetobacter_malorum
Acetobacter_aceti
Proteobacteria
deltaepsilon_subdivisions
Sulfuricurvum_kujiense
Deltaproteobacteria
Desulforegula_conservatrix
Syntrophaceae_bacterium_CG2_30_58_14
Desulfovibrio_sp_DV
Deltaproteobacteria
Deltaproteobacteria_bacterium_HGW-Deltaproteobacteria-17
Deltaproteobacteria_bacterium_CG11_big_fil_rev_8_21_14_0_20_49_13
Geobacter
Geobacter_lovleyi
Geobacter_bremensis
Gammaproteobacteria
Marinimicrobium_agarilyticum
Methylohalobius_crimeensis
Legionellales
Legionella_adelaidensis
Coxiella
Coxiella_sp_DG_40
Coxiella_burnetii_RSA_493
Chromatiaceae
Candidatus_Thiosymbion_oneisti
Pararheinheimera_mesophila
Thiocapsa_sp_KS1
Pasteurellaceae
Gallibacterium_genomosp_3
Aggregatibacter_actinomycetemcomitans
Glaesserella_parasuis
Haemophilus
Haemophilus
Haemophilus_influenzae_R3021
Haemophilus_haemoglobinophilus
Avibacterium
Avibacterium_endocarditidis
Avibacterium_paragallinarum
Avibacterium_gallinarum
Rodentibacter
Rodentibacter_heidelbergensis
Rodentibacter_rarus
Rodentibacter_ratti
Rodentibacter_pneumotropicus

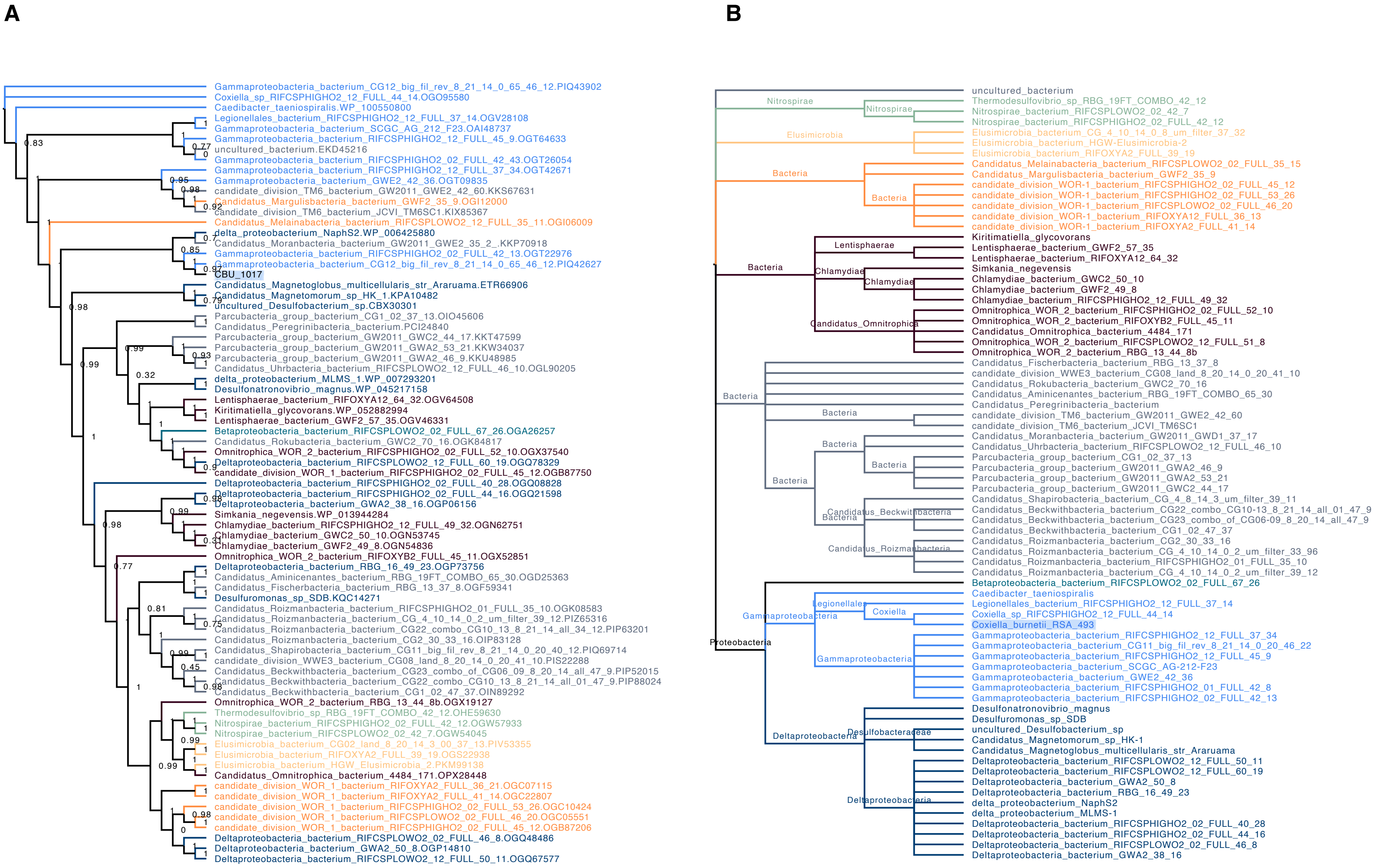
A
Gammaproteobacteria_bacterium_CG12_big_fil_rev_8_21_14_0_65_46_12.PIQ43902
Coxiella_sp_RIFCSPHIGHO2_12_FULL_44_14.OGO95580
Caedibacter_taeniospiralis.WP_100550800
Legionellales_bacterium_RIFCSPHIGHO2_12_FULL_37_14.OGV28108
Gammaproteobacteria_bacterium_SCGC_AG_212_F23.OAI48737
Gammaproteobacteria_bacterium_RIFCSPHIGHO2_12_FULL_45_9.OGT64633
uncultured_bacterium.EKD45216
Gammaproteobacteria_bacterium_RIFCSPHIGHO2_02_FULL_42_43.OGT26054
Gammaproteobacteria_bacterium_RIFCSPHIGHO2_12_FULL_37_34.OGT42671
Gammaproteobacteria_bacterium_GWE2_42_36.OGT09835
candidate_division_TM6_bacterium_GW2011_GWE2_42_60.KKS67631
Candidatus_Margulisbacteria_bacterium_GWF2_35_9.OGI12000
candidate_division_TM6_bacterium_JCVI_TM6SC1.KIX85367
Candidatus_Melainabacteria_bacterium_RIFCSPLOWO2_12_FULL_35_11.OGI06009
delta_proteobacterium_NaphS2.WP_006425880
Candidatus_Moranbacteria_bacterium_GW2011_GWE2_35_2_.KKP70918
Gammaproteobacteria_bacterium_RIFCSPHIGHO2_02_FULL_42_13.OGT22976
Gammaproteobacteria_bacterium_CG12_big_fil_rev_8_21_14_0_65_46_12.PIQ42627
CBU_1017
Candidatus_Magnetoglobus_multicellularis_str_Araruama.ETR66906
Candidatus_Magnetomorum_sp_HK_1.KPA10482
uncultured_Desulfobacterium_sp.CBX30301
Parcubacteria_group_bacterium_CG1_02_37_13.OIO45606
Candidatus_Peregrinibacteria_bacterium.PCI24840
Parcubacteria_group_bacterium_GW2011_GWC2_44_17.KKT47599
Parcubacteria_group_bacterium_GW2011_GWA2_53_21.KKW34037
Parcubacteria_group_bacterium_GW2011_GWA2_46_9.KKU48985
Candidatus_Uhrbacteria_bacterium_RIFCSPLOWO2_12_FULL_46_10.OGL90205
delta_proteobacterium_MLMS_1.WP_007293201
Desulfonatronovibrio_magnus.WP_045217158
Lentisphaerae_bacterium_RIFOXYA12_64_32.OGV64508
Kiritimatiella_glycovorans.WP_052882994
Lentisphaerae_bacterium_GWF2_57_35.OGV46331
Betaproteobacteria_bacterium_RIFCSPLOWO2_02_FULL_67_26.OGA26257
Candidatus_Rokubacteria_bacterium_GWC2_70_16.OGK84817
Omnitrophica_WOR_2_bacterium_RIFCSPHIGHO2_02_FULL_52_10.OGX37540
Deltaproteobacteria_bacterium_RIFCSPLOWO2_12_FULL_60_19.OGQ78329
candidate_division_WOR_1_bacterium_RIFCSPHIGHO2_02_FULL_45_12.OGB87750
Deltaproteobacteria_bacterium_RIFCSPHIGHO2_02_FULL_40_28.OGQ08828
Deltaproteobacteria_bacterium_RIFCSPHIGHO2_02_FULL_44_16.OGQ21598
Deltaproteobacteria_bacterium_GWA2_38_16.OGP06156
Simkania_negevensis.WP_013944284
Chlamydiae_bacterium_RIFCSPHIGHO2_12_FULL_49_32.OGN62751
Chlamydiae_bacterium_GWC2_50_10.OGN53745
Chlamydiae_bacterium_GWF2_49_8.OGN54836
Omnitrophica_WOR_2_bacterium_RIFOXYB2_FULL_45_11.OGX52851
Deltaproteobacteria_bacterium_RBG_16_49_23.OGP73756
Candidatus_Aminicenantes_bacterium_RBG_19FT_COMBO_65_30.OGD25363
Candidatus_Fischerbacteria_bacterium_RBG_13_37_8.OGF59341
Desulfuromonas_sp_SDB.KQC14271
Candidatus_Roizmanbacteria_bacterium_RIFCSPHIGHO2_01_FULL_35_10.OGK08583
Candidatus_Roizmanbacteria_bacterium_CG_4_10_14_0_2_um_filter_39_12.PIZ65316
Candidatus_Roizmanbacteria_bacterium_CG22_combo_CG10_13_8_21_14_all_34_12.PIP63201
Candidatus_Roizmanbacteria_bacterium_CG2_30_33_16.OIP83128
Candidatus_Shapirobacteria_bacterium_CG11_big_fil_rev_8_21_14_0_20_40_12.PIQ69714
candidate_division_WWE3_bacterium_CG08_land_8_20_14_0_20_41_10.PIS22288
Candidatus_Beckwithbacteria_bacterium_CG23_combo_of_CG06_09_8_20_14_all_47_9.PIP52015
Candidatus_Beckwithbacteria_bacterium_CG22_combo_CG10_13_8_21_14_all_01_47_9.PIP88024
Candidatus_Beckwithbacteria_bacterium_CG1_02_47_37.OIN89292
Omnitrophica_WOR_2_bacterium_RBG_13_44_8b.OGX19127
Thermodesulfovibrio_sp_RBG_19FT_COMBO_42_12.OHE59630
Nitrospirae_bacterium_RIFCSPHIGHO2_02_FULL_42_12.OGW57933
Nitrospirae_bacterium_RIFCSPLOWO2_02_42_7.OGW54045
Elusimicrobia_bacterium_CG02_land_8_20_14_3_00_37_13.PIV53355
Elusimicrobia_bacterium_RIFOXYA2_FULL_39_19.OGS22938
Elusimicrobia_bacterium_HGW_Elusimicrobia_2.PKM99138
Candidatus_Omnitrophica_bacterium_4484_171.OPX28448
candidate_division_WOR_1_bacterium_RIFOXYA2_FULL_36_21.OGC07115
candidate_division_WOR_1_bacterium_RIFOXYA2_FULL_41_14.OGC22807
candidate_division_WOR_1_bacterium_RIFCSPHIGHO2_02_FULL_53_26.OGC10424
candidate_division_WOR_1_bacterium_RIFCSPLOWO2_02_FULL_46_20.OGC05551
candidate_division_WOR_1_bacterium_RIFCSPHIGHO2_02_FULL_45_12.OGB87206
Deltaproteobacteria_bacterium_RIFCSPLOWO2_02_FULL_46_8.OGQ48486
Deltaproteobacteria_bacterium_GWA2_50_8.OGP14810
Deltaproteobacteria_bacterium_RIFCSPLOWO2_12_FULL_50_11.OGQ67577
B
uncultured_bacterium
Nitrospirae
Thermodesulfovibrio_sp_RBG_19FT_COMBO_42_12
Nitrospirae_bacterium_RIFCSPLOWO2_02_42_7
Nitrospirae_bacterium_RIFCSPHIGHO2_02_FULL_42_12
Elusimicrobia
Elusimicrobia_bacterium_CG_4_10_14_0_8_um_filter_37_32
Elusimicrobia_bacterium_HGW-Elusimicrobia-2
Elusimicrobia_bacterium_RIFOXYA2_FULL_39_19
Bacteria
Candidatus_Melainabacteria_bacterium_RIFCSPLOWO2_02_FULL_35_15
Candidatus_Margulisbacteria_bacterium_GWF2_35_9
candidate_division_WOR-1_bacterium_RIFCSPHIGHO2_02_FULL_45_12
candidate_division_WOR-1_bacterium_RIFCSPHIGHO2_02_FULL_53_26
candidate_division_WOR-1_bacterium_RIFCSPLOWO2_02_FULL_46_20
candidate_division_WOR-1_bacterium_RIFOXYA12_FULL_36_13
candidate_division_WOR-1_bacterium_RIFOXYA2_FULL_41_14
Lentisphaerae
Chlamydiae
Candidatus_Omnitrophica
Kiritimatiella_glycovorans
Lentisphaerae_bacterium_GWF2_57_35
Lentisphaerae_bacterium_RIFOXYA12_64_32
Simkania_negevensis
Chlamydiae_bacterium_GWC2_50_10
Chlamydiae_bacterium_GWF2_49_8
Chlamydiae_bacterium_RIFCSPHIGHO2_12_FULL_49_32
Omnitrophica_WOR_2_bacterium_RIFCSPHIGHO2_02_FULL_52_10
Omnitrophica_WOR_2_bacterium_RIFOXYB2_FULL_45_11
Candidatus_Omnitrophica_bacterium_4484_171
Omnitrophica_WOR_2_bacterium_RIFCSPLOWO2_12_FULL_51_8
Omnitrophica_WOR_2_bacterium_RBG_13_44_8b
Candidatus_Fischerbacteria_bacterium_RBG_13_37_8
candidate_division_WWE3_bacterium_CG08_land_8_20_14_0_20_41_10
Candidatus_Rokubacteria_bacterium_GWC2_70_16
Candidatus_Aminicenantes_bacterium_RBG_19FT_COMBO_65_30
Candidatus_Peregrinibacteria_bacterium
candidate_division_TM6_bacterium_GW2011_GWE2_42_60
candidate_division_TM6_bacterium_JCVI_TM6SC1
Candidatus_Moranbacteria_bacterium_GW2011_GWD1_37_17
Candidatus_Uhrbacteria_bacterium_RIFCSPLOWO2_12_FULL_46_10
Parcubacteria_group_bacterium_CG1_02_37_13
Parcubacteria_group_bacterium_GW2011_GWA2_46_9
Parcubacteria_group_bacterium_GW2011_GWA2_53_21
Parcubacteria_group_bacterium_GW2011_GWC2_44_17
Candidatus_Beckwithbacteria
Candidatus_Roizmanbacteria
Candidatus_Shapirobacteria_bacterium_CG_4_8_14_3_um_filter_39_11
Candidatus_Beckwithbacteria_bacterium_CG22_combo_CG10-13_8_21_14_all_01_47_9
Candidatus_Beckwithbacteria_bacterium_CG23_combo_of_CG06-09_8_20_14_all_47_9
Candidatus_Beckwithbacteria_bacterium_CG1_02_47_37
Candidatus_Roizmanbacteria_bacterium_CG2_30_33_16
Candidatus_Roizmanbacteria_bacterium_CG_4_10_14_0_2_um_filter_33_96
Candidatus_Roizmanbacteria_bacterium_RIFCSPHIGHO2_01_FULL_35_10
Candidatus_Roizmanbacteria_bacterium_CG_4_10_14_0_2_um_filter_39_12
Proteobacteria
Gammaproteobacteria
Legionellales
Coxiella
Betaproteobacteria_bacterium_RIFCSPLOWO2_02_FULL_67_26
Caedibacter_taeniospiralis
Legionellales_bacterium_RIFCSPHIGHO2_12_FULL_37_14
Coxiella_sp_RIFCSPHIGHO2_12_FULL_44_14
Coxiella_burnetii_RSA_493
Gammaproteobacteria_bacterium_RIFCSPHIGHO2_12_FULL_37_34
Gammaproteobacteria_bacterium_CG11_big_fil_rev_8_21_14_0_20_46_22
Gammaproteobacteria_bacterium_RIFCSPHIGHO2_12_FULL_45_9
Gammaproteobacteria_bacterium_SCGC_AG-212-F23
Gammaproteobacteria_bacterium_GWE2_42_36
Gammaproteobacteria_bacterium_RIFCSPHIGHO2_01_FULL_42_8
Gammaproteobacteria_bacterium_RIFCSPHIGHO2_02_FULL_42_13
Deltaproteobacteria
Desulfobacteraceae
Desulfonatronovibrio_magnus
Desulfuromonas_sp_SDB
uncultured_Desulfobacterium_sp
Candidatus_Magnetomorum_sp_HK-1
Candidatus_Magnetoglobus_multicellularis_str_Araruama
Deltaproteobacteria_bacterium_RIFCSPLOWO2_12_FULL_50_11
Deltaproteobacteria_bacterium_RIFCSPLOWO2_12_FULL_60_19
Deltaproteobacteria_bacterium_GWA2_50_8
Deltaproteobacteria_bacterium_RBG_16_49_23
delta_proteobacterium_NaphS2
delta_proteobacterium_MLMS-1
Deltaproteobacteria_bacterium_RIFCSPHIGHO2_02_FULL_40_28
Deltaproteobacteria_bacterium_RIFCSPHIGHO2_02_FULL_44_16
Deltaproteobacteria_bacterium_RIFCSPLOWO2_02_FULL_46_8
Deltaproteobacteria_bacterium_GWA2_38_16

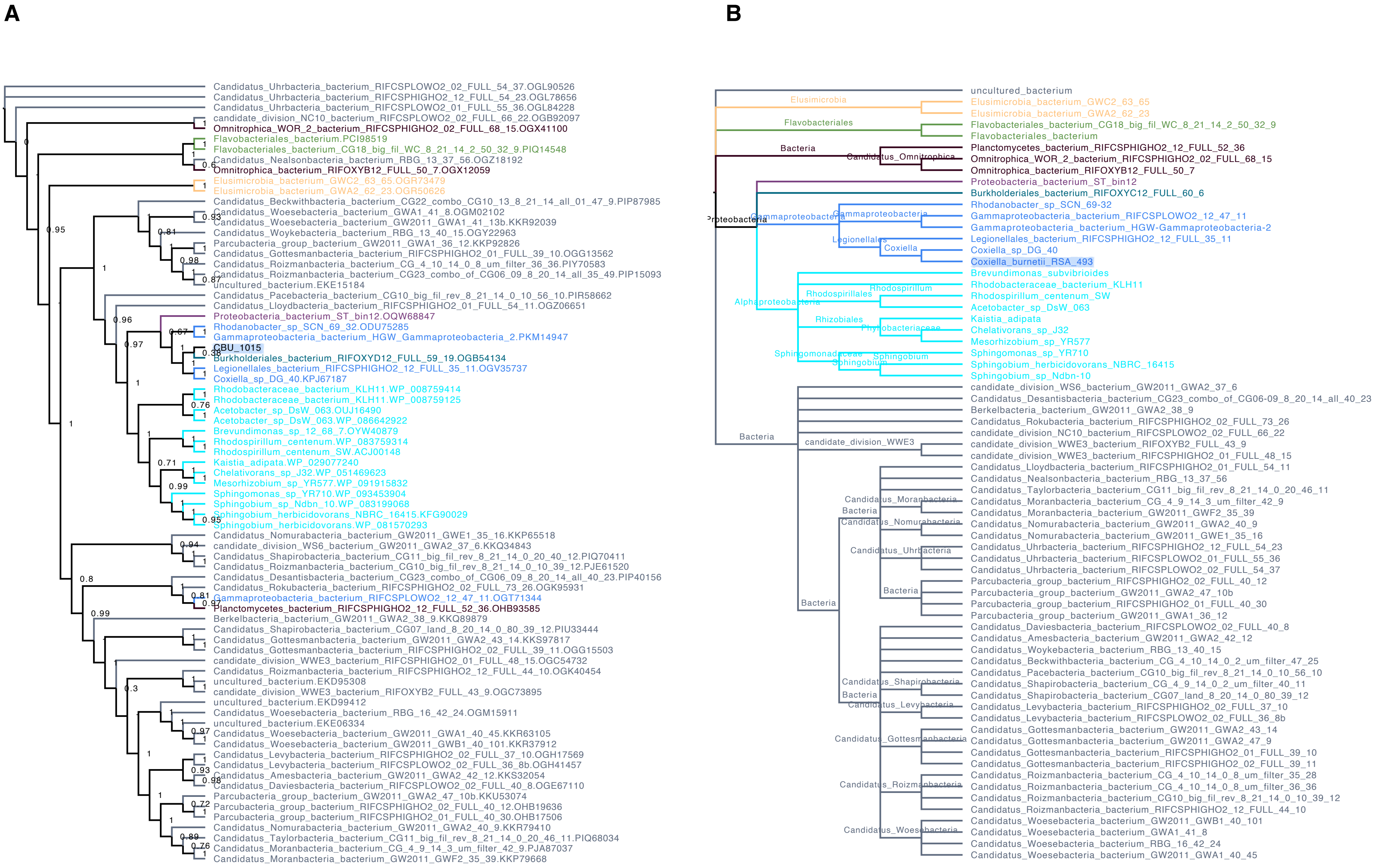
A
Candidatus_Uhrbacteria_bacterium_RIFCSPLOWO2_02_FULL_54_37.OGL90526
Candidatus_Uhrbacteria_bacterium_RIFCSPHIGHO2_12_FULL_54_23.OGL78656
Candidatus_Uhrbacteria_bacterium_RIFCSPLOWO2_01_FULL_55_36.OGL84228
candidate_division_NC10_bacterium_RIFCSPLOWO2_02_FULL_66_22.OGB92097
Omnitrophica_WOR_2_bacterium_RIFCSPHIGHO2_02_FULL_68_15.OGX41100
Flavobacteriales_bacterium.PCI98519
Flavobacteriales_bacterium_CG18_big_fil_WC_8_21_14_2_50_32_9.PIQ14548
Candidatus_Nealsonbacteria_bacterium_RBG_13_37_56.OGZ18192
Omnitrophica_bacterium_RIFOXYB12_FULL_50_7.OGX12059
Elusimicrobia_bacterium_GWC2_63_65.OGR73479
Elusimicrobia_bacterium_GWA2_62_23.OGR50626
Candidatus_Beckwithbacteria_bacterium_CG22_combo_CG10_13_8_21_14_all_01_47_9.PIP87985
Candidatus_Woesebacteria_bacterium_GWA1_41_8.OGM02102
Candidatus_Woesebacteria_bacterium_GW2011_GWA1_41_13b.KKR92039
Candidatus_Woykebacteria_bacterium_RBG_13_40_15.OGY22963
Parcubacteria_group_bacterium_GW2011_GWA1_36_12.KKP92826
Candidatus_Gottesmanbacteria_bacterium_RIFCSPHIGHO2_01_FULL_39_10.OGG13562
Candidatus_Roizmanbacteria_bacterium_CG_4_10_14_0_8_um_filter_36_36.PIY70583
Candidatus_Roizmanbacteria_bacterium_CG23_combo_of_CG06_09_8_20_14_all_35_49.PIP15093
uncultured_bacterium.EKE15184
Candidatus_Pacebacteria_bacterium_CG10_big_fil_rev_8_21_14_0_10_56_10.PIR58662
Candidatus_Lloydbacteria_bacterium_RIFCSPHIGHO2_01_FULL_54_11.OGZ06651
Proteobacteria_bacterium_ST_bin12.OQW68847
Rhodanobacter_sp_SCN_69_32.ODU75285
Gammaproteobacteria_bacterium_HGW_Gammaproteobacteria_2.PKM14947
CBU_1015
Burkholderiales_bacterium_RIFOXYD12_FULL_59_19.OGB54134
Legionellales_bacterium_RIFCSPHIGHO2_12_FULL_35_11.OGV35737
Coxiella_sp_DG_40.KPJ67187
Rhodobacteraceae_bacterium_KLH11.WP_008759414
Rhodobacteraceae_bacterium_KLH11.WP_008759125
Acetobacter_sp_DsW_063.OUJ16490
Acetobacter_sp_DsW_063.WP_086642922
Brevundimonas_sp_12_68_7.OYW40879
Rhodospirillum_centenum.WP_083759314
Rhodospirillum_centenum_SW.ACJ00148
Kaistia_adipata.WP_029077240
Chelativorans_sp_J32.WP_051469623
Mesorhizobium_sp_YR577.WP_091915832
Sphingomonas_sp_YR710.WP_093453904
Sphingobium_sp_Ndbn_10.WP_083199068
Sphingobium_herbicidovorans_NBRC_16415.KFG90029
Sphingobium_herbicidovorans.WP_081570293
Candidatus_Nomurabacteria_bacterium_GW2011_GWE1_35_16.KKP65518
candidate_division_WS6_bacterium_GW2011_GWA2_37_6.KKQ34843
Candidatus_Shapirobacteria_bacterium_CG11_big_fil_rev_8_21_14_0_20_40_12.PIQ70411
Candidatus_Roizmanbacteria_bacterium_CG10_big_fil_rev_8_21_14_0_10_39_12.PJE61520
Candidatus_Desantisbacteria_bacterium_CG23_combo_of_CG06_09_8_20_14_all_40_23.PIP40156
Candidatus_Rokubacteria_bacterium_RIFCSPHIGHO2_02_FULL_73_26.OGK95931
Gammaproteobacteria_bacterium_RIFCSPLOWO2_12_47_11.OGT71344
Planctomycetes_bacterium_RIFCSPHIGHO2_12_FULL_52_36.OHB93585
Berkelbacteria_bacterium_GW2011_GWA2_38_9.KKQ89879
Candidatus_Shapirobacteria_bacterium_CG07_land_8_20_14_0_80_39_12.PIU33444
Candidatus_Gottesmanbacteria_bacterium_GW2011_GWA2_43_14.KKS97817
Candidatus_Gottesmanbacteria_bacterium_RIFCSPHIGHO2_02_FULL_39_11.OGG15503
candidate_division_WWE3_bacterium_RIFCSPHIGHO2_01_FULL_48_15.OGC54732
Candidatus_Roizmanbacteria_bacterium_RIFCSPHIGHO2_12_FULL_44_10.OGK40454
uncultured_bacterium.EKD95308
candidate_division_WWE3_bacterium_RIFOXYB2_FULL_43_9.OGC73895
uncultured_bacterium.EKD99412
Candidatus_Woesebacteria_bacterium_RBG_16_42_24.OGM15911
uncultured_bacterium.EKE06334
Candidatus_Woesebacteria_bacterium_GW2011_GWA1_40_45.KKR63105
Candidatus_Woesebacteria_bacterium_GW2011_GWB1_40_101.KKR37912
Candidatus_Levybacteria_bacterium_RIFCSPHIGHO2_02_FULL_37_10.OGH17569
Candidatus_Levybacteria_bacterium_RIFCSPLOWO2_02_FULL_36_8b.OGH41457
Candidatus_Amesbacteria_bacterium_GW2011_GWA2_42_12.KKS32054
Candidatus_Daviesbacteria_bacterium_RIFCSPLOWO2_02_FULL_40_8.OGE67110
Parcubacteria_group_bacterium_GW2011_GWA2_47_10b.KKU53074
Parcubacteria_group_bacterium_RIFCSPHIGHO2_02_FULL_40_12.OHB19636
Parcubacteria_group_bacterium_RIFCSPHIGHO2_01_FULL_40_30.OHB17506
Candidatus_Nomurabacteria_bacterium_GW2011_GWA2_40_9.KKR79410
Candidatus_Taylorbacteria_bacterium_CG11_big_fil_rev_8_21_14_0_20_46_11.PIQ68034
Candidatus_Moranbacteria_bacterium_CG_4_9_14_3_um_filter_42_9.PJA87037
Candidatus_Moranbacteria_bacterium_GW2011_GWF2_35_39.KKP79668
B
uncultured_bacterium
Elusimicrobia
Elusimicrobia_bacterium_GWC2_63_65
Elusimicrobia_bacterium_GWA2_62_23
Flavobacteriales
Flavobacteriales_bacterium_CG18_big_fil_WC_8_21_14_2_50_32_9
Flavobacteriales_bacterium
Bacteria
Candidatus_Omnitrophica
Planctomycetes_bacterium_RIFCSPHIGHO2_12_FULL_52_36
Omnitrophica_WOR_2_bacterium_RIFCSPHIGHO2_02_FULL_68_15
Omnitrophica_bacterium_RIFOXYB12_FULL_50_7
Proteobacteria
Proteobacteria_bacterium_ST_bin12
Burkholderiales_bacterium_RIFOXYC12_FULL_60_6
Gammaproteobacteria
Rhodanobacter_sp_SCN-69-32
Gammaproteobacteria_bacterium_RIFCSPLOWO2_12_47_11
Gammaproteobacteria_bacterium_HGW-Gammaproteobacteria-2
Legionellales
Legionellales_bacterium_RIFCSPHIGHO2_12_FULL_35_11
Coxiella
Coxiella_sp_DG_40
Coxiella_burnetii_RSA_493
Alphaproteobacteria
Brevundimonas_subvibrioides
Rhodospirillales
Rhodospirillum
Rhodobacteraceae_bacterium_KLH11
Rhodospirillum_centenum_SW
Acetobacter_sp_DsW_063
Rhizobiales
Phyllobacteriaceae
Kaistia_adipata
Chelativorans_sp_J32
Mesorhizobium_sp_YR577
Sphingomonadaceae
Sphingobium
Sphingomonas_sp_YR710
Sphingobium_herbicidovorans_NBRC_16415
Sphingobium_sp_Ndbn-10
candidate_division_WS6_bacterium_GW2011_GWA2_37_6
Candidatus_Desantisbacteria_bacterium_CG23_combo_of_CG06-09_8_20_14_all_40_23
Berkelbacteria_bacterium_GW2011_GWA2_38_9
Candidatus_Rokubacteria_bacterium_RIFCSPHIGHO2_02_FULL_73_26
candidate_division_NC10_bacterium_RIFCSPLOWO2_02_FULL_66_22
candidate_division_WWE3
candidate_division_WWE3_bacterium_RIFOXYB2_FULL_43_9
candidate_division_WWE3_bacterium_RIFCSPHIGHO2_01_FULL_48_15
Candidatus_Lloydbacteria_bacterium_RIFCSPHIGHO2_01_FULL_54_11
Candidatus_Nealsonbacteria_bacterium_RBG_13_37_56
Candidatus_Taylorbacteria_bacterium_CG11_big_fil_rev_8_21_14_0_20_46_11
Candidatus_Moranbacteria
Candidatus_Moranbacteria_bacterium_CG_4_9_14_3_um_filter_42_9
Candidatus_Moranbacteria_bacterium_GW2011_GWF2_35_39
Candidatus_Nomurabacteria
Candidatus_Nomurabacteria_bacterium_GW2011_GWA2_40_9
Candidatus_Nomurabacteria_bacterium_GW2011_GWE1_35_16
Candidatus_Uhrbacteria
Candidatus_Uhrbacteria_bacterium_RIFCSPHIGHO2_12_FULL_54_23
Candidatus_Uhrbacteria_bacterium_RIFCSPLOWO2_01_FULL_55_36
Candidatus_Uhrbacteria_bacterium_RIFCSPLOWO2_02_FULL_54_37
Parcubacteria_group_bacterium_RIFCSPHIGHO2_02_FULL_40_12
Parcubacteria_group_bacterium_GW2011_GWA2_47_10b
Parcubacteria_group_bacterium_RIFCSPHIGHO2_01_FULL_40_30
Parcubacteria_group_bacterium_GW2011_GWA1_36_12
Candidatus_Daviesbacteria_bacterium_RIFCSPLOWO2_02_FULL_40_8
Candidatus_Amesbacteria_bacterium_GW2011_GWA2_42_12
Candidatus_Woykebacteria_bacterium_RBG_13_40_15
Candidatus_Beckwithbacteria_bacterium_CG_4_10_14_0_2_um_filter_47_25
Candidatus_Pacebacteria_bacterium_CG10_big_fil_rev_8_21_14_0_10_56_10
Candidatus_Shapirobacteria
Candidatus_Shapirobacteria_bacterium_CG_4_9_14_0_2_um_filter_40_11
Candidatus_Shapirobacteria_bacterium_CG07_land_8_20_14_0_80_39_12
Candidatus_Levybacteria
Candidatus_Levybacteria_bacterium_RIFCSPHIGHO2_02_FULL_37_10
Candidatus_Levybacteria_bacterium_RIFCSPLOWO2_02_FULL_36_8b
Candidatus_Gottesmanbacteria
Candidatus_Gottesmanbacteria_bacterium_GW2011_GWA2_43_14
Candidatus_Gottesmanbacteria_bacterium_GW2011_GWA2_47_9
Candidatus_Gottesmanbacteria_bacterium_RIFCSPHIGHO2_01_FULL_39_10
Candidatus_Gottesmanbacteria_bacterium_RIFCSPHIGHO2_02_FULL_39_11
Candidatus_Roizmanbacteria
Candidatus_Roizmanbacteria_bacterium_CG_4_10_14_0_8_um_filter_35_28
Candidatus_Roizmanbacteria_bacterium_CG_4_10_14_0_8_um_filter_36_36
Candidatus_Roizmanbacteria_bacterium_CG10_big_fil_rev_8_21_14_0_10_39_12
Candidatus_Roizmanbacteria_bacterium_RIFCSPHIGHO2_12_FULL_44_10
Candidatus_Woesebacteria
Candidatus_Woesebacteria_bacterium_GW2011_GWB1_40_101
Candidatus_Woesebacteria_bacterium_GWA1_41_8
Candidatus_Woesebacteria_bacterium_RBG_16_42_24
Candidatus_Woesebacteria_bacterium_GW2011_GWA1_40_45

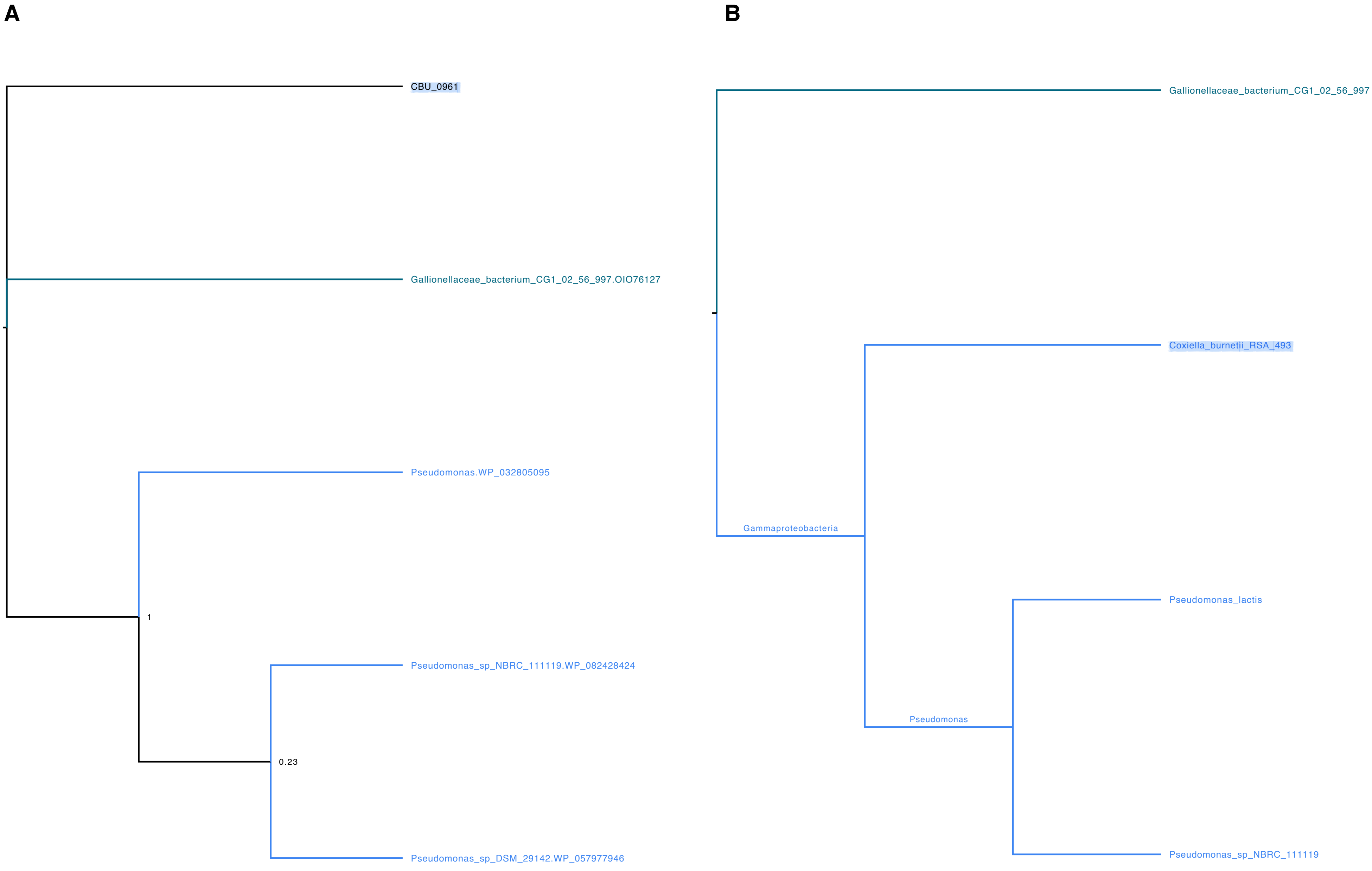
A
CBU_0961
Gallionellaceae_bacterium_CG1_02_56_997.OIO76127
Pseudomonas.WP_032805095
1
Pseudomonas_sp_NBRC_111119.WP_082428424
0.23
Pseudomonas_sp_DSM_29142.WP_057977946
B
Gallionellaceae_bacterium_CG1_02_56_997
Coxiella_burnetii_RSA_493
Gammaproteobacteria
Pseudomonas_lactis
Pseudomonas
Pseudomonas_sp_NBRC_111119

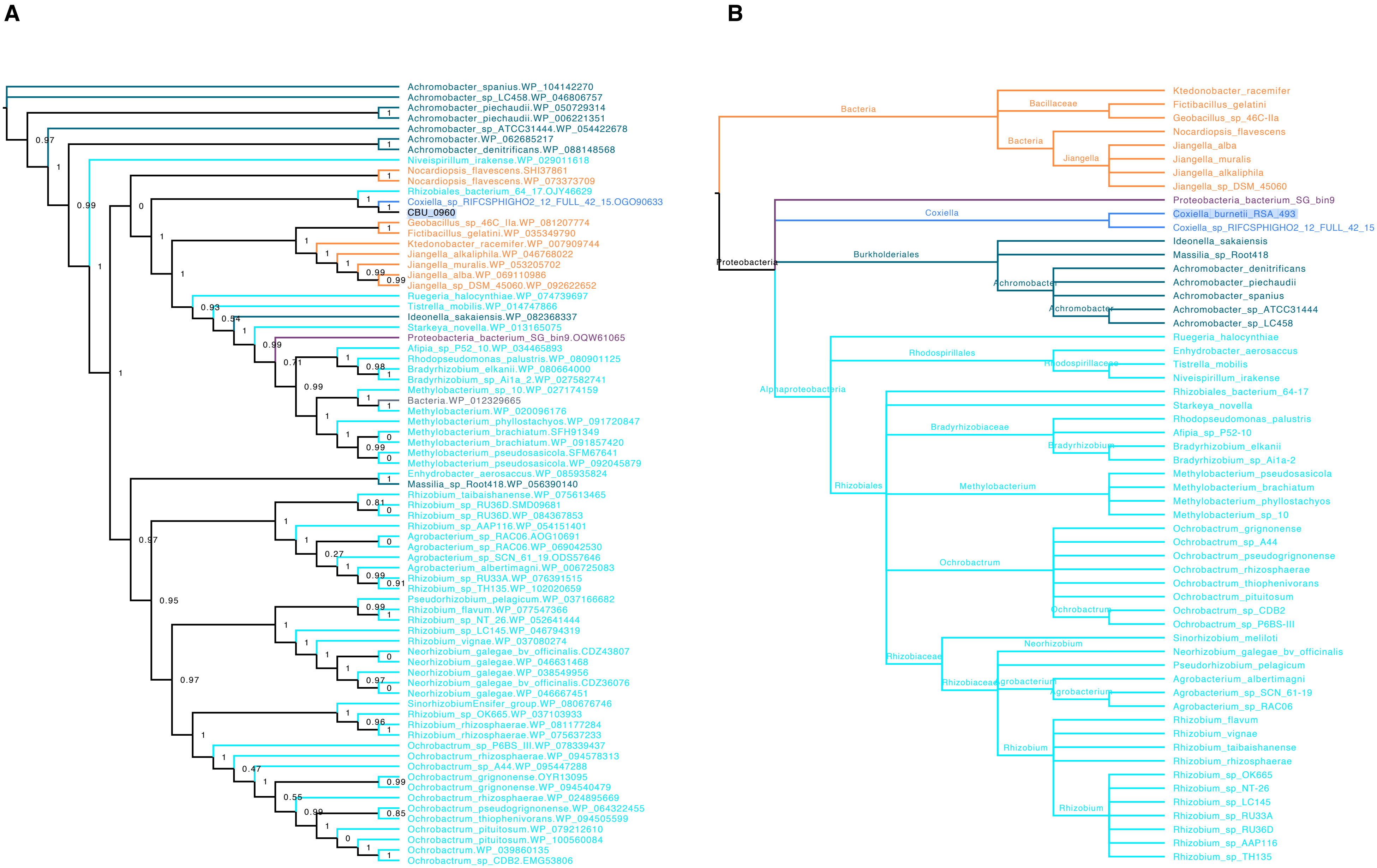
A
Achromobacter_spanius.WP_104142270
Achromobacter_sp_LC458.WP_046806757
Achromobacter_piechaudii.WP_050729314
Achromobacter_piechaudii.WP_006221351
Achromobacter_sp_ATCC31444.WP_054422678
Achromobacter.WP_062685217
Achromobacter_denitrificans.WP_088148568
Niveispirillum_irakense.WP_029011618
Nocardiopsis_flavescens.SHI37861
Nocardiopsis_flavescens.WP_073373709
Rhizobiales_bacterium_64_17.OJY46629
Coxiella_sp_RIFCSPHIGHO2_12_FULL_42_15.OGO90633
CBU_0960
Geobacillus_sp_46C_IIa.WP_081207774
Fictibacillus_gelatini.WP_035349790
Ktedonobacter_racemifer.WP_007909744
Jiangella_alkaliphila.WP_046768022
Jiangella_muralis.WP_053205702
Jiangella_alba.WP_069110986
Jiangella_sp_DSM_45060.WP_092622652
Ruegeria_halocynthiae.WP_074739697
Tistrella_mobilis.WP_014747866
Ideonella_sakaiensis.WP_082368337
Starkeya_novella.WP_013165075
Proteobacteria_bacterium_SG_bin9.OQW61065
Afipia_sp_P52_10.WP_034465893
Rhodopseudomonas_palustris.WP_080901125
Bradyrhizobium_elkanii.WP_080664000
Bradyrhizobium_sp_Ai1a_2.WP_027582741
Methylobacterium_sp_10.WP_027174159
Bacteria.WP_012329665
Methylobacterium.WP_020096176
Methylobacterium_phyllostachyos.WP_091720847
Methylobacterium_brachiatum.SFH91349
Methylobacterium_brachiatum.WP_091857420
Methylobacterium_pseudosasicola.SFM67641
Methylobacterium_pseudosasicola.WP_092045879
Enhydrobacter_aerosaccus.WP_085935824
Massilia_sp_Root418.WP_056390140
Rhizobium_taibaishanense.WP_075613465
Rhizobium_sp_RU36D.SMD09681
Rhizobium_sp_RU36D.WP_084367853
Rhizobium_sp_AAP116.WP_054151401
Agrobacterium_sp_RAC06.AOG10691
Agrobacterium_sp_RAC06.WP_069042530
Agrobacterium_sp_SCN_61_19.ODS57646
Agrobacterium_albertimagni.WP_006725083
Rhizobium_sp_RU33A.WP_076391515
Rhizobium_sp_TH135.WP_102020659
Pseudorhizobium_pelagicum.WP_037166682
Rhizobium_flavum.WP_077547366
Rhizobium_sp_NT_26.WP_052641444
Rhizobium_sp_LC145.WP_046794319
Rhizobium_vignae.WP_037080274
Neorhizobium_galegae_bv_officinalis.CDZ43807
Neorhizobium_galegae.WP_046631468
Neorhizobium_galegae.WP_038549956
Neorhizobium_galegae_bv_officinalis.CDZ36076
Neorhizobium_galegae.WP_046667451
SinorhizobiumEnsifer_group.WP_080676746
Rhizobium_sp_OK665.WP_037103933
Rhizobium_rhizosphaerae.WP_081177284
Rhizobium_rhizosphaerae.WP_075637233
Ochrobactrum_sp_P6BS_III.WP_078339437
Ochrobactrum_rhizosphaerae.WP_094578313
Ochrobactrum_sp_A44.WP_095447288
Ochrobactrum_grignonense.OYR13095
Ochrobactrum_grignonense.WP_094540479
Ochrobactrum_rhizosphaerae.WP_024895669
Ochrobactrum_pseudogrignonense.WP_064322455
Ochrobactrum_thiophenivorans.WP_094505599
Ochrobactrum_pituitosum.WP_079212610
Ochrobactrum_pituitosum.WP_100560084
Ochrobactrum.WP_039860135
Ochrobactrum_sp_CDB2.EMG53806
B
Bacteria
Bacillaceae
Jiangella
Proteobacteria
Coxiella
Burkholderiales
Achromobacter
Alphaproteobacteria
Rhodospirillales
Rhodospirillaceae
Rhizobiales
Bradyrhizobiaceae
Bradyrhizobium
Methylobacterium
Ochrobactrum
Rhizobiaceae
Neorhizobium
Agrobacterium
Rhizobium
Ktedonobacter_racemifer
Fictibacillus_gelatini
Geobacillus_sp_46C-IIa
Nocardiopsis_flavescens
Jiangella_alba
Jiangella_muralis
Jiangella_alkaliphila
Jiangella_sp_DSM_45060
Proteobacteria_bacterium_SG_bin9
Coxiella_burnetii_RSA_493
Coxiella_sp_RIFCSPHIGHO2_12_FULL_42_15
Ideonella_sakaiensis
Massilia_sp_Root418
Achromobacter_denitrificans
Achromobacter_piechaudii
Achromobacter_spanius
Achromobacter_sp_ATCC31444
Achromobacter_sp_LC458
Ruegeria_halocynthiae
Enhydrobacter_aerosaccus
Tistrella_mobilis
Niveispirillum_irakense
Rhizobiales_bacterium_64-17
Starkeya_novella
Rhodopseudomonas_palustris
Afipia_sp_P52-10
Bradyrhizobium_elkanii
Bradyrhizobium_sp_Ai1a-2
Methylobacterium_pseudosasicola
Methylobacterium_brachiatum
Methylobacterium_phyllostachyos
Methylobacterium_sp_10
Ochrobactrum_grignonense
Ochrobactrum_sp_A44
Ochrobactrum_pseudogrignonense
Ochrobactrum_rhizosphaerae
Ochrobactrum_thiophenivorans
Ochrobactrum_pituitosum
Ochrobactrum_sp_CDB2
Ochrobactrum_sp_P6BS-III
Sinorhizobium_meliloti
Neorhizobium_galegae_bv_officinalis
Pseudorhizobium_pelagicum
Agrobacterium_albertimagni
Agrobacterium_sp_SCN_61-19
Agrobacterium_sp_RAC06
Rhizobium_flavum
Rhizobium_vignae
Rhizobium_taibaishanense
Rhizobium_rhizosphaerae
Rhizobium_sp_OK665
Rhizobium_sp_NT-26
Rhizobium_sp_LC145
Rhizobium_sp_RU33A
Rhizobium_sp_RU36D
Rhizobium_sp_AAP116
Rhizobium_sp_TH135

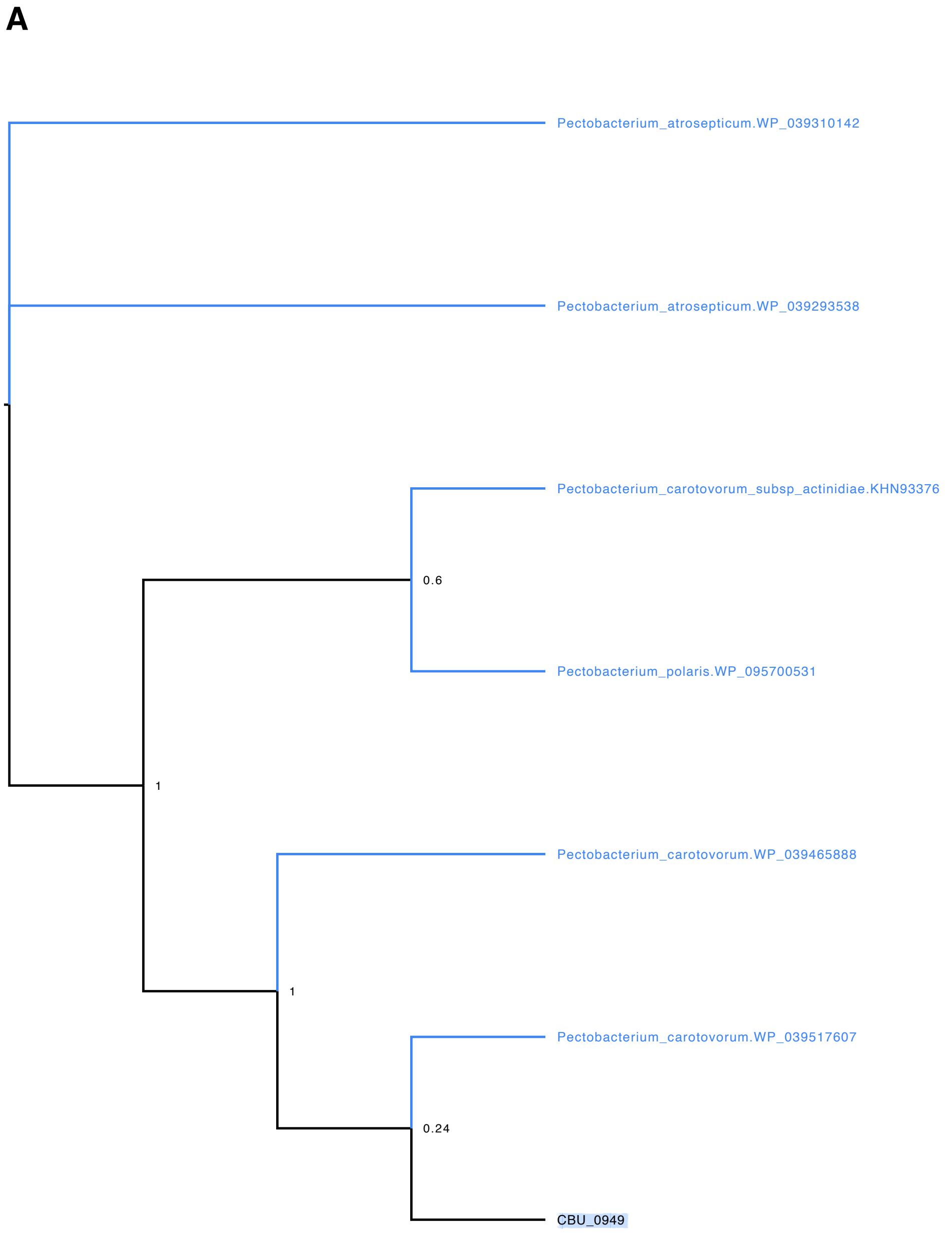
A
Pectobacterium_atrosepticum.WP_039310142
Pectobacterium_atrosepticum.WP_039293538
Pectobacterium_carotovorum_subsp_actinidiae.KHN93376
0.6
Pectobacterium_polaris.WP_095700531
1
Pectobacterium_carotovorum.WP_039465888
1
Pectobacterium_carotovorum.WP_039517607
0.24
CBU_0949

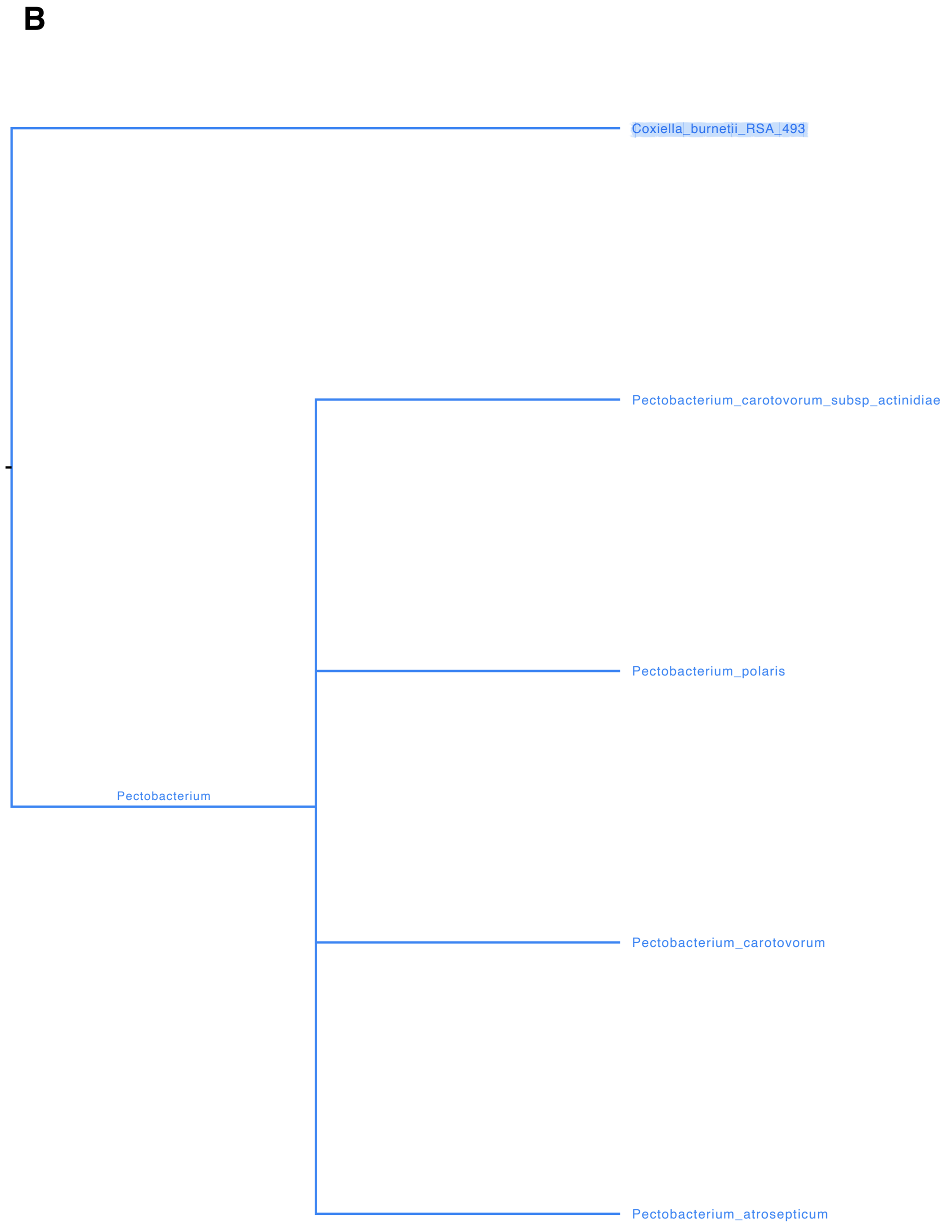
B
Coxiella_burnetii_RSA_493
Pectobacterium_carotovorum_subsp_actinidiae
Pectobacterium_polaris
Pectobacterium
Pectobacterium_carotovorum
Pectobacterium_atrosepticum

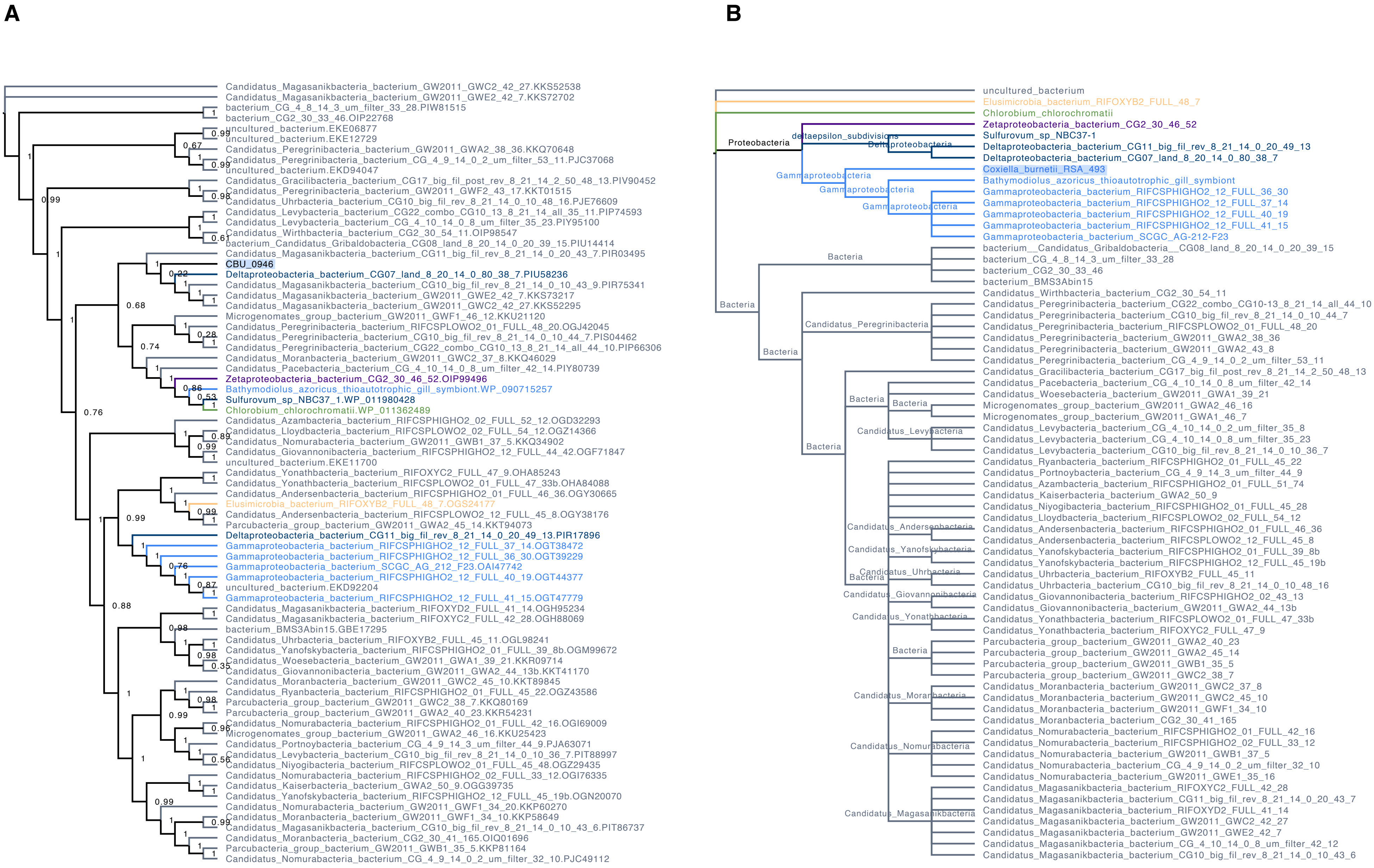
A
Candidatus_Magasanikbacteria_bacterium_GW2011_GWC2_42_27.KKS52538
Candidatus_Magasanikbacteria_bacterium_GW2011_GWE2_42_7.KKS72702
bacterium_CG_4_8_14_3_um_filter_33_28.PIW81515
bacterium_CG2_30_33_46.OIP22768
uncultured_bacterium.EKE06877
uncultured_bacterium.EKE12729
Candidatus_Peregrinibacteria_bacterium_GW2011_GWA2_38_36.KKQ70648
Candidatus_Peregrinibacteria_bacterium_CG_4_9_14_0_2_um_filter_53_11.PJC37068
uncultured_bacterium.EKD94047
Candidatus_Gracilibacteria_bacterium_CG17_big_fil_post_rev_8_21_14_2_50_48_13.PIV90452
Candidatus_Peregrinibacteria_bacterium_GW2011_GWF2_43_17.KKT01515
Candidatus_Uhrbacteria_bacterium_CG10_big_fil_rev_8_21_14_0_10_48_16.PJE76609
Candidatus_Levybacteria_bacterium_CG22_combo_CG10_13_8_21_14_all_35_11.PIP74593
Candidatus_Levybacteria_bacterium_CG_4_10_14_0_8_um_filter_35_23.PIY95100
Candidatus_Wirthbacteria_bacterium_CG2_30_54_11.OIP98547
bacterium_Candidatus_Gribaldobacteria_CG08_land_8_20_14_0_20_39_15.PIU14414
Candidatus_Magasanikbacteria_bacterium_CG11_big_fil_rev_8_21_14_0_20_43_7.PIR03495
CBU_0946
Deltaproteobacteria_bacterium_CG07_land_8_20_14_0_80_38_7.PIU58236
Candidatus_Magasanikbacteria_bacterium_CG10_big_fil_rev_8_21_14_0_10_43_9.PIR75341
Candidatus_Magasanikbacteria_bacterium_GW2011_GWE2_42_7.KKS73217
Candidatus_Magasanikbacteria_bacterium_GW2011_GWC2_42_27.KKS52295
Microgenomates_group_bacterium_GW2011_GWF1_46_12.KKU21120
Candidatus_Peregrinibacteria_bacterium_RIFCSPLOWO2_01_FULL_48_20.OGJ42045
Candidatus_Peregrinibacteria_bacterium_CG10_big_fil_rev_8_21_14_0_10_44_7.PIS04462
Candidatus_Peregrinibacteria_bacterium_CG22_combo_CG10_13_8_21_14_all_44_10.PIP66306
Candidatus_Moranbacteria_bacterium_GW2011_GWC2_37_8.KKQ46029
Candidatus_Pacebacteria_bacterium_CG_4_10_14_0_8_um_filter_42_14.PIY80739
Zetaproteobacteria_bacterium_CG2_30_46_52.OIP99496
Bathymodiolus_azoricus_thioautotrophic_gill_symbiont.WP_090715257
Sulfurovum_sp_NBC37_1.WP_011980428
Chlorobium_chlorochromatii.WP_011362489
Candidatus_Azambacteria_bacterium_RIFCSPHIGHO2_02_FULL_52_12.OGD32293
Candidatus_Lloydbacteria_bacterium_RIFCSPLOWO2_02_FULL_54_12.OGZ14366
Candidatus_Nomurabacteria_bacterium_GW2011_GWB1_37_5.KKQ34902
Candidatus_Giovannonibacteria_bacterium_RIFCSPHIGHO2_12_FULL_44_42.OGF71847
uncultured_bacterium.EKE11700
Candidatus_Yonathbacteria_bacterium_RIFOXYC2_FULL_47_9.OHA85243
Candidatus_Yonathbacteria_bacterium_RIFCSPLOWO2_01_FULL_47_33b.OHA84088
Candidatus_Andersenbacteria_bacterium_RIFCSPLOWO2_01_FULL_46_36.OGY30665
Elusimicrobia_bacterium_RIFOXYB2_FULL_48_7.OGS24177
Candidatus_Andersenbacteria_bacterium_RIFCSPLOWO2_12_FULL_45_8.OGY38176
Parcubacteria_group_bacterium_GW2011_GWA2_45_14.KKT94073
Deltaproteobacteria_bacterium_CG11_big_fil_rev_8_21_14_0_20_49_13.PIR17896
Gammaproteobacteria_bacterium_RIFCSPHIGHO2_12_FULL_37_14.OGT38472
Gammaproteobacteria_bacterium_RIFCSPHIGHO2_12_FULL_36_30.OGT39229
Gammaproteobacteria_bacterium_SCGC_AG_212_F23.OAI47742
Gammaproteobacteria_bacterium_RIFCSPHIGHO2_12_FULL_40_19.OGT44377
uncultured_bacterium.EKD92204
Gammaproteobacteria_bacterium_RIFCSPHIGHO2_12_FULL_41_15.OGT47779
Candidatus_Magasanikbacteria_bacterium_RIFOXYD2_FULL_41_14.OGH95234
Candidatus_Magasanikbacteria_bacterium_RIFOXYC2_FULL_42_28.OGH88069
bacterium_BMS3Abin15.GBE17295
Candidatus_Uhrbacteria_bacterium_RIFOXYB2_FULL_45_11.OGL98241
Candidatus_Yanofskybacteria_bacterium_RIFCSPHIGHO2_01_FULL_39_8b.OGM99672
Candidatus_Woesebacteria_bacterium_GW2011_GWA1_39_21.KKR09714
Candidatus_Giovannonibacteria_bacterium_GW2011_GWA2_44_13b.KKT41170
Candidatus_Moranbacteria_bacterium_GW2011_GWC2_45_10.KKT89845
Candidatus_Ryanbacteria_bacterium_RIFCSPHIGHO2_01_FULL_45_22.OGZ43586
Parcubacteria_group_bacterium_GW2011_GWC2_38_7.KKQ80169
Parcubacteria_group_bacterium_GW2011_GWA2_40_23.KKR54231
Candidatus_Nomurabacteria_bacterium_RIFCSPHIGHO2_01_FULL_42_16.OGI69009
Microgenomates_group_bacterium_GW2011_GWA2_46_16.KKU25423
Candidatus_Portnoybacteria_bacterium_CG_4_9_14_3_um_filter_44_9.PJA63071
Candidatus_Levybacteria_bacterium_CG10_big_fil_rev_8_21_14_0_10_36_7.PIT88997
Candidatus_Niyogibacteria_bacterium_RIFCSPLOWO2_01_FULL_45_48.OGZ29435
Candidatus_Nomurabacteria_bacterium_RIFCSPHIGHO2_02_FULL_33_12.OGI76335
Candidatus_Kaiserbacteria_bacterium_GWA2_50_9.OGG39735
Candidatus_Yanofskybacteria_bacterium_RIFCSPHIGHO2_12_FULL_45_19b.OGN20070
Candidatus_Nomurabacteria_bacterium_GW2011_GWF1_34_20.KKP60270
Candidatus_Moranbacteria_bacterium_GW2011_GWF1_34_10.KKP58649
Candidatus_Magasanikbacteria_bacterium_CG10_big_fil_rev_8_21_14_0_10_43_6.PIT86737
Candidatus_Moranbacteria_bacterium_CG2_30_41_165.OIQ01696
Parcubacteria_group_bacterium_GW2011_GWB1_35_5.KKP81164
Candidatus_Nomurabacteria_bacterium_CG_4_9_14_0_2_um_filter_32_10.PJC49112
B
uncultured_bacterium
Elusimicrobia_bacterium_RIFOXYB2_FULL_48_7
Chlorobium_chlorochromatii
Zetaproteobacteria_bacterium_CG2_30_46_52
Sulfurovum_sp_NBC37-1
Deltaproteobacteria_bacterium_CG11_big_fil_rev_8_21_14_0_20_49_13
Deltaproteobacteria_bacterium_CG07_land_8_20_14_0_80_38_7
Coxiella_burnetii_RSA_493
Bathymodiolus_azoricus_thioautotrophic_gill_symbiont
Gammaproteobacteria_bacterium_RIFCSPHIGHO2_12_FULL_36_30
Gammaproteobacteria_bacterium_RIFCSPHIGHO2_12_FULL_37_14
Gammaproteobacteria_bacterium_RIFCSPHIGHO2_12_FULL_40_19
Gammaproteobacteria_bacterium_RIFCSPHIGHO2_12_FULL_41_15
Gammaproteobacteria_bacterium_SCGC_AG-212-F23
bacterium__Candidatus_Gribaldobacteria__CG08_land_8_20_14_0_20_39_15
bacterium_CG_4_8_14_3_um_filter_33_28
bacterium_CG2_30_33_46
bacterium_BMS3Abin15
Candidatus_Wirthbacteria_bacterium_CG2_30_54_11
Candidatus_Peregrinibacteria_bacterium_CG22_combo_CG10-13_8_21_14_all_44_10
Candidatus_Peregrinibacteria_bacterium_CG10_big_fil_rev_8_21_14_0_10_44_7
Candidatus_Peregrinibacteria_bacterium_RIFCSPLOWO2_01_FULL_48_20
Candidatus_Peregrinibacteria_bacterium_GW2011_GWA2_38_36
Candidatus_Peregrinibacteria_bacterium_GW2011_GWA2_43_8
Candidatus_Peregrinibacteria_bacterium_CG_4_9_14_0_2_um_filter_53_11
Candidatus_Gracilibacteria_bacterium_CG17_big_fil_post_rev_8_21_14_2_50_48_13
Candidatus_Pacebacteria_bacterium_CG_4_10_14_0_8_um_filter_42_14
Candidatus_Woesebacteria_bacterium_GW2011_GWA1_39_21
Microgenomates_group_bacterium_GW2011_GWA2_46_16
Microgenomates_group_bacterium_GW2011_GWA1_46_7
Candidatus_Levybacteria_bacterium_CG_4_10_14_0_2_um_filter_35_8
Candidatus_Levybacteria_bacterium_CG_4_10_14_0_8_um_filter_35_23
Candidatus_Levybacteria_bacterium_CG10_big_fil_rev_8_21_14_0_10_36_7
Candidatus_Ryanbacteria_bacterium_RIFCSPHIGHO2_01_FULL_45_22
Candidatus_Portnoybacteria_bacterium_CG_4_9_14_3_um_filter_44_9
Candidatus_Azambacteria_bacterium_RIFCSPHIGHO2_01_FULL_51_74
Candidatus_Kaiserbacteria_bacterium_GWA2_50_9
Candidatus_Niyogibacteria_bacterium_RIFCSPHIGHO2_01_FULL_45_28
Candidatus_Lloydbacteria_bacterium_RIFCSPLOWO2_02_FULL_54_12
Candidatus_Andersenbacteria_bacterium_RIFCSPHIGHO2_01_FULL_46_36
Candidatus_Andersenbacteria_bacterium_RIFCSPLOWO2_12_FULL_45_8
Candidatus_Yanofskybacteria_bacterium_RIFCSPHIGHO2_01_FULL_39_8b
Candidatus_Yanofskybacteria_bacterium_RIFCSPHIGHO2_12_FULL_45_19b
Candidatus_Uhrbacteria_bacterium_RIFOXYB2_FULL_45_11
Candidatus_Uhrbacteria_bacterium_CG10_big_fil_rev_8_21_14_0_10_48_16
Candidatus_Giovannonibacteria_bacterium_RIFCSPHIGHO2_02_43_13
Candidatus_Giovannonibacteria_bacterium_GW2011_GWA2_44_13b
Candidatus_Yonathbacteria_bacterium_RIFCSPLOWO2_01_FULL_47_33b
Candidatus_Yonathbacteria_bacterium_RIFOXYC2_FULL_47_9
Parcubacteria_group_bacterium_GW2011_GWA2_40_23
Parcubacteria_group_bacterium_GW2011_GWA2_45_14
Parcubacteria_group_bacterium_GW2011_GWB1_35_5
Parcubacteria_group_bacterium_GW2011_GWC2_38_7
Candidatus_Moranbacteria_bacterium_GW2011_GWC2_37_8
Candidatus_Moranbacteria_bacterium_GW2011_GWC2_45_10
Candidatus_Moranbacteria_bacterium_GW2011_GWF1_34_10
Candidatus_Moranbacteria_bacterium_CG2_30_41_165
Candidatus_Nomurabacteria_bacterium_RIFCSPHIGHO2_01_FULL_42_16
Candidatus_Nomurabacteria_bacterium_RIFCSPHIGHO2_02_FULL_33_12
Candidatus_Nomurabacteria_bacterium_GW2011_GWB1_37_5
Candidatus_Nomurabacteria_bacterium_CG_4_9_14_0_2_um_filter_32_10
Candidatus_Nomurabacteria_bacterium_GW2011_GWE1_35_16
Candidatus_Magasanikbacteria_bacterium_RIFOXYC2_FULL_42_28
Candidatus_Magasanikbacteria_bacterium_CG11_big_fil_rev_8_21_14_0_20_43_7
Candidatus_Magasanikbacteria_bacterium_RIFOXYD2_FULL_41_14
Candidatus_Magasanikbacteria_bacterium_GW2011_GWC2_42_27
Candidatus_Magasanikbacteria_bacterium_GW2011_GWE2_42_7
Candidatus_Magasanikbacteria_bacterium_CG_4_10_14_0_8_um_filter_42_12
Candidatus_Magasanikbacteria_bacterium_CG10_big_fil_rev_8_21_14_0_10_43_6
Proteobacteria
deltaepsilon_subdivisions
Deltaproteobacteria
Gammaproteobacteria
Bacteria
Candidatus_Peregrinibacteria
Candidatus_Levybacteria
Candidatus_Andersenbacteria
Candidatus_Yanofskybacteria
Candidatus_Uhrbacteria
Candidatus_Giovannonibacteria
Candidatus_Yonathbacteria
Candidatus_Moranbacteria
Candidatus_Nomurabacteria
Candidatus_Magasanikbacteria

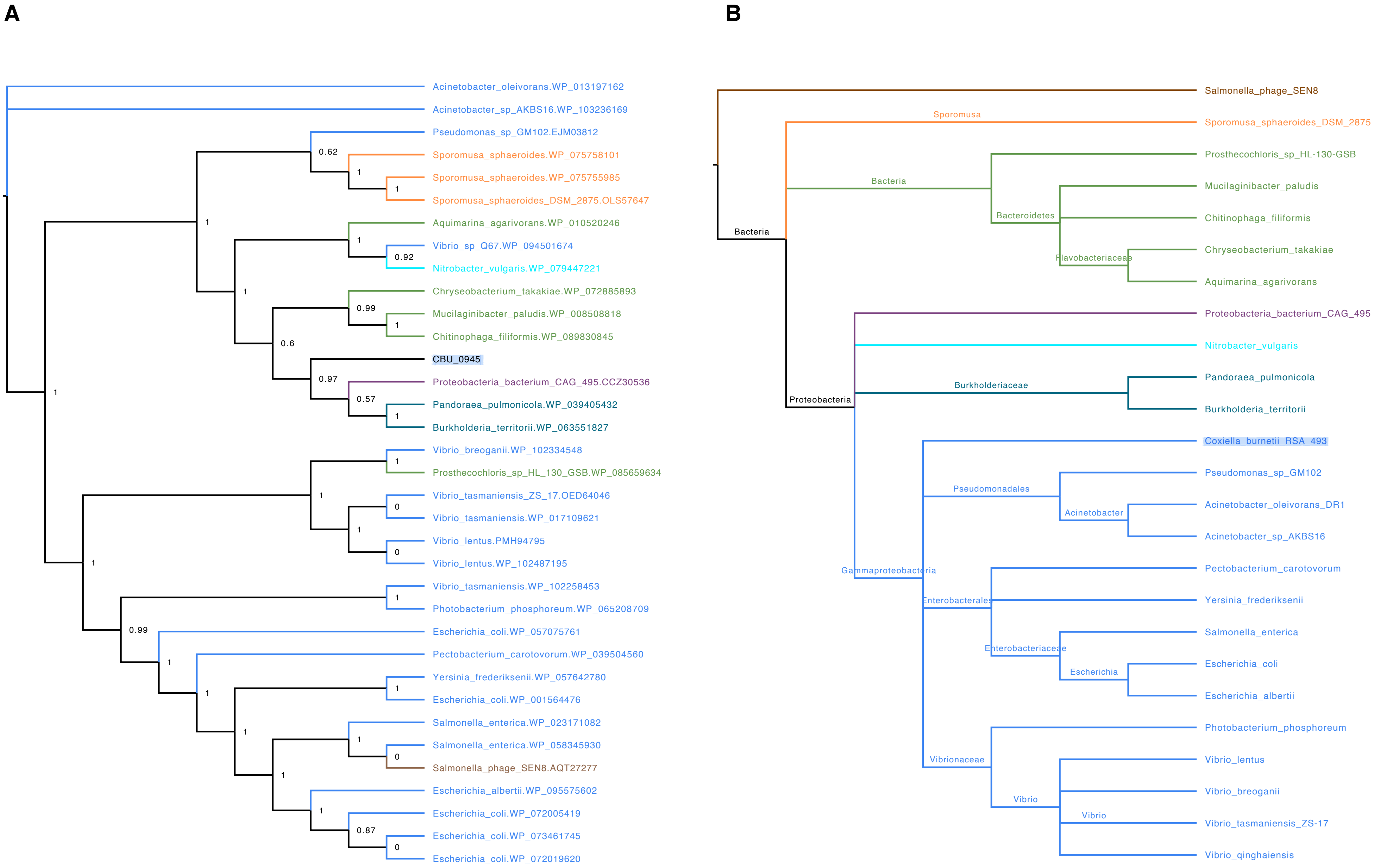
A
Acinetobacter_oleivorans.WP_013197162
Acinetobacter_sp_AKBS16.WP_103236169
Pseudomonas_sp_GM102.EJM03812
Sporomusa_sphaeroides.WP_075758101
Sporomusa_sphaeroides.WP_075755985
Sporomusa_sphaeroides_DSM_2875.OLS57647
Aquimarina_agarivorans.WP_010520246
Vibrio_sp_Q67.WP_094501674
Nitrobacter_vulgaris.WP_079447221
Chryseobacterium_takakiae.WP_072885893
Mucilaginibacter_paludis.WP_008508818
Chitinophaga_filiformis.WP_089830845
CBU_0945
Proteobacteria_bacterium_CAG_495.CCZ30536
Pandoraea_pulmonicola.WP_039405432
Burkholderia_territorii.WP_063551827
Vibrio_breoganii.WP_102334548
Prosthecochloris_sp_HL_130_GSB.WP_085659634
Vibrio_tasmaniensis_ZS_17.OED64046
Vibrio_tasmaniensis.WP_017109621
Vibrio_lentus.PMH94795
Vibrio_lentus.WP_102487195
Vibrio_tasmaniensis.WP_102258453
Photobacterium_phosphoreum.WP_065208709
Escherichia_coli.WP_057075761
Pectobacterium_carotovorum.WP_039504560
Yersinia_frederiksenii.WP_057642780
Escherichia_coli.WP_001564476
Salmonella_enterica.WP_023171082
Salmonella_enterica.WP_058345930
Salmonella_phage_SEN8.AQT27277
Escherichia_albertii.WP_095575602
Escherichia_coli.WP_072005419
Escherichia_coli.WP_073461745
Escherichia_coli.WP_072019620
B
Salmonella_phage_SEN8
Sporomusa
Sporomusa_sphaeroides_DSM_2875
Bacteria
Prosthecochloris_sp_HL-130-GSB
Mucilaginibacter_paludis
Bacteroidetes
Chitinophaga_filiformis
Bacteria
Chryseobacterium_takakiae
Flavobacteriaceae
Aquimarina_agarivorans
Proteobacteria_bacterium_CAG_495
Nitrobacter_vulgaris
Burkholderiaceae
Pandoraea_pulmonicola
Proteobacteria
Burkholderia_territorii
Coxiella_burnetii_RSA_493
Pseudomonadales
Pseudomonas_sp_GM102
Acinetobacter
Acinetobacter_oleivorans_DR1
Acinetobacter_sp_AKBS16
Gammaproteobacteria
Pectobacterium_carotovorum
Enterobacterales
Yersinia_frederiksenii
Salmonella_enterica
Enterobacteriaceae
Escherichia
Escherichia_coli
Escherichia_albertii
Photobacterium_phosphoreum
Vibrionaceae
Vibrio_lentus
Vibrio_breoganii
Vibrio
Vibrio
Vibrio_tasmaniensis_ZS-17
Vibrio_qinghaiensis

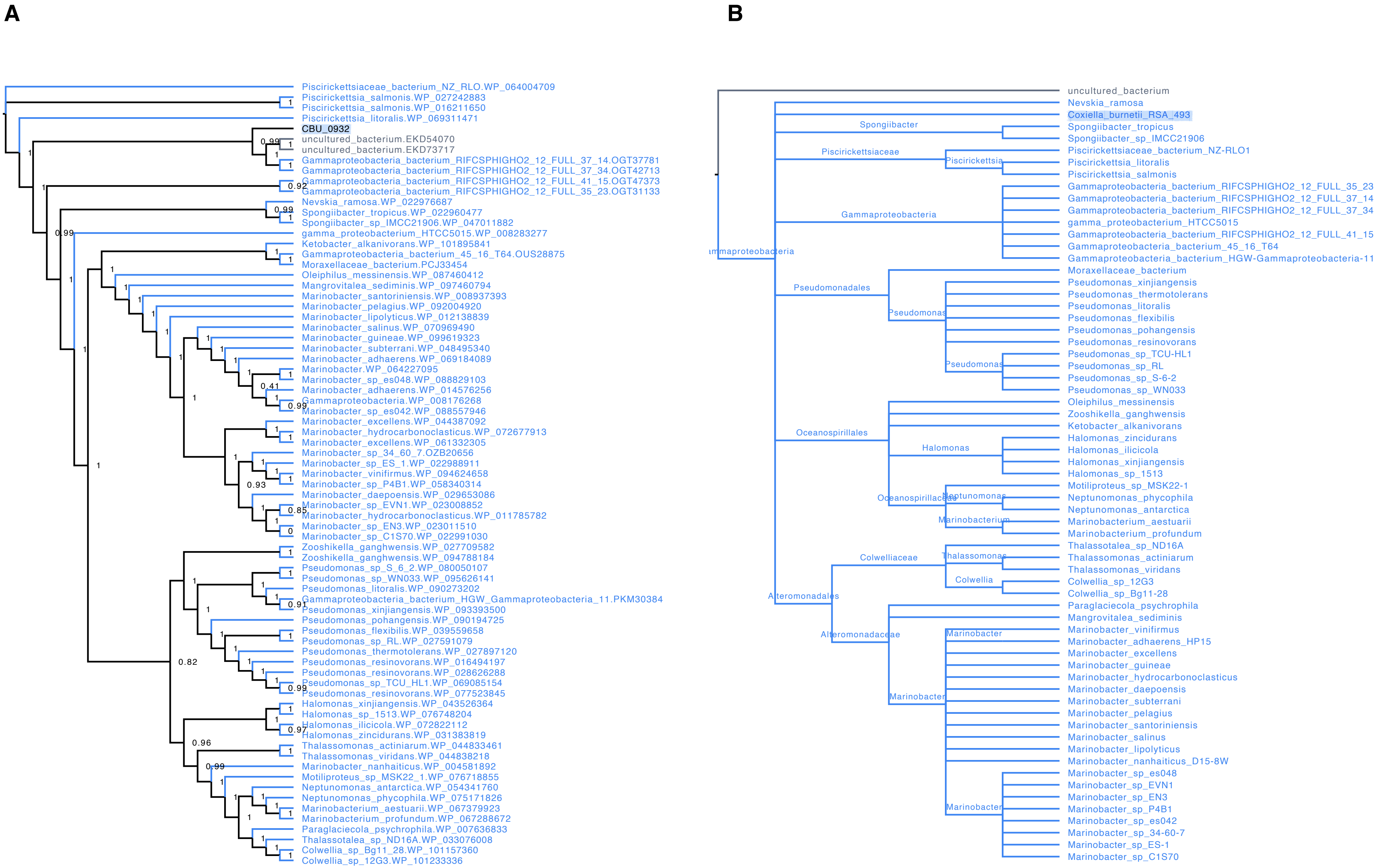
A
Piscirickettsiaceae_bacterium_NZ_RLO.WP_064004709
Piscirickettsia_salmonis.WP_027242883
Piscirickettsia_salmonis.WP_016211650
Piscirickettsia_litoralis.WP_069311471
CBU_0932
uncultured_bacterium.EKD54070
uncultured_bacterium.EKD73717
Gammaproteobacteria_bacterium_RIFCSPHIGHO2_12_FULL_37_14.OGT37781
Gammaproteobacteria_bacterium_RIFCSPHIGHO2_12_FULL_37_34.OGT42713
Gammaproteobacteria_bacterium_RIFCSPHIGHO2_12_FULL_41_15.OGT47373
Gammaproteobacteria_bacterium_RIFCSPHIGHO2_12_FULL_35_23.OGT31133
Nevskia_ramosa.WP_022976687
Spongiibacter_tropicus.WP_022960477
Spongiibacter_sp_IMCC21906.WP_047011882
gamma_proteobacterium_HTCC5015.WP_008283277
Ketobacter_alkanivorans.WP_101895841
Gammaproteobacteria_bacterium_45_16_T64.OUS28875
Moraxellaceae_bacterium.PCJ33454
Oleiphilus_messinensis.WP_087460412
Mangrovitalea_sediminis.WP_097460794
Marinobacter_santoriniensis.WP_008937393
Marinobacter_pelagius.WP_092004920
Marinobacter_lipolyticus.WP_012138839
Marinobacter_salinus.WP_070969490
Marinobacter_guineae.WP_099619323
Marinobacter_subterrani.WP_048495340
Marinobacter_adhaerens.WP_069184089
Marinobacter.WP_064227095
Marinobacter_sp_es048.WP_088829103
Marinobacter_adhaerens.WP_014576256
Gammaproteobacteria.WP_008176268
Marinobacter_sp_es042.WP_088557946
Marinobacter_excellens.WP_044387092
Marinobacter_hydrocarbonoclasticus.WP_072677913
Marinobacter_excellens.WP_061332305
Marinobacter_sp_34_60_7.OZB20656
Marinobacter_sp_ES_1.WP_022988911
Marinobacter_vinifirmus.WP_094624658
Marinobacter_sp_P4B1.WP_058340314
Marinobacter_daepoensis.WP_029653086
Marinobacter_sp_EVN1.WP_023008852
Marinobacter_hydrocarbonoclasticus.WP_011785782
Marinobacter_sp_EN3.WP_023011510
Marinobacter_sp_C1S70.WP_022991030
Zooshikella_ganghwensis.WP_027709582
Zooshikella_ganghwensis.WP_094788184
Pseudomonas_sp_S_6_2.WP_080050107
Pseudomonas_sp_WN033.WP_095626141
Pseudomonas_litoralis.WP_090273202
Gammaproteobacteria_bacterium_HGW_Gammaproteobacteria_11.PKM30384
Pseudomonas_xinjiangensis.WP_093393500
Pseudomonas_pohangensis.WP_090194725
Pseudomonas_flexibilis.WP_039559658
Pseudomonas_sp_RL.WP_027591079
Pseudomonas_thermotolerans.WP_027897120
Pseudomonas_resinovorans.WP_016494197
Pseudomonas_resinovorans.WP_028626288
Pseudomonas_sp_TCU_HL1.WP_069085154
Pseudomonas_resinovorans.WP_077523845
Halomonas_xinjiangensis.WP_043526364
Halomonas_sp_1513.WP_076748204
Halomonas_ilicicola.WP_072822112
Halomonas_zincidurans.WP_031383819
Thalassomonas_actiniarum.WP_044833461
Thalassomonas_viridans.WP_044838218
Marinobacter_nanhaiticus.WP_004581892
Motiliproteus_sp_MSK22_1.WP_076718855
Neptunomonas_antarctica.WP_054341760
Neptunomonas_phycophila.WP_075171826
Marinobacterium_aestuarii.WP_067379923
Marinobacterium_profundum.WP_067288672
Paraglaciecola_psychrophila.WP_007636833
Thalassotalea_sp_ND16A.WP_033076008
Colwellia_sp_Bg11_28.WP_101157360
Colwellia_sp_12G3.WP_101233336
B
uncultured_bacterium
Nevskia_ramosa
Coxiella_burnetii_RSA_493
Spongiibacter
Spongiibacter_tropicus
Spongiibacter_sp_IMCC21906
Piscirickettsiaceae
Piscirickettsia
Piscirickettsiaceae_bacterium_NZ-RLO1
Piscirickettsia_litoralis
Piscirickettsia_salmonis
Gammaproteobacteria
Gammaproteobacteria_bacterium_RIFCSPHIGHO2_12_FULL_35_23
Gammaproteobacteria_bacterium_RIFCSPHIGHO2_12_FULL_37_14
Gammaproteobacteria_bacterium_RIFCSPHIGHO2_12_FULL_37_34
gamma_proteobacterium_HTCC5015
Gammaproteobacteria_bacterium_RIFCSPHIGHO2_12_FULL_41_15
Gammaproteobacteria_bacterium_45_16_T64
Gammaproteobacteria_bacterium_HGW-Gammaproteobacteria-11
Pseudomonadales
Moraxellaceae_bacterium
Pseudomonas
Pseudomonas_xinjiangensis
Pseudomonas_thermotolerans
Pseudomonas_litoralis
Pseudomonas_flexibilis
Pseudomonas_pohangensis
Pseudomonas_resinovorans
Pseudomonas_sp_TCU-HL1
Pseudomonas_sp_RL
Pseudomonas_sp_S-6-2
Pseudomonas_sp_WN033
Oceanospirillales
Oleiphilus_messinensis
Zooshikella_ganghwensis
Ketobacter_alkanivorans
Halomonas
Halomonas_zincidurans
Halomonas_ilicicola
Halomonas_xinjiangensis
Halomonas_sp_1513
Oceanospirillaceae
Neptunomonas
Motiliproteus_sp_MSK22-1
Neptunomonas_phycophila
Neptunomonas_antarctica
Marinobacterium
Marinobacterium_aestuarii
Marinobacterium_profundum
Alteromonadales
Colwelliaceae
Thalassomonas
Thalassotalea_sp_ND16A
Thalassomonas_actiniarum
Thalassomonas_viridans
Colwellia
Colwellia_sp_12G3
Colwellia_sp_Bg11-28
Alteromonadaceae
Paraglaciecola_psychrophila
Mangrovitalea_sediminis
Marinobacter
Marinobacter_vinifirmus
Marinobacter_adhaerens_HP15
Marinobacter_excellens
Marinobacter_guineae
Marinobacter_hydrocarbonoclasticus
Marinobacter_daepoensis
Marinobacter_subterrani
Marinobacter_pelagius
Marinobacter_santoriniensis
Marinobacter_salinus
Marinobacter_lipolyticus
Marinobacter_nanhaiticus_D15-8W
Marinobacter_sp_es048
Marinobacter_sp_EVN1
Marinobacter_sp_EN3
Marinobacter_sp_P4B1
Marinobacter_sp_es042
Marinobacter_sp_34-60-7
Marinobacter_sp_ES-1
Marinobacter_sp_C1S70

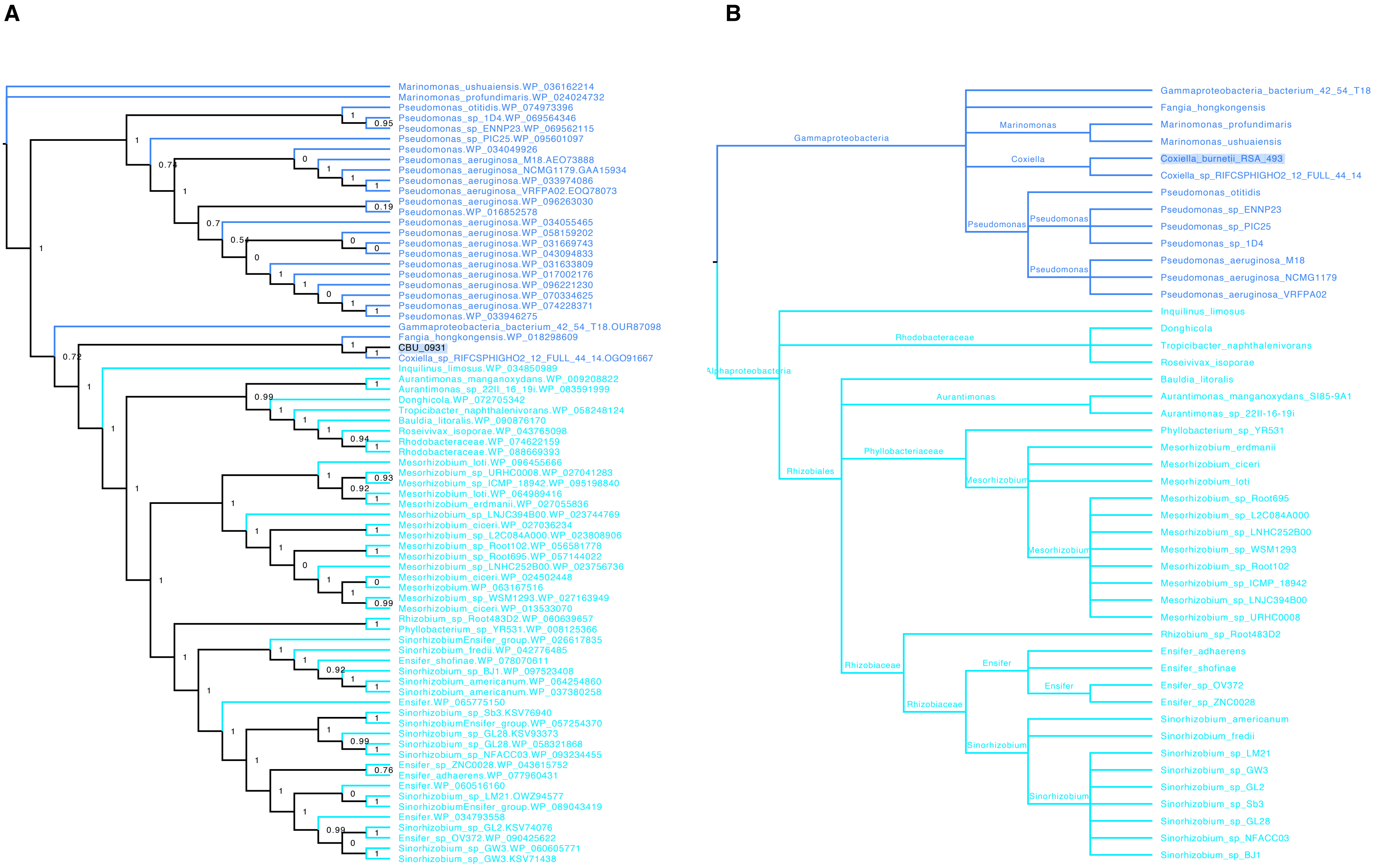

A
Marinomonas_ushuaiensis.WP_036162214
Marinomonas_profundimaris.WP_024024732
Pseudomonas_otitidis.WP_074973396
Pseudomonas_sp_1D4.WP_069564346
Pseudomonas_sp_ENNP23.WP_069562115
Pseudomonas_sp_PIC25.WP_095601097
Pseudomonas.WP_034049926
Pseudomonas_aeruginosa_M18.AEO73888
Pseudomonas_aeruginosa_NCMG1179.GAA15934
Pseudomonas_aeruginosa.WP_033974086
Pseudomonas_aeruginosa_VRFPA02.EOQ78073
Pseudomonas_aeruginosa.WP_096263030
Pseudomonas.WP_016852578
Pseudomonas_aeruginosa.WP_034055465
Pseudomonas_aeruginosa.WP_058159202
Pseudomonas_aeruginosa.WP_031669743
Pseudomonas_aeruginosa.WP_043094833
Pseudomonas_aeruginosa.WP_031633809
Pseudomonas_aeruginosa.WP_017002176
Pseudomonas_aeruginosa.WP_096221230
Pseudomonas_aeruginosa.WP_070334625
Pseudomonas_aeruginosa.WP_074228371
Pseudomonas.WP_033946275
Gammaproteobacteria_bacterium_42_54_T18.OUR87098
Fangia_hongkongensis.WP_018298609
CBU_0931
Coxiella_sp_RIFCSPHIGHO2_12_FULL_44_14.OGO91667
Inquilinus_limosus.WP_034850989
Aurantimonas_manganoxydans.WP_009208822
Aurantimonas_sp_22II_16_19i.WP_083591999
Donghicola.WP_072705342
Tropicibacter_naphthalenivorans.WP_058248124
Bauldia_litoralis.WP_090876170
Roseivivax_isoporae.WP_043765098
Rhodobacteraceae.WP_074622159
Rhodobacteraceae.WP_088669393
Mesorhizobium_loti.WP_096455666
Mesorhizobium_sp_URHC0008.WP_027041283
Mesorhizobium_sp_ICMP_18942.WP_095198840
Mesorhizobium_loti.WP_064989416
Mesorhizobium_erdmanii.WP_027055836
Mesorhizobium_sp_LNJC394B00.WP_023744769
Mesorhizobium_ciceri.WP_027036234
Mesorhizobium_sp_L2C084A000.WP_023808906
Mesorhizobium_sp_Root102.WP_056581778
Mesorhizobium_sp_Root695.WP_057144022
Mesorhizobium_sp_LNHC252B00.WP_023756736
Mesorhizobium_ciceri.WP_024502448
Mesorhizobium.WP_063167516
Mesorhizobium_sp_WSM1293.WP_027163949
Mesorhizobium_ciceri.WP_013533070
Rhizobium_sp_Root483D2.WP_060639657
Phyllobacterium_sp_YR531.WP_008125366
SinorhizobiumEnsifer_group.WP_026617835
Sinorhizobium_fredii.WP_042776485
Ensifer_shofinae.WP_078070611
Sinorhizobium_sp_BJ1.WP_097523408
Sinorhizobium_americanum.WP_064254860
Sinorhizobium_americanum.WP_037380258
Ensifer.WP_065775150
Sinorhizobium_sp_Sb3.KSV76940
SinorhizobiumEnsifer_group.WP_057254370
Sinorhizobium_sp_GL28.KSV93373
Sinorhizobium_sp_GL28.WP_058321868
Sinorhizobium_sp_NFACC03.WP_093234455
Ensifer_sp_ZNC0028.WP_043615752
Ensifer_adhaerens.WP_077960431
Ensifer.WP_060516160
Sinorhizobium_sp_LM21.OWZ94577
SinorhizobiumEnsifer_group.WP_089043419
Ensifer.WP_034793558
Sinorhizobium_sp_GL2.KSV74076
Ensifer_sp_OV372.WP_090425622
Sinorhizobium_sp_GW3.WP_060605771
Sinorhizobium_sp_GW3.KSV71438
B
Gammaproteobacteria
Alphaproteobacteria
Marinomonas
Coxiella
Pseudomonas
Rhodobacteraceae
Aurantimonas
Phyllobacteriaceae
Rhizobiales
Mesorhizobium
Rhizobiaceae
Ensifer
Sinorhizobium
Gammaproteobacteria_bacterium_42_54_T18
Fangia_hongkongensis
Marinomonas_profundimaris
Marinomonas_ushuaiensis
Coxiella_burnetii_RSA_493
Coxiella_sp_RIFCSPHIGHO2_12_FULL_44_14
Pseudomonas_otitidis
Pseudomonas_sp_ENNP23
Pseudomonas_sp_PIC25
Pseudomonas_sp_1D4
Pseudomonas_aeruginosa_M18
Pseudomonas_aeruginosa_NCMG1179
Pseudomonas_aeruginosa_VRFPA02
Inquilinus_limosus
Donghicola
Tropicibacter_naphthalenivorans
Roseivivax_isoporae
Bauldia_litoralis
Aurantimonas_manganoxydans_SI85-9A1
Aurantimonas_sp_22II-16-19i
Phyllobacterium_sp_YR531
Mesorhizobium_erdmanii
Mesorhizobium_ciceri
Mesorhizobium_loti
Mesorhizobium_sp_Root695
Mesorhizobium_sp_L2C084A000
Mesorhizobium_sp_LNHC252B00
Mesorhizobium_sp_WSM1293
Mesorhizobium_sp_Root102
Mesorhizobium_sp_ICMP_18942
Mesorhizobium_sp_LNJC394B00
Mesorhizobium_sp_URHC0008
Rhizobium_sp_Root483D2
Ensifer_adhaerens
Ensifer_shofinae
Ensifer_sp_OV372
Ensifer_sp_ZNC0028
Sinorhizobium_americanum
Sinorhizobium_fredii
Sinorhizobium_sp_LM21
Sinorhizobium_sp_GW3
Sinorhizobium_sp_GL2
Sinorhizobium_sp_Sb3
Sinorhizobium_sp_GL28
Sinorhizobium_sp_NFACC03
Sinorhizobium_sp_BJ1

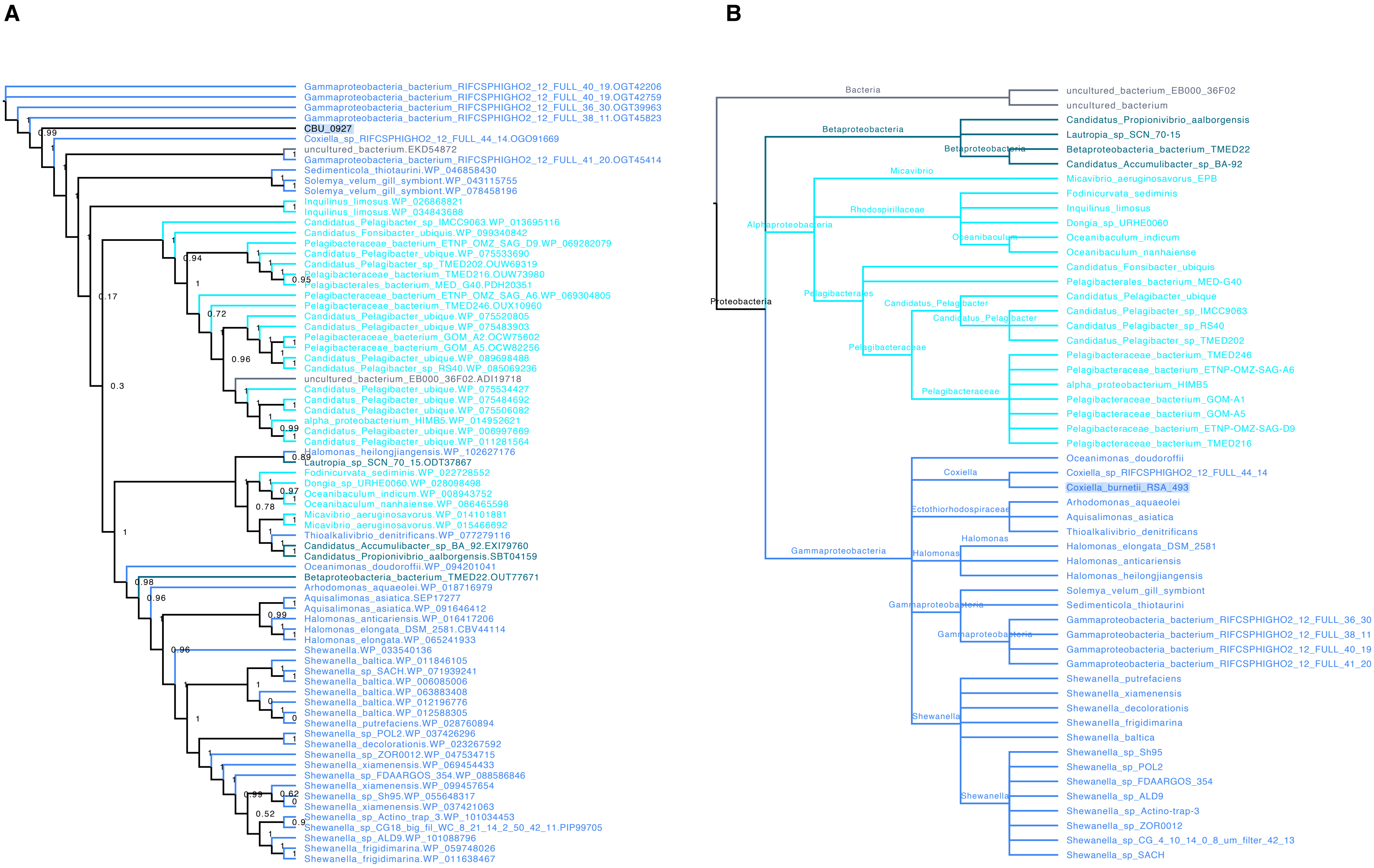
A
Gammaproteobacteria_bacterium_RIFCSPHIGHO2_12_FULL_40_19.OGT42206
Gammaproteobacteria_bacterium_RIFCSPHIGHO2_12_FULL_40_19.OGT42759
Gammaproteobacteria_bacterium_RIFCSPHIGHO2_12_FULL_36_30.OGT39963
Gammaproteobacteria_bacterium_RIFCSPHIGHO2_12_FULL_38_11.OGT45823
CBU_0927
Coxiella_sp_RIFCSPHIGHO2_12_FULL_44_14.OGO91669
uncultured_bacterium.EKD54872
Gammaproteobacteria_bacterium_RIFCSPHIGHO2_12_FULL_41_20.OGT45414
Sedimenticola_thiotaurini.WP_046858430
Solemya_velum_gill_symbiont.WP_043115755
Solemya_velum_gill_symbiont.WP_078458196
Inquilinus_limosus.WP_026868821
Inquilinus_limosus.WP_034843688
Candidatus_Pelagibacter_sp_IMCC9063.WP_013695116
Candidatus_Fonsibacter_ubiquis.WP_099340842
Pelagibacteraceae_bacterium_ETNP_OMZ_SAG_D9.WP_069282079
Candidatus_Pelagibacter_ubique.WP_075533690
Candidatus_Pelagibacter_sp_TMED202.OUW69319
Pelagibacteraceae_bacterium_TMED216.OUW73980
Pelagibacterales_bacterium_MED_G40.PDH20351
Pelagibacteraceae_bacterium_ETNP_OMZ_SAG_A6.WP_069304805
Pelagibacteraceae_bacterium_TMED246.OUX10960
Candidatus_Pelagibacter_ubique.WP_075520805
Candidatus_Pelagibacter_ubique.WP_075483903
Pelagibacteraceae_bacterium_GOM_A2.OCW75602
Pelagibacteraceae_bacterium_GOM_A5.OCW82256
Candidatus_Pelagibacter_ubique.WP_089698488
Candidatus_Pelagibacter_sp_RS40.WP_085069236
uncultured_bacterium_EB000_36F02.ADI19718
Candidatus_Pelagibacter_ubique.WP_075534427
Candidatus_Pelagibacter_ubique.WP_075484692
Candidatus_Pelagibacter_ubique.WP_075506082
alpha_proteobacterium_HIMB5.WP_014952621
Candidatus_Pelagibacter_ubique.WP_006997669
Candidatus_Pelagibacter_ubique.WP_011281564
Halomonas_heilongjiangensis.WP_102627176
Lautropia_sp_SCN_70_15.ODT37867
Fodinicurvata_sediminis.WP_022728552
Dongia_sp_URHE0060.WP_028098498
Oceanibaculum_indicum.WP_008943752
Oceanibaculum_nanhaiense.WP_086465598
Micavibrio_aeruginosavorus.WP_014101881
Micavibrio_aeruginosavorus.WP_015466692
Thioalkalivibrio_denitrificans.WP_077279116
Candidatus_Accumulibacter_sp_BA_92.EXI79760
Candidatus_Propionivibrio_aalborgensis.SBT04159
Oceanimonas_doudoroffii.WP_094201041
Betaproteobacteria_bacterium_TMED22.OUT77671
Arhodomonas_aquaeolei.WP_018716979
Aquisalimonas_asiatica.SEP17277
Aquisalimonas_asiatica.WP_091646412
Halomonas_anticariensis.WP_016417206
Halomonas_elongata_DSM_2581.CBV44114
Halomonas_elongata.WP_065241933
Shewanella.WP_033540136
Shewanella_baltica.WP_011846105
Shewanella_sp_SACH.WP_071939241
Shewanella_baltica.WP_006085006
Shewanella_baltica.WP_063883408
Shewanella_baltica.WP_012196776
Shewanella_baltica.WP_012588305
Shewanella_putrefaciens.WP_028760894
Shewanella_sp_POL2.WP_037426296
Shewanella_decolorationis.WP_023267592
Shewanella_sp_ZOR0012.WP_047534715
Shewanella_xiamenensis.WP_069454433
Shewanella_sp_FDAARGOS_354.WP_088586846
Shewanella_xiamenensis.WP_099457654
Shewanella_sp_Sh95.WP_055648317
Shewanella_xiamenensis.WP_037421063
Shewanella_sp_Actino_trap_3.WP_101034453
Shewanella_sp_CG18_big_fil_WC_8_21_14_2_50_42_11.PIP99705
Shewanella_sp_ALD9.WP_101088796
Shewanella_frigidimarina.WP_059748026
Shewanella_frigidimarina.WP_011638467
B
Bacteria
uncultured_bacterium_EB000_36F02
uncultured_bacterium
Betaproteobacteria
Candidatus_Propionivibrio_aalborgensis
Lautropia_sp_SCN_70-15
Betaproteobacteria
Betaproteobacteria_bacterium_TMED22
Candidatus_Accumulibacter_sp_BA-92
Micavibrio
Micavibrio_aeruginosavorus_EPB
Fodinicurvata_sediminis
Rhodospirillaceae
Inquilinus_limosus
Alphaproteobacteria
Dongia_sp_URHE0060
Oceanibaculum
Oceanibaculum_indicum
Oceanibaculum_nanhaiense
Candidatus_Fonsibacter_ubiquis
Pelagibacterales_bacterium_MED-G40
Pelagibacterales
Candidatus_Pelagibacter_ubique
Proteobacteria
Candidatus_Pelagibacter
Candidatus_Pelagibacter_sp_IMCC9063
Candidatus_Pelagibacter
Candidatus_Pelagibacter_sp_RS40
Candidatus_Pelagibacter_sp_TMED202
Pelagibacteraceae
Pelagibacteraceae_bacterium_TMED246
Pelagibacteraceae_bacterium_ETNP-OMZ-SAG-A6
alpha_proteobacterium_HIMB5
Pelagibacteraceae
Pelagibacteraceae_bacterium_GOM-A1
Pelagibacteraceae_bacterium_GOM-A5
Pelagibacteraceae_bacterium_ETNP-OMZ-SAG-D9
Pelagibacteraceae_bacterium_TMED216
Oceanimonas_doudoroffii
Coxiella
Coxiella_sp_RIFCSPHIGHO2_12_FULL_44_14
Coxiella_burnetii_RSA_493
Arhodomonas_aquaeolei
Ectothiorhodospiraceae
Aquisalimonas_asiatica
Thioalkalivibrio_denitrificans
Halomonas
Halomonas_elongata_DSM_2581
Halomonas
Halomonas_anticariensis
Gammaproteobacteria
Halomonas_heilongjiangensis
Solemya_velum_gill_symbiont
Gammaproteobacteria
Sedimenticola_thiotaurini
Gammaproteobacteria_bacterium_RIFCSPHIGHO2_12_FULL_36_30
Gammaproteobacteria
Gammaproteobacteria_bacterium_RIFCSPHIGHO2_12_FULL_38_11
Gammaproteobacteria_bacterium_RIFCSPHIGHO2_12_FULL_40_19
Gammaproteobacteria_bacterium_RIFCSPHIGHO2_12_FULL_41_20
Shewanella_putrefaciens
Shewanella_xiamenensis
Shewanella_decolorationis
Shewanella
Shewanella_frigidimarina
Shewanella_baltica
Shewanella_sp_Sh95
Shewanella_sp_POL2
Shewanella_sp_FDAARGOS_354
Shewanella
Shewanella_sp_ALD9
Shewanella_sp_Actino-trap-3
Shewanella_sp_ZOR0012
Shewanella_sp_CG_4_10_14_0_8_um_filter_42_13
Shewanella_sp_SACH

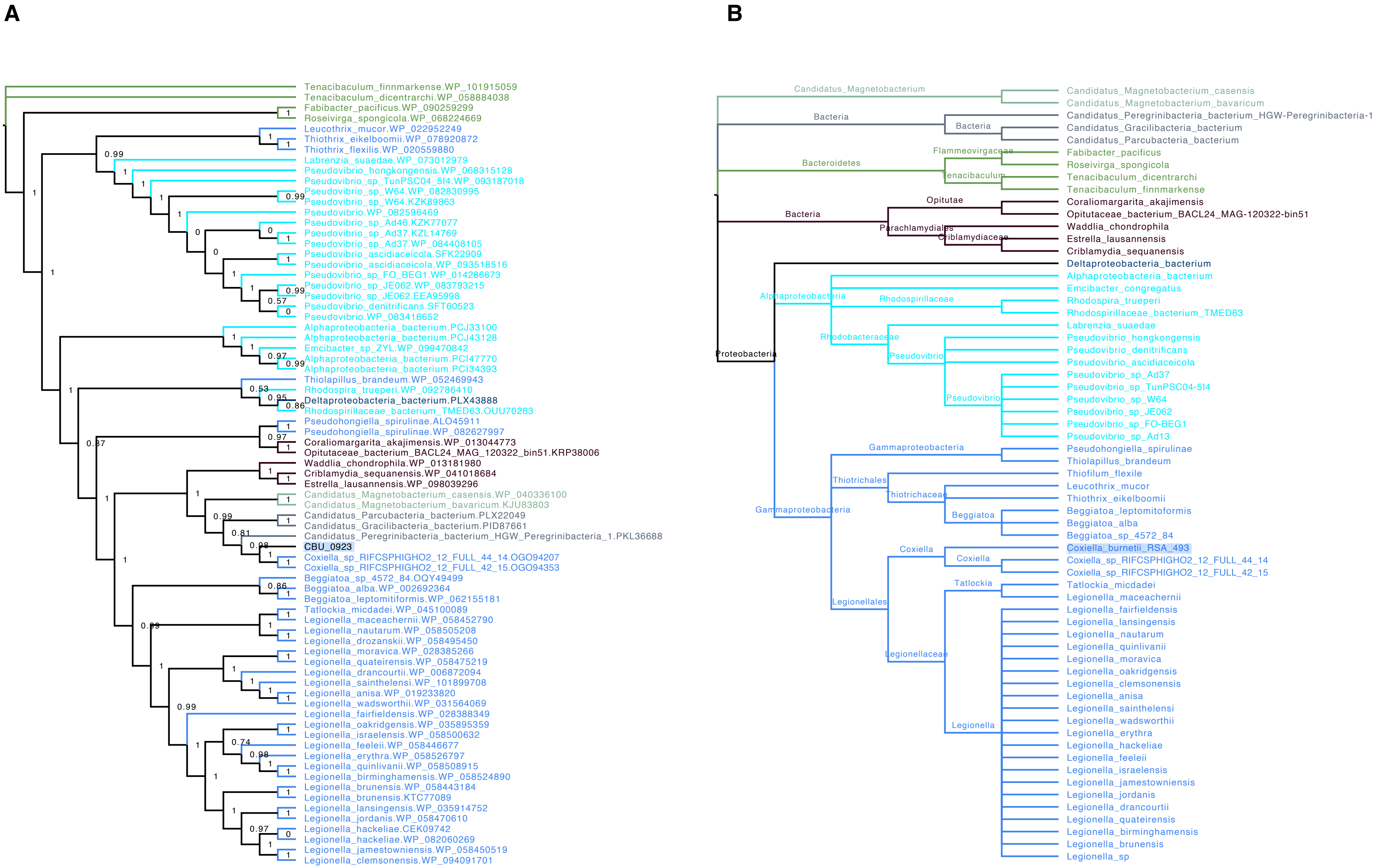
A
Tenacibaculum_finnmarkense.WP_101915059
Tenacibaculum_dicentrarchi.WP_058884038
Fabibacter_pacificus.WP_090259299
Roseivirga_spongicola.WP_068224669
Leucothrix_mucor.WP_022952249
Thiothrix_eikelboomii.WP_078920872
Thiothrix_flexilis.WP_020559880
Labrenzia_suaedae.WP_073012979
Pseudovibrio_hongkongensis.WP_068315128
Pseudovibrio_sp_TunPSC04_5I4.WP_093187018
Pseudovibrio_sp_W64.WP_082830995
Pseudovibrio_sp_W64.KZK89863
Pseudovibrio.WP_082596469
Pseudovibrio_sp_Ad46.KZK77077
Pseudovibrio_sp_Ad37.KZL14769
Pseudovibrio_sp_Ad37.WP_084408105
Pseudovibrio_ascidiaceicola.SFK22909
Pseudovibrio_ascidiaceicola.WP_093518516
Pseudovibrio_sp_FO_BEG1.WP_014286673
Pseudovibrio_sp_JE062.WP_083793215
Pseudovibrio_sp_JE062.EEA95998
Pseudovibrio_denitrificans.SFT60523
Pseudovibrio.WP_083416652
Alphaproteobacteria_bacterium.PCJ33100
Alphaproteobacteria_bacterium.PCJ43128
Emcibacter_sp_ZYL.WP_099470842
Alphaproteobacteria_bacterium.PCI47770
Alphaproteobacteria_bacterium.PCI34393
Thiolapillus_brandeum.WP_052469943
Rhodospira_trueperi.WP_092786410
Deltaproteobacteria_bacterium.PLX43888
Rhodospirillaceae_bacterium_TMED63.OUU70263
Pseudohongiella_spirulinae.ALO45911
Pseudohongiella_spirulinae.WP_082627997
Coraliomargarita_akajimensis.WP_013044773
Opitutaceae_bacterium_BACL24_MAG_120322_bin51.KRP38006
Waddlia_chondrophila.WP_013181980
Criblamydia_sequanensis.WP_041018684
Estrella_lausannensis.WP_098039296
Candidatus_Magnetobacterium_casensis.WP_040336100
Candidatus_Magnetobacterium_bavaricum.KJU83803
Candidatus_Parcubacteria_bacterium.PLX22049
Candidatus_Gracilibacteria_bacterium.PID87661
Candidatus_Peregrinibacteria_bacterium_HGW_Peregrinibacteria_1.PKL36688
CBU_0923
Coxiella_sp_RIFCSPHIGHO2_12_FULL_44_14.OGO94207
Coxiella_sp_RIFCSPHIGHO2_12_FULL_42_15.OGO94353
Beggiatoa_sp_4572_84.OQY49499
Beggiatoa_alba.WP_002692364
Beggiatoa_leptomitiformis.WP_062155181
Tatlockia_micdadei.WP_045100089
Legionella_maceachernii.WP_058452790
Legionella_nautarum.WP_058505208
Legionella_drozanskii.WP_058495450
Legionella_moravica.WP_028385266
Legionella_quateirensis.WP_058475219
Legionella_drancourtii.WP_006872094
Legionella_sainthelensi.WP_101899708
Legionella_anisa.WP_019233820
Legionella_wadsworthii.WP_031564069
Legionella_fairfieldensis.WP_028388349
Legionella_oakridgensis.WP_035895359
Legionella_israelensis.WP_058500632
Legionella_feeleii.WP_058446677
Legionella_erythra.WP_058526797
Legionella_quinlivanii.WP_058508915
Legionella_birminghamensis.WP_058524890
Legionella_brunensis.WP_058443184
Legionella_brunensis.KTC77089
Legionella_lansingensis.WP_035914752
Legionella_jordanis.WP_058470610
Legionella_hackeliae.CEK09742
Legionella_hackeliae.WP_082060269
Legionella_jamestowniensis.WP_058450519
Legionella_clemsonensis.WP_094091701
B
Candidatus_Magnetobacterium
Bacteria
Bacteria
Flammeovirgaceae
Bacteroidetes
Tenacibaculum
Opitutae
Bacteria
Parachlamydiales
Criblamydiaceae
Proteobacteria
Alphaproteobacteria
Rhodospirillaceae
Rhodobacteraceae
Pseudovibrio
Pseudovibrio
Gammaproteobacteria
Thiotrichales
Thiotrichaceae
Gammaproteobacteria
Beggiatoa
Coxiella
Coxiella
Tatlockia
Legionellales
Legionellaceae
Legionella
Candidatus_Magnetobacterium_casensis
Candidatus_Magnetobacterium_bavaricum
Candidatus_Peregrinibacteria_bacterium_HGW-Peregrinibacteria-1
Candidatus_Gracilibacteria_bacterium
Candidatus_Parcubacteria_bacterium
Fabibacter_pacificus
Roseivirga_spongicola
Tenacibaculum_dicentrarchi
Tenacibaculum_finnmarkense
Coraliomargarita_akajimensis
Opitutaceae_bacterium_BACL24_MAG-120322-bin51
Waddlia_chondrophila
Estrella_lausannensis
Criblamydia_sequanensis
Deltaproteobacteria_bacterium
Alphaproteobacteria_bacterium
Emcibacter_congregatus
Rhodospira_trueperi
Rhodospirillaceae_bacterium_TMED63
Labrenzia_suaedae
Pseudovibrio_hongkongensis
Pseudovibrio_denitrificans
Pseudovibrio_ascidiaceicola
Pseudovibrio_sp_Ad37
Pseudovibrio_sp_TunPSC04-5I4
Pseudovibrio_sp_W64
Pseudovibrio_sp_JE062
Pseudovibrio_sp_FO-BEG1
Pseudovibrio_sp_Ad13
Pseudohongiella_spirulinae
Thiolapillus_brandeum
Thiofilum_flexile
Leucothrix_mucor
Thiothrix_eikelboomii
Beggiatoa_leptomitoformis
Beggiatoa_alba
Beggiatoa_sp_4572_84
Coxiella_burnetii_RSA_493
Coxiella_sp_RIFCSPHIGHO2_12_FULL_44_14
Coxiella_sp_RIFCSPHIGHO2_12_FULL_42_15
Tatlockia_micdadei
Legionella_maceachernii
Legionella_fairfieldensis
Legionella_lansingensis
Legionella_nautarum
Legionella_quinlivanii
Legionella_moravica
Legionella_oakridgensis
Legionella_clemsonensis
Legionella_anisa
Legionella_sainthelensi
Legionella_wadsworthii
Legionella_erythra
Legionella_hackeliae
Legionella_feeleii
Legionella_israelensis
Legionella_jamestowniensis
Legionella_jordanis
Legionella_drancourtii
Legionella_quateirensis
Legionella_birminghamensis
Legionella_brunensis
Legionella_sp

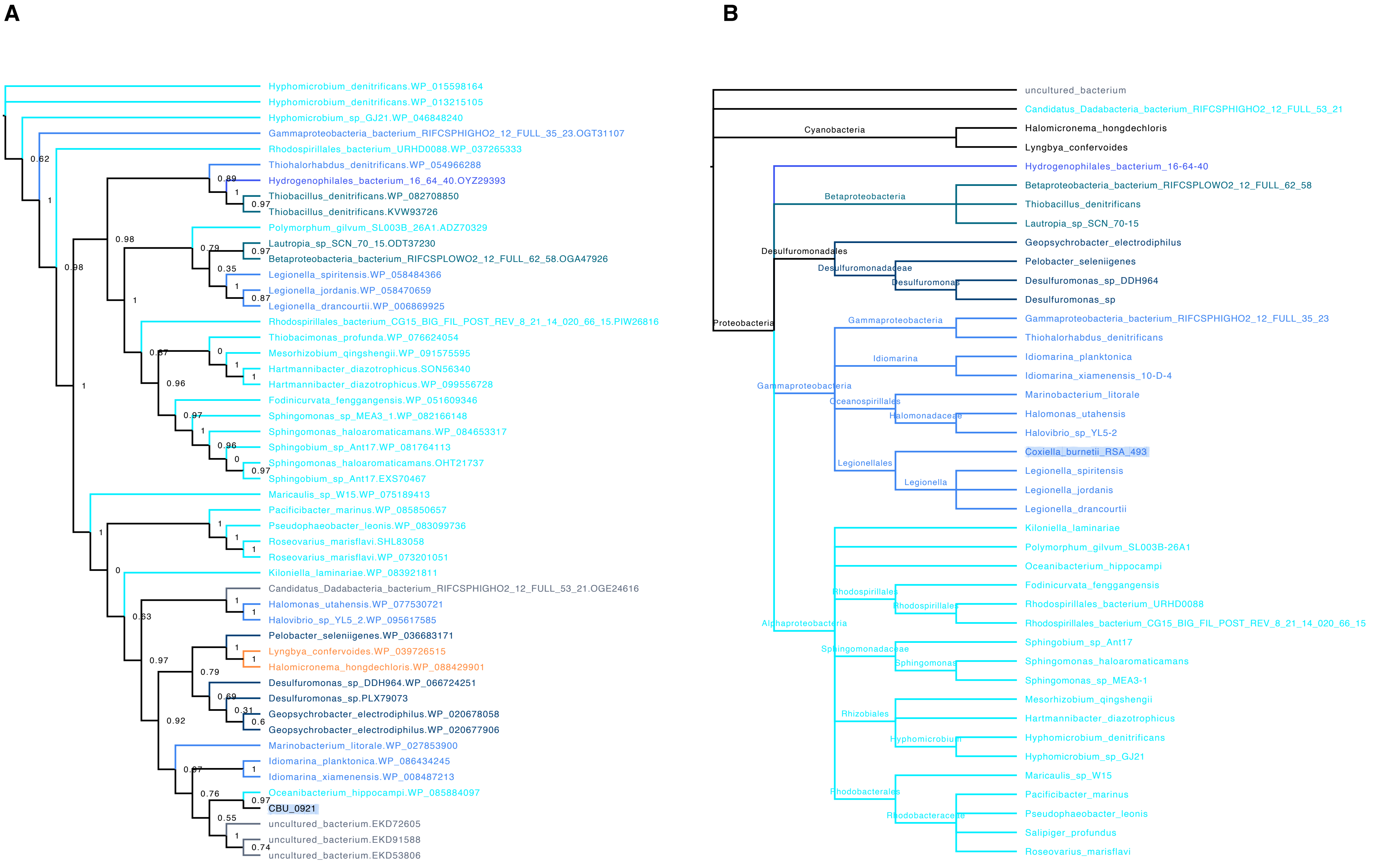
A
Hyphomicrobium_denitrificans.WP_015598164
Hyphomicrobium_denitrificans.WP_013215105
Hyphomicrobium_sp_GJ21.WP_046848240
Gammaproteobacteria_bacterium_RIFCSPHIGHO2_12_FULL_35_23.OGT31107
Rhodospirillales_bacterium_URHD0088.WP_037265333
Thiohalorhabdus_denitrificans.WP_054966288
Hydrogenophilales_bacterium_16_64_40.OYZ29393
Thiobacillus_denitrificans.WP_082708850
Thiobacillus_denitrificans.KVW93726
Polymorphum_gilvum_SL003B_26A1.ADZ70329
Lautropia_sp_SCN_70_15.ODT37230
Betaproteobacteria_bacterium_RIFCSPLOWO2_12_FULL_62_58.OGA47926
Legionella_spiritensis.WP_058484366
Legionella_jordanis.WP_058470659
Legionella_drancourtii.WP_006869925
Rhodospirillales_bacterium_CG15_BIG_FIL_POST_REV_8_21_14_020_66_15.PIW26816
Thiobacimonas_profunda.WP_076624054
Mesorhizobium_qingshengii.WP_091575595
Hartmannibacter_diazotrophicus.SON56340
Hartmannibacter_diazotrophicus.WP_099556728
Fodinicurvata_fenggangensis.WP_051609346
Sphingomonas_sp_MEA3_1.WP_082166148
Sphingomonas_haloaromaticamans.WP_084653317
Sphingobium_sp_Ant17.WP_081764113
Sphingomonas_haloaromaticamans.OHT21737
Sphingobium_sp_Ant17.EXS70467
Maricaulis_sp_W15.WP_075189413
Pacificibacter_marinus.WP_085850657
Pseudophaeobacter_leonis.WP_083099736
Roseovarius_marisflavi.SHL83058
Roseovarius_marisflavi.WP_073201051
Kiloniella_laminariae.WP_083921811
Candidatus_Dadabacteria_bacterium_RIFCSPHIGHO2_12_FULL_53_21.OGE24616
Halomonas_utahensis.WP_077530721
Halovibrio_sp_YL5_2.WP_095617585
Pelobacter_seleniigenes.WP_036683171
Lyngbya_confervoides.WP_039726515
Halomicronema_hongdechloris.WP_088429901
Desulfuromonas_sp_DDH964.WP_066724251
Desulfuromonas_sp.PLX79073
Geopsychrobacter_electrodiphilus.WP_020678058
Geopsychrobacter_electrodiphilus.WP_020677906
Marinobacterium_litorale.WP_027853900
Idiomarina_planktonica.WP_086434245
Idiomarina_xiamenensis.WP_008487213
Oceanibacterium_hippocampi.WP_085884097
CBU_0921
uncultured_bacterium.EKD72605
uncultured_bacterium.EKD91588
uncultured_bacterium.EKD53806
B
uncultured_bacterium
Candidatus_Dadabacteria_bacterium_RIFCSPHIGHO2_12_FULL_53_21
Cyanobacteria
Halomicronema_hongdechloris
Lyngbya_confervoides
Hydrogenophilales_bacterium_16-64-40
Betaproteobacteria
Betaproteobacteria_bacterium_RIFCSPLOWO2_12_FULL_62_58
Thiobacillus_denitrificans
Lautropia_sp_SCN_70-15
Desulfuromonadales
Desulfuromonadaceae
Desulfuromonas
Geopsychrobacter_electrodiphilus
Pelobacter_seleniigenes
Desulfuromonas_sp_DDH964
Desulfuromonas_sp
Proteobacteria
Gammaproteobacteria
Gammaproteobacteria_bacterium_RIFCSPHIGHO2_12_FULL_35_23
Thiohalorhabdus_denitrificans
Idiomarina
Idiomarina_planktonica
Idiomarina_xiamenensis_10-D-4
Oceanospirillales
Halomonadaceae
Marinobacterium_litorale
Halomonas_utahensis
Halovibrio_sp_YL5-2
Legionellales
Legionella
Coxiella_burnetii_RSA_493
Legionella_spiritensis
Legionella_jordanis
Legionella_drancourtii
Alphaproteobacteria
Kiloniella_laminariae
Polymorphum_gilvum_SL003B-26A1
Oceanibacterium_hippocampi
Rhodospirillales
Fodinicurvata_fenggangensis
Rhodospirillales_bacterium_URHD0088
Rhodospirillales_bacterium_CG15_BIG_FIL_POST_REV_8_21_14_020_66_15
Sphingomonadaceae
Sphingomonas
Sphingobium_sp_Ant17
Sphingomonas_haloaromaticamans
Sphingomonas_sp_MEA3-1
Rhizobiales
Hyphomicrobium
Mesorhizobium_qingshengii
Hartmannibacter_diazotrophicus
Hyphomicrobium_denitrificans
Hyphomicrobium_sp_GJ21
Rhodobacterales
Rhodobacteraceae
Maricaulis_sp_W15
Pacificibacter_marinus
Pseudophaeobacter_leonis
Salipiger_profundus
Roseovarius_marisflavi

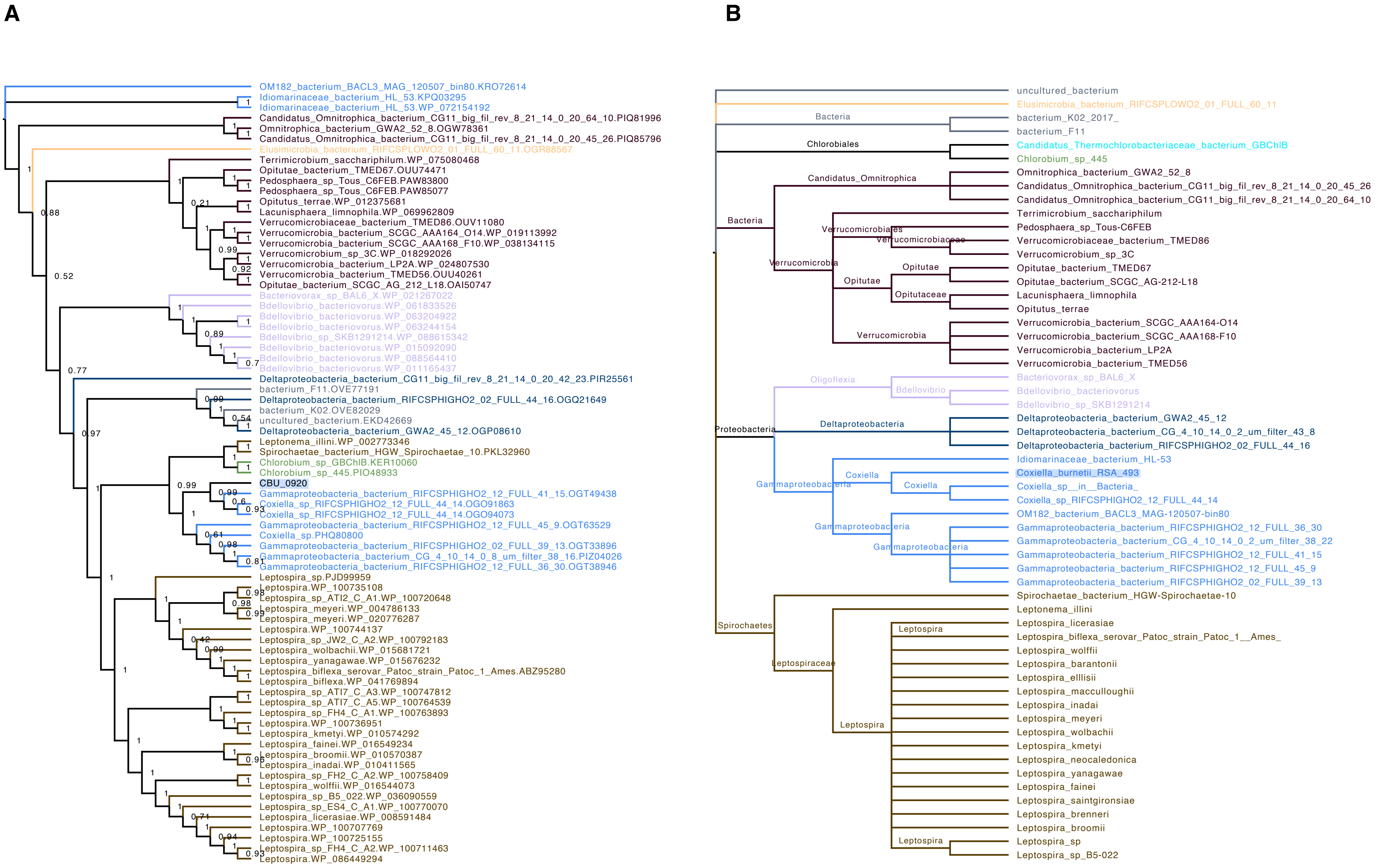
A
OM182_bacterium_BACL3_MAG_120507_bin80.KRO72614
Idiomarinaceae_bacterium_HL_53.KPQ03295
Idiomarinaceae_bacterium_HL_53.WP_072154192
Candidatus_Omnitrophica_bacterium_CG11_big_fil_rev_8_21_14_0_20_64_10.PIQ81996
Omnitrophica_bacterium_GWA2_52_8.OGW78361
Candidatus_Omnitrophica_bacterium_CG11_big_fil_rev_8_21_14_0_20_45_26.PIQ85796
Elusimicrobia_bacterium_RIFCSPLOWO2_01_FULL_60_11.OGR88567
Terrimicrobium_sacchariphilum.WP_075080468
Opitutae_bacterium_TMED67.OUU74471
Pedosphaera_sp_Tous_C6FEB.PAW83800
Pedosphaera_sp_Tous_C6FEB.PAW85077
Opitutus_terrae.WP_012375681
Lacunisphaera_limnophila.WP_069962809
Verrucomicrobiaceae_bacterium_TMED86.OUV11080
Verrucomicrobia_bacterium_SCGC_AAA164_O14.WP_019113992
Verrucomicrobia_bacterium_SCGC_AAA168_F10.WP_038134115
Verrucomicrobium_sp_3C.WP_018292026
Verrucomicrobia_bacterium_LP2A.WP_024807530
Verrucomicrobia_bacterium_TMED56.OUU40261
Opitutae_bacterium_SCGC_AG_212_L18.OAI50747
Bacteriovorax_sp_BAL6_X.WP_021267022
Bdellovibrio_bacteriovorus.WP_061833526
Bdellovibrio_bacteriovorus.WP_063204922
Bdellovibrio_bacteriovorus.WP_063244154
Bdellovibrio_sp_SKB1291214.WP_088615342
Bdellovibrio_bacteriovorus.WP_015092090
Bdellovibrio_bacteriovorus.WP_088564410
Bdellovibrio_bacteriovorus.WP_011165437
Deltaproteobacteria_bacterium_CG11_big_fil_rev_8_21_14_0_20_42_23.PIR25561
bacterium_F11.OVE77191
Deltaproteobacteria_bacterium_RIFCSPHIGHO2_02_FULL_44_16.OGQ21649
bacterium_K02.OVE82029
uncultured_bacterium.EKD42669
Deltaproteobacteria_bacterium_GWA2_45_12.OGP08610
Leptonema_illini.WP_002773346
Spirochaetae_bacterium_HGW_Spirochaetae_10.PKL32960
Chlorobium_sp_GBChlB.KER10060
Chlorobium_sp_445.PIO48933
CBU_0920
Gammaproteobacteria_bacterium_RIFCSPHIGHO2_12_FULL_41_15.OGT49438
Coxiella_sp_RIFCSPHIGHO2_12_FULL_44_14.OGO91863
Coxiella_sp_RIFCSPHIGHO2_12_FULL_44_14.OGO94073
Gammaproteobacteria_bacterium_RIFCSPHIGHO2_12_FULL_45_9.OGT63529
Coxiella_sp.PHQ80800
Gammaproteobacteria_bacterium_RIFCSPHIGHO2_02_FULL_39_13.OGT33896
Gammaproteobacteria_bacterium_CG_4_10_14_0_8_um_filter_38_16.PIZ04026
Gammaproteobacteria_bacterium_RIFCSPHIGHO2_12_FULL_36_30.OGT38946
Leptospira_sp.PJD99959
Leptospira.WP_100735108
Leptospira_sp_ATI2_C_A1.WP_100720648
Leptospira_meyeri.WP_004786133
Leptospira_meyeri.WP_020776287
Leptospira.WP_100744137
Leptospira_sp_JW2_C_A2.WP_100792183
Leptospira_wolbachii.WP_015681721
Leptospira_yanagawae.WP_015676232
Leptospira_biflexa_serovar_Patoc_strain_Patoc_1_Ames.ABZ95280
Leptospira_biflexa.WP_041769894
Leptospira_sp_ATI7_C_A3.WP_100747812
Leptospira_sp_ATI7_C_A5.WP_100764539
Leptospira_sp_FH4_C_A1.WP_100763893
Leptospira.WP_100736951
Leptospira_kmetyi.WP_010574292
Leptospira_fainei.WP_016549234
Leptospira_broomii.WP_010570387
Leptospira_inadai.WP_010411565
Leptospira_sp_FH2_C_A2.WP_100758409
Leptospira_wolffii.WP_016544073
Leptospira_sp_B5_022.WP_036090559
Leptospira_sp_ES4_C_A1.WP_100770070
Leptospira_licerasiae.WP_008591484
Leptospira.WP_100707769
Leptospira.WP_100725155
Leptospira_sp_FH4_C_A2.WP_100711463
Leptospira.WP_086449294
B
uncultured_bacterium
Elusimicrobia_bacterium_RIFCSPLOWO2_01_FULL_60_11
Bacteria
bacterium_K02_2017_
bacterium_F11
Chlorobiales
Candidatus_Thermochlorobacteriaceae_bacterium_GBChlB
Chlorobium_sp_445
Candidatus_Omnitrophica
Omnitrophica_bacterium_GWA2_52_8
Candidatus_Omnitrophica_bacterium_CG11_big_fil_rev_8_21_14_0_20_45_26
Candidatus_Omnitrophica_bacterium_CG11_big_fil_rev_8_21_14_0_20_64_10
Bacteria
Terrimicrobium_sacchariphilum
Verrucomicrobiales
Pedosphaera_sp_Tous-C6FEB
Verrucomicrobiaceae
Verrucomicrobiaceae_bacterium_TMED86
Verrucomicrobium_sp_3C
Verrucomicrobia
Opitutae
Opitutae_bacterium_TMED67
Opitutae
Opitutae_bacterium_SCGC_AG-212-L18
Opitutaceae
Lacunisphaera_limnophila
Opitutus_terrae
Verrucomicrobia_bacterium_SCGC_AAA164-O14
Verrucomicrobia
Verrucomicrobia_bacterium_SCGC_AAA168-F10
Verrucomicrobia_bacterium_LP2A
Verrucomicrobia_bacterium_TMED56
Oligoflexia
Bacteriovorax_sp_BAL6_X
Bdellovibrio
Bdellovibrio_bacteriovorus
Bdellovibrio_sp_SKB1291214
Deltaproteobacteria
Deltaproteobacteria_bacterium_GWA2_45_12
Proteobacteria
Deltaproteobacteria_bacterium_CG_4_10_14_0_2_um_filter_43_8
Deltaproteobacteria_bacterium_RIFCSPHIGHO2_02_FULL_44_16
Idiomarinaceae_bacterium_HL-53
Coxiella
Coxiella_burnetii_RSA_493
Gammaproteobacteria
Coxiella
Coxiella_sp__in__Bacteria_
Coxiella_sp_RIFCSPHIGHO2_12_FULL_44_14
OM182_bacterium_BACL3_MAG-120507-bin80
Gammaproteobacteria
Gammaproteobacteria_bacterium_RIFCSPHIGHO2_12_FULL_36_30
Gammaproteobacteria_bacterium_CG_4_10_14_0_2_um_filter_38_22
Gammaproteobacteria
Gammaproteobacteria_bacterium_RIFCSPHIGHO2_12_FULL_41_15
Gammaproteobacteria_bacterium_RIFCSPHIGHO2_12_FULL_45_9
Gammaproteobacteria_bacterium_RIFCSPHIGHO2_02_FULL_39_13
Spirochaetae_bacterium_HGW-Spirochaetae-10
Leptonema_illini
Spirochaetes
Leptospira
Leptospira_licerasiae
Leptospira_biflexa_serovar_Patoc_strain_Patoc_1__Ames_
Leptospira_wolffii
Leptospiraceae
Leptospira_barantonii
Leptospira_ellisii
Leptospira_macculloughii
Leptospira_inadai
Leptospira_meyeri
Leptospira
Leptospira_wolbachii
Leptospira_kmetyi
Leptospira_neocaledonica
Leptospira_yanagawae
Leptospira_fainei
Leptospira_saintgironsiae
Leptospira_brenneri
Leptospira_broomii
Leptospira
Leptospira_sp
Leptospira_sp_B5-022

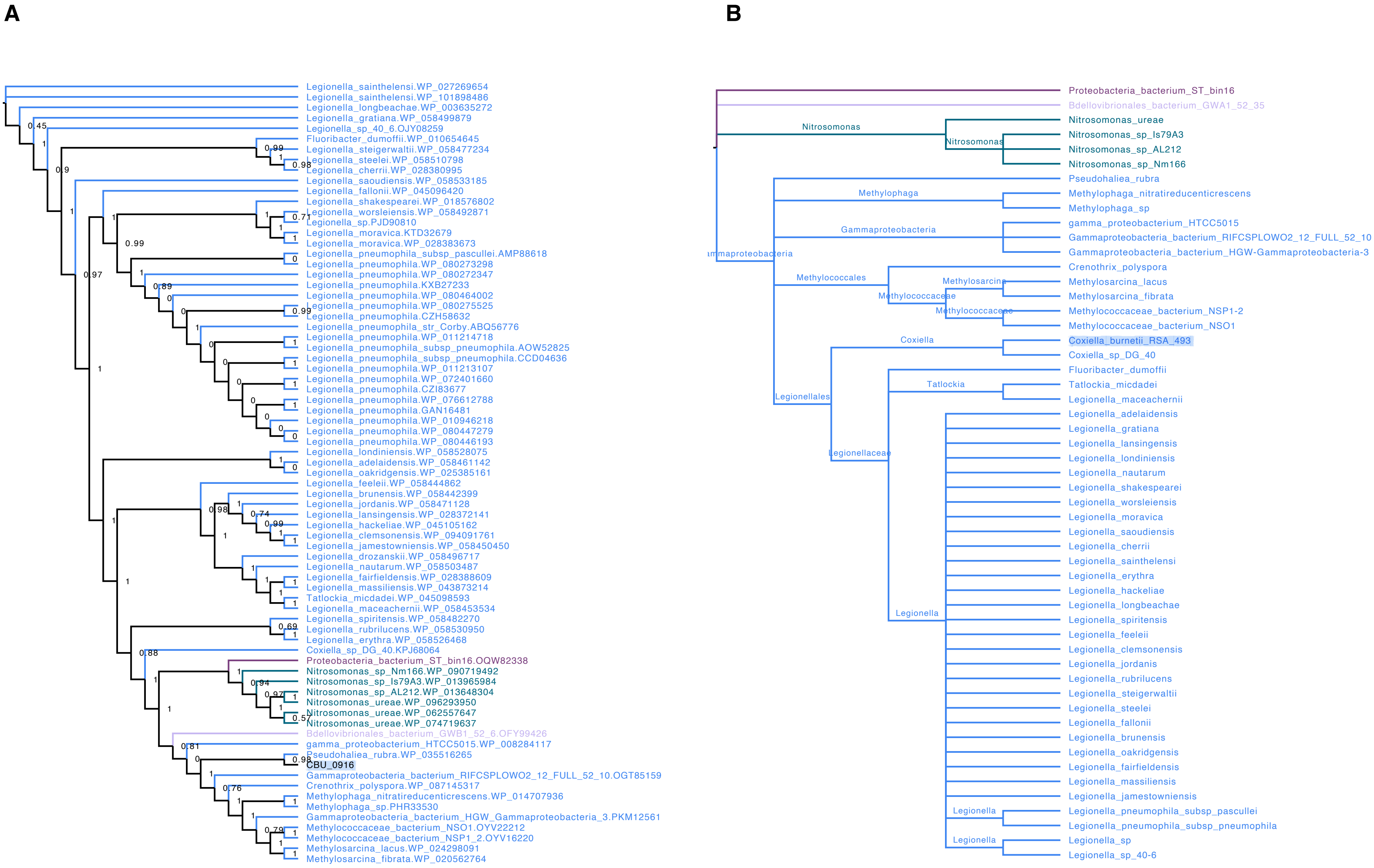
A
Legionella_sainthelensi.WP_027269654
Legionella_sainthelensi.WP_101898486
Legionella_longbeachae.WP_003635272
Legionella_gratiana.WP_058499879
Legionella_sp_40_6.OJY08259
Fluoribacter_dumoffii.WP_010654645
Legionella_steigerwaltii.WP_058477234
Legionella_steelei.WP_058510798
Legionella_cherrii.WP_028380995
Legionella_saoudiensis.WP_058533185
Legionella_fallonii.WP_045096420
Legionella_shakespearei.WP_018576802
Legionella_worsleiensis.WP_058492871
Legionella_sp.PJD90810
Legionella_moravica.KTD32679
Legionella_moravica.WP_028383673
Legionella_pneumophila_subsp_pascullei.AMP88618
Legionella_pneumophila.WP_080273298
Legionella_pneumophila.WP_080272347
Legionella_pneumophila.KXB27233
Legionella_pneumophila.WP_080464002
Legionella_pneumophila.WP_080275525
Legionella_pneumophila.CZH58632
Legionella_pneumophila_str_Corby.ABQ56776
Legionella_pneumophila.WP_011214718
Legionella_pneumophila_subsp_pneumophila.AOW52825
Legionella_pneumophila_subsp_pneumophila.CCD04636
Legionella_pneumophila.WP_011213107
Legionella_pneumophila.WP_072401660
Legionella_pneumophila.CZI83677
Legionella_pneumophila.WP_076612788
Legionella_pneumophila.GAN16481
Legionella_pneumophila.WP_010946218
Legionella_pneumophila.WP_080447279
Legionella_pneumophila.WP_080446193
Legionella_londiniensis.WP_058528075
Legionella_adelaidensis.WP_058461142
Legionella_oakridgensis.WP_025385161
Legionella_feeleii.WP_058444862
Legionella_brunensis.WP_058442399
Legionella_jordanis.WP_058471128
Legionella_lansingensis.WP_028372141
Legionella_hackeliae.WP_045105162
Legionella_clemsonensis.WP_094091761
Legionella_jamestowniensis.WP_058450450
Legionella_drozanskii.WP_058496717
Legionella_nautarum.WP_058503487
Legionella_fairfieldensis.WP_028388609
Legionella_massiliensis.WP_043873214
Tatlockia_micdadei.WP_045098593
Legionella_maceachernii.WP_058453534
Legionella_spiritensis.WP_058482270
Legionella_rubrilucens.WP_058530950
Legionella_erythra.WP_058526468
Coxiella_sp_DG_40.KPJ68064
Proteobacteria_bacterium_ST_bin16.OQW82338
Nitrosomonas_sp_Nm166.WP_090719492
Nitrosomonas_sp_Is79A3.WP_013965984
Nitrosomonas_sp_AL212.WP_013648304
Nitrosomonas_ureae.WP_096293950
Nitrosomonas_ureae.WP_062557647
Nitrosomonas_ureae.WP_074719637
Bdellovibrionales_bacterium_GWB1_52_6.OFY99426
gamma_proteobacterium_HTCC5015.WP_008284117
Pseudohaliea_rubra.WP_035516265
CBU_0916
Gammaproteobacteria_bacterium_RIFCSPLOWO2_12_FULL_52_10.OGT85159
Crenothrix_polyspora.WP_087145317
Methylophaga_nitratireducenticrescens.WP_014707936
Methylophaga_sp.PHR33530
Gammaproteobacteria_bacterium_HGW_Gammaproteobacteria_3.PKM12561
Methylococcaceae_bacterium_NSO1.OYV22212
Methylococcaceae_bacterium_NSP1_2.OYV16220
Methylosarcina_lacus.WP_024298091
Methylosarcina_fibrata.WP_020562764
B
Proteobacteria_bacterium_ST_bin16
Bdellovibrionales_bacterium_GWA1_52_35
Nitrosomonas
Nitrosomonas
Nitrosomonas_ureae
Nitrosomonas_sp_Is79A3
Nitrosomonas_sp_AL212
Nitrosomonas_sp_Nm166
Pseudohaliea_rubra
Methylophaga
Methylophaga_nitratireducenticrescens
Methylophaga_sp
Gammaproteobacteria
gamma_proteobacterium_HTCC5015
Gammaproteobacteria_bacterium_RIFCSPLOWO2_12_FULL_52_10
Gammaproteobacteria_bacterium_HGW-Gammaproteobacteria-3
Methylococcales
Crenothrix_polyspora
Methylosarcina
Methylosarcina_lacus
Methylosarcina_fibrata
Methylococcaceae
Methylococcaceae
Methylococcaceae_bacterium_NSP1-2
Methylococcaceae_bacterium_NSO1
Coxiella
Coxiella_burnetii_RSA_493
Coxiella_sp_DG_40
Fluoribacter_dumoffii
Tatlockia
Tatlockia_micdadei
Legionella_maceachernii
Legionellales
Legionellaceae
Legionella_adelaidensis
Legionella_gratiana
Legionella_lansingensis
Legionella_londiniensis
Legionella_nautarum
Legionella_shakespearei
Legionella_worsleiensis
Legionella_moravica
Legionella_saoudiensis
Legionella_cherrii
Legionella_sainthelensi
Legionella_erythra
Legionella_hackeliae
Legionella_longbeachae
Legionella_spiritensis
Legionella
Legionella_feeleii
Legionella_clemsonensis
Legionella_jordanis
Legionella_rubrilucens
Legionella_steigerwaltii
Legionella_steelei
Legionella_fallonii
Legionella_brunensis
Legionella_oakridgensis
Legionella_fairfieldensis
Legionella_massiliensis
Legionella_jamestowniensis
Legionella
Legionella_pneumophila_subsp_pascullei
Legionella_pneumophila_subsp_pneumophila
Legionella
Legionella_sp
Legionella_sp_40-6

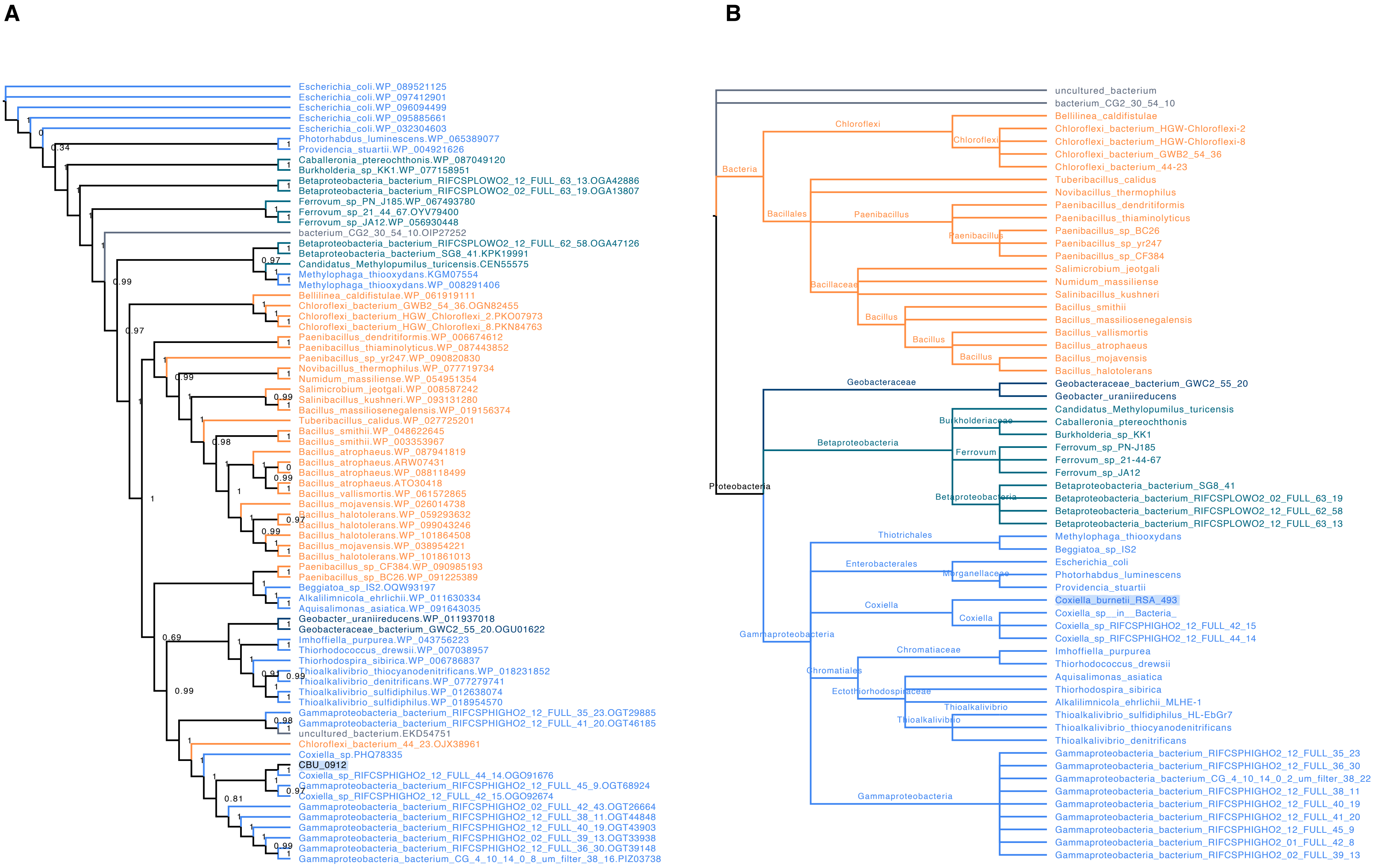
A
Escherichia_coli.WP_089521125
Escherichia_coli.WP_097412901
Escherichia_coli.WP_096094499
Escherichia_coli.WP_095885661
Escherichia_coli.WP_032304603
Photorhabdus_luminescens.WP_065389077
Providencia_stuartii.WP_004921626
Caballeronia_ptereochthonis.WP_087049120
Burkholderia_sp_KK1.WP_077158951
Betaproteobacteria_bacterium_RIFCSPLOWO2_12_FULL_63_13.OGA42886
Betaproteobacteria_bacterium_RIFCSPLOWO2_02_FULL_63_19.OGA13807
Ferrovum_sp_PN_J185.WP_067493780
Ferrovum_sp_21_44_67.OYV79400
Ferrovum_sp_JA12.WP_056930448
bacterium_CG2_30_54_10.OIP27252
Betaproteobacteria_bacterium_RIFCSPLOWO2_12_FULL_62_58.OGA47126
Betaproteobacteria_bacterium_SG8_41.KPK19991
Candidatus_Methylopumilus_turicensis.CEN55575
Methylophaga_thiooxydans.KGM07554
Methylophaga_thiooxydans.WP_008291406
Bellilinea_caldifistulae.WP_061919111
Chloroflexi_bacterium_GWB2_54_36.OGN82455
Chloroflexi_bacterium_HGW_Chloroflexi_2.PKO07973
Chloroflexi_bacterium_HGW_Chloroflexi_8.PKN84763
Paenibacillus_dendritiformis.WP_006674612
Paenibacillus_thiaminolyticus.WP_087443852
Paenibacillus_sp_yr247.WP_090820830
Novibacillus_thermophilus.WP_077719734
Numidum_massiliense.WP_054951354
Salimicrobium_jeotgali.WP_008587242
Salinibacillus_kushneri.WP_093131280
Bacillus_massiliosenegalensis.WP_019156374
Tuberibacillus_calidus.WP_027725201
Bacillus_smithii.WP_048622645
Bacillus_smithii.WP_003353967
Bacillus_atrophaeus.WP_087941819
Bacillus_atrophaeus.ARW07431
Bacillus_atrophaeus.WP_088118499
Bacillus_atrophaeus.ATO30418
Bacillus_vallismortis.WP_061572865
Bacillus_mojavensis.WP_026014738
Bacillus_halotolerans.WP_059293632
Bacillus_halotolerans.WP_099043246
Bacillus_halotolerans.WP_101864508
Bacillus_mojavensis.WP_038954221
Bacillus_halotolerans.WP_101861013
Paenibacillus_sp_CF384.WP_090985193
Paenibacillus_sp_BC26.WP_091225389
Beggiatoa_sp_IS2.OQW93197
Alkalilimnicola_ehrlichii.WP_011630334
Aquisalimonas_asiatica.WP_091643035
Geobacter_uraniireducens.WP_011937018
Geobacteraceae_bacterium_GWC2_55_20.OGU01622
Imhoffiella_purpurea.WP_043756223
Thiorhodococcus_drewsii.WP_007038957
Thiorhodospira_sibirica.WP_006786837
Thioalkalivibrio_thiocyanodenitrificans.WP_018231852
Thioalkalivibrio_denitrificans.WP_077279741
Thioalkalivibrio_sulfidiphilus.WP_012638074
Thioalkalivibrio_sulfidiphilus.WP_018954570
Gammaproteobacteria_bacterium_RIFCSPHIGHO2_12_FULL_35_23.OGT29885
Gammaproteobacteria_bacterium_RIFCSPHIGHO2_12_FULL_41_20.OGT46185
uncultured_bacterium.EKD54751
Chloroflexi_bacterium_44_23.OJX38961
Coxiella_sp.PHQ78335
CBU_0912
Coxiella_sp_RIFCSPHIGHO2_12_FULL_44_14.OGO91676
Gammaproteobacteria_bacterium_RIFCSPHIGHO2_12_FULL_45_9.OGT68924
Coxiella_sp_RIFCSPHIGHO2_12_FULL_42_15.OGO92674
Gammaproteobacteria_bacterium_RIFCSPHIGHO2_02_FULL_42_43.OGT26664
Gammaproteobacteria_bacterium_RIFCSPHIGHO2_12_FULL_38_11.OGT44848
Gammaproteobacteria_bacterium_RIFCSPHIGHO2_12_FULL_40_19.OGT43903
Gammaproteobacteria_bacterium_RIFCSPHIGHO2_02_FULL_39_13.OGT33938
Gammaproteobacteria_bacterium_RIFCSPHIGHO2_12_FULL_36_30.OGT39148
Gammaproteobacteria_bacterium_CG_4_10_14_0_8_um_filter_38_16.PIZ03738
B
uncultured_bacterium
bacterium_CG2_30_54_10
Bacteria
Chloroflexi
Bellilinea_caldifistulae
Chloroflexi_bacterium_HGW-Chloroflexi-2
Chloroflexi_bacterium_HGW-Chloroflexi-8
Chloroflexi_bacterium_GWB2_54_36
Chloroflexi_bacterium_44-23
Bacillales
Tuberibacillus_calidus
Novibacillus_thermophilus
Paenibacillus
Paenibacillus_dendritiformis
Paenibacillus_thiaminolyticus
Paenibacillus_sp_BC26
Paenibacillus_sp_yr247
Paenibacillus_sp_CF384
Bacillaceae
Salimicrobium_jeotgali
Numidum_massiliense
Salinibacillus_kushneri
Bacillus
Bacillus_smithii
Bacillus_massiliosenegalensis
Bacillus_vallismortis
Bacillus_atrophaeus
Bacillus_mojavensis
Bacillus_halotolerans
Proteobacteria
Geobacteraceae
Geobacteraceae_bacterium_GWC2_55_20
Geobacter_uraniireducens
Betaproteobacteria
Burkholderiaceae
Candidatus_Methylopumilus_turicensis
Caballeronia_ptereochthonis
Burkholderia_sp_KK1
Ferrovum
Ferrovum_sp_PN-J185
Ferrovum_sp_21-44-67
Ferrovum_sp_JA12
Betaproteobacteria_bacterium_SG8_41
Betaproteobacteria_bacterium_RIFCSPLOWO2_02_FULL_63_19
Betaproteobacteria_bacterium_RIFCSPLOWO2_12_FULL_62_58
Betaproteobacteria_bacterium_RIFCSPLOWO2_12_FULL_63_13
Gammaproteobacteria
Thiotrichales
Methylophaga_thiooxydans
Beggiatoa_sp_IS2
Enterobacterales
Morganellaceae
Escherichia_coli
Photorhabdus_luminescens
Providencia_stuartii
Coxiella
Coxiella_burnetii_RSA_493
Coxiella_sp__in__Bacteria_
Coxiella_sp_RIFCSPHIGHO2_12_FULL_42_15
Coxiella_sp_RIFCSPHIGHO2_12_FULL_44_14
Chromatiales
Chromatiaceae
Imhoffiella_purpurea
Thiorhodococcus_drewsii
Ectothiorhodospiraceae
Aquisalimonas_asiatica
Thiorhodospira_sibirica
Alkalilimnicola_ehrlichii_MLHE-1
Thioalkalivibrio
Thioalkalivibrio_sulfidiphilus_HL-EbGr7
Thioalkalivibrio_thiocyanodenitrificans
Thioalkalivibrio_denitrificans
Gammaproteobacteria_bacterium_RIFCSPHIGHO2_12_FULL_35_23
Gammaproteobacteria_bacterium_RIFCSPHIGHO2_12_FULL_36_30
Gammaproteobacteria_bacterium_CG_4_10_14_0_2_um_filter_38_22
Gammaproteobacteria_bacterium_RIFCSPHIGHO2_12_FULL_38_11
Gammaproteobacteria_bacterium_RIFCSPHIGHO2_12_FULL_40_19
Gammaproteobacteria_bacterium_RIFCSPHIGHO2_12_FULL_41_20
Gammaproteobacteria_bacterium_RIFCSPHIGHO2_12_FULL_45_9
Gammaproteobacteria_bacterium_RIFCSPHIGHO2_01_FULL_42_8
Gammaproteobacteria_bacterium_RIFCSPHIGHO2_02_FULL_39_13

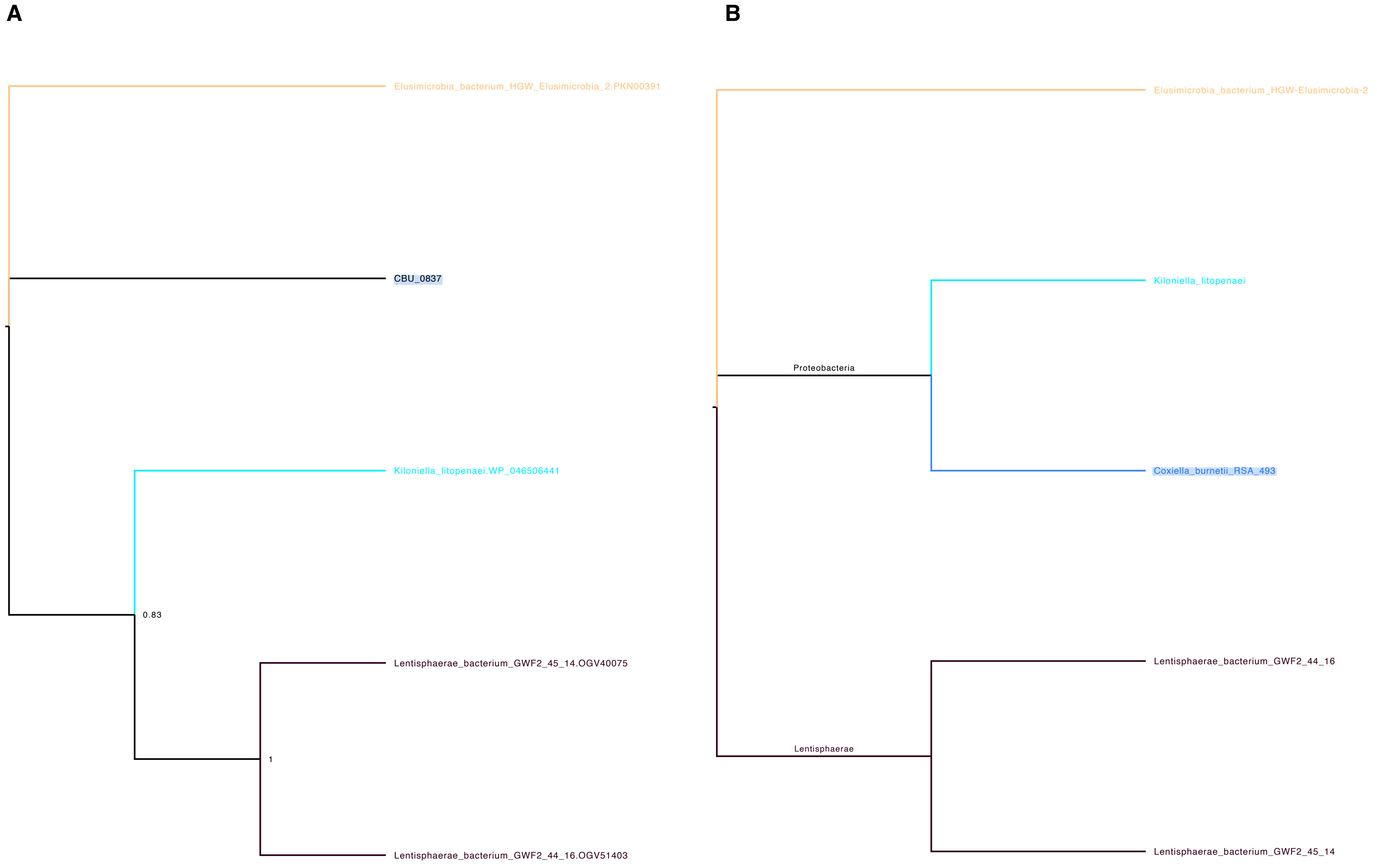
A
Elusimicrobia_bacterium_HGW_Elusimicrobia_2.PKN00391
CBU_0837
Kiloniella_litopenaei.WP_046506441
0.83
Lentisphaerae_bacterium_GWF2_45_14.OGV40075
1
Lentisphaerae_bacterium_GWF2_44_16.OGV51403
B
Elusimicrobia_bacterium_HGW-Elusimicrobia-2
Kiloniella_litopenaei
Proteobacteria
Coxiella_burnetii_RSA_493
Lentisphaerae_bacterium_GWF2_44_16
Lentisphaerae
Lentisphaerae_bacterium_GWF2_45_14

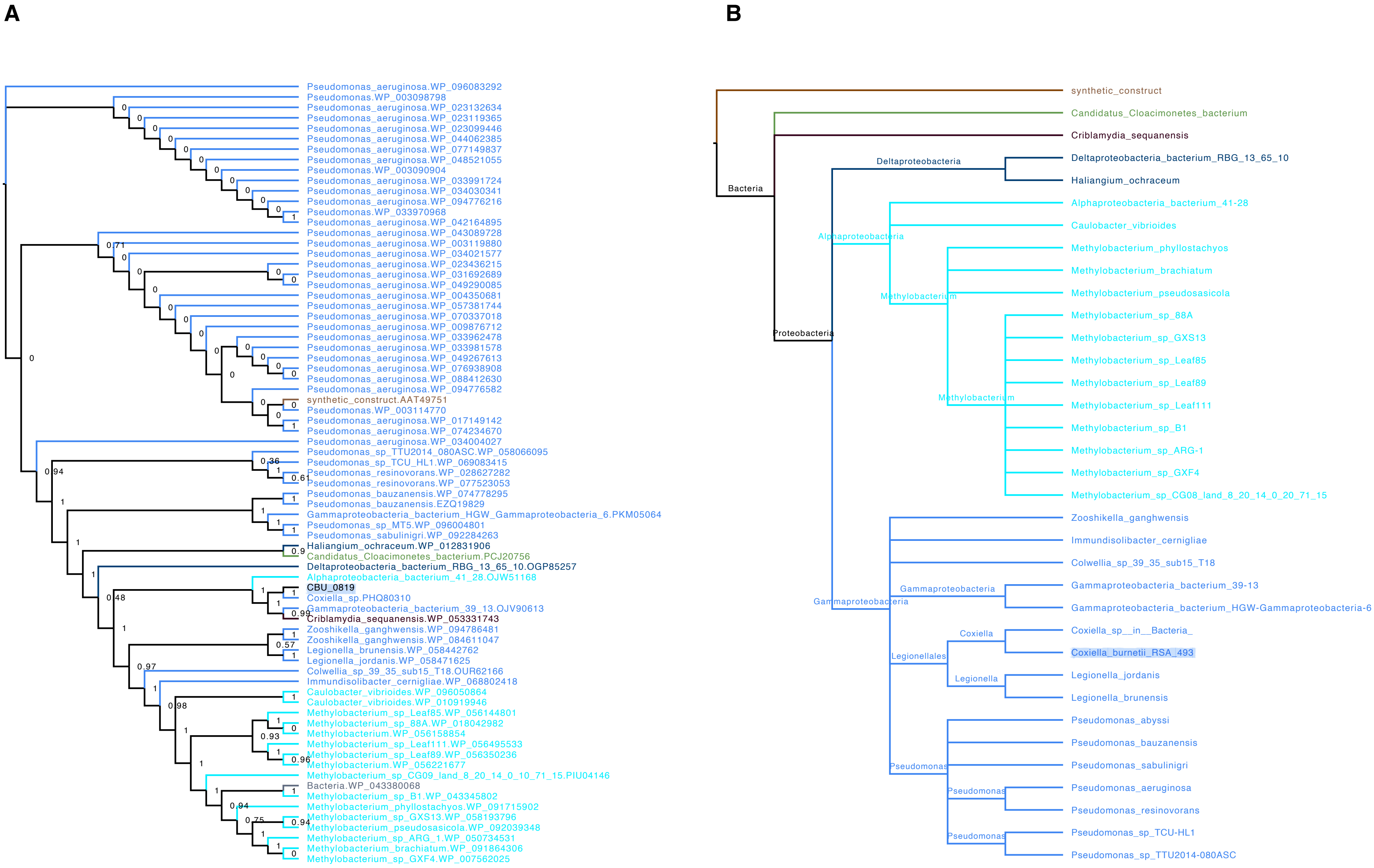
A
Pseudomonas_aeruginosa.WP_096083292
Pseudomonas.WP_003098798
Pseudomonas_aeruginosa.WP_023132634
Pseudomonas_aeruginosa.WP_023119365
Pseudomonas_aeruginosa.WP_023099446
Pseudomonas_aeruginosa.WP_044062385
Pseudomonas_aeruginosa.WP_077149837
Pseudomonas_aeruginosa.WP_048521055
Pseudomonas.WP_003090904
Pseudomonas_aeruginosa.WP_033991724
Pseudomonas_aeruginosa.WP_034030341
Pseudomonas_aeruginosa.WP_094776216
Pseudomonas.WP_033970968
Pseudomonas_aeruginosa.WP_042164895
Pseudomonas_aeruginosa.WP_043089728
Pseudomonas_aeruginosa.WP_003119880
Pseudomonas_aeruginosa.WP_034021577
Pseudomonas_aeruginosa.WP_023436215
Pseudomonas_aeruginosa.WP_031692689
Pseudomonas_aeruginosa.WP_049290085
Pseudomonas_aeruginosa.WP_004350681
Pseudomonas_aeruginosa.WP_057381744
Pseudomonas_aeruginosa.WP_070337018
Pseudomonas_aeruginosa.WP_009876712
Pseudomonas_aeruginosa.WP_033962478
Pseudomonas_aeruginosa.WP_033981578
Pseudomonas_aeruginosa.WP_049267613
Pseudomonas_aeruginosa.WP_076938908
Pseudomonas_aeruginosa.WP_088412630
Pseudomonas_aeruginosa.WP_094776582
synthetic_construct.AAT49751
Pseudomonas.WP_003114770
Pseudomonas_aeruginosa.WP_017149142
Pseudomonas_aeruginosa.WP_074234670
Pseudomonas_aeruginosa.WP_034004027
Pseudomonas_sp_TTU2014_080ASC.WP_058066095
Pseudomonas_sp_TCU_HL1.WP_069083415
Pseudomonas_resinovorans.WP_028627282
Pseudomonas_resinovorans.WP_077523053
Pseudomonas_bauzanensis.WP_074778295
Pseudomonas_bauzanensis.EZQ19829
Gammaproteobacteria_bacterium_HGW_Gammaproteobacteria_6.PKM05064
Pseudomonas_sp_MT5.WP_096004801
Pseudomonas_sabulinigri.WP_092284263
Haliangium_ochraceum.WP_012831906
Candidatus_Cloacimonetes_bacterium.PCJ20756
Deltaproteobacteria_bacterium_RBG_13_65_10.OGP85257
Alphaproteobacteria_bacterium_41_28.OJW51168
CBU_0819
Coxiella_sp.PHQ80310
Gammaproteobacteria_bacterium_39_13.OJV90613
Criblamydia_sequanensis.WP_053331743
Zooshikella_ganghwensis.WP_094786481
Zooshikella_ganghwensis.WP_084611047
Legionella_brunensis.WP_058442762
Legionella_jordanis.WP_058471625
Colwellia_sp_39_35_sub15_T18.OUR62166
Immundisolibacter_cernigliae.WP_068802418
Caulobacter_vibrioides.WP_096050864
Caulobacter_vibrioides.WP_010919946
Methylobacterium_sp_Leaf85.WP_056144801
Methylobacterium_sp_88A.WP_018042982
Methylobacterium.WP_056158854
Methylobacterium_sp_Leaf111.WP_056495533
Methylobacterium_sp_Leaf89.WP_056350236
Methylobacterium.WP_056221677
Methylobacterium_sp_CG09_land_8_20_14_0_10_71_15.PIU04146
Bacteria.WP_043380068
Methylobacterium_sp_B1.WP_043345802
Methylobacterium_phyllostachyos.WP_091715902
Methylobacterium_sp_GXS13.WP_058193796
Methylobacterium_pseudosasicola.WP_092039348
Methylobacterium_sp_ARG_1.WP_050734531
Methylobacterium_brachiatum.WP_091864306
Methylobacterium_sp_GXF4.WP_007562025
B
synthetic_construct
Candidatus_Cloacimonetes_bacterium
Criblamydia_sequanensis
Bacteria
Deltaproteobacteria
Deltaproteobacteria_bacterium_RBG_13_65_10
Haliangium_ochraceum
Alphaproteobacteria_bacterium_41-28
Caulobacter_vibrioides
Alphaproteobacteria
Methylobacterium_phyllostachyos
Methylobacterium_brachiatum
Methylobacterium_pseudosasicola
Methylobacterium
Methylobacterium_sp_88A
Methylobacterium_sp_GXS13
Methylobacterium_sp_Leaf85
Methylobacterium_sp_Leaf89
Methylobacterium_sp_Leaf111
Methylobacterium
Methylobacterium_sp_B1
Methylobacterium_sp_ARG-1
Methylobacterium_sp_GXF4
Methylobacterium_sp_CG08_land_8_20_14_0_20_71_15
Proteobacteria
Zooshikella_ganghwensis
Immundisolibacter_cernigliae
Colwellia_sp_39_35_sub15_T18
Gammaproteobacteria
Gammaproteobacteria_bacterium_39-13
Gammaproteobacteria
Gammaproteobacteria_bacterium_HGW-Gammaproteobacteria-6
Coxiella
Coxiella_sp__in__Bacteria_
Legionellales
Coxiella_burnetii_RSA_493
Legionella
Legionella_jordanis
Legionella_brunensis
Pseudomonas_abyssi
Pseudomonas_bauzanensis
Pseudomonas
Pseudomonas_sabulinigri
Pseudomonas
Pseudomonas_aeruginosa
Pseudomonas_resinovorans
Pseudomonas
Pseudomonas_sp_TCU-HL1
Pseudomonas_sp_TTU2014-080ASC

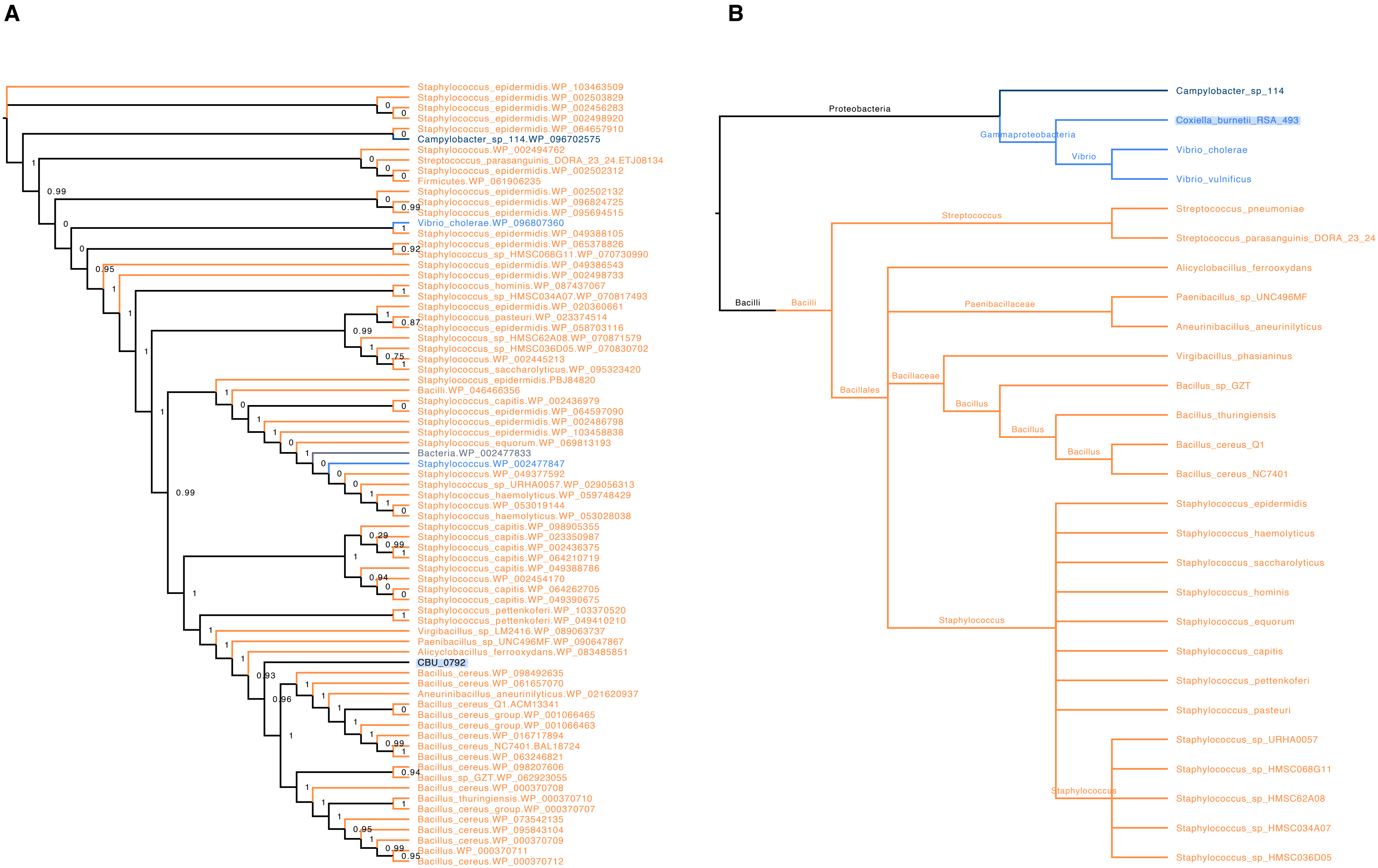
A
Staphylococcus_epidermidis.WP_103463509
Staphylococcus_epidermidis.WP_002503829
Staphylococcus_epidermidis.WP_002456283
Staphylococcus_epidermidis.WP_002498920
Staphylococcus_epidermidis.WP_064657910
Campylobacter_sp_114.WP_096702575
Staphylococcus.WP_002494762
Streptococcus_parasanguinis_DORA_23_24.ETJ08134
Staphylococcus_epidermidis.WP_002502312
Firmicutes.WP_061906235
Staphylococcus_epidermidis.WP_002502132
Staphylococcus_epidermidis.WP_096824725
Staphylococcus_epidermidis.WP_095694515
Vibrio_cholerae.WP_096807360
Staphylococcus_epidermidis.WP_049388105
Staphylococcus_epidermidis.WP_065378826
Staphylococcus_sp_HMSC068G11.WP_070730990
Staphylococcus_epidermidis.WP_049386543
Staphylococcus_epidermidis.WP_002498733
Staphylococcus_hominis.WP_087437067
Staphylococcus_sp_HMSC034A07.WP_070817493
Staphylococcus_epidermidis.WP_020360661
Staphylococcus_pasteuri.WP_023374514
Staphylococcus_epidermidis.WP_058703116
Staphylococcus_sp_HMSC62A08.WP_070871579
Staphylococcus_sp_HMSC036D05.WP_070830702
Staphylococcus.WP_002445213
Staphylococcus_saccharolyticus.WP_095323420
Staphylococcus_epidermidis.PBJ84820
Bacilli.WP_046466356
Staphylococcus_capitis.WP_002436979
Staphylococcus_epidermidis.WP_064597090
Staphylococcus_epidermidis.WP_002486798
Staphylococcus_epidermidis.WP_103458838
Staphylococcus_equorum.WP_069813193
Bacteria.WP_002477833
Staphylococcus.WP_002477847
Staphylococcus.WP_049377592
Staphylococcus_sp_URHA0057.WP_029056313
Staphylococcus_haemolyticus.WP_059748429
Staphylococcus.WP_053019144
Staphylococcus_haemolyticus.WP_053028038
Staphylococcus_capitis.WP_098905355
Staphylococcus_capitis.WP_023350987
Staphylococcus_capitis.WP_002436375
Staphylococcus_capitis.WP_064210719
Staphylococcus_capitis.WP_049388786
Staphylococcus.WP_002454170
Staphylococcus_capitis.WP_064262705
Staphylococcus_capitis.WP_049390675
Staphylococcus_pettenkoferi.WP_103370520
Staphylococcus_pettenkoferi.WP_049410210
Virgibacillus_sp_LM2416.WP_089063737
Paenibacillus_sp_UNC496MF.WP_090647867
Alicyclobacillus_ferrooxydans.WP_083485851
CBU_0792
Bacillus_cereus.WP_098492635
Bacillus_cereus.WP_061657070
Aneurinibacillus_aneurinilyticus.WP_021620937
Bacillus_cereus_Q1.ACM13341
Bacillus_cereus_group.WP_001066465
Bacillus_cereus_group.WP_001066463
Bacillus_cereus.WP_016717894
Bacillus_cereus_NC7401.BAL18724
Bacillus_cereus.WP_063246821
Bacillus_cereus.WP_098207606
Bacillus_sp_GZT.WP_062923055
Bacillus_cereus.WP_000370708
Bacillus_thuringiensis.WP_000370710
Bacillus_cereus_group.WP_000370707
Bacillus_cereus.WP_073542135
Bacillus_cereus.WP_095843104
Bacillus_cereus.WP_000370709
Bacillus.WP_000370711
Bacillus_cereus.WP_000370712
B
Proteobacteria
Gammaproteobacteria
Vibrio
Campylobacter_sp_114
Coxiella_burnetii_RSA_493
Vibrio_cholerae
Vibrio_vulnificus
Bacilli
Streptococcus
Streptococcus_pneumoniae
Streptococcus_parasanguinis_DORA_23_24
Alicyclobacillus_ferrooxydans
Paenibacillaceae
Paenibacillus_sp_UNC496MF
Aneurinibacillus_aneurinilyticus
Bacillales
Bacillaceae
Bacillus
Virgibacillus_phasianinus
Bacillus_sp_GZT
Bacillus_thuringiensis
Bacillus_cereus_Q1
Bacillus_cereus_NC7401
Staphylococcus
Staphylococcus_epidermidis
Staphylococcus_haemolyticus
Staphylococcus_saccharolyticus
Staphylococcus_hominis
Staphylococcus_equorum
Staphylococcus_capitis
Staphylococcus_pettenkoferi
Staphylococcus_pasteuri
Staphylococcus_sp_URHA0057
Staphylococcus_sp_HMSC068G11
Staphylococcus_sp_HMSC62A08
Staphylococcus_sp_HMSC034A07
Staphylococcus_sp_HMSC036D05

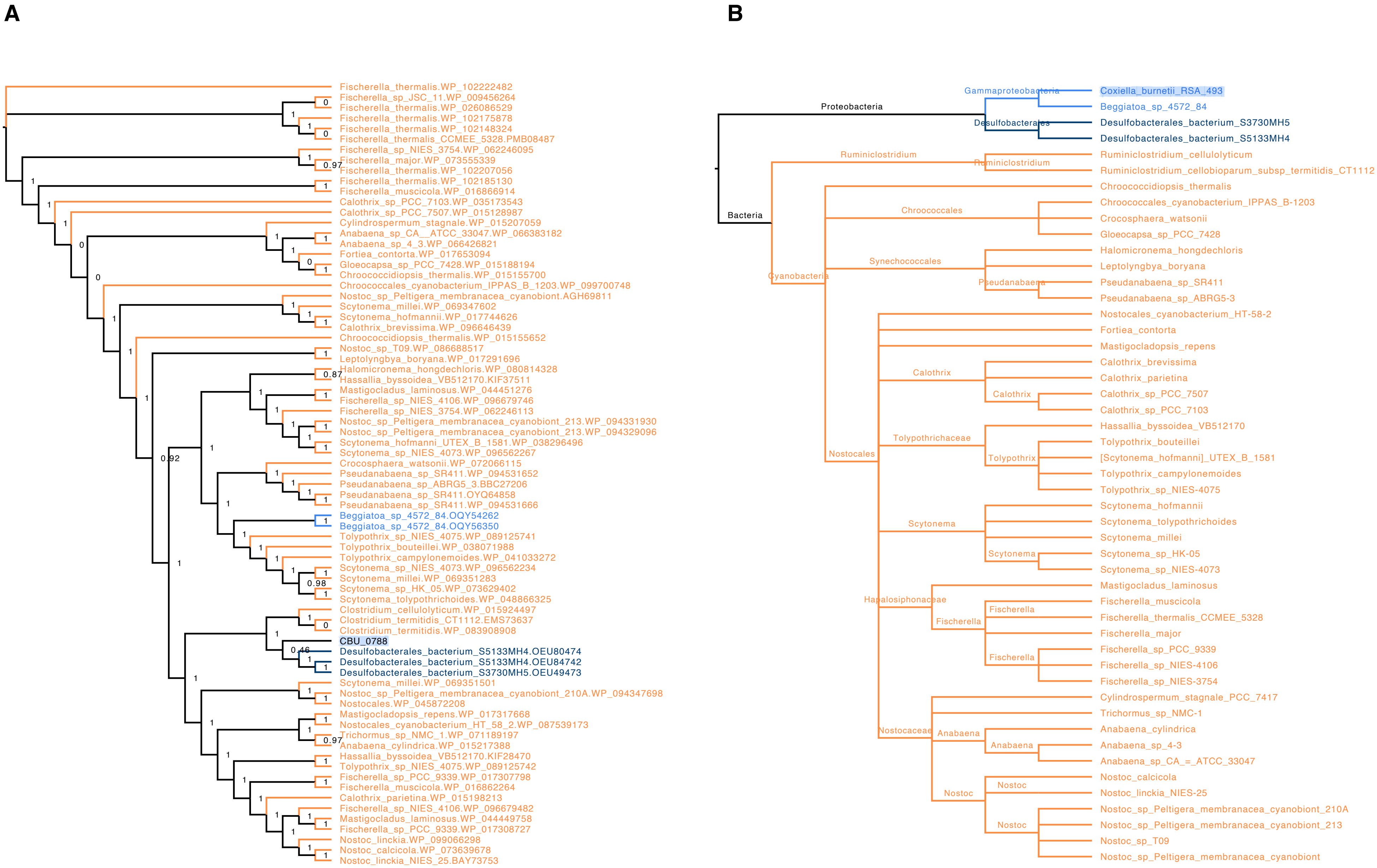
A
Fischerella_thermalis.WP_102222482
Fischerella_sp_JSC_11.WP_009456264
Fischerella_thermalis.WP_026086529
Fischerella_thermalis.WP_102175878
Fischerella_thermalis.WP_102148324
Fischerella_thermalis_CCMEE_5328.PMB08487
Fischerella_sp_NIES_3754.WP_062246095
Fischerella_major.WP_073555339
Fischerella_thermalis.WP_102207056
Fischerella_thermalis.WP_102185130
Fischerella_muscicola.WP_016866914
Calothrix_sp_PCC_7103.WP_035173543
Calothrix_sp_PCC_7507.WP_015128987
Cylindrospermum_stagnale.WP_015207059
Anabaena_sp_CA__ATCC_33047.WP_066383182
Anabaena_sp_4_3.WP_066426821
Fortiea_contorta.WP_017653094
Gloeocapsa_sp_PCC_7428.WP_015188194
Chroococcidiopsis_thermalis.WP_015155700
Chroococcales_cyanobacterium_IPPAS_B_1203.WP_099700748
Nostoc_sp_Peltigera_membranacea_cyanobiont.AGH69811
Scytonema_millei.WP_069347602
Scytonema_hofmannii.WP_017744626
Calothrix_brevissima.WP_096646439
Chroococcidiopsis_thermalis.WP_015155652
Nostoc_sp_T09.WP_086688517
Leptolyngbya_boryana.WP_017291696
Halomicronema_hongdechloris.WP_080814328
Hassallia_byssoidea_VB512170.KIF37511
Mastigocladus_laminosus.WP_044451276
Fischerella_sp_NIES_4106.WP_096679746
Fischerella_sp_NIES_3754.WP_062246113
Nostoc_sp_Peltigera_membranacea_cyanobiont_213.WP_094331930
Nostoc_sp_Peltigera_membranacea_cyanobiont_213.WP_094329096
Scytonema_hofmanni_UTEX_B_1581.WP_038296496
Scytonema_sp_NIES_4073.WP_096562267
Crocosphaera_watsonii.WP_072066115
Pseudanabaena_sp_SR411.WP_094531652
Pseudanabaena_sp_ABRG5_3.BBC27206
Pseudanabaena_sp_SR411.OYQ64858
Pseudanabaena_sp_SR411.WP_094531666
Beggiatoa_sp_4572_84.OQY54262
Beggiatoa_sp_4572_84.OQY56350
Tolypothrix_sp_NIES_4075.WP_089125741
Tolypothrix_bouteillei.WP_038071988
Tolypothrix_campylonemoides.WP_041033272
Scytonema_sp_NIES_4073.WP_096562234
Scytonema_millei.WP_069351283
Scytonema_sp_HK_05.WP_073629402
Scytonema_tolypothrichoides.WP_048866325
Clostridium_cellulolyticum.WP_015924497
Clostridium_termitidis_CT1112.EMS73637
Clostridium_termitidis.WP_083908908
CBU_0788
Desulfobacterales_bacterium_S5133MH4.OEU80474
Desulfobacterales_bacterium_S5133MH4.OEU84742
Desulfobacterales_bacterium_S3730MH5.OEU49473
Scytonema_millei.WP_069351501
Nostoc_sp_Peltigera_membranacea_cyanobiont_210A.WP_094347698
Nostocales.WP_045872208
Mastigocladopsis_repens.WP_017317668
Nostocales_cyanobacterium_HT_58_2.WP_087539173
Trichormus_sp_NMC_1.WP_071189197
Anabaena_cylindrica.WP_015217388
Hassallia_byssoidea_VB512170.KIF28470
Tolypothrix_sp_NIES_4075.WP_089125742
Fischerella_sp_PCC_9339.WP_017307798
Fischerella_muscicola.WP_016862264
Calothrix_parietina.WP_015198213
Fischerella_sp_NIES_4106.WP_096679482
Mastigocladus_laminosus.WP_044449758
Fischerella_sp_PCC_9339.WP_017308727
Nostoc_linckia.WP_099066298
Nostoc_calcicola.WP_073639678
Nostoc_linckia_NIES_25.BAY73753
B
Proteobacteria
Gammaproteobacteria
Desulfobacterales
Bacteria
Ruminiclostridium
Cyanobacteria
Chroococcales
Synechococcales
Pseudanabaena
Nostocales
Calothrix
Tolypothrichaceae
Tolypothrix
Scytonema
Hapalosiphonaceae
Fischerella
Nostocaceae
Anabaena
Nostoc
Coxiella_burnetii_RSA_493
Beggiatoa_sp_4572_84
Desulfobacterales_bacterium_S3730MH5
Desulfobacterales_bacterium_S5133MH4
Ruminiclostridium_cellulolyticum
Ruminiclostridium_cellobioparum_subsp_termitidis_CT1112
Chroococcidiopsis_thermalis
Chroococcales_cyanobacterium_IPPAS_B-1203
Crocosphaera_watsonii
Gloeocapsa_sp_PCC_7428
Halomicronema_hongdechloris
Leptolyngbya_boryana
Pseudanabaena_sp_SR411
Pseudanabaena_sp_ABRG5-3
Nostocales_cyanobacterium_HT-58-2
Fortiea_contorta
Mastigocladopsis_repens
Calothrix_brevissima
Calothrix_parietina
Calothrix_sp_PCC_7507
Calothrix_sp_PCC_7103
Hassallia_byssoidea_VB512170
Tolypothrix_bouteillei
[Scytonema_hofmanni]_UTEX_B_1581
Tolypothrix_campylonemoides
Tolypothrix_sp_NIES-4075
Scytonema_hofmannii
Scytonema_tolypothrichoides
Scytonema_millei
Scytonema_sp_HK-05
Scytonema_sp_NIES-4073
Mastigocladus_laminosus
Fischerella_muscicola
Fischerella_thermalis_CCMEE_5328
Fischerella_major
Fischerella_sp_PCC_9339
Fischerella_sp_NIES-4106
Fischerella_sp_NIES-3754
Cylindrospermum_stagnale_PCC_7417
Trichormus_sp_NMC-1
Anabaena_cylindrica
Anabaena_sp_4-3
Anabaena_sp_CA_=_ATCC_33047
Nostoc_calcicola
Nostoc_linckia_NIES-25
Nostoc_sp_Peltigera_membranacea_cyanobiont_210A
Nostoc_sp_Peltigera_membranacea_cyanobiont_213
Nostoc_sp_T09
Nostoc_sp_Peltigera_membranacea_cyanobiont

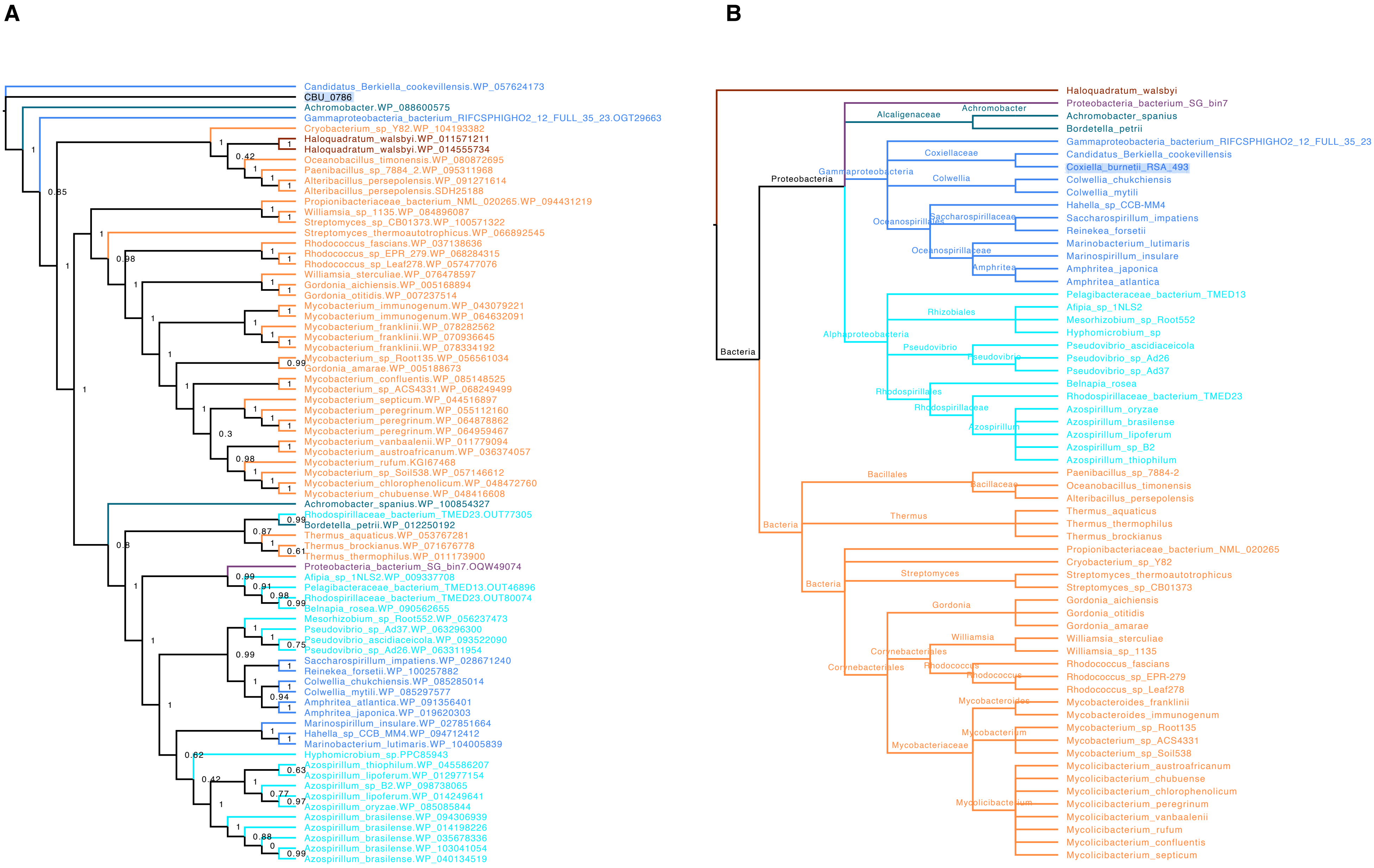
A
Candidatus_Berkiella_cookevillensis.WP_057624173
CBU_0786
Achromobacter.WP_088600575
Gammaproteobacteria_bacterium_RIFCSPHIGHO2_12_FULL_35_23.OGT29663
Cryobacterium_sp_Y82.WP_104193382
Haloquadratum_walsbyi.WP_011571211
Haloquadratum_walsbyi.WP_014555734
Oceanobacillus_timonensis.WP_080872695
Paenibacillus_sp_7884_2.WP_095311968
Alteribacillus_persepolensis.WP_091271614
Alteribacillus_persepolensis.SDH25188
Propionibacteriaceae_bacterium_NML_020265.WP_094431219
Williamsia_sp_1135.WP_084896087
Streptomyces_sp_CB01373.WP_100571322
Streptomyces_thermoautotrophicus.WP_066892545
Rhodococcus_fascians.WP_037138636
Rhodococcus_sp_EPR_279.WP_068284315
Rhodococcus_sp_Leaf278.WP_057477076
Williamsia_sterculiae.WP_076478597
Gordonia_aichiensis.WP_005168894
Gordonia_otitidis.WP_007237514
Mycobacterium_immunogenum.WP_043079221
Mycobacterium_immunogenum.WP_064632091
Mycobacterium_franklinii.WP_078282562
Mycobacterium_franklinii.WP_070936645
Mycobacterium_franklinii.WP_078334192
Mycobacterium_sp_Root135.WP_056561034
Gordonia_amarae.WP_005188673
Mycobacterium_confluentis.WP_085148525
Mycobacterium_sp_ACS4331.WP_068249499
Mycobacterium_septicum.WP_044516897
Mycobacterium_peregrinum.WP_055112160
Mycobacterium_peregrinum.WP_064878862
Mycobacterium_peregrinum.WP_064959467
Mycobacterium_vanbaalenii.WP_011779094
Mycobacterium_austroafricanum.WP_036374057
Mycobacterium_rufum.KGI67468
Mycobacterium_sp_Soil538.WP_057146612
Mycobacterium_chlorophenolicum.WP_048472760
Mycobacterium_chubuense.WP_048416608
Achromobacter_spanius.WP_100854327
Rhodospirillaceae_bacterium_TMED23.OUT77305
Bordetella_petrii.WP_012250192
Thermus_aquaticus.WP_053767281
Thermus_brockianus.WP_071676778
Thermus_thermophilus.WP_011173900
Proteobacteria_bacterium_SG_bin7.OQW49074
Afipia_sp_1NLS2.WP_009337708
Pelagibacteraceae_bacterium_TMED13.OUT46896
Rhodospirillaceae_bacterium_TMED23.OUT80074
Belnapia_rosea.WP_090562655
Mesorhizobium_sp_Root552.WP_056237473
Pseudovibrio_sp_Ad37.WP_063296300
Pseudovibrio_ascidiaceicola.WP_093522090
Pseudovibrio_sp_Ad26.WP_063311954
Saccharospirillum_impatiens.WP_028671240
Reinekea_forsetii.WP_100257882
Colwellia_chukchiensis.WP_085285014
Colwellia_mytili.WP_085297577
Amphritea_atlantica.WP_091356401
Amphritea_japonica.WP_019620303
Marinospirillum_insulare.WP_027851664
Hahella_sp_CCB_MM4.WP_094712412
Marinobacterium_lutimaris.WP_104005839
Hyphomicrobium_sp.PPC85943
Azospirillum_thiophilum.WP_045586207
Azospirillum_lipoferum.WP_012977154
Azospirillum_sp_B2.WP_098738065
Azospirillum_lipoferum.WP_014249641
Azospirillum_oryzae.WP_085085844
Azospirillum_brasilense.WP_094306939
Azospirillum_brasilense.WP_014198226
Azospirillum_brasilense.WP_035678336
Azospirillum_brasilense.WP_103041054
Azospirillum_brasilense.WP_040134519
B
Haloquadratum_walsbyi
Proteobacteria_bacterium_SG_bin7
Achromobacter_spanius
Bordetella_petrii
Gammaproteobacteria_bacterium_RIFCSPHIGHO2_12_FULL_35_23
Candidatus_Berkiella_cookevillensis
Coxiella_burnetii_RSA_493
Colwellia_chukchiensis
Colwellia_mytili
Hahella_sp_CCB-MM4
Saccharospirillum_impatiens
Reinekea_forsetii
Marinobacterium_lutimaris
Marinospirillum_insulare
Amphritea_japonica
Amphritea_atlantica
Pelagibacteraceae_bacterium_TMED13
Afipia_sp_1NLS2
Mesorhizobium_sp_Root552
Hyphomicrobium_sp
Pseudovibrio_ascidiaceicola
Pseudovibrio_sp_Ad26
Pseudovibrio_sp_Ad37
Belnapia_rosea
Rhodospirillaceae_bacterium_TMED23
Azospirillum_oryzae
Azospirillum_brasilense
Azospirillum_lipoferum
Azospirillum_sp_B2
Azospirillum_thiophilum
Paenibacillus_sp_7884-2
Oceanobacillus_timonensis
Alteribacillus_persepolensis
Thermus_aquaticus
Thermus_thermophilus
Thermus_brockianus
Propionibacteriaceae_bacterium_NML_020265
Cryobacterium_sp_Y82
Streptomyces_thermoautotrophicus
Streptomyces_sp_CB01373
Gordonia_aichiensis
Gordonia_otitidis
Gordonia_amarae
Williamsia_sterculiae
Williamsia_sp_1135
Rhodococcus_fascians
Rhodococcus_sp_EPR-279
Rhodococcus_sp_Leaf278
Mycobacteroides_franklinii
Mycobacteroides_immunogenum
Mycobacterium_sp_Root135
Mycobacterium_sp_ACS4331
Mycobacterium_sp_Soil538
Mycolicibacterium_austroafricanum
Mycolicibacterium_chubuense
Mycolicibacterium_chlorophenolicum
Mycolicibacterium_peregrinum
Mycolicibacterium_vanbaalenii
Mycolicibacterium_rufum
Mycolicibacterium_confluentis
Mycolicibacterium_septicum
Alcaligenaceae
Achromobacter
Proteobacteria
Gammaproteobacteria
Coxiellaceae
Colwellia
Oceanospirillales
Saccharospirillaceae
Oceanospirillaceae
Amphritea
Bacteria
Alphaproteobacteria
Rhizobiales
Pseudovibrio
Rhodospirillales
Rhodospirillaceae
Azospirillum
Bacillales
Bacillaceae
Thermus
Streptomyces
Gordonia
Williamsia
Corynebacteriales
Rhodococcus
Mycobacteroides
Mycobacterium
Mycobacteriaceae
Mycolicibacterium

A

B

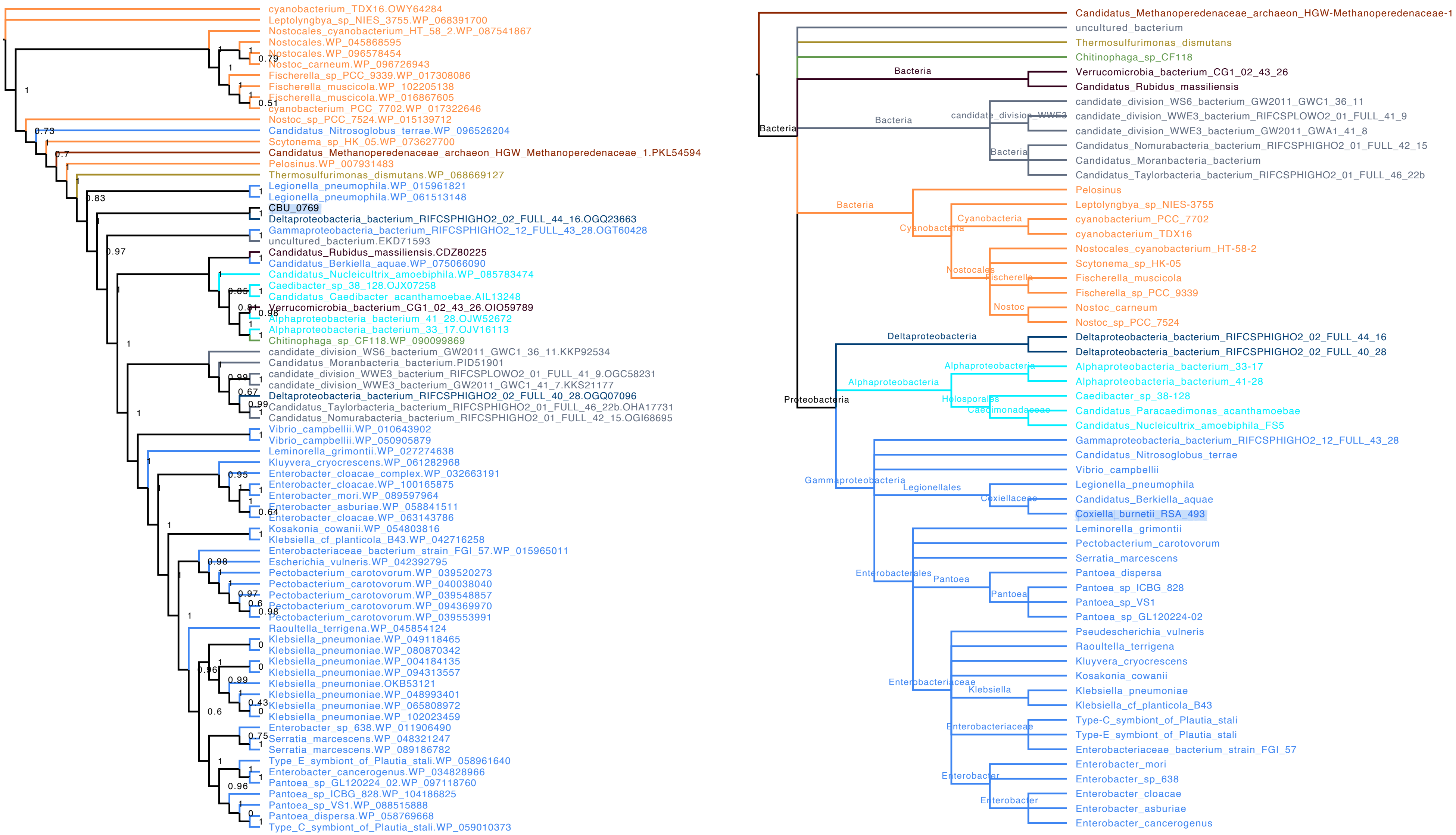

cyanobacterium_TDX16.OWY64284
Leptolyngbya_sp_NIES_3755.WP_068391700
Nostocales_cyanobacterium_HT_58_2.WP_087541867
Nostocales.WP_045868595
Nostocales.WP_096578454
Nostoc_carneum.WP_096726943
Fischerella_sp_PCC_9339.WP_017308086
Fischerella_muscicola.WP_102205138
Fischerella_muscicola.WP_016867605
cyanobacterium_PCC_7702.WP_017322646
Nostoc_sp_PCC_7524.WP_015139712
Candidatus_Nitrosoglobus_terrae.WP_096526204
Scytonema_sp_HK_05.WP_073627700
Candidatus_Methanoperedenaceae_archaeon_HGW_Methanoperedenaceae_1.PKL54594
Pelosinus.WP_007931483
Thermosulfurimonas_dismutans.WP_068669127
Legionella_pneumophila.WP_015961821
Legionella_pneumophila.WP_061513148
CBU_0769
Deltaproteobacteria_bacterium_RIFCSPHIGHO2_02_FULL_44_16.OGQ23663
Gammaproteobacteria_bacterium_RIFCSPHIGHO2_12_FULL_43_28.OGT60428
uncultured_bacterium.EKD71593
Candidatus_Rubidus_massiliensis.CDZ80225
Candidatus_Berkiella_aquae.WP_075066090
Candidatus_Nucleicultrix_amoebiphila.WP_085783474
Caedibacter_sp_38_128.OJX07258
Candidatus_Caedibacter_acanthamoebae.AIL13248
Verrucomicrobia_bacterium_CG1_02_43_26.OIO59789
Alphaproteobacteria_bacterium_41_28.OJW52672
Alphaproteobacteria_bacterium_33_17.OJV16113
Chitinophaga_sp_CF118.WP_090099869
candidate_division_WS6_bacterium_GW2011_GWC1_36_11.KKP92534
Candidatus_Moranbacteria_bacterium.PID51901
candidate_division_WWE3_bacterium_RIFCSPLOWO2_01_FULL_41_9.OGC58231
candidate_division_WWE3_bacterium_GW2011_GWC1_41_7.KKS21177
Deltaproteobacteria_bacterium_RIFCSPHIGHO2_02_FULL_40_28.OGQ07096
Candidatus_Taylorbacteria_bacterium_RIFCSPHIGHO2_01_FULL_46_22b.OHA17731
Candidatus_Nomurabacteria_bacterium_RIFCSPHIGHO2_01_FULL_42_15.OGI68695
Vibrio_campbellii.WP_010643902
Vibrio_campbellii.WP_050905879
Leminorella_grimontii.WP_027274638
Kluyvera_cryocrescens.WP_061282968
Enterobacter_cloacae_complex.WP_032663191
Enterobacter_cloacae.WP_100165875
Enterobacter_mori.WP_089597964
Enterobacter_asburiae.WP_058841511
Enterobacter_cloacae.WP_063143786
Kosakonia_cowanii.WP_054803816
Klebsiella_cf_planticola_B43.WP_042716258
Enterobacteriaceae_bacterium_strain_FGI_57.WP_015965011
Escherichia_vulneris.WP_042392795
Pectobacterium_carotovorum.WP_039520273
Pectobacterium_carotovorum.WP_040038040
Pectobacterium_carotovorum.WP_039548857
Pectobacterium_carotovorum.WP_094369970
Pectobacterium_carotovorum.WP_039553991
Raoultella_terrigena.WP_045854124
Klebsiella_pneumoniae.WP_049118465
Klebsiella_pneumoniae.WP_080870342
Klebsiella_pneumoniae.WP_004184135
Klebsiella_pneumoniae.WP_094313557
Klebsiella_pneumoniae.OKB53121
Klebsiella_pneumoniae.WP_048993401
Klebsiella_pneumoniae.WP_065808972
Klebsiella_pneumoniae.WP_102023459
Enterobacter_sp_638.WP_011906490
Serratia_marcescens.WP_048321247
Serratia_marcescens.WP_089186782
Type_E_symbiont_of_Plautia_stali.WP_058961640
Enterobacter_cancerogenus.WP_034828966
Pantoea_sp_GL120224_02.WP_097118760
Pantoea_sp_ICBG_828.WP_104186825
Pantoea_sp_VS1.WP_088515888
Pantoea_dispersa.WP_058769668
Type_C_symbiont_of_Plautia_stali.WP_059010373
Candidatus_Methanoperedenaceae_archaeon_HGW-Methanoperedenaceae-1
uncultured_bacterium
Thermosulfurimonas_dismutans
Chitinophaga_sp_CF118
Bacteria
Verrucomicrobia_bacterium_CG1_02_43_26
Candidatus_Rubidus_massiliensis
candidate_division_WS6_bacterium_GW2011_GWC1_36_11
candidate_division_WWE3
candidate_division_WWE3_bacterium_RIFCSPLOWO2_01_FULL_41_9
candidate_division_WWE3_bacterium_GW2011_GWA1_41_8
Candidatus_Nomurabacteria_bacterium_RIFCSPHIGHO2_01_FULL_42_15
Candidatus_Moranbacteria_bacterium
Candidatus_Taylorbacteria_bacterium_RIFCSPHIGHO2_01_FULL_46_22b
Pelosinus
Cyanobacteria
Leptolyngbya_sp_NIES-3755
cyanobacterium_PCC_7702
cyanobacterium_TDX16
Nostocales
Nostocales_cyanobacterium_HT-58-2
Scytonema_sp_HK-05
Fischerella
Fischerella_muscicola
Fischerella_sp_PCC_9339
Nostoc
Nostoc_carneum
Nostoc_sp_PCC_7524
Proteobacteria
Deltaproteobacteria
Deltaproteobacteria_bacterium_RIFCSPHIGHO2_02_FULL_44_16
Deltaproteobacteria_bacterium_RIFCSPHIGHO2_02_FULL_40_28
Alphaproteobacteria
Alphaproteobacteria_bacterium_33-17
Alphaproteobacteria_bacterium_41-28
Holosporales
Caedimonadaceae
Caedibacter_sp_38-128
Candidatus_Paracaedimonas_acanthamoebae
Candidatus_Nucleicultrix_amoebiphila_FS5
Gammaproteobacteria
Gammaproteobacteria_bacterium_RIFCSPHIGHO2_12_FULL_43_28
Candidatus_Nitrosoglobus_terrae
Vibrio_campbellii
Legionellales
Coxiellaceae
Legionella_pneumophila
Candidatus_Berkiella_aquae
Coxiella_burnetii_RSA_493
Enterobacterales
Leminorella_grimontii
Pectobacterium_carotovorum
Serratia_marcescens
Pantoea
Pantoea_dispersa
Pantoea_sp_ICBG_828
Pantoea_sp_VS1
Pantoea_sp_GL120224-02
Enterobacteriaceae
Pseudescherichia_vulneris
Raoultella_terrigena
Kluyvera_cryocrescens
Kosakonia_cowanii
Klebsiella
Klebsiella_pneumoniae
Klebsiella_cf_planticola_B43
Type-C_symbiont_of_Plautia_stali
Type-E_symbiont_of_Plautia_stali
Enterobacteriaceae_bacterium_strain_FGI_57
Enterobacter
Enterobacter_mori
Enterobacter_sp_638
Enterobacter_cloacae
Enterobacter_asburiae
Enterobacter_cancerogenus

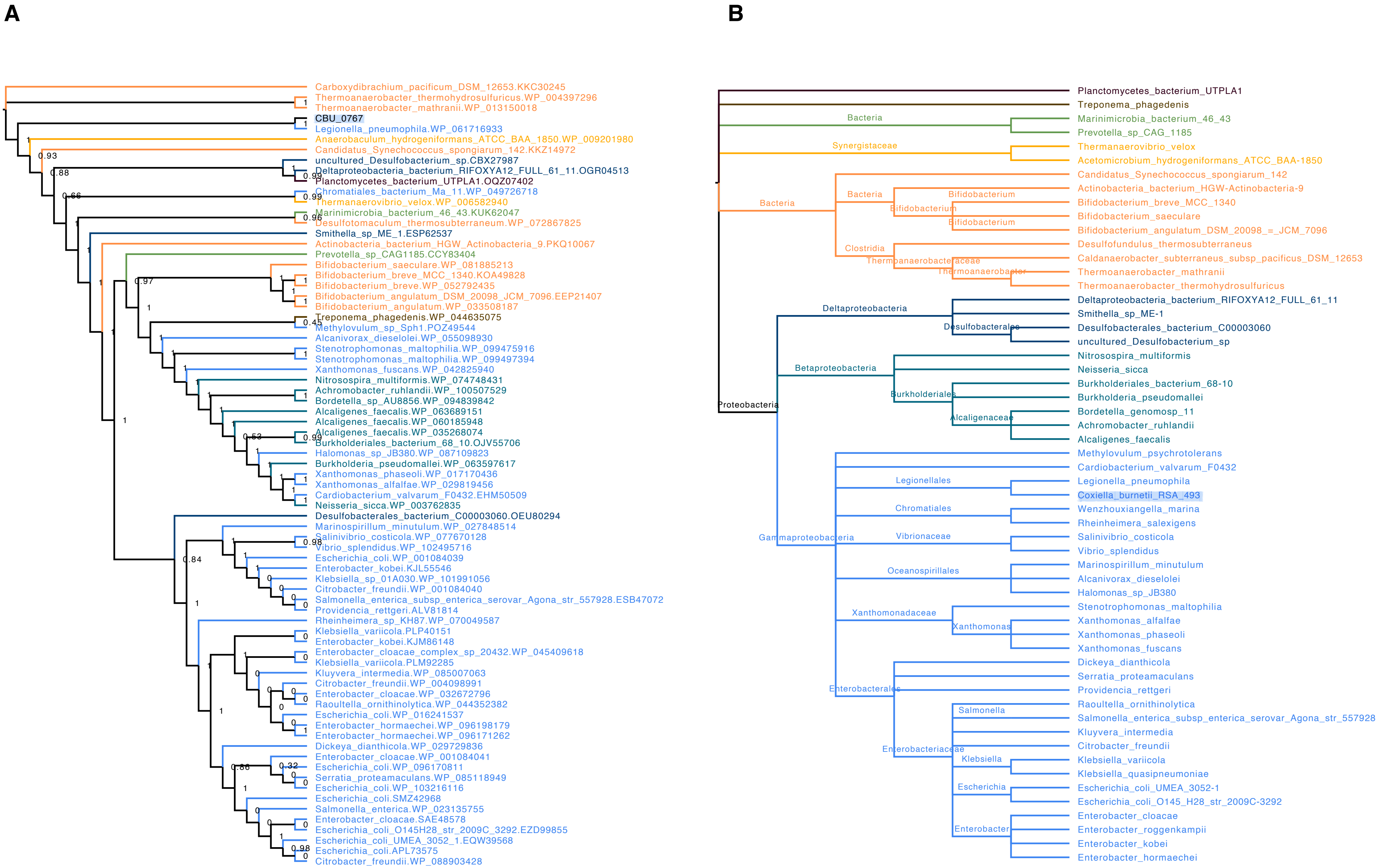
A
Carboxydibrachium_pacificum_DSM_12653.KKC30245
Thermoanaerobacter_thermohydrosulfuricus.WP_004397296
Thermoanaerobacter_mathranii.WP_013150018
CBU_0767
Legionella_pneumophila.WP_061716933
Anaerobaculum_hydrogeniformans_ATCC_BAA_1850.WP_009201980
Candidatus_Synechococcus_spongiarum_142.KKZ14972
uncultured_Desulfobacterium_sp.CBX27987
Deltaproteobacteria_bacterium_RIFOXYA12_FULL_61_11.OGR04513
Planctomycetes_bacterium_UTPLA1.OQZ07402
Chromatiales_bacterium_Ma_11.WP_049726718
Thermanaerovibrio_velox.WP_006582940
Marinimicrobia_bacterium_46_43.KUK62047
Desulfotomaculum_thermosubterraneum.WP_072867825
Smithella_sp_ME_1.ESP62537
Actinobacteria_bacterium_HGW_Actinobacteria_9.PKQ10067
Prevotella_sp_CAG1185.CCY83404
Bifidobacterium_saeculare.WP_081885213
Bifidobacterium_breve_MCC_1340.KOA49828
Bifidobacterium_breve.WP_052792435
Bifidobacterium_angulatum_DSM_20098_JCM_7096.EEP21407
Bifidobacterium_angulatum.WP_033508187
Treponema_phagedenis.WP_044635075
Methylovulum_sp_Sph1.POZ49544
Alcanivorax_dieselolei.WP_055098930
Stenotrophomonas_maltophilia.WP_099475916
Stenotrophomonas_maltophilia.WP_099497394
Xanthomonas_fuscans.WP_042825940
Nitrosospira_multiformis.WP_074748431
Achromobacter_ruhlandii.WP_100507529
Bordetella_sp_AU8856.WP_094839842
Alcaligenes_faecalis.WP_063689151
Alcaligenes_faecalis.WP_060185948
Alcaligenes_faecalis.WP_035268074
Burkholderiales_bacterium_68_10.OJV55706
Halomonas_sp_JB380.WP_087109823
Burkholderia_pseudomallei.WP_063597617
Xanthomonas_phaseoli.WP_017170436
Xanthomonas_alfalfae.WP_029819456
Cardiobacterium_valvarum_F0432.EHM50509
Neisseria_sicca.WP_003762835
Desulfobacterales_bacterium_C00003060.OEU80294
Marinospirillum_minutulum.WP_027848514
Salinivibrio_costicola.WP_077670128
Vibrio_splendidus.WP_102495716
Escherichia_coli.WP_001084039
Enterobacter_kobei.KJL55546
Klebsiella_sp_01A030.WP_101991056
Citrobacter_freundii.WP_001084040
Salmonella_enterica_subsp_enterica_serovar_Agona_str_557928.ESB47072
Providencia_rettgeri.ALV81814
Rheinheimera_sp_KH87.WP_070049587
Klebsiella_variicola.PLP40151
Enterobacter_kobei.KJM86148
Enterobacter_cloacae_complex_sp_20432.WP_045409618
Klebsiella_variicola.PLM92285
Kluyvera_intermedia.WP_085007063
Citrobacter_freundii.WP_004098991
Enterobacter_cloacae.WP_032672796
Raoultella_ornithinolytica.WP_044352382
Escherichia_coli.WP_016241537
Enterobacter_hormaechei.WP_096198179
Enterobacter_hormaechei.WP_096171262
Dickeya_dianthicola.WP_029729836
Enterobacter_cloacae.WP_001084041
Escherichia_coli.WP_096170811
Serratia_proteamaculans.WP_085118949
Escherichia_coli.WP_103216116
Escherichia_coli.SMZ42968
Salmonella_enterica.WP_023135755
Enterobacter_cloacae.SAE48578
Escherichia_coli_O145H28_str_2009C_3292.EZD99855
Escherichia_coli_UMEA_3052_1.EQW39568
Escherichia_coli.APL73575
Citrobacter_freundii.WP_088903428
B
Planctomycetes_bacterium_UTPLA1
Treponema_phagedenis
Bacteria
Marinimicrobia_bacterium_46_43
Prevotella_sp_CAG_1185
Synergistaceae
Thermanaerovibrio_velox
Acetomicrobium_hydrogeniformans_ATCC_BAA-1850
Bacteria
Candidatus_Synechococcus_spongiarum_142
Bacteria
Bifidobacterium
Actinobacteria_bacterium_HGW-Actinobacteria-9
Bifidobacterium
Bifidobacterium_breve_MCC_1340
Bifidobacterium
Bifidobacterium_saeculare
Bifidobacterium_angulatum_DSM_20098_=_JCM_7096
Clostridia
Desulfofundulus_thermosubterraneus
Thermoanaerobacteraceae
Caldanaerobacter_subterraneus_subsp_pacificus_DSM_12653
Thermoanaerobacter
Thermoanaerobacter_mathranii
Thermoanaerobacter_thermohydrosulfuricus
Deltaproteobacteria
Deltaproteobacteria_bacterium_RIFOXYA12_FULL_61_11
Smithella_sp_ME-1
Desulfobacterales
Desulfobacterales_bacterium_C00003060
uncultured_Desulfobacterium_sp
Nitrosospira_multiformis
Betaproteobacteria
Neisseria_sicca
Burkholderiales_bacterium_68-10
Burkholderiales
Burkholderia_pseudomallei
Proteobacteria
Bordetella_genomosp_11
Alcaligenaceae
Achromobacter_ruhlandii
Alcaligenes_faecalis
Methylovulum_psychrotolerans
Cardiobacterium_valvarum_F0432
Legionellales
Legionella_pneumophila
Coxiella_burnetii_RSA_493
Chromatiales
Wenzhouxiangella_marina
Rheinheimera_salexigens
Gammaproteobacteria
Vibrionaceae
Salinivibrio_costicola
Vibrio_splendidus
Oceanospirillales
Marinospirillum_minutulum
Alcanivorax_dieselolei
Halomonas_sp_JB380
Xanthomonadaceae
Stenotrophomonas_maltophilia
Xanthomonas
Xanthomonas_alfalfae
Xanthomonas_phaseoli
Xanthomonas_fuscans
Dickeya_dianthicola
Serratia_proteamaculans
Enterobacterales
Providencia_rettgeri
Raoultella_ornithinolytica
Salmonella
Salmonella_enterica_subsp_enterica_serovar_Agona_str_557928
Kluyvera_intermedia
Enterobacteriaceae
Citrobacter_freundii
Klebsiella
Klebsiella_variicola
Klebsiella_quasipneumoniae
Escherichia
Escherichia_coli_UMEA_3052-1
Escherichia_coli_O145_H28_str_2009C-3292
Enterobacter
Enterobacter_cloacae
Enterobacter_roggenkampii
Enterobacter_kobei
Enterobacter_hormaechei

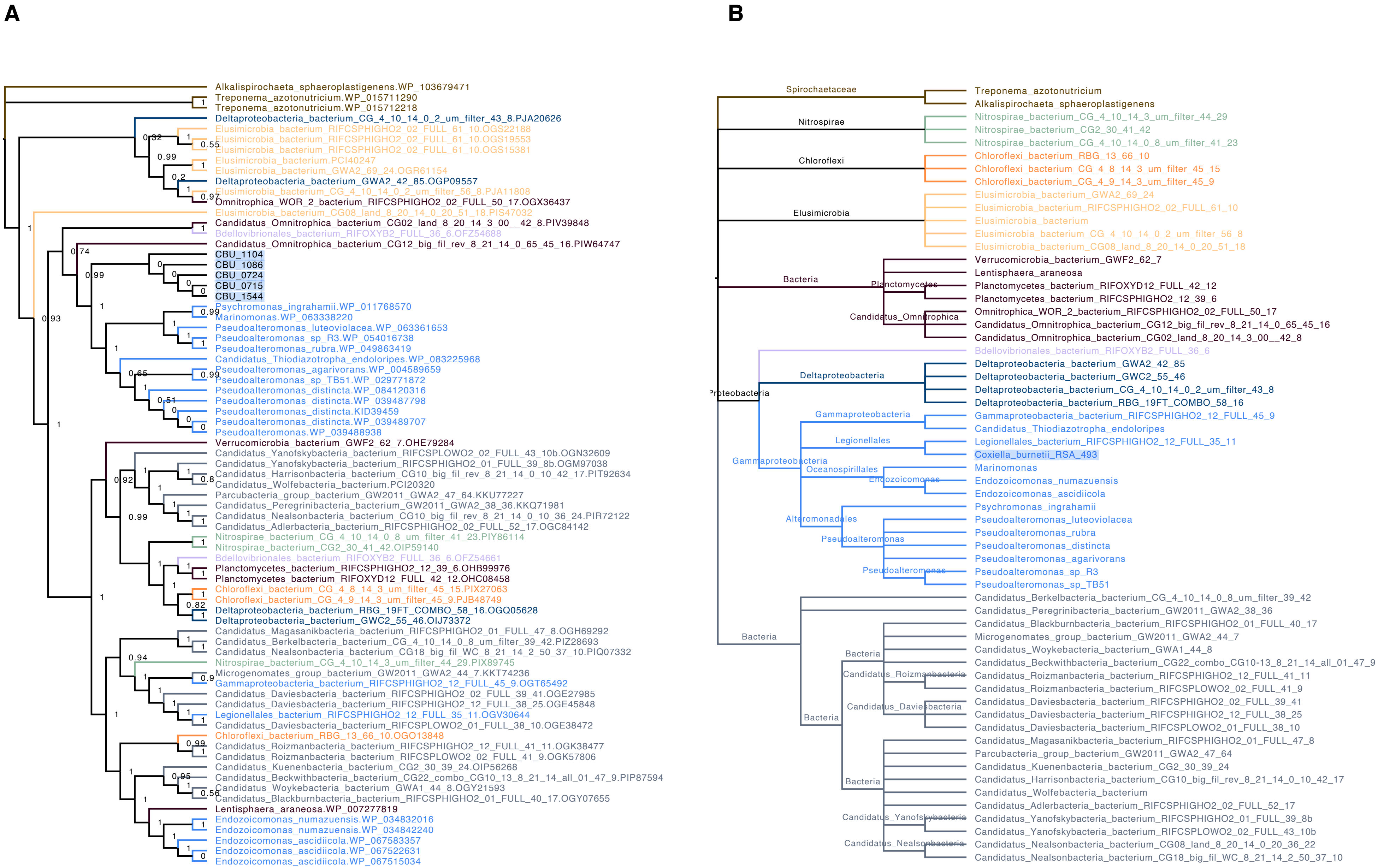
A
Alkalispirochaeta_sphaeroplastigenens.WP_103679471
Treponema_azotonutricium.WP_015711290
Treponema_azotonutricium.WP_015712218
Deltaproteobacteria_bacterium_CG_4_10_14_0_2_um_filter_43_8.PJA20626
Elusimicrobia_bacterium_RIFCSPHIGHO2_02_FULL_61_10.OGS22188
Elusimicrobia_bacterium_RIFCSPHIGHO2_02_FULL_61_10.OGS19553
Elusimicrobia_bacterium_RIFCSPHIGHO2_02_FULL_61_10.OGS15381
Elusimicrobia_bacterium.PCI40247
Elusimicrobia_bacterium_GWA2_69_24.OGR61154
Deltaproteobacteria_bacterium_GWA2_42_85.OGP09557
Elusimicrobia_bacterium_CG_4_10_14_0_2_um_filter_56_8.PJA11808
Omnitrophica_WOR_2_bacterium_RIFCSPHIGHO2_02_FULL_50_17.OGX36437
Elusimicrobia_bacterium_CG08_land_8_20_14_0_20_51_18.PIS47032
Candidatus_Omnitrophica_bacterium_CG02_land_8_20_14_3_00__42_8.PIV39848
Bdellovibrionales_bacterium_RIFOXYB2_FULL_36_6.OFZ54688
Candidatus_Omnitrophica_bacterium_CG12_big_fil_rev_8_21_14_0_65_45_16.PIW64747
CBU_1104
CBU_1086
CBU_0724
CBU_0715
CBU_1544
Psychromonas_ingrahamii.WP_011768570
Marinomonas.WP_063338220
Pseudoalteromonas_luteoviolacea.WP_063361653
Pseudoalteromonas_sp_R3.WP_054016738
Pseudoalteromonas_rubra.WP_049863419
Candidatus_Thiodiazotropha_endoloripes.WP_083225968
Pseudoalteromonas_agarivorans.WP_004589659
Pseudoalteromonas_sp_TB51.WP_029771872
Pseudoalteromonas_distincta.WP_084120316
Pseudoalteromonas_distincta.WP_039487798
Pseudoalteromonas_distincta.KID39459
Pseudoalteromonas_distincta.WP_039489707
Pseudoalteromonas.WP_039488938
Verrucomicrobia_bacterium_GWF2_62_7.OHE79284
Candidatus_Yanofskybacteria_bacterium_RIFCSPLOWO2_02_FULL_43_10b.OGN32609
Candidatus_Yanofskybacteria_bacterium_RIFCSPHIGHO2_01_FULL_39_8b.OGM97038
Candidatus_Harrisonbacteria_bacterium_CG10_big_fil_rev_8_21_14_0_10_42_17.PIT92634
Candidatus_Wolfebacteria_bacterium.PCI20320
Parcubacteria_group_bacterium_GW2011_GWA2_47_64.KKU77227
Candidatus_Peregrinibacteria_bacterium_GW2011_GWA2_38_36.KKQ71981
Candidatus_Nealsonbacteria_bacterium_CG10_big_fil_rev_8_21_14_0_10_36_24.PIR72122
Candidatus_Adlerbacteria_bacterium_RIFCSPHIGHO2_02_FULL_52_17.OGC84142
Nitrospirae_bacterium_CG_4_10_14_0_8_um_filter_41_23.PIY86114
Nitrospirae_bacterium_CG2_30_41_42.OIP59140
Bdellovibrionales_bacterium_RIFOXYB2_FULL_36_6.OFZ54661
Planctomycetes_bacterium_RIFCSPHIGHO2_12_39_6.OHB99976
Planctomycetes_bacterium_RIFOXYD12_FULL_42_12.OHC08458
Chloroflexi_bacterium_CG_4_8_14_3_um_filter_45_15.PIX27063
Chloroflexi_bacterium_CG_4_9_14_3_um_filter_45_9.PJB48749
Deltaproteobacteria_bacterium_RBG_19FT_COMBO_58_16.OGQ05628
Deltaproteobacteria_bacterium_GWC2_55_46.OIJ73372
Candidatus_Magasanikbacteria_bacterium_RIFCSPHIGHO2_01_FULL_47_8.OGH69292
Candidatus_Berkelbacteria_bacterium_CG_4_10_14_0_8_um_filter_39_42.PIZ28693
Candidatus_Nealsonbacteria_bacterium_CG18_big_fil_WC_8_21_14_2_50_37_10.PIQ07332
Nitrospirae_bacterium_CG_4_10_14_3_um_filter_44_29.PIX89745
Microgenomates_group_bacterium_GW2011_GWA2_44_7.KKT74236
Gammaproteobacteria_bacterium_RIFCSPHIGHO2_12_FULL_45_9.OGT65492
Candidatus_Daviesbacteria_bacterium_RIFCSPHIGHO2_02_FULL_39_41.OGE27985
Candidatus_Daviesbacteria_bacterium_RIFCSPHIGHO2_12_FULL_38_25.OGE45848
Legionellales_bacterium_RIFCSPHIGHO2_12_FULL_35_11.OGV30644
Candidatus_Daviesbacteria_bacterium_RIFCSPLOWO2_01_FULL_38_10.OGE38472
Chloroflexi_bacterium_RBG_13_66_10.OGO13848
Candidatus_Roizmanbacteria_bacterium_RIFCSPHIGHO2_12_FULL_41_11.OGK38477
Candidatus_Roizmanbacteria_bacterium_RIFCSPLOWO2_02_FULL_41_9.OGK57806
Candidatus_Kuenenbacteria_bacterium_CG2_30_39_24.OIP56268
Candidatus_Beckwithbacteria_bacterium_CG22_combo_CG10_13_8_21_14_all_01_47_9.PIP87594
Candidatus_Woykebacteria_bacterium_GWA1_44_8.OGY21593
Candidatus_Blackburnbacteria_bacterium_RIFCSPHIGHO2_01_FULL_40_17.OGY07655
Lentisphaera_araneosa.WP_007277819
Endozoicomonas_numazuensis.WP_034832016
Endozoicomonas_numazuensis.WP_034842240
Endozoicomonas_ascidiicola.WP_067583357
Endozoicomonas_ascidiicola.WP_067522631
Endozoicomonas_ascidiicola.WP_067515034
B
Spirochaetaceae
Nitrospirae
Chloroflexi
Elusimicrobia
Bacteria
Planctomycetes
Candidatus_Omnitrophica
Proteobacteria
Deltaproteobacteria
Gammaproteobacteria
Legionellales
Oceanospirillales
Endozoicomonas
Alteromonadales
Pseudoalteromonas
Candidatus_Roizmanbacteria
Candidatus_Daviesbacteria
Candidatus_Yanofskybacteria
Candidatus_Nealsonbacteria
Treponema_azotonutricium
Alkalispirochaeta_sphaeroplastigenens
Nitrospirae_bacterium_CG_4_10_14_3_um_filter_44_29
Nitrospirae_bacterium_CG2_30_41_42
Nitrospirae_bacterium_CG_4_10_14_0_8_um_filter_41_23
Chloroflexi_bacterium_RBG_13_66_10
Chloroflexi_bacterium_CG_4_8_14_3_um_filter_45_15
Chloroflexi_bacterium_CG_4_9_14_3_um_filter_45_9
Elusimicrobia_bacterium_GWA2_69_24
Elusimicrobia_bacterium_RIFCSPHIGHO2_02_FULL_61_10
Elusimicrobia_bacterium
Elusimicrobia_bacterium_CG_4_10_14_0_2_um_filter_56_8
Elusimicrobia_bacterium_CG08_land_8_20_14_0_20_51_18
Verrucomicrobia_bacterium_GWF2_62_7
Lentisphaera_araneosa
Planctomycetes_bacterium_RIFOXYD12_FULL_42_12
Planctomycetes_bacterium_RIFCSPHIGHO2_12_39_6
Omnitrophica_WOR_2_bacterium_RIFCSPHIGHO2_02_FULL_50_17
Candidatus_Omnitrophica_bacterium_CG12_big_fil_rev_8_21_14_0_65_45_16
Candidatus_Omnitrophica_bacterium_CG02_land_8_20_14_3_00__42_8
Bdellovibrionales_bacterium_RIFOXYB2_FULL_36_6
Deltaproteobacteria_bacterium_GWA2_42_85
Deltaproteobacteria_bacterium_GWC2_55_46
Deltaproteobacteria_bacterium_CG_4_10_14_0_2_um_filter_43_8
Deltaproteobacteria_bacterium_RBG_19FT_COMBO_58_16
Gammaproteobacteria_bacterium_RIFCSPHIGHO2_12_FULL_45_9
Candidatus_Thiodiazotropha_endoloripes
Legionellales_bacterium_RIFCSPHIGHO2_12_FULL_35_11
Coxiella_burnetii_RSA_493
Marinomonas
Endozoicomonas_numazuensis
Endozoicomonas_ascidiicola
Psychromonas_ingrahamii
Pseudoalteromonas_luteoviolacea
Pseudoalteromonas_rubra
Pseudoalteromonas_distincta
Pseudoalteromonas_agarivorans
Pseudoalteromonas_sp_R3
Pseudoalteromonas_sp_TB51
Candidatus_Berkelbacteria_bacterium_CG_4_10_14_0_8_um_filter_39_42
Candidatus_Peregrinibacteria_bacterium_GW2011_GWA2_38_36
Candidatus_Blackburnbacteria_bacterium_RIFCSPHIGHO2_01_FULL_40_17
Microgenomates_group_bacterium_GW2011_GWA2_44_7
Candidatus_Woykebacteria_bacterium_GWA1_44_8
Candidatus_Beckwithbacteria_bacterium_CG22_combo_CG10-13_8_21_14_all_01_47_9
Candidatus_Roizmanbacteria_bacterium_RIFCSPHIGHO2_12_FULL_41_11
Candidatus_Roizmanbacteria_bacterium_RIFCSPLOWO2_02_FULL_41_9
Candidatus_Daviesbacteria_bacterium_RIFCSPHIGHO2_02_FULL_39_41
Candidatus_Daviesbacteria_bacterium_RIFCSPHIGHO2_12_FULL_38_25
Candidatus_Daviesbacteria_bacterium_RIFCSPLOWO2_01_FULL_38_10
Candidatus_Magasanikbacteria_bacterium_RIFCSPHIGHO2_01_FULL_47_8
Parcubacteria_group_bacterium_GW2011_GWA2_47_64
Candidatus_Kuenenbacteria_bacterium_CG2_30_39_24
Candidatus_Harrisonbacteria_bacterium_CG10_big_fil_rev_8_21_14_0_10_42_17
Candidatus_Wolfebacteria_bacterium
Candidatus_Adlerbacteria_bacterium_RIFCSPHIGHO2_02_FULL_52_17
Candidatus_Yanofskybacteria_bacterium_RIFCSPHIGHO2_01_FULL_39_8b
Candidatus_Yanofskybacteria_bacterium_RIFCSPLOWO2_02_FULL_43_10b
Candidatus_Nealsonbacteria_bacterium_CG08_land_8_20_14_0_20_36_22
Candidatus_Nealsonbacteria_bacterium_CG18_big_fil_WC_8_21_14_2_50_37_10

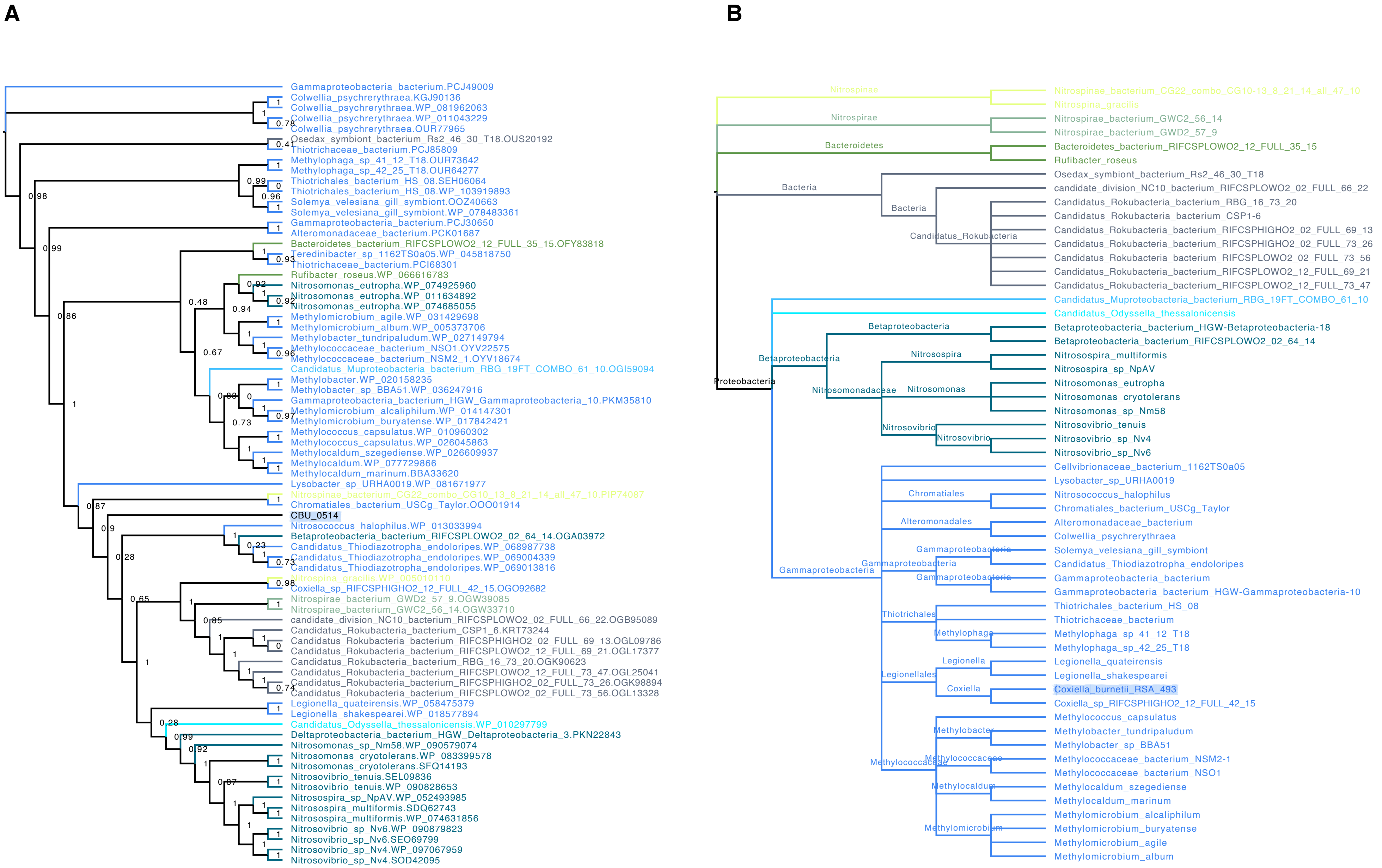
A
Gammaproteobacteria_bacterium.PCJ49009
Colwellia_psychrerythraea.KGJ90136
Colwellia_psychrerythraea.WP_081962063
Colwellia_psychrerythraea.WP_011043229
Colwellia_psychrerythraea.OUR77965
Osedax_symbiont_bacterium_Rs2_46_30_T18.OUS20192
Thiotrichaceae_bacterium.PCJ85809
Methylophaga_sp_41_12_T18.OUR73642
Methylophaga_sp_42_25_T18.OUR64277
Thiotrichales_bacterium_HS_08.SEH06064
Thiotrichales_bacterium_HS_08.WP_103919893
Solemya_velesiana_gill_symbiont.OOZ40663
Solemya_velesiana_gill_symbiont.WP_078483361
Gammaproteobacteria_bacterium.PCJ30650
Alteromonadaceae_bacterium.PCK01687
Bacteroidetes_bacterium_RIFCSPLOWO2_12_FULL_35_15.OFY83818
Teredinibacter_sp_1162TS0a05.WP_045818750
Thiotrichaceae_bacterium.PCI68301
Rufibacter_roseus.WP_066616783
Nitrosomonas_eutropha.WP_074925960
Nitrosomonas_eutropha.WP_011634892
Nitrosomonas_eutropha.WP_074685055
Methylomicrobium_agile.WP_031429698
Methylomicrobium_album.WP_005373706
Methylobacter_tundripaludum.WP_027149794
Methylococcaceae_bacterium_NSO1.OYV22575
Methylococcaceae_bacterium_NSM2_1.OYV18674
Candidatus_Muproteobacteria_bacterium_RBG_19FT_COMBO_61_10.OGI59094
Methylobacter.WP_020158235
Methylobacter_sp_BBA51.WP_036247916
Gammaproteobacteria_bacterium_HGW_Gammaproteobacteria_10.PKM35810
Methylomicrobium_alcaliphilum.WP_014147301
Methylomicrobium_buryatense.WP_017842421
Methylococcus_capsulatus.WP_010960302
Methylococcus_capsulatus.WP_026045863
Methylocaldum_szegediense.WP_026609937
Methylocaldum.WP_077729866
Methylocaldum_marinum.BBA33620
Lysobacter_sp_URHA0019.WP_081671977
Nitrospinae_bacterium_CG22_combo_CG10_13_8_21_14_all_47_10.PIP74087
Chromatiales_bacterium_USCg_Taylor.OOO01914
CBU_0514
Nitrosococcus_halophilus.WP_013033994
Betaproteobacteria_bacterium_RIFCSPLOWO2_02_64_14.OGA03972
Candidatus_Thiodiazotropha_endoloripes.WP_068987738
Candidatus_Thiodiazotropha_endoloripes.WP_069004339
Candidatus_Thiodiazotropha_endoloripes.WP_069013816
Nitrospina_gracilis.WP_005010110
Coxiella_sp_RIFCSPHIGHO2_12_FULL_42_15.OGO92682
Nitrospirae_bacterium_GWD2_57_9.OGW39085
Nitrospirae_bacterium_GWC2_56_14.OGW33710
candidate_division_NC10_bacterium_RIFCSPLOWO2_02_FULL_66_22.OGB95089
Candidatus_Rokubacteria_bacterium_CSP1_6.KRT73244
Candidatus_Rokubacteria_bacterium_RIFCSPHIGHO2_02_FULL_69_13.OGL09786
Candidatus_Rokubacteria_bacterium_RIFCSPLOWO2_12_FULL_69_21.OGL17377
Candidatus_Rokubacteria_bacterium_RBG_16_73_20.OGK90623
Candidatus_Rokubacteria_bacterium_RIFCSPLOWO2_12_FULL_73_47.OGL25041
Candidatus_Rokubacteria_bacterium_RIFCSPHIGHO2_02_FULL_73_26.OGK98894
Candidatus_Rokubacteria_bacterium_RIFCSPLOWO2_02_FULL_73_56.OGL13328
Legionella_quateirensis.WP_058475379
Legionella_shakespearei.WP_018577894
Candidatus_Odyssella_thessalonicensis.WP_010297799
Deltaproteobacteria_bacterium_HGW_Deltaproteobacteria_3.PKN22843
Nitrosomonas_sp_Nm58.WP_090579074
Nitrosomonas_cryotolerans.WP_083399578
Nitrosomonas_cryotolerans.SFQ14193
Nitrosovibrio_tenuis.SEL09836
Nitrosovibrio_tenuis.WP_090828653
Nitrosospira_sp_NpAV.WP_052493985
Nitrosospira_multiformis.SDQ62743
Nitrosospira_multiformis.WP_074631856
Nitrosovibrio_sp_Nv6.WP_090879823
Nitrosovibrio_sp_Nv6.SEO69799
Nitrosovibrio_sp_Nv4.WP_097067959
Nitrosovibrio_sp_Nv4.SOD42095
B
Nitrospinae
Nitrospinae_bacterium_CG22_combo_CG10-13_8_21_14_all_47_10
Nitrospina_gracilis
Nitrospirae
Nitrospirae_bacterium_GWC2_56_14
Nitrospirae_bacterium_GWD2_57_9
Bacteroidetes
Bacteroidetes_bacterium_RIFCSPLOWO2_12_FULL_35_15
Rufibacter_roseus
Bacteria
Osedax_symbiont_bacterium_Rs2_46_30_T18
candidate_division_NC10_bacterium_RIFCSPLOWO2_02_FULL_66_22
Candidatus_Rokubacteria
Candidatus_Rokubacteria_bacterium_RBG_16_73_20
Candidatus_Rokubacteria_bacterium_CSP1-6
Candidatus_Rokubacteria_bacterium_RIFCSPHIGHO2_02_FULL_69_13
Candidatus_Rokubacteria_bacterium_RIFCSPHIGHO2_02_FULL_73_26
Candidatus_Rokubacteria_bacterium_RIFCSPLOWO2_02_FULL_73_56
Candidatus_Rokubacteria_bacterium_RIFCSPLOWO2_12_FULL_69_21
Candidatus_Rokubacteria_bacterium_RIFCSPLOWO2_12_FULL_73_47
Proteobacteria
Candidatus_Muproteobacteria_bacterium_RBG_19FT_COMBO_61_10
Candidatus_Odyssella_thessalonicensis
Betaproteobacteria
Betaproteobacteria_bacterium_HGW-Betaproteobacteria-18
Betaproteobacteria_bacterium_RIFCSPLOWO2_02_64_14
Nitrosomonadaceae
Nitrosospira
Nitrosospira_multiformis
Nitrosospira_sp_NpAV
Nitrosomonas
Nitrosomonas_eutropha
Nitrosomonas_cryotolerans
Nitrosomonas_sp_Nm58
Nitrosovibrio
Nitrosovibrio_tenuis
Nitrosovibrio_sp_Nv4
Nitrosovibrio_sp_Nv6
Gammaproteobacteria
Cellvibrionaceae_bacterium_1162TS0a05
Lysobacter_sp_URHA0019
Chromatiales
Nitrosococcus_halophilus
Chromatiales_bacterium_USCg_Taylor
Alteromonadales
Alteromonadaceae_bacterium
Colwellia_psychrerythraea
Solemya_velesiana_gill_symbiont
Candidatus_Thiodiazotropha_endoloripes
Gammaproteobacteria_bacterium
Gammaproteobacteria_bacterium_HGW-Gammaproteobacteria-10
Thiotrichales
Thiotrichales_bacterium_HS_08
Thiotrichaceae_bacterium
Methylophaga
Methylophaga_sp_41_12_T18
Methylophaga_sp_42_25_T18
Legionellales
Legionella
Legionella_quateirensis
Legionella_shakespearei
Coxiella
Coxiella_burnetii_RSA_493
Coxiella_sp_RIFCSPHIGHO2_12_FULL_42_15
Methylococcaceae
Methylococcus_capsulatus
Methylobacter
Methylobacter_tundripaludum
Methylobacter_sp_BBA51
Methylococcaceae_bacterium_NSM2-1
Methylococcaceae_bacterium_NSO1
Methylocaldum
Methylocaldum_szegediense
Methylocaldum_marinum
Methylomicrobium
Methylomicrobium_alcaliphilum
Methylomicrobium_buryatense
Methylomicrobium_agile
Methylomicrobium_album

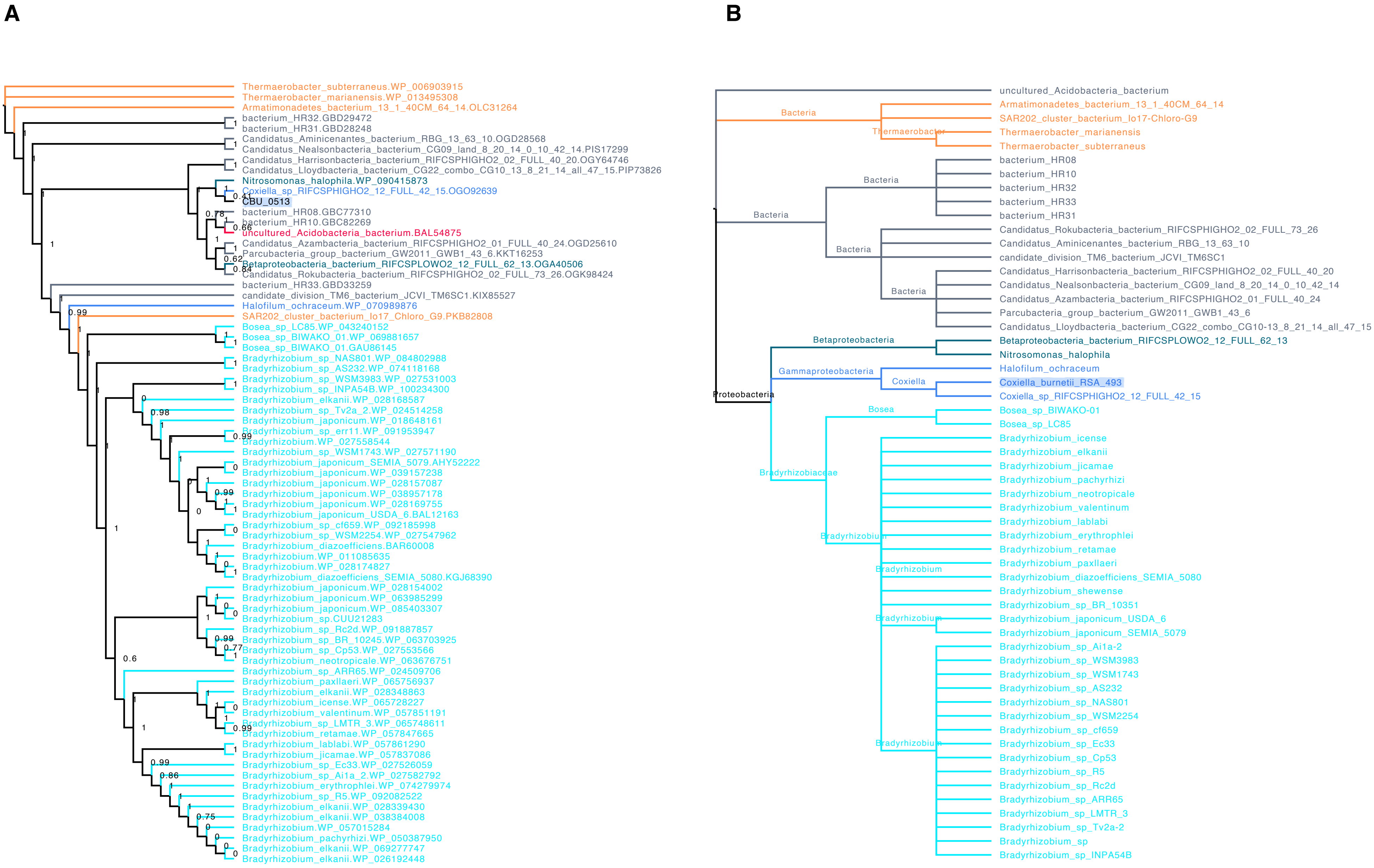
A
Thermaerobacter_subterraneus.WP_006903915
Thermaerobacter_marianensis.WP_013495308
Armatimonadetes_bacterium_13_1_40CM_64_14.OLC31264
bacterium_HR32.GBD29472
bacterium_HR31.GBD28248
Candidatus_Aminicenantes_bacterium_RBG_13_63_10.OGD28568
Candidatus_Nealsonbacteria_bacterium_CG09_land_8_20_14_0_10_42_14.PIS17299
Candidatus_Harrisonbacteria_bacterium_RIFCSPHIGHO2_02_FULL_40_20.OGY64746
Candidatus_Lloydbacteria_bacterium_CG22_combo_CG10_13_8_21_14_all_47_15.PIP73826
Nitrosomonas_halophila.WP_090415873
Coxiella_sp_RIFCSPHIGHO2_12_FULL_42_15.OGO92639
CBU_0513
bacterium_HR08.GBC77310
bacterium_HR10.GBC82269
uncultured_Acidobacteria_bacterium.BAL54875
Candidatus_Azambacteria_bacterium_RIFCSPHIGHO2_01_FULL_40_24.OGD25610
Parcubacteria_group_bacterium_GW2011_GWB1_43_6.KKT16253
Betaproteobacteria_bacterium_RIFCSPLOWO2_12_FULL_62_13.OGA40506
Candidatus_Rokubacteria_bacterium_RIFCSPHIGHO2_02_FULL_73_26.OGK98424
bacterium_HR33.GBD33259
candidate_division_TM6_bacterium_JCVI_TM6SC1.KIX85527
Halofilum_ochraceum.WP_070989876
SAR202_cluster_bacterium_Io17_Chloro_G9.PKB82808
Bosea_sp_LC85.WP_043240152
Bosea_sp_BIWAKO_01.WP_069881657
Bosea_sp_BIWAKO_01.GAU86145
Bradyrhizobium_sp_NAS801.WP_084802988
Bradyrhizobium_sp_AS232.WP_074118168
Bradyrhizobium_sp_WSM3983.WP_027531003
Bradyrhizobium_sp_INPA54B.WP_100234300
Bradyrhizobium_elkanii.WP_028168587
Bradyrhizobium_sp_Tv2a_2.WP_024514258
Bradyrhizobium_japonicum.WP_018648161
Bradyrhizobium_sp_err11.WP_091953947
Bradyrhizobium.WP_027558544
Bradyrhizobium_sp_WSM1743.WP_027571190
Bradyrhizobium_japonicum_SEMIA_5079.AHY52222
Bradyrhizobium_japonicum.WP_039157238
Bradyrhizobium_japonicum.WP_028157087
Bradyrhizobium_japonicum.WP_038957178
Bradyrhizobium_japonicum.WP_028169755
Bradyrhizobium_japonicum_USDA_6.BAL12163
Bradyrhizobium_sp_cf659.WP_092185998
Bradyrhizobium_sp_WSM2254.WP_027547962
Bradyrhizobium_diazoefficiens.BAR60008
Bradyrhizobium.WP_011085635
Bradyrhizobium.WP_028174827
Bradyrhizobium_diazoefficiens_SEMIA_5080.KGJ68390
Bradyrhizobium_japonicum.WP_028154002
Bradyrhizobium_japonicum.WP_063985299
Bradyrhizobium_japonicum.WP_085403307
Bradyrhizobium_sp.CUU21283
Bradyrhizobium_sp_Rc2d.WP_091887857
Bradyrhizobium_sp_BR_10245.WP_063703925
Bradyrhizobium_sp_Cp53.WP_027553566
Bradyrhizobium_neotropicale.WP_063676751
Bradyrhizobium_sp_ARR65.WP_024509706
Bradyrhizobium_paxllaeri.WP_065756937
Bradyrhizobium_elkanii.WP_028348863
Bradyrhizobium_icense.WP_065728227
Bradyrhizobium_valentinum.WP_057851191
Bradyrhizobium_sp_LMTR_3.WP_065748611
Bradyrhizobium_retamae.WP_057847665
Bradyrhizobium_lablabi.WP_057861290
Bradyrhizobium_jicamae.WP_057837086
Bradyrhizobium_sp_Ec33.WP_027526059
Bradyrhizobium_sp_Ai1a_2.WP_027582792
Bradyrhizobium_erythrophlei.WP_074279974
Bradyrhizobium_sp_R5.WP_092082522
Bradyrhizobium_elkanii.WP_028339430
Bradyrhizobium_elkanii.WP_038384008
Bradyrhizobium.WP_057015284
Bradyrhizobium_pachyrhizi.WP_050387950
Bradyrhizobium_elkanii.WP_069277747
Bradyrhizobium_elkanii.WP_026192448
B
uncultured_Acidobacteria_bacterium
Bacteria
Armatimonadetes_bacterium_13_1_40CM_64_14
SAR202_cluster_bacterium_Io17-Chloro-G9
Thermaerobacter
Thermaerobacter_marianensis
Thermaerobacter_subterraneus
Bacteria
bacterium_HR08
bacterium_HR10
bacterium_HR32
bacterium_HR33
bacterium_HR31
Bacteria
Candidatus_Rokubacteria_bacterium_RIFCSPHIGHO2_02_FULL_73_26
Bacteria
Candidatus_Aminicenantes_bacterium_RBG_13_63_10
candidate_division_TM6_bacterium_JCVI_TM6SC1
Candidatus_Harrisonbacteria_bacterium_RIFCSPHIGHO2_02_FULL_40_20
Candidatus_Nealsonbacteria_bacterium_CG09_land_8_20_14_0_10_42_14
Bacteria
Candidatus_Azambacteria_bacterium_RIFCSPHIGHO2_01_FULL_40_24
Parcubacteria_group_bacterium_GW2011_GWB1_43_6
Candidatus_Lloydbacteria_bacterium_CG22_combo_CG10-13_8_21_14_all_47_15
Betaproteobacteria
Betaproteobacteria_bacterium_RIFCSPLOWO2_12_FULL_62_13
Nitrosomonas_halophila
Gammaproteobacteria
Halofilum_ochraceum
Coxiella
Coxiella_burnetii_RSA_493
Proteobacteria
Coxiella_sp_RIFCSPHIGHO2_12_FULL_42_15
Bosea
Bosea_sp_BIWAKO-01
Bosea_sp_LC85
Bradyrhizobium_icense
Bradyrhizobium_elkanii
Bradyrhizobium_jicamae
Bradyrhizobium_pachyrhizi
Bradyrhizobiaceae
Bradyrhizobium_neotropicale
Bradyrhizobium_valentinum
Bradyrhizobium_lablabi
Bradyrhizobium_erythrophlei
Bradyrhizobium_retamae
Bradyrhizobium_paxllaeri
Bradyrhizobium_diazoefficiens_SEMIA_5080
Bradyrhizobium_shewense
Bradyrhizobium_sp_BR_10351
Bradyrhizobium_japonicum_USDA_6
Bradyrhizobium_japonicum_SEMIA_5079
Bradyrhizobium_sp_Ai1a-2
Bradyrhizobium_sp_WSM3983
Bradyrhizobium_sp_WSM1743
Bradyrhizobium_sp_AS232
Bradyrhizobium_sp_NAS801
Bradyrhizobium_sp_WSM2254
Bradyrhizobium_sp_cf659
Bradyrhizobium_sp_Ec33
Bradyrhizobium_sp_Cp53
Bradyrhizobium_sp_R5
Bradyrhizobium_sp_Rc2d
Bradyrhizobium_sp_ARR65
Bradyrhizobium_sp_LMTR_3
Bradyrhizobium_sp_Tv2a-2
Bradyrhizobium_sp
Bradyrhizobium_sp_INPA54B

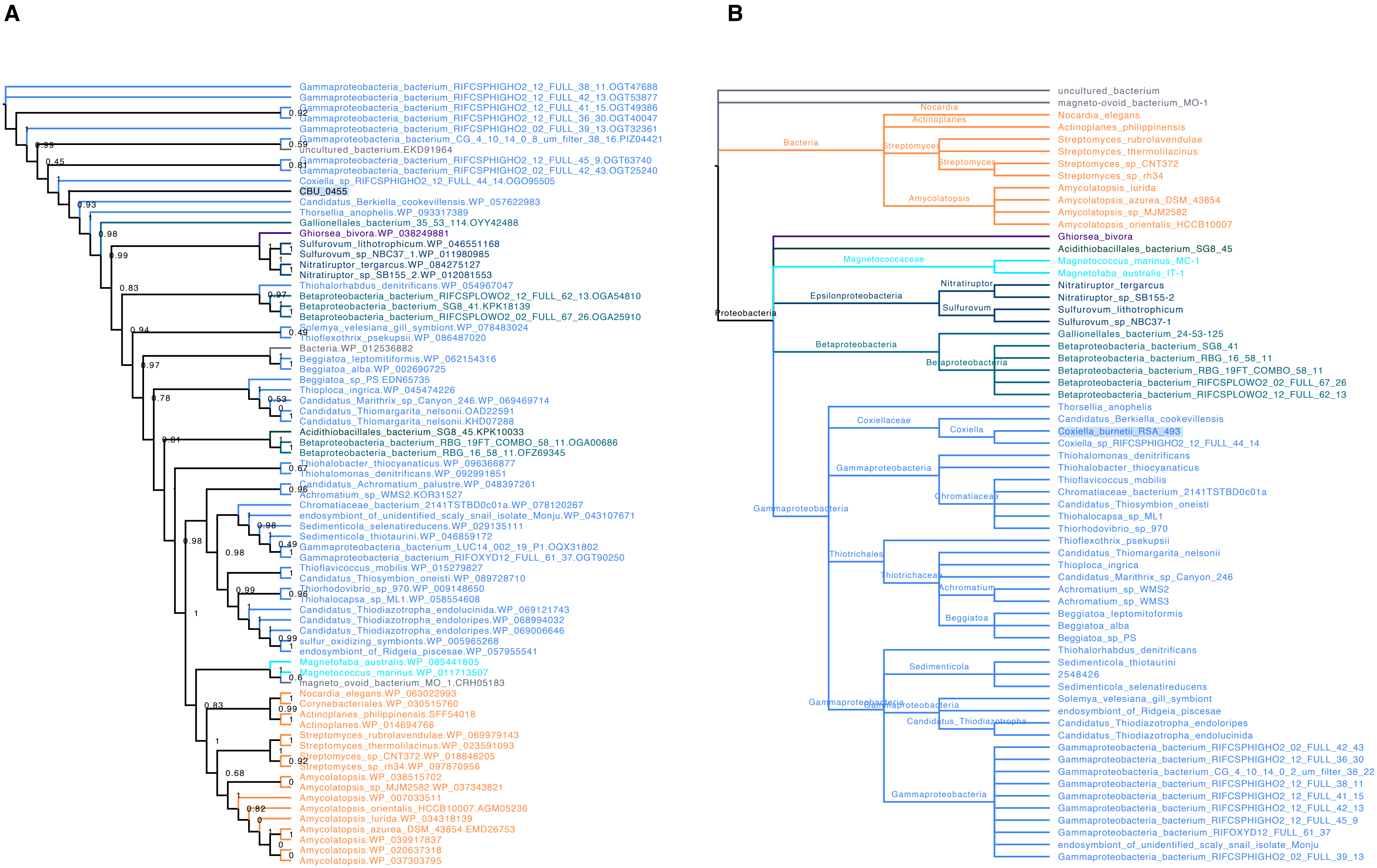
A
Gammaproteobacteria_bacterium_RIFCSPHIGHO2_12_FULL_38_11.OGT47688
Gammaproteobacteria_bacterium_RIFCSPHIGHO2_12_FULL_42_13.OGT53877
Gammaproteobacteria_bacterium_RIFCSPHIGHO2_12_FULL_41_15.OGT49386
Gammaproteobacteria_bacterium_RIFCSPHIGHO2_12_FULL_36_30.OGT40047
Gammaproteobacteria_bacterium_RIFCSPHIGHO2_02_FULL_39_13.OGT32361
Gammaproteobacteria_bacterium_CG_4_10_14_0_8_um_filter_38_16.PIZ04421
uncultured_bacterium.EKD91964
Gammaproteobacteria_bacterium_RIFCSPHIGHO2_12_FULL_45_9.OGT63740
Gammaproteobacteria_bacterium_RIFCSPHIGHO2_02_FULL_42_43.OGT25240
Coxiella_sp_RIFCSPHIGHO2_12_FULL_44_14.OGO95505
CBU_0455
Candidatus_Berkiella_cookevillensis.WP_057622983
Thorsellia_anophelis.WP_093317389
Gallionellales_bacterium_35_53_114.OYY42488
Ghiorsea_bivora.WP_038249881
Sulfurovum_lithotrophicum.WP_046551168
Sulfurovum_sp_NBC37_1.WP_011980985
Nitratiruptor_tergarcus.WP_084275127
Nitratiruptor_sp_SB155_2.WP_012081553
Thiohalorhabdus_denitrificans.WP_054967047
Betaproteobacteria_bacterium_RIFCSPLOWO2_12_FULL_62_13.OGA54810
Betaproteobacteria_bacterium_SG8_41.KPK18139
Betaproteobacteria_bacterium_RIFCSPLOWO2_02_FULL_67_26.OGA25910
Solemya_velesiana_gill_symbiont.WP_078483024
Thioflexothrix_psekupsii.WP_086487020
Bacteria.WP_012536882
Beggiatoa_leptomitiformis.WP_062154316
Beggiatoa_alba.WP_002690725
Beggiatoa_sp_PS.EDN65735
Thioploca_ingrica.WP_045474226
Candidatus_Marithrix_sp_Canyon_246.WP_069469714
Candidatus_Thiomargarita_nelsonii.OAD22591
Candidatus_Thiomargarita_nelsonii.KHD07288
Acidithiobacillales_bacterium_SG8_45.KPK10033
Betaproteobacteria_bacterium_RBG_19FT_COMBO_58_11.OGA00686
Betaproteobacteria_bacterium_RBG_16_58_11.OFZ69345
Thiohalobacter_thiocyanaticus.WP_096366877
Thiohalomonas_denitrificans.WP_092991851
Candidatus_Achromatium_palustre.WP_048397261
Achromatium_sp_WMS2.KOR31527
Chromatiaceae_bacterium_2141TSTBD0c01a.WP_078120267
endosymbiont_of_unidentified_scaly_snail_isolate_Monju.WP_043107671
Sedimenticola_selenatireducens.WP_029135111
Sedimenticola_thiotaurini.WP_046859172
Gammaproteobacteria_bacterium_LUC14_002_19_P1.OQX31802
Gammaproteobacteria_bacterium_RIFOXYD12_FULL_61_37.OGT90250
Thioflavicoccus_mobilis.WP_015279827
Candidatus_Thiosymbion_oneisti.WP_089728710
Thiorhodovibrio_sp_970.WP_009148650
Thiohalocapsa_sp_ML1.WP_058554608
Candidatus_Thiodiazotropha_endolucinida.WP_069121743
Candidatus_Thiodiazotropha_endoloripes.WP_068994032
Candidatus_Thiodiazotropha_endoloripes.WP_069006646
sulfur_oxidizing_symbionts.WP_005965268
endosymbiont_of_Ridgeia_piscesae.WP_057955541
Magnetofaba_australis.WP_085441805
Magnetococcus_marinus.WP_011713507
magneto_ovoid_bacterium_MO_1.CRH05183
Nocardia_elegans.WP_063022993
Corynebacteriales.WP_030515760
Actinoplanes_philippinensis.SFF54018
Actinoplanes.WP_014694766
Streptomyces_rubrolavendulae.WP_069979143
Streptomyces_thermolilacinus.WP_023591093
Streptomyces_sp_CNT372.WP_018848205
Streptomyces_sp_rh34.WP_097870956
Amycolatopsis.WP_038515702
Amycolatopsis_sp_MJM2582.WP_037343821
Amycolatopsis.WP_007033511
Amycolatopsis_orientalis_HCCB10007.AGM05236
Amycolatopsis_lurida.WP_034318139
Amycolatopsis_azurea_DSM_43854.EMD26753
Amycolatopsis.WP_039917837
Amycolatopsis.WP_020637318
Amycolatopsis.WP_037303795
B
uncultured_bacterium
magneto-ovoid_bacterium_MO-1
Bacteria
Nocardia
Actinoplanes
Streptomyces
Amycolatopsis
Nocardia_elegans
Actinoplanes_philippinensis
Streptomyces_rubrolavendulae
Streptomyces_thermolilacinus
Streptomyces_sp_CNT372
Streptomyces_sp_rh34
Amycolatopsis_lurida
Amycolatopsis_azurea_DSM_43854
Amycolatopsis_sp_MJM2582
Amycolatopsis_orientalis_HCCB10007
Proteobacteria
Ghiorsea_bivora
Acidithiobacillales_bacterium_SG8_45
Magnetococcaceae
Magnetococcus_marinus_MC-1
Magnetofaba_australis_IT-1
Epsilonproteobacteria
Nitratiruptor
Sulfurovum
Nitratiruptor_tergarcus
Nitratiruptor_sp_SB155-2
Sulfurovum_lithotrophicum
Sulfurovum_sp_NBC37-1
Betaproteobacteria
Gallionellales_bacterium_24-53-125
Betaproteobacteria_bacterium_SG8_41
Betaproteobacteria_bacterium_RBG_16_58_11
Betaproteobacteria_bacterium_RBG_19FT_COMBO_58_11
Betaproteobacteria_bacterium_RIFCSPLOWO2_02_FULL_67_26
Betaproteobacteria_bacterium_RIFCSPLOWO2_12_FULL_62_13
Gammaproteobacteria
Coxiellaceae
Coxiella
Thorsellia_anophelis
Candidatus_Berkiella_cookevillensis
Coxiella_burnetii_RSA_493
Coxiella_sp_RIFCSPHIGHO2_12_FULL_44_14
Chromatiaceae
Thiohalomonas_denitrificans
Thiohalobacter_thiocyanaticus
Thioflavicoccus_mobilis
Chromatiaceae_bacterium_2141TSTBD0c01a
Candidatus_Thiosymbion_oneisti
Thiohalocapsa_sp_ML1
Thiorhodovibrio_sp_970
Thiotrichales
Thiotrichaceae
Achromatium
Beggiatoa
Thioflexothrix_psekupsii
Candidatus_Thiomargarita_nelsonii
Thioploca_ingrica
Candidatus_Marithrix_sp_Canyon_246
Achromatium_sp_WMS2
Achromatium_sp_WMS3
Beggiatoa_leptomitoformis
Beggiatoa_alba
Beggiatoa_sp_PS
Sedimenticola
Candidatus_Thiodiazotropha
Thiohalorhabdus_denitrificans
Sedimenticola_thiotaurini
2548426
Sedimenticola_selenatireducens
Solemya_velesiana_gill_symbiont
endosymbiont_of_Ridgeia_piscesae
Candidatus_Thiodiazotropha_endoloripes
Candidatus_Thiodiazotropha_endolucinida
Gammaproteobacteria_bacterium_RIFCSPHIGHO2_02_FULL_42_43
Gammaproteobacteria_bacterium_RIFCSPHIGHO2_12_FULL_36_30
Gammaproteobacteria_bacterium_CG_4_10_14_0_2_um_filter_38_22
Gammaproteobacteria_bacterium_RIFCSPHIGHO2_12_FULL_38_11
Gammaproteobacteria_bacterium_RIFCSPHIGHO2_12_FULL_41_15
Gammaproteobacteria_bacterium_RIFCSPHIGHO2_12_FULL_42_13
Gammaproteobacteria_bacterium_RIFCSPHIGHO2_12_FULL_45_9
Gammaproteobacteria_bacterium_RIFOXYD12_FULL_61_37
endosymbiont_of_unidentified_scaly_snail_isolate_Monju
Gammaproteobacteria_bacterium_RIFCSPHIGHO2_02_FULL_39_13

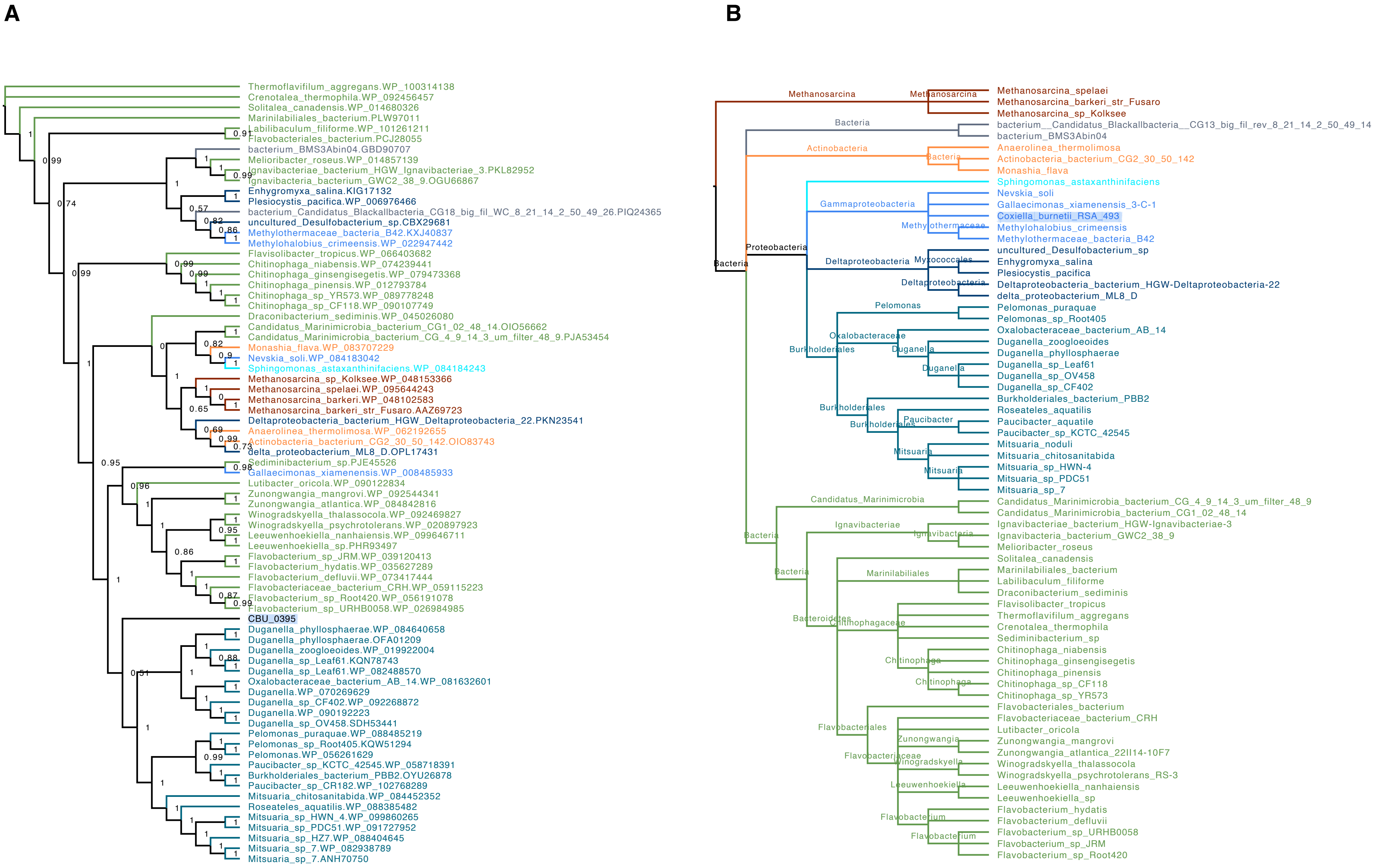
A
Thermoflavifilum_aggregans.WP_100314138
Crenotalea_thermophila.WP_092456457
Solitalea_canadensis.WP_014680326
Marinilabiliales_bacterium.PLW97011
Labilibaculum_filiforme.WP_101261211
Flavobacteriales_bacterium.PCJ28055
bacterium_BMS3Abin04.GBD90707
Melioribacter_roseus.WP_014857139
Ignavibacteriae_bacterium_HGW_Ignavibacteriae_3.PKL82952
Ignavibacteria_bacterium_GWC2_38_9.OGU66867
Enhygromyxa_salina.KIG17132
Plesiocystis_pacifica.WP_006976466
bacterium_Candidatus_Blackallbacteria_CG18_big_fil_WC_8_21_14_2_50_49_26.PIQ24365
uncultured_Desulfobacterium_sp.CBX29681
Methylothermaceae_bacteria_B42.KXJ40837
Methylohalobius_crimeensis.WP_022947442
Flavisolibacter_tropicus.WP_066403682
Chitinophaga_niabensis.WP_074239441
Chitinophaga_ginsengisegetis.WP_079473368
Chitinophaga_pinensis.WP_012793784
Chitinophaga_sp_YR573.WP_089778248
Chitinophaga_sp_CF118.WP_090107749
Draconibacterium_sediminis.WP_045026080
Candidatus_Marinimicrobia_bacterium_CG1_02_48_14.OIO56662
Candidatus_Marinimicrobia_bacterium_CG_4_9_14_3_um_filter_48_9.PJA53454
Monashia_flava.WP_083707229
Nevskia_soli.WP_084183042
Sphingomonas_astaxanthinifaciens.WP_084184243
Methanosarcina_sp_Kolksee.WP_048153366
Methanosarcina_spelaei.WP_095644243
Methanosarcina_barkeri.WP_048102583
Methanosarcina_barkeri_str_Fusaro.AAZ69723
Deltaproteobacteria_bacterium_HGW_Deltaproteobacteria_22.PKN23541
Anaerolinea_thermolimosa.WP_062192655
Actinobacteria_bacterium_CG2_30_50_142.OIO83743
delta_proteobacterium_ML8_D.OPL17431
Sediminibacterium_sp.PJE45526
Gallaecimonas_xiamenensis.WP_008485933
Lutibacter_oricola.WP_090122834
Zunongwangia_mangrovi.WP_092544341
Zunongwangia_atlantica.WP_084842816
Winogradskyella_thalassocola.WP_092469827
Winogradskyella_psychrotolerans.WP_020897923
Leeuwenhoekiella_nanhaiensis.WP_099646711
Leeuwenhoekiella_sp.PHR93497
Flavobacterium_sp_JRM.WP_039120413
Flavobacterium_hydatis.WP_035627289
Flavobacterium_defluvii.WP_073417444
Flavobacteriaceae_bacterium_CRH.WP_059115223
Flavobacterium_sp_Root420.WP_056191078
Flavobacterium_sp_URHB0058.WP_026984985
CBU_0395
Duganella_phyllosphaerae.WP_084640658
Duganella_phyllosphaerae.OFA01209
Duganella_zoogloeoides.WP_019922004
Duganella_sp_Leaf61.KQN78743
Duganella_sp_Leaf61.WP_082488570
Oxalobacteraceae_bacterium_AB_14.WP_081632601
Duganella.WP_070269629
Duganella_sp_CF402.WP_092268872
Duganella.WP_090192223
Duganella_sp_OV458.SDH53441
Pelomonas_puraquae.WP_088485219
Pelomonas_sp_Root405.KQW51294
Pelomonas.WP_056261629
Paucibacter_sp_KCTC_42545.WP_058718391
Burkholderiales_bacterium_PBB2.OYU26878
Paucibacter_sp_CR182.WP_102768289
Mitsuaria_chitosanitabida.WP_084452352
Roseateles_aquatilis.WP_088385482
Mitsuaria_sp_HWN_4.WP_099860265
Mitsuaria_sp_PDC51.WP_091727952
Mitsuaria_sp_HZ7.WP_088404645
Mitsuaria_sp_7.WP_082938789
Mitsuaria_sp_7.ANH70750
B
Methanosarcina_spelaei
Methanosarcina_barkeri_str_Fusaro
Methanosarcina_sp_Kolksee
bacterium__Candidatus_Blackallbacteria__CG13_big_fil_rev_8_21_14_2_50_49_14
bacterium_BMS3Abin04
Anaerolinea_thermolimosa
Actinobacteria_bacterium_CG2_30_50_142
Monashia_flava
Sphingomonas_astaxanthinifaciens
Nevskia_soli
Gallaecimonas_xiamenensis_3-C-1
Coxiella_burnetii_RSA_493
Methylohalobius_crimeensis
Methylothermaceae_bacteria_B42
uncultured_Desulfobacterium_sp
Enhygromyxa_salina
Plesiocystis_pacifica
Deltaproteobacteria_bacterium_HGW-Deltaproteobacteria-22
delta_proteobacterium_ML8_D
Pelomonas_puraquae
Pelomonas_sp_Root405
Oxalobacteraceae_bacterium_AB_14
Duganella_zoogloeoides
Duganella_phyllosphaerae
Duganella_sp_Leaf61
Duganella_sp_OV458
Duganella_sp_CF402
Burkholderiales_bacterium_PBB2
Roseateles_aquatilis
Paucibacter_aquatile
Paucibacter_sp_KCTC_42545
Mitsuaria_noduli
Mitsuaria_chitosanitabida
Mitsuaria_sp_HWN-4
Mitsuaria_sp_PDC51
Mitsuaria_sp_7
Candidatus_Marinimicrobia_bacterium_CG_4_9_14_3_um_filter_48_9
Candidatus_Marinimicrobia_bacterium_CG1_02_48_14
Ignavibacteriae_bacterium_HGW-Ignavibacteriae-3
Ignavibacteria_bacterium_GWC2_38_9
Melioribacter_roseus
Solitalea_canadensis
Marinilabiliales_bacterium
Labilibaculum_filiforme
Draconibacterium_sediminis
Flavisolibacter_tropicus
Thermoflavifilum_aggregans
Crenotalea_thermophila
Sediminibacterium_sp
Chitinophaga_niabensis
Chitinophaga_ginsengisegetis
Chitinophaga_pinensis
Chitinophaga_sp_CF118
Chitinophaga_sp_YR573
Flavobacteriales_bacterium
Flavobacteriaceae_bacterium_CRH
Lutibacter_oricola
Zunongwangia_mangrovi
Zunongwangia_atlantica_22II14-10F7
Winogradskyella_thalassocola
Winogradskyella_psychrotolerans_RS-3
Leeuwenhoekiella_nanhaiensis
Leeuwenhoekiella_sp
Flavobacterium_hydatis
Flavobacterium_defluvii
Flavobacterium_sp_URHB0058
Flavobacterium_sp_JRM
Flavobacterium_sp_Root420
Methanosarcina
Bacteria
Actinobacteria
Proteobacteria
Gammaproteobacteria
Methylothermaceae
Deltaproteobacteria
Myxococcales
Pelomonas
Oxalobacteraceae
Burkholderiales
Duganella
Paucibacter
Mitsuaria
Candidatus_Marinimicrobia
Ignavibacteriae
Ignavibacteria
Marinilabiliales
Bacteroidetes
Chitinophagaceae
Chitinophaga
Flavobacteriales
Zunongwangia
Flavobacteriaceae
Winogradskyella
Leeuwenhoekiella
Flavobacterium

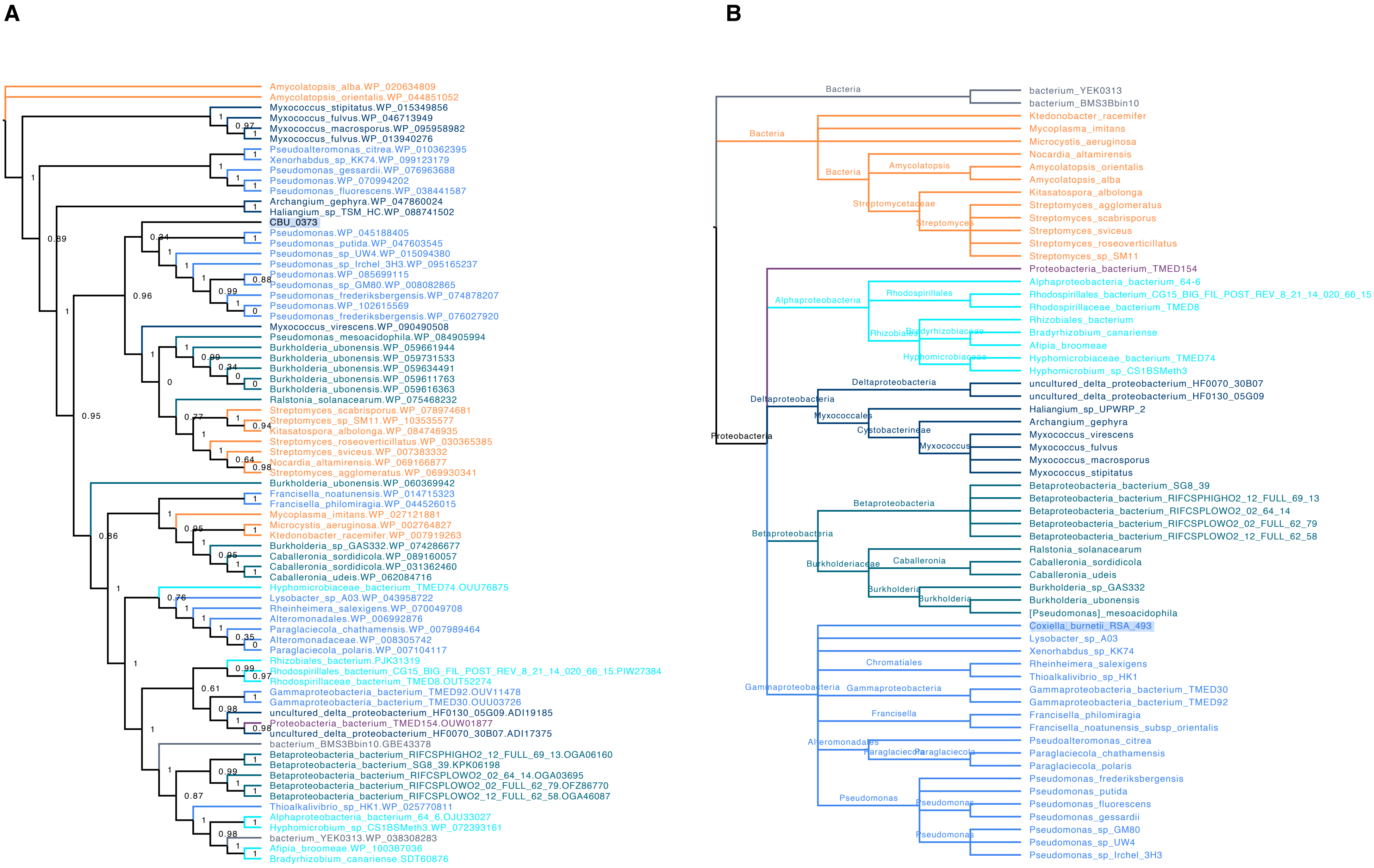

A
Amycolatopsis_alba.WP_020634809
Amycolatopsis_orientalis.WP_044851052
Myxococcus_stipitatus.WP_015349856
Myxococcus_fulvus.WP_046713949
Myxococcus_macrosporus.WP_095958982
Myxococcus_fulvus.WP_013940276
Pseudoalteromonas_citrea.WP_010362395
Xenorhabdus_sp_KK74.WP_099123179
Pseudomonas_gessardii.WP_076963688
Pseudomonas.WP_070994202
Pseudomonas_fluorescens.WP_038441587
Archangium_gephyra.WP_047860024
Haliangium_sp_TSM_HC.WP_088741502
CBU_0373
Pseudomonas.WP_045188405
Pseudomonas_putida.WP_047603545
Pseudomonas_sp_UW4.WP_015094380
Pseudomonas_sp_Irchel_3H3.WP_095165237
Pseudomonas.WP_085699115
Pseudomonas_sp_GM80.WP_008082865
Pseudomonas_frederiksbergensis.WP_074878207
Pseudomonas.WP_102615569
Pseudomonas_frederiksbergensis.WP_076027920
Myxococcus_virescens.WP_090490508
Pseudomonas_mesoacidophila.WP_084905994
Burkholderia_ubonensis.WP_059661944
Burkholderia_ubonensis.WP_059731533
Burkholderia_ubonensis.WP_059634491
Burkholderia_ubonensis.WP_059611763
Burkholderia_ubonensis.WP_059616363
Ralstonia_solanacearum.WP_075468232
Streptomyces_scabrisporus.WP_078974681
Streptomyces_sp_SM11.WP_103535577
Kitasatospora_albolonga.WP_084746935
Streptomyces_roseoverticillatus.WP_030365385
Streptomyces_sviceus.WP_007383332
Nocardia_altamirensis.WP_069166877
Streptomyces_agglomeratus.WP_069930341
Burkholderia_ubonensis.WP_060369942
Francisella_noatunensis.WP_014715323
Francisella_philomiragia.WP_044526015
Mycoplasma_imitans.WP_027121881
Microcystis_aeruginosa.WP_002764827
Ktedonobacter_racemifer.WP_007919263
Burkholderia_sp_GAS332.WP_074286677
Caballeronia_sordidicola.WP_089160057
Caballeronia_sordidicola.WP_031362460
Caballeronia_udeis.WP_062084716
Hyphomicrobiaceae_bacterium_TMED74.OUU76875
Lysobacter_sp_A03.WP_043958722
Rheinheimera_salexigens.WP_070049708
Alteromonadales.WP_006992876
Paraglaciecola_chathamensis.WP_007989464
Alteromonadaceae.WP_008305742
Paraglaciecola_polaris.WP_007104117
Rhizobiales_bacterium.PJK31319
Rhodospirillales_bacterium_CG15_BIG_FIL_POST_REV_8_21_14_020_66_15.PIW27384
Rhodospirillaceae_bacterium_TMED8.OUT52274
Gammaproteobacteria_bacterium_TMED92.OUV11478
Gammaproteobacteria_bacterium_TMED30.OUU03726
uncultured_delta_proteobacterium_HF0130_05G09.ADI19185
Proteobacteria_bacterium_TMED154.OUW01877
uncultured_delta_proteobacterium_HF0070_30B07.ADI17375
bacterium_BMS3Bbin10.GBE43378
Betaproteobacteria_bacterium_RIFCSPHIGHO2_12_FULL_69_13.OGA06160
Betaproteobacteria_bacterium_SG8_39.KPK06198
Betaproteobacteria_bacterium_RIFCSPLOWO2_02_64_14.OGA03695
Betaproteobacteria_bacterium_RIFCSPLOWO2_02_FULL_62_79.OFZ86770
Betaproteobacteria_bacterium_RIFCSPLOWO2_12_FULL_62_58.OGA46087
Thioalkalivibrio_sp_HK1.WP_025770811
Alphaproteobacteria_bacterium_64_6.OJU33027
Hyphomicrobium_sp_CS1BSMeth3.WP_072393161
bacterium_YEK0313.WP_038308283
Afipia_broomeae.WP_100387036
Bradyrhizobium_canariense.SDT60876
B
Bacteria
bacterium_YEK0313
bacterium_BMS3Bbin10
Bacteria
Ktedonobacter_racemifer
Mycoplasma_imitans
Microcystis_aeruginosa
Bacteria
Amycolatopsis
Nocardia_altamirensis
Amycolatopsis_orientalis
Amycolatopsis_alba
Streptomycetaceae
Kitasatospora_albolonga
Streptomyces
Streptomyces_agglomeratus
Streptomyces_scabrisporus
Streptomyces_sviceus
Streptomyces_roseoverticillatus
Streptomyces_sp_SM11
Proteobacteria_bacterium_TMED154
Alphaproteobacteria
Rhodospirillales
Alphaproteobacteria_bacterium_64-6
Rhodospirillales_bacterium_CG15_BIG_FIL_POST_REV_8_21_14_020_66_15
Rhodospirillaceae_bacterium_TMED8
Rhizobiales
Bradyrhizobiaceae
Rhizobiales_bacterium
Bradyrhizobium_canariense
Afipia_broomeae
Hyphomicrobiaceae
Hyphomicrobiaceae_bacterium_TMED74
Hyphomicrobium_sp_CS1BSMeth3
Proteobacteria
Deltaproteobacteria
Deltaproteobacteria
uncultured_delta_proteobacterium_HF0070_30B07
uncultured_delta_proteobacterium_HF0130_05G09
Myxococcales
Haliangium_sp_UPWRP_2
Cystobacterineae
Archangium_gephyra
Myxococcus
Myxococcus_virescens
Myxococcus_fulvus
Myxococcus_macrosporus
Myxococcus_stipitatus
Betaproteobacteria
Betaproteobacteria
Betaproteobacteria_bacterium_SG8_39
Betaproteobacteria_bacterium_RIFCSPHIGHO2_12_FULL_69_13
Betaproteobacteria_bacterium_RIFCSPLOWO2_02_64_14
Betaproteobacteria_bacterium_RIFCSPLOWO2_02_FULL_62_79
Betaproteobacteria_bacterium_RIFCSPLOWO2_12_FULL_62_58
Burkholderiaceae
Ralstonia_solanacearum
Caballeronia
Caballeronia_sordidicola
Caballeronia_udeis
Burkholderia
Burkholderia
Burkholderia_sp_GAS332
Burkholderia_ubonensis
[Pseudomonas]_mesoacidophila
Gammaproteobacteria
Coxiella_burnetii_RSA_493
Lysobacter_sp_A03
Xenorhabdus_sp_KK74
Chromatiales
Rheinheimera_salexigens
Thioalkalivibrio_sp_HK1
Gammaproteobacteria
Gammaproteobacteria_bacterium_TMED30
Gammaproteobacteria_bacterium_TMED92
Francisella
Francisella_philomiragia
Francisella_noatunensis_subsp_orientalis
Alteromonadales
Pseudoalteromonas_citrea
Paraglaciecola
Paraglaciecola
Paraglaciecola_chathamensis
Paraglaciecola_polaris
Pseudomonas
Pseudomonas
Pseudomonas_frederiksbergensis
Pseudomonas_putida
Pseudomonas_fluorescens
Pseudomonas_gessardii
Pseudomonas
Pseudomonas_sp_GM80
Pseudomonas_sp_UW4
Pseudomonas_sp_Irchel_3H3

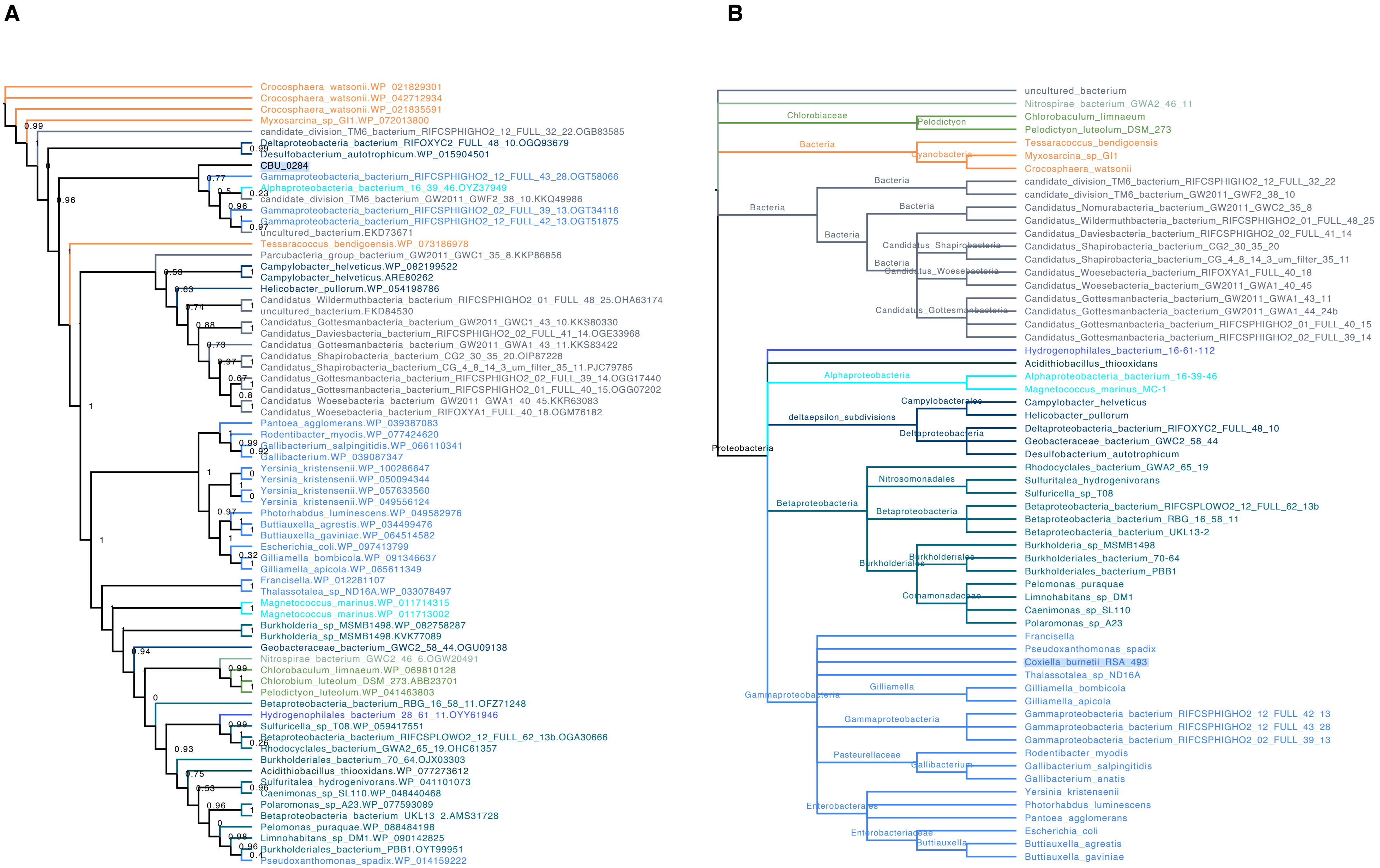
A
Crocosphaera_watsonii.WP_021829301
Crocosphaera_watsonii.WP_042712934
Crocosphaera_watsonii.WP_021835591
Myxosarcina_sp_GI1.WP_072013800
candidate_division_TM6_bacterium_RIFCSPHIGHO2_12_FULL_32_22.OGB83585
Deltaproteobacteria_bacterium_RIFOXYC2_FULL_48_10.OGQ93679
Desulfobacterium_autotrophicum.WP_015904501
CBU_0284
Gammaproteobacteria_bacterium_RIFCSPHIGHO2_12_FULL_43_28.OGT58066
Alphaproteobacteria_bacterium_16_39_46.OYZ37949
candidate_division_TM6_bacterium_GW2011_GWF2_38_10.KKQ49986
Gammaproteobacteria_bacterium_RIFCSPHIGHO2_02_FULL_39_13.OGT34116
Gammaproteobacteria_bacterium_RIFCSPHIGHO2_12_FULL_42_13.OGT51875
uncultured_bacterium.EKD73671
Tessaracoccus_bendigoensis.WP_073186978
Parcubacteria_group_bacterium_GW2011_GWC1_35_8.KKP86856
Campylobacter_helveticus.WP_082199522
Campylobacter_helveticus.ARE80262
Helicobacter_pullorum.WP_054198786
Candidatus_Wildermuthbacteria_bacterium_RIFCSPHIGHO2_01_FULL_48_25.OHA63174
uncultured_bacterium.EKD84530
Candidatus_Gottesmanbacteria_bacterium_GW2011_GWC1_43_10.KKS80330
Candidatus_Daviesbacteria_bacterium_RIFCSPHIGHO2_02_FULL_41_14.OGE33968
Candidatus_Gottesmanbacteria_bacterium_GW2011_GWA1_43_11.KKS83422
Candidatus_Shapirobacteria_bacterium_CG2_30_35_20.OIP87228
Candidatus_Shapirobacteria_bacterium_CG_4_8_14_3_um_filter_35_11.PJC79785
Candidatus_Gottesmanbacteria_bacterium_RIFCSPHIGHO2_02_FULL_39_14.OGG17440
Candidatus_Gottesmanbacteria_bacterium_RIFCSPHIGHO2_01_FULL_40_15.OGG07202
Candidatus_Woesebacteria_bacterium_GW2011_GWA1_40_45.KKR63083
Candidatus_Woesebacteria_bacterium_RIFOXYA1_FULL_40_18.OGM76182
Pantoea_agglomerans.WP_039387083
Rodentibacter_myodis.WP_077424620
Gallibacterium_salpingitidis.WP_066110341
Gallibacterium.WP_039087347
Yersinia_kristensenii.WP_100286647
Yersinia_kristensenii.WP_050094344
Yersinia_kristensenii.WP_057633560
Yersinia_kristensenii.WP_049556124
Photorhabdus_luminescens.WP_049582976
Buttiauxella_agrestis.WP_034499476
Buttiauxella_gaviniae.WP_064514582
Escherichia_coli.WP_097413799
Gilliamella_bombicola.WP_091346637
Gilliamella_apicola.WP_065611349
Francisella.WP_012281107
Thalassotalea_sp_ND16A.WP_033078497
Magnetococcus_marinus.WP_011714315
Magnetococcus_marinus.WP_011713002
Burkholderia_sp_MSMB1498.WP_082758287
Burkholderia_sp_MSMB1498.KVK77089
Geobacteraceae_bacterium_GWC2_58_44.OGU09138
Nitrospirae_bacterium_GWC2_46_6.OGW20491
Chlorobaculum_limnaeum.WP_069810128
Chlorobium_luteolum_DSM_273.ABB23701
Pelodictyon_luteolum.WP_041463803
Betaproteobacteria_bacterium_RBG_16_58_11.OFZ71248
Hydrogenophilales_bacterium_28_61_11.OYY61946
Sulfuricella_sp_T08.WP_059417551
Betaproteobacteria_bacterium_RIFCSPLOWO2_12_FULL_62_13b.OGA30666
Rhodocyclales_bacterium_GWA2_65_19.OHC61357
Burkholderiales_bacterium_70_64.OJX03303
Acidithiobacillus_thiooxidans.WP_077273612
Sulfuritalea_hydrogenivorans.WP_041101073
Caenimonas_sp_SL110.WP_048440468
Polaromonas_sp_A23.WP_077593089
Betaproteobacteria_bacterium_UKL13_2.AMS31728
Pelomonas_puraquae.WP_088484198
Limnohabitans_sp_DM1.WP_090142825
Burkholderiales_bacterium_PBB1.OYT99951
Pseudoxanthomonas_spadix.WP_014159222
B
uncultured_bacterium
Nitrospirae_bacterium_GWA2_46_11
Chlorobiaceae
Pelodictyon
Chlorobaculum_limnaeum
Pelodictyon_luteolum_DSM_273
Bacteria
Cyanobacteria
Tessaracoccus_bendigoensis
Myxosarcina_sp_GI1
Crocosphaera_watsonii
candidate_division_TM6_bacterium_RIFCSPHIGHO2_12_FULL_32_22
candidate_division_TM6_bacterium_GW2011_GWF2_38_10
Candidatus_Nomurabacteria_bacterium_GW2011_GWC2_35_8
Candidatus_Wildermuthbacteria_bacterium_RIFCSPHIGHO2_01_FULL_48_25
Candidatus_Daviesbacteria_bacterium_RIFCSPHIGHO2_02_FULL_41_14
Candidatus_Shapirobacteria
Candidatus_Shapirobacteria_bacterium_CG2_30_35_20
Candidatus_Shapirobacteria_bacterium_CG_4_8_14_3_um_filter_35_11
Candidatus_Woesebacteria
Candidatus_Woesebacteria_bacterium_RIFOXYA1_FULL_40_18
Candidatus_Woesebacteria_bacterium_GW2011_GWA1_40_45
Candidatus_Gottesmanbacteria
Candidatus_Gottesmanbacteria_bacterium_GW2011_GWA1_43_11
Candidatus_Gottesmanbacteria_bacterium_GW2011_GWA1_44_24b
Candidatus_Gottesmanbacteria_bacterium_RIFCSPHIGHO2_01_FULL_40_15
Candidatus_Gottesmanbacteria_bacterium_RIFCSPHIGHO2_02_FULL_39_14
Proteobacteria
Hydrogenophilales_bacterium_16-61-112
Acidithiobacillus_thiooxidans
Alphaproteobacteria
Alphaproteobacteria_bacterium_16-39-46
Magnetococcus_marinus_MC-1
deltaepsilon_subdivisions
Campylobacterales
Campylobacter_helveticus
Helicobacter_pullorum
Deltaproteobacteria
Deltaproteobacteria_bacterium_RIFOXYC2_FULL_48_10
Geobacteraceae_bacterium_GWC2_58_44
Desulfobacterium_autotrophicum
Betaproteobacteria
Rhodocyclales_bacterium_GWA2_65_19
Nitrosomonadales
Sulfuritalea_hydrogenivorans
Sulfuricella_sp_T08
Betaproteobacteria_bacterium_RIFCSPLOWO2_12_FULL_62_13b
Betaproteobacteria_bacterium_RBG_16_58_11
Betaproteobacteria_bacterium_UKL13-2
Burkholderiales
Burkholderia_sp_MSMB1498
Burkholderiales_bacterium_70-64
Burkholderiales_bacterium_PBB1
Comamonadaceae
Pelomonas_puraquae
Limnohabitans_sp_DM1
Caenimonas_sp_SL110
Polaromonas_sp_A23
Gammaproteobacteria
Francisella
Pseudoxanthomonas_spadix
Coxiella_burnetii_RSA_493
Thalassotalea_sp_ND16A
Gilliamella
Gilliamella_bombicola
Gilliamella_apicola
Gammaproteobacteria_bacterium_RIFCSPHIGHO2_12_FULL_42_13
Gammaproteobacteria_bacterium_RIFCSPHIGHO2_12_FULL_43_28
Gammaproteobacteria_bacterium_RIFCSPHIGHO2_02_FULL_39_13
Pasteurellaceae
Gallibacterium
Rodentibacter_myodis
Gallibacterium_salpingitidis
Gallibacterium_anatis
Enterobacterales
Enterobacteriaceae
Buttiauxella
Yersinia_kristensenii
Photorhabdus_luminescens
Pantoea_agglomerans
Escherichia_coli
Buttiauxella_agrestis
Buttiauxella_gaviniae

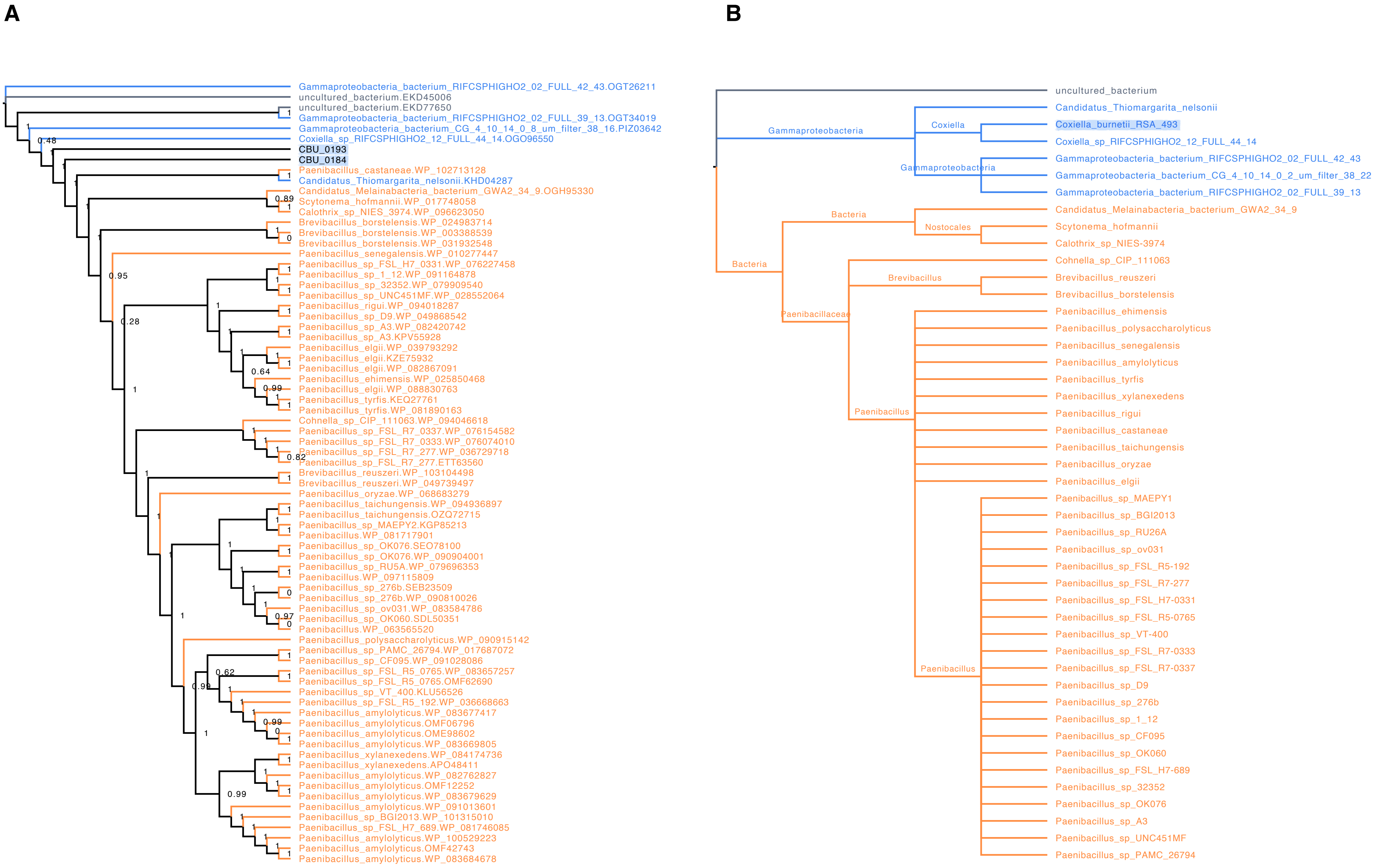
A
Gammaproteobacteria_bacterium_RIFCSPHIGHO2_02_FULL_42_43.OGT26211
uncultured_bacterium.EKD45006
uncultured_bacterium.EKD77650
Gammaproteobacteria_bacterium_RIFCSPHIGHO2_02_FULL_39_13.OGT34019
Gammaproteobacteria_bacterium_CG_4_10_14_0_8_um_filter_38_16.PIZ03642
Coxiella_sp_RIFCSPHIGHO2_12_FULL_44_14.OGO96550
CBU_0193
CBU_0184
Paenibacillus_castaneae.WP_102713128
Candidatus_Thiomargarita_nelsonii.KHD04287
Candidatus_Melainabacteria_bacterium_GWA2_34_9.OGH95330
Scytonema_hofmannii.WP_017748058
Calothrix_sp_NIES_3974.WP_096623050
Brevibacillus_borstelensis.WP_024983714
Brevibacillus_borstelensis.WP_003388539
Brevibacillus_borstelensis.WP_031932548
Paenibacillus_senegalensis.WP_010277447
Paenibacillus_sp_FSL_H7_0331.WP_076227458
Paenibacillus_sp_1_12.WP_091164878
Paenibacillus_sp_32352.WP_079909540
Paenibacillus_sp_UNC451MF.WP_028552064
Paenibacillus_rigui.WP_094018287
Paenibacillus_sp_D9.WP_049868542
Paenibacillus_sp_A3.WP_082420742
Paenibacillus_sp_A3.KPV55928
Paenibacillus_elgii.WP_039793292
Paenibacillus_elgii.KZE75932
Paenibacillus_elgii.WP_082867091
Paenibacillus_ehimensis.WP_025850468
Paenibacillus_elgii.WP_088830763
Paenibacillus_tyrfis.KEQ27761
Paenibacillus_tyrfis.WP_081890163
Cohnella_sp_CIP_111063.WP_094046618
Paenibacillus_sp_FSL_R7_0337.WP_076154582
Paenibacillus_sp_FSL_R7_0333.WP_076074010
Paenibacillus_sp_FSL_R7_277.WP_036729718
Paenibacillus_sp_FSL_R7_277.ETT63560
Brevibacillus_reuszeri.WP_103104498
Brevibacillus_reuszeri.WP_049739497
Paenibacillus_oryzae.WP_068683279
Paenibacillus_taichungensis.WP_094936897
Paenibacillus_taichungensis.OZQ72715
Paenibacillus_sp_MAEPY2.KGP85213
Paenibacillus.WP_081717901
Paenibacillus_sp_OK076.SEO78100
Paenibacillus_sp_OK076.WP_090904001
Paenibacillus_sp_RU5A.WP_079696353
Paenibacillus.WP_097115809
Paenibacillus_sp_276b.SEB23509
Paenibacillus_sp_276b.WP_090810026
Paenibacillus_sp_ov031.WP_083584786
Paenibacillus_sp_OK060.SDL50351
Paenibacillus.WP_063565520
Paenibacillus_polysaccharolyticus.WP_090915142
Paenibacillus_sp_PAMC_26794.WP_017687072
Paenibacillus_sp_CF095.WP_091028086
Paenibacillus_sp_FSL_R5_0765.WP_083657257
Paenibacillus_sp_FSL_R5_0765.OMF62690
Paenibacillus_sp_VT_400.KLU56526
Paenibacillus_sp_FSL_R5_192.WP_036668663
Paenibacillus_amylolyticus.WP_083677417
Paenibacillus_amylolyticus.OMF06796
Paenibacillus_amylolyticus.OME98602
Paenibacillus_amylolyticus.WP_083669805
Paenibacillus_xylanexedens.WP_084174736
Paenibacillus_xylanexedens.APO48411
Paenibacillus_amylolyticus.WP_082762827
Paenibacillus_amylolyticus.OMF12252
Paenibacillus_amylolyticus.WP_083679629
Paenibacillus_amylolyticus.WP_091013601
Paenibacillus_sp_BGI2013.WP_101315010
Paenibacillus_sp_FSL_H7_689.WP_081746085
Paenibacillus_amylolyticus.WP_100529223
Paenibacillus_amylolyticus.OMF42743
Paenibacillus_amylolyticus.WP_083684678
B
uncultured_bacterium
Gammaproteobacteria
Candidatus_Thiomargarita_nelsonii
Coxiella
Coxiella_burnetii_RSA_493
Coxiella_sp_RIFCSPHIGHO2_12_FULL_44_14
Gammaproteobacteria
Gammaproteobacteria_bacterium_RIFCSPHIGHO2_02_FULL_42_43
Gammaproteobacteria_bacterium_CG_4_10_14_0_2_um_filter_38_22
Gammaproteobacteria_bacterium_RIFCSPHIGHO2_02_FULL_39_13
Bacteria
Candidatus_Melainabacteria_bacterium_GWA2_34_9
Nostocales
Scytonema_hofmannii
Calothrix_sp_NIES-3974
Bacteria
Cohnella_sp_CIP_111063
Brevibacillus
Brevibacillus_reuszeri
Brevibacillus_borstelensis
Paenibacillaceae
Paenibacillus_ehimensis
Paenibacillus_polysaccharolyticus
Paenibacillus_senegalensis
Paenibacillus_amylolyticus
Paenibacillus_tyrfis
Paenibacillus_xylanexedens
Paenibacillus
Paenibacillus_rigui
Paenibacillus_castaneae
Paenibacillus_taichungensis
Paenibacillus_oryzae
Paenibacillus_elgii
Paenibacillus_sp_MAEPY1
Paenibacillus_sp_BGI2013
Paenibacillus_sp_RU26A
Paenibacillus_sp_ov031
Paenibacillus_sp_FSL_R5-192
Paenibacillus_sp_FSL_R7-277
Paenibacillus_sp_FSL_H7-0331
Paenibacillus_sp_FSL_R5-0765
Paenibacillus_sp_VT-400
Paenibacillus_sp_FSL_R7-0333
Paenibacillus
Paenibacillus_sp_FSL_R7-0337
Paenibacillus_sp_D9
Paenibacillus_sp_276b
Paenibacillus_sp_1_12
Paenibacillus_sp_CF095
Paenibacillus_sp_OK060
Paenibacillus_sp_FSL_H7-689
Paenibacillus_sp_32352
Paenibacillus_sp_OK076
Paenibacillus_sp_A3
Paenibacillus_sp_UNC451MF
Paenibacillus_sp_PAMC_26794

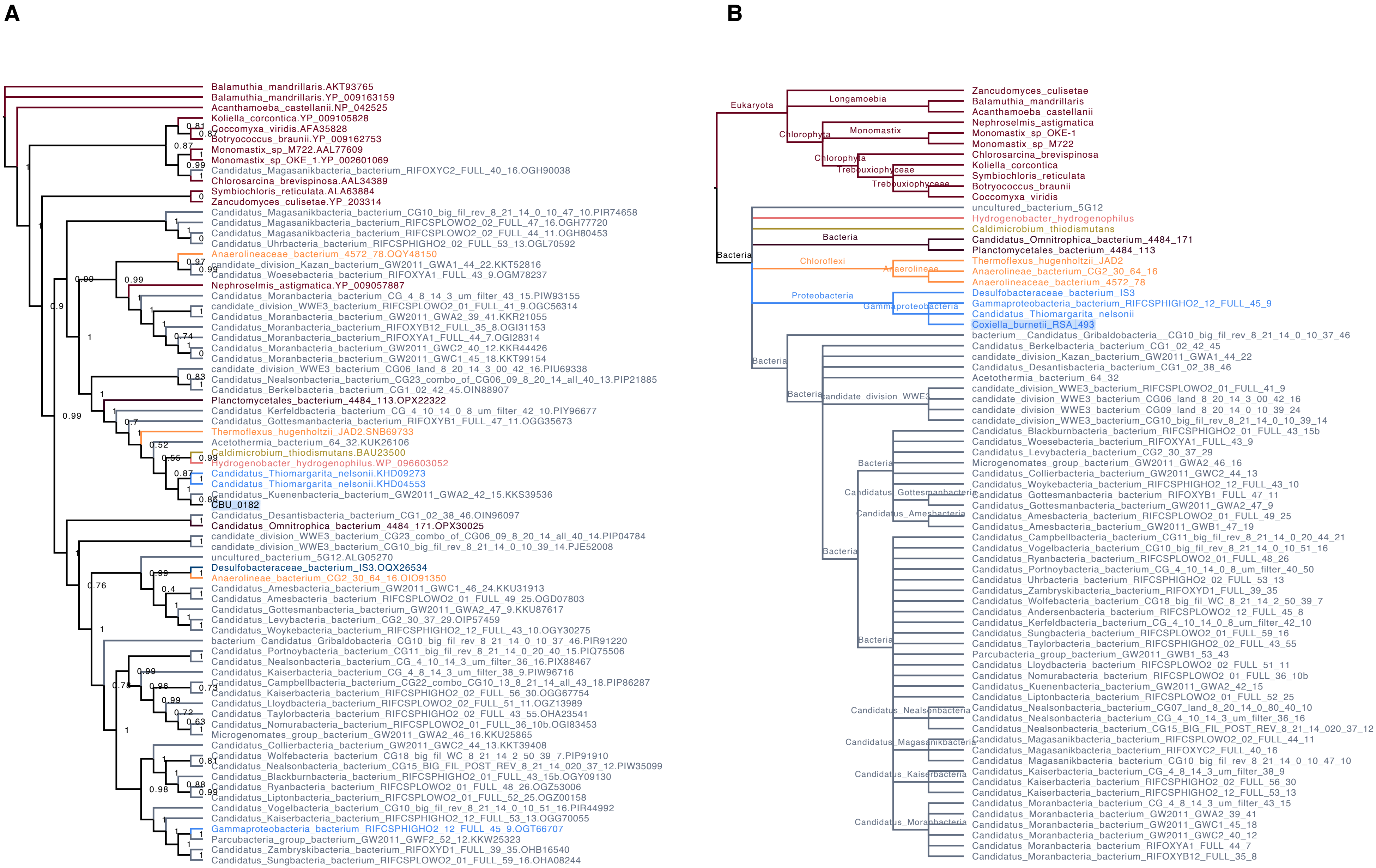
A
Balamuthia_mandrillaris.AKT93765
Balamuthia_mandrillaris.YP_009163159
Acanthamoeba_castellanii.NP_042525
Koliella_corcontica.YP_009105828
Coccomyxa_viridis.AFA35828
Botryococcus_braunii.YP_009162753
Monomastix_sp_M722.AAL77609
Monomastix_sp_OKE_1.YP_002601069
Candidatus_Magasanikbacteria_bacterium_RIFOXYC2_FULL_40_16.OGH90038
Chlorosarcina_brevispinosa.AAL34389
Symbiochloris_reticulata.ALA63884
Zancudomyces_culisetae.YP_203314
Candidatus_Magasanikbacteria_bacterium_CG10_big_fil_rev_8_21_14_0_10_47_10.PIR74658
Candidatus_Magasanikbacteria_bacterium_RIFCSPLOWO2_02_FULL_47_16.OGH77720
Candidatus_Magasanikbacteria_bacterium_RIFCSPLOWO2_02_FULL_44_11.OGH80453
Candidatus_Uhrbacteria_bacterium_RIFCSPHIGHO2_02_FULL_53_13.OGL70592
Anaerolineaceae_bacterium_4572_78.OQY48150
candidate_division_Kazan_bacterium_GW2011_GWA1_44_22.KKT52816
Candidatus_Woesebacteria_bacterium_RIFOXYA1_FULL_43_9.OGM78237
Nephroselmis_astigmatica.YP_009057887
Candidatus_Moranbacteria_bacterium_CG_4_8_14_3_um_filter_43_15.PIW93155
candidate_division_WWE3_bacterium_RIFCSPLOWO2_01_FULL_41_9.OGC56314
Candidatus_Moranbacteria_bacterium_GW2011_GWA2_39_41.KKR21055
Candidatus_Moranbacteria_bacterium_RIFOXYB12_FULL_35_8.OGI31153
Candidatus_Moranbacteria_bacterium_RIFOXYA1_FULL_44_7.OGI28314
Candidatus_Moranbacteria_bacterium_GW2011_GWC2_40_12.KKR44426
Candidatus_Moranbacteria_bacterium_GW2011_GWC1_45_18.KKT99154
candidate_division_WWE3_bacterium_CG06_land_8_20_14_3_00_42_16.PIU69338
Candidatus_Nealsonbacteria_bacterium_CG23_combo_of_CG06_09_8_20_14_all_40_13.PIP21885
Candidatus_Berkelbacteria_bacterium_CG1_02_42_45.OIN88907
Planctomycetales_bacterium_4484_113.OPX22322
Candidatus_Kerfeldbacteria_bacterium_CG_4_10_14_0_8_um_filter_42_10.PIY96677
Candidatus_Gottesmanbacteria_bacterium_RIFOXYB1_FULL_47_11.OGG35673
Thermoflexus_hugenholtzii_JAD2.SNB69733
Acetothermia_bacterium_64_32.KUK26106
Caldimicrobium_thiodismutans.BAU23500
Hydrogenobacter_hydrogenophilus.WP_096603052
Candidatus_Thiomargarita_nelsonii.KHD09273
Candidatus_Thiomargarita_nelsonii.KHD04553
Candidatus_Kuenenbacteria_bacterium_GW2011_GWA2_42_15.KKS39536
CBU_0182
Candidatus_Desantisbacteria_bacterium_CG1_02_38_46.OIN96097
Candidatus_Omnitrophica_bacterium_4484_171.OPX30025
candidate_division_WWE3_bacterium_CG23_combo_of_CG06_09_8_20_14_all_40_14.PIP04784
candidate_division_WWE3_bacterium_CG10_big_fil_rev_8_21_14_0_10_39_14.PJE52008
uncultured_bacterium_5G12.ALG05270
Desulfobacteraceae_bacterium_IS3.OQX26534
Anaerolineae_bacterium_CG2_30_64_16.OIO91350
Candidatus_Amesbacteria_bacterium_GW2011_GWC1_46_24.KKU31913
Candidatus_Amesbacteria_bacterium_RIFCSPLOWO2_01_FULL_49_25.OGD07803
Candidatus_Gottesmanbacteria_bacterium_GW2011_GWA2_47_9.KKU87617
Candidatus_Levybacteria_bacterium_CG2_30_37_29.OIP57459
Candidatus_Woykebacteria_bacterium_RIFCSPHIGHO2_12_FULL_43_10.OGY30275
bacterium_Candidatus_Gribaldobacteria_CG10_big_fil_rev_8_21_14_0_10_37_46.PIR91220
Candidatus_Portnoybacteria_bacterium_CG11_big_fil_rev_8_21_14_0_20_40_15.PIQ75506
Candidatus_Nealsonbacteria_bacterium_CG_4_10_14_3_um_filter_36_16.PIX88467
Candidatus_Kaiserbacteria_bacterium_CG_4_8_14_3_um_filter_38_9.PIW96716
Candidatus_Campbellbacteria_bacterium_CG22_combo_CG10_13_8_21_14_all_43_18.PIP86287
Candidatus_Kaiserbacteria_bacterium_RIFCSPHIGHO2_02_FULL_56_30.OGG67754
Candidatus_Lloydbacteria_bacterium_RIFCSPLOWO2_02_FULL_51_11.OGZ13989
Candidatus_Taylorbacteria_bacterium_RIFCSPHIGHO2_02_FULL_43_55.OHA23541
Candidatus_Nomurabacteria_bacterium_RIFCSPLOWO2_01_FULL_36_10b.OGI83453
Microgenomates_group_bacterium_GW2011_GWA2_46_16.KKU25865
Candidatus_Collierbacteria_bacterium_GW2011_GWC2_44_13.KKT39408
Candidatus_Wolfebacteria_bacterium_CG18_big_fil_WC_8_21_14_2_50_39_7.PIP91910
Candidatus_Nealsonbacteria_bacterium_CG15_BIG_FIL_POST_REV_8_21_14_020_37_12.PIW35099
Candidatus_Blackburnbacteria_bacterium_RIFCSPHIGHO2_01_FULL_43_15b.OGY09130
Candidatus_Ryanbacteria_bacterium_RIFCSPLOWO2_01_FULL_48_26.OGZ53006
Candidatus_Liptonbacteria_bacterium_RIFCSPLOWO2_01_FULL_52_25.OGZ00158
Candidatus_Vogelbacteria_bacterium_CG10_big_fil_rev_8_21_14_0_10_51_16.PIR44992
Candidatus_Kaiserbacteria_bacterium_RIFCSPHIGHO2_12_FULL_53_13.OGG70055
Gammaproteobacteria_bacterium_RIFCSPHIGHO2_12_FULL_45_9.OGT66707
Parcubacteria_group_bacterium_GW2011_GWF2_52_12.KKW25323
Candidatus_Zambryskibacteria_bacterium_RIFOXYD1_FULL_39_35.OHB16540
Candidatus_Sungbacteria_bacterium_RIFCSPLOWO2_01_FULL_59_16.OHA08244
B
Eukaryota
Longamoebia
Chlorophyta
Monomastix
Trebouxiophyceae
Bacteria
Chloroflexi
Anaerolineae
Proteobacteria
Gammaproteobacteria
candidate_division_WWE3
Candidatus_Gottesmanbacteria
Candidatus_Amesbacteria
Candidatus_Nealsonbacteria
Candidatus_Magasanikbacteria
Candidatus_Kaiserbacteria
Candidatus_Moranbacteria
Zancudomyces_culisetae
Balamuthia_mandrillaris
Acanthamoeba_castellanii
Nephroselmis_astigmatica
Monomastix_sp_OKE-1
Monomastix_sp_M722
Chlorosarcina_brevispinosa
Koliella_corcontica
Symbiochloris_reticulata
Botryococcus_braunii
Coccomyxa_viridis
uncultured_bacterium_5G12
Hydrogenobacter_hydrogenophilus
Caldimicrobium_thiodismutans
Candidatus_Omnitrophica_bacterium_4484_171
Planctomycetales_bacterium_4484_113
Thermoflexus_hugenholtzii_JAD2
Anaerolineae_bacterium_CG2_30_64_16
Anaerolineaceae_bacterium_4572_78
Desulfobacteraceae_bacterium_IS3
Gammaproteobacteria_bacterium_RIFCSPHIGHO2_12_FULL_45_9
Candidatus_Thiomargarita_nelsonii
Coxiella_burnetii_RSA_493
bacterium__Candidatus_Gribaldobacteria__CG10_big_fil_rev_8_21_14_0_10_37_46
Candidatus_Berkelbacteria_bacterium_CG1_02_42_45
candidate_division_Kazan_bacterium_GW2011_GWA1_44_22
Candidatus_Desantisbacteria_bacterium_CG1_02_38_46
Acetothermia_bacterium_64_32
candidate_division_WWE3_bacterium_RIFCSPLOWO2_01_FULL_41_9
candidate_division_WWE3_bacterium_CG06_land_8_20_14_3_00_42_16
candidate_division_WWE3_bacterium_CG09_land_8_20_14_0_10_39_24
candidate_division_WWE3_bacterium_CG10_big_fil_rev_8_21_14_0_10_39_14
Candidatus_Blackburnbacteria_bacterium_RIFCSPHIGHO2_01_FULL_43_15b
Candidatus_Woesebacteria_bacterium_RIFOXYA1_FULL_43_9
Candidatus_Levybacteria_bacterium_CG2_30_37_29
Microgenomates_group_bacterium_GW2011_GWA2_46_16
Candidatus_Collierbacteria_bacterium_GW2011_GWC2_44_13
Candidatus_Woykebacteria_bacterium_RIFCSPHIGHO2_12_FULL_43_10
Candidatus_Gottesmanbacteria_bacterium_RIFOXYB1_FULL_47_11
Candidatus_Gottesmanbacteria_bacterium_GW2011_GWA2_47_9
Candidatus_Amesbacteria_bacterium_RIFCSPLOWO2_01_FULL_49_25
Candidatus_Amesbacteria_bacterium_GW2011_GWB1_47_19
Candidatus_Campbellbacteria_bacterium_CG11_big_fil_rev_8_21_14_0_20_44_21
Candidatus_Vogelbacteria_bacterium_CG10_big_fil_rev_8_21_14_0_10_51_16
Candidatus_Ryanbacteria_bacterium_RIFCSPLOWO2_01_FULL_48_26
Candidatus_Portnoybacteria_bacterium_CG_4_10_14_0_8_um_filter_40_50
Candidatus_Uhrbacteria_bacterium_RIFCSPHIGHO2_02_FULL_53_13
Candidatus_Zambryskibacteria_bacterium_RIFOXYD1_FULL_39_35
Candidatus_Wolfebacteria_bacterium_CG18_big_fil_WC_8_21_14_2_50_39_7
Candidatus_Andersenbacteria_bacterium_RIFCSPHIGHO2_12_FULL_45_8
Candidatus_Kerfeldbacteria_bacterium_CG_4_10_14_0_8_um_filter_42_10
Candidatus_Sungbacteria_bacterium_RIFCSPLOWO2_01_FULL_59_16
Candidatus_Taylorbacteria_bacterium_RIFCSPHIGHO2_02_FULL_43_55
Parcubacteria_group_bacterium_GW2011_GWB1_53_43
Candidatus_Lloydbacteria_bacterium_RIFCSPLOWO2_02_FULL_51_11
Candidatus_Nomurabacteria_bacterium_RIFCSPLOWO2_01_FULL_36_10b
Candidatus_Kuenenbacteria_bacterium_GW2011_GWA2_42_15
Candidatus_Liptonbacteria_bacterium_RIFCSPLOWO2_01_FULL_52_25
Candidatus_Nealsonbacteria_bacterium_CG07_land_8_20_14_0_80_40_10
Candidatus_Nealsonbacteria_bacterium_CG_4_10_14_3_um_filter_36_16
Candidatus_Nealsonbacteria_bacterium_CG15_BIG_FIL_POST_REV_8_21_14_020_37_12
Candidatus_Magasanikbacteria_bacterium_RIFCSPLOWO2_02_FULL_44_11
Candidatus_Magasanikbacteria_bacterium_RIFOXYC2_FULL_40_16
Candidatus_Magasanikbacteria_bacterium_CG10_big_fil_rev_8_21_14_0_10_47_10
Candidatus_Kaiserbacteria_bacterium_CG_4_8_14_3_um_filter_38_9
Candidatus_Kaiserbacteria_bacterium_RIFCSPHIGHO2_02_FULL_56_30
Candidatus_Kaiserbacteria_bacterium_RIFCSPHIGHO2_12_FULL_53_13
Candidatus_Moranbacteria_bacterium_CG_4_8_14_3_um_filter_43_15
Candidatus_Moranbacteria_bacterium_GW2011_GWA2_39_41
Candidatus_Moranbacteria_bacterium_GW2011_GWC1_45_18
Candidatus_Moranbacteria_bacterium_GW2011_GWC2_40_12
Candidatus_Moranbacteria_bacterium_RIFOXYA1_FULL_44_7
Candidatus_Moranbacteria_bacterium_RIFOXYB12_FULL_35_8

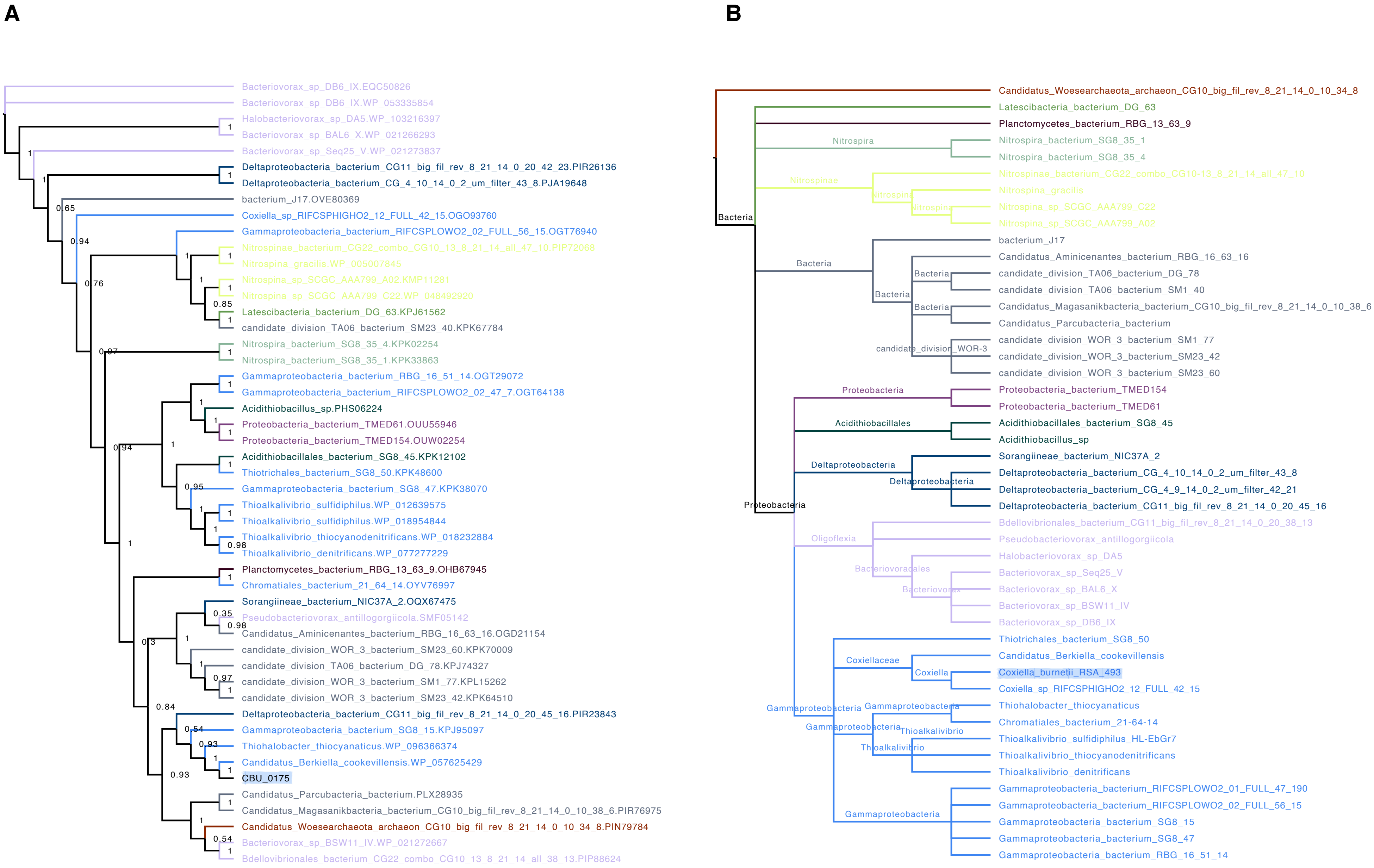
A
Bacteriovorax_sp_DB6_IX.EQC50826
Bacteriovorax_sp_DB6_IX.WP_053335854
Halobacteriovorax_sp_DA5.WP_103216397
Bacteriovorax_sp_BAL6_X.WP_021266293
Bacteriovorax_sp_Seq25_V.WP_021273837
Deltaproteobacteria_bacterium_CG11_big_fil_rev_8_21_14_0_20_42_23.PIR26136
Deltaproteobacteria_bacterium_CG_4_10_14_0_2_um_filter_43_8.PJA19648
bacterium_J17.OVE80369
Coxiella_sp_RIFCSPHIGHO2_12_FULL_42_15.OGO93760
Gammaproteobacteria_bacterium_RIFCSPLOWO2_02_FULL_56_15.OGT76940
Nitrospinae_bacterium_CG22_combo_CG10_13_8_21_14_all_47_10.PIP72088
Nitrospina_gracilis.WP_005007845
Nitrospina_sp_SCGC_AAA799_A02.KMP11281
Nitrospina_sp_SCGC_AAA799_C22.WP_048492920
Latescibacteria_bacterium_DG_63.KPJ61562
candidate_division_TA06_bacterium_SM23_40.KPK67784
Nitrospira_bacterium_SG8_35_4.KPK02254
Nitrospira_bacterium_SG8_35_1.KPK33863
Gammaproteobacteria_bacterium_RBG_16_51_14.OGT29072
Gammaproteobacteria_bacterium_RIFCSPLOWO2_02_47_7.OGT64138
Acidithiobacillus_sp.PHS06224
Proteobacteria_bacterium_TMED61.OUU55946
Proteobacteria_bacterium_TMED154.OUW02254
Acidithiobacillales_bacterium_SG8_45.KPK12102
Thiotrichales_bacterium_SG8_50.KPK48600
Gammaproteobacteria_bacterium_SG8_47.KPK38070
Thioalkalivibrio_sulfidiphilus.WP_012639575
Thioalkalivibrio_sulfidiphilus.WP_018954844
Thioalkalivibrio_thiocyanodenitrificans.WP_018232884
Thioalkalivibrio_denitrificans.WP_077277229
Planctomycetes_bacterium_RBG_13_63_9.OHB67945
Chromatiales_bacterium_21_64_14.OYV76997
Sorangiineae_bacterium_NIC37A_2.OQX67475
Pseudobacteriovorax_antillogorgiicola.SMF05142
Candidatus_Aminicenantes_bacterium_RBG_16_63_16.OGD21154
candidate_division_WOR_3_bacterium_SM23_60.KPK70009
candidate_division_TA06_bacterium_DG_78.KPJ74327
candidate_division_WOR_3_bacterium_SM1_77.KPL15262
candidate_division_WOR_3_bacterium_SM23_42.KPK64510
Deltaproteobacteria_bacterium_CG11_big_fil_rev_8_21_14_0_20_45_16.PIR23843
Gammaproteobacteria_bacterium_SG8_15.KPJ95097
Thiohalobacter_thiocyanaticus.WP_096366374
Candidatus_Berkiella_cookevillensis.WP_057625429
CBU_0175
Candidatus_Parcubacteria_bacterium.PLX28935
Candidatus_Magasanikbacteria_bacterium_CG10_big_fil_rev_8_21_14_0_10_38_6.PIR76975
Candidatus_Woesearchaeota_archaeon_CG10_big_fil_rev_8_21_14_0_10_34_8.PIN79784
Bacteriovorax_sp_BSW11_IV.WP_021272667
Bdellovibrionales_bacterium_CG22_combo_CG10_13_8_21_14_all_38_13.PIP88624
B
Candidatus_Woesearchaeota_archaeon_CG10_big_fil_rev_8_21_14_0_10_34_8
Latescibacteria_bacterium_DG_63
Planctomycetes_bacterium_RBG_13_63_9
Nitrospira_bacterium_SG8_35_1
Nitrospira_bacterium_SG8_35_4
Nitrospinae_bacterium_CG22_combo_CG10-13_8_21_14_all_47_10
Nitrospina_gracilis
Nitrospina_sp_SCGC_AAA799_C22
Nitrospina_sp_SCGC_AAA799_A02
bacterium_J17
Candidatus_Aminicenantes_bacterium_RBG_16_63_16
candidate_division_TA06_bacterium_DG_78
candidate_division_TA06_bacterium_SM1_40
Candidatus_Magasanikbacteria_bacterium_CG10_big_fil_rev_8_21_14_0_10_38_6
Candidatus_Parcubacteria_bacterium
candidate_division_WOR_3_bacterium_SM1_77
candidate_division_WOR_3_bacterium_SM23_42
candidate_division_WOR_3_bacterium_SM23_60
Proteobacteria_bacterium_TMED154
Proteobacteria_bacterium_TMED61
Acidithiobacillales_bacterium_SG8_45
Acidithiobacillus_sp
Sorangiineae_bacterium_NIC37A_2
Deltaproteobacteria_bacterium_CG_4_10_14_0_2_um_filter_43_8
Deltaproteobacteria_bacterium_CG_4_9_14_0_2_um_filter_42_21
Deltaproteobacteria_bacterium_CG11_big_fil_rev_8_21_14_0_20_45_16
Bdellovibrionales_bacterium_CG11_big_fil_rev_8_21_14_0_20_38_13
Pseudobacteriovorax_antillogorgiicola
Halobacteriovorax_sp_DA5
Bacteriovorax_sp_Seq25_V
Bacteriovorax_sp_BAL6_X
Bacteriovorax_sp_BSW11_IV
Bacteriovorax_sp_DB6_IX
Thiotrichales_bacterium_SG8_50
Candidatus_Berkiella_cookevillensis
Coxiella_burnetii_RSA_493
Coxiella_sp_RIFCSPHIGHO2_12_FULL_42_15
Thiohalobacter_thiocyanaticus
Chromatiales_bacterium_21-64-14
Thioalkalivibrio_sulfidiphilus_HL-EbGr7
Thioalkalivibrio_thiocyanodenitrificans
Thioalkalivibrio_denitrificans
Gammaproteobacteria_bacterium_RIFCSPLOWO2_01_FULL_47_190
Gammaproteobacteria_bacterium_RIFCSPLOWO2_02_FULL_56_15
Gammaproteobacteria_bacterium_SG8_15
Gammaproteobacteria_bacterium_SG8_47
Gammaproteobacteria_bacterium_RBG_16_51_14

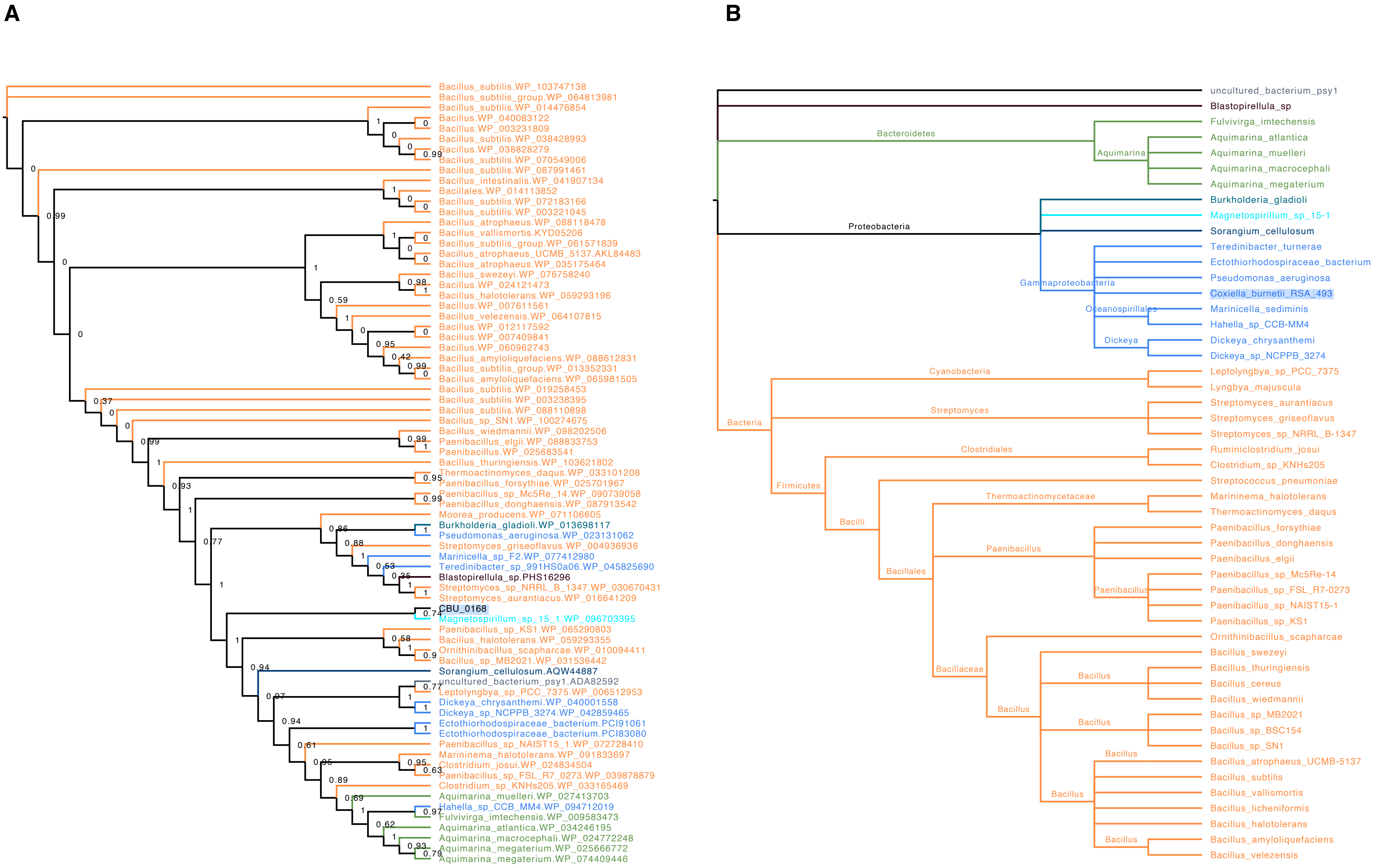
A
Bacillus_subtilis.WP_103747138
Bacillus_subtilis_group.WP_064813981
Bacillus_subtilis.WP_014476854
Bacillus.WP_040083122
Bacillus.WP_003231809
Bacillus_subtilis.WP_038428993
Bacillus.WP_038828279
Bacillus_subtilis.WP_070549006
Bacillus_subtilis.WP_087991461
Bacillus_intestinalis.WP_041907134
Bacillales.WP_014113852
Bacillus_subtilis.WP_072183166
Bacillus_subtilis.WP_003221045
Bacillus_atrophaeus.WP_088118478
Bacillus_vallismortis.KYD05206
Bacillus_subtilis_group.WP_061571839
Bacillus_atrophaeus_UCMB_5137.AKL84483
Bacillus_atrophaeus.WP_035175464
Bacillus_swezeyi.WP_076758240
Bacillus.WP_024121473
Bacillus_halotolerans.WP_059293196
Bacillus.WP_007611561
Bacillus_velezensis.WP_064107815
Bacillus.WP_012117592
Bacillus.WP_007409841
Bacillus.WP_060962743
Bacillus_amyloliquefaciens.WP_088612831
Bacillus_subtilis_group.WP_013352331
Bacillus_amyloliquefaciens.WP_065981505
Bacillus_subtilis.WP_019258453
Bacillus_subtilis.WP_003238395
Bacillus_subtilis.WP_088110898
Bacillus_sp_SN1.WP_100274675
Bacillus_wiedmannii.WP_098202506
Paenibacillus_elgii.WP_088833753
Paenibacillus.WP_025683541
Bacillus_thuringiensis.WP_103621802
Thermoactinomyces_daqus.WP_033101208
Paenibacillus_forsythiae.WP_025701967
Paenibacillus_sp_Mc5Re_14.WP_090739058
Paenibacillus_donghaensis.WP_087913542
Moorea_producens.WP_071106605
Burkholderia_gladioli.WP_013698117
Pseudomonas_aeruginosa.WP_023131062
Streptomyces_griseoflavus.WP_004936936
Marinicella_sp_F2.WP_077412980
Teredinibacter_sp_991HS0a06.WP_045825690
Blastopirellula_sp.PHS16296
Streptomyces_sp_NRRL_B_1347.WP_030670431
Streptomyces_aurantiacus.WP_016641209
CBU_0168
Magnetospirillum_sp_15_1.WP_096703395
Paenibacillus_sp_KS1.WP_065290803
Bacillus_halotolerans.WP_059293355
Ornithinibacillus_scapharcae.WP_010094411
Bacillus_sp_MB2021.WP_031536442
Sorangium_cellulosum.AQW44887
uncultured_bacterium_psy1.ADA82592
Leptolyngbya_sp_PCC_7375.WP_006512953
Dickeya_chrysanthemi.WP_040001558
Dickeya_sp_NCPPB_3274.WP_042859465
Ectothiorhodospiraceae_bacterium.PCI91061
Ectothiorhodospiraceae_bacterium.PCI83080
Paenibacillus_sp_NAIST15_1.WP_072728410
Marininema_halotolerans.WP_091833697
Clostridium_josui.WP_024834504
Paenibacillus_sp_FSL_R7_0273.WP_039878879
Clostridium_sp_KNHs205.WP_033165469
Aquimarina_muelleri.WP_027413703
Hahella_sp_CCB_MM4.WP_094712019
Fulvivirga_imtechensis.WP_009583473
Aquimarina_atlantica.WP_034246195
Aquimarina_macrocephali.WP_024772248
Aquimarina_megaterium.WP_025666772
Aquimarina_megaterium.WP_074409446
B
uncultured_bacterium_psy1
Blastopirellula_sp
Bacteroidetes
Aquimarina
Fulvivirga_imtechensis
Aquimarina_atlantica
Aquimarina_muelleri
Aquimarina_macrocephali
Aquimarina_megaterium
Proteobacteria
Burkholderia_gladioli
Magnetospirillum_sp_15-1
Sorangium_cellulosum
Gammaproteobacteria
Teredinibacter_turnerae
Ectothiorhodospiraceae_bacterium
Pseudomonas_aeruginosa
Coxiella_burnetii_RSA_493
Oceanospirillales
Marinicella_sedimenis
Hahella_sp_CCB-MM4
Dickeya
Dickeya_chrysanthemi
Dickeya_sp_NCPPB_3274
Bacteria
Cyanobacteria
Leptolyngbya_sp_PCC_7375
Lyngbya_majuscula
Streptomyces
Streptomyces_aurantiacus
Streptomyces_griseoflavus
Streptomyces_sp_NRRL_B-1347
Firmicutes
Clostridiales
Ruminiclostridium_josui
Clostridium_sp_KNHs205
Bacilli
Streptococcus_pneumoniae
Bacillales
Thermoactinomycetaceae
Marininema_halotolerans
Thermoactinomyces_daqus
Paenibacillus
Paenibacillus_forsythiae
Paenibacillus_donghaensis
Paenibacillus_elgii
Paenibacillus_sp_Mc5Re-14
Paenibacillus_sp_FSL_R7-0273
Paenibacillus_sp_NAIST15-1
Paenibacillus_sp_KS1
Bacillaceae
Ornithinibacillus_scapharcae
Bacillus
Bacillus_swezeyi
Bacillus_thuringiensis
Bacillus_cereus
Bacillus_wiedmannii
Bacillus_sp_MB2021
Bacillus_sp_BSC154
Bacillus_sp_SN1
Bacillus_atrophaeus_UCMB-5137
Bacillus_subtilis
Bacillus_vallismortis
Bacillus_licheniformis
Bacillus_halotolerans
Bacillus_amyloliquefaciens
Bacillus_velezensis

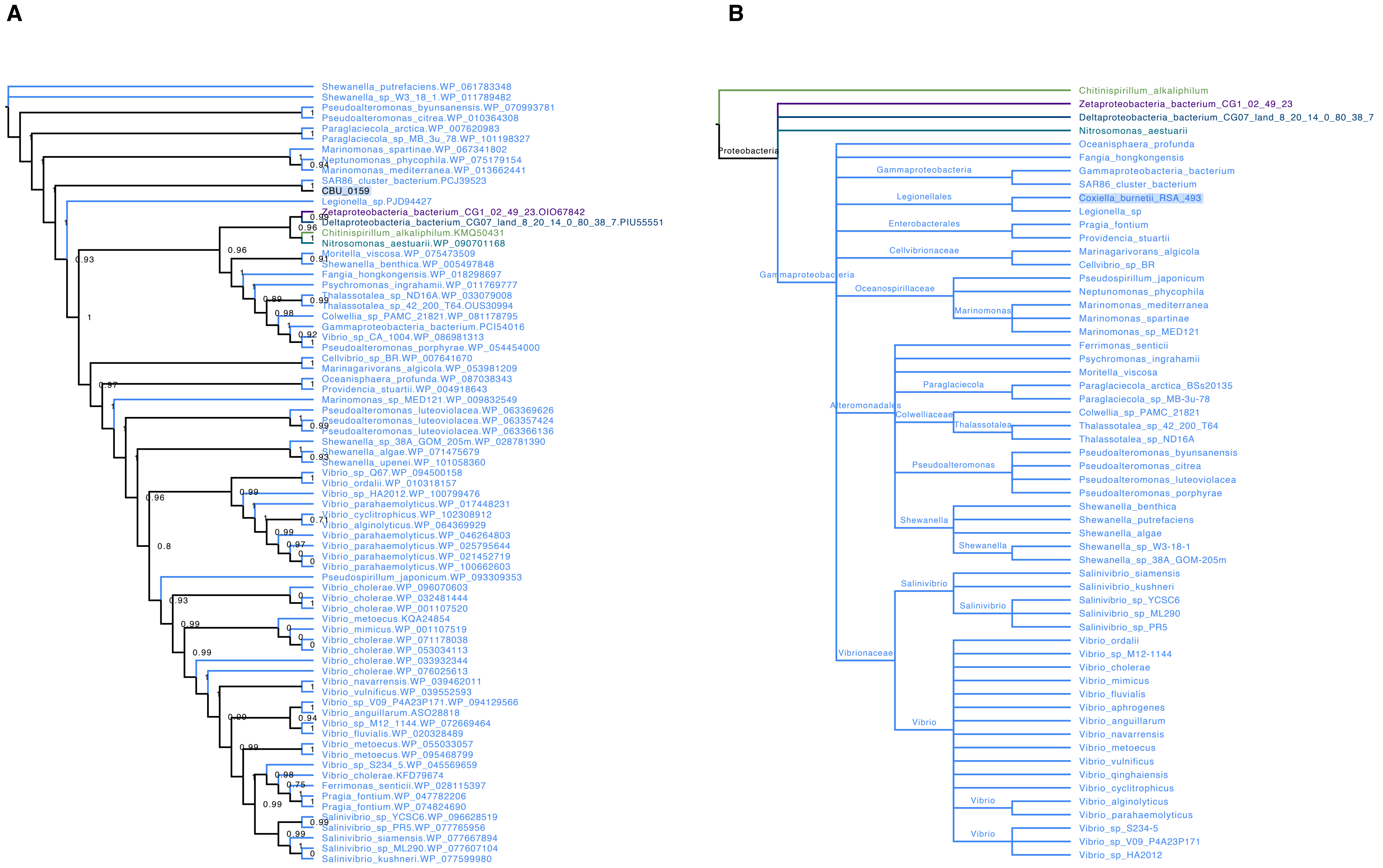
A
Shewanella_putrefaciens.WP_061783348
Shewanella_sp_W3_18_1.WP_011789482
Pseudoalteromonas_byunsanensis.WP_070993781
Pseudoalteromonas_citrea.WP_010364308
Paraglaciecola_arctica.WP_007620983
Paraglaciecola_sp_MB_3u_78.WP_101198327
Marinomonas_spartinae.WP_067341802
Neptunomonas_phycophila.WP_075179154
Marinomonas_mediterranea.WP_013662441
SAR86_cluster_bacterium.PCJ39523
CBU_0159
Legionella_sp.PJD94427
Zetaproteobacteria_bacterium_CG1_02_49_23.OIO67842
Deltaproteobacteria_bacterium_CG07_land_8_20_14_0_80_38_7.PIU55551
Chitinispirillum_alkaliphilum.KMQ50431
Nitrosomonas_aestuarii.WP_090701168
Moritella_viscosa.WP_075473509
Shewanella_benthica.WP_005497848
Fangia_hongkongensis.WP_018298697
Psychromonas_ingrahamii.WP_011769777
Thalassotalea_sp_ND16A.WP_033079008
Thalassotalea_sp_42_200_T64.OUS30994
Colwellia_sp_PAMC_21821.WP_081178795
Gammaproteobacteria_bacterium.PCI54016
Vibrio_sp_CA_1004.WP_086981313
Pseudoalteromonas_porphyrae.WP_054454000
Cellvibrio_sp_BR.WP_007641670
Marinagarivorans_algicola.WP_053981209
Oceanisphaera_profunda.WP_087038343
Providencia_stuartii.WP_004918643
Marinomonas_sp_MED121.WP_009832549
Pseudoalteromonas_luteoviolacea.WP_063369626
Pseudoalteromonas_luteoviolacea.WP_063357424
Pseudoalteromonas_luteoviolacea.WP_063366136
Shewanella_sp_38A_GOM_205m.WP_028781390
Shewanella_algae.WP_071475679
Shewanella_upenei.WP_101058360
Vibrio_sp_Q67.WP_094500158
Vibrio_ordalii.WP_010318157
Vibrio_sp_HA2012.WP_100799476
Vibrio_parahaemolyticus.WP_017448231
Vibrio_cyclitrophicus.WP_102308912
Vibrio_alginolyticus.WP_064369929
Vibrio_parahaemolyticus.WP_046264803
Vibrio_parahaemolyticus.WP_025795644
Vibrio_parahaemolyticus.WP_021452719
Vibrio_parahaemolyticus.WP_100662603
Pseudospirillum_japonicum.WP_093309353
Vibrio_cholerae.WP_096070603
Vibrio_cholerae.WP_032481444
Vibrio_cholerae.WP_001107520
Vibrio_metoecus.KQA24854
Vibrio_mimicus.WP_001107519
Vibrio_cholerae.WP_071178038
Vibrio_cholerae.WP_053034113
Vibrio_cholerae.WP_033932344
Vibrio_cholerae.WP_076025613
Vibrio_navarrensis.WP_039462011
Vibrio_vulnificus.WP_039552593
Vibrio_sp_V09_P4A23P171.WP_094129566
Vibrio_anguillarum.ASO28818
Vibrio_sp_M12_1144.WP_072669464
Vibrio_fluvialis.WP_020328489
Vibrio_metoecus.WP_055033057
Vibrio_metoecus.WP_095468799
Vibrio_sp_S234_5.WP_045569659
Vibrio_cholerae.KFD79674
Ferrimonas_senticii.WP_028115397
Pragia_fontium.WP_047782206
Pragia_fontium.WP_074824690
Salinivibrio_sp_YCSC6.WP_096628519
Salinivibrio_sp_PR5.WP_077765956
Salinivibrio_siamensis.WP_077667894
Salinivibrio_sp_ML290.WP_077607104
Salinivibrio_kushneri.WP_077599980
B
Proteobacteria
Gammaproteobacteria
Legionellales
Enterobacterales
Cellvibrionaceae
Oceanospirillaceae
Marinomonas
Alteromonadales
Paraglaciecola
Colwelliaceae
Thalassotalea
Pseudoalteromonas
Shewanella
Salinivibrio
Vibrionaceae
Vibrio
Chitinispirillum_alkaliphilum
Zetaproteobacteria_bacterium_CG1_02_49_23
Deltaproteobacteria_bacterium_CG07_land_8_20_14_0_80_38_7
Nitrosomonas_aestuarii
Oceanisphaera_profunda
Fangia_hongkongensis
Gammaproteobacteria_bacterium
SAR86_cluster_bacterium
Coxiella_burnetii_RSA_493
Legionella_sp
Pragia_fontium
Providencia_stuartii
Marinagarivorans_algicola
Cellvibrio_sp_BR
Pseudospirillum_japonicum
Neptunomonas_phycophila
Marinomonas_mediterranea
Marinomonas_spartinae
Marinomonas_sp_MED121
Ferrimonas_senticii
Psychromonas_ingrahamii
Moritella_viscosa
Paraglaciecola_arctica_BSs20135
Paraglaciecola_sp_MB-3u-78
Colwellia_sp_PAMC_21821
Thalassotalea_sp_42_200_T64
Thalassotalea_sp_ND16A
Pseudoalteromonas_byunsanensis
Pseudoalteromonas_citrea
Pseudoalteromonas_luteoviolacea
Pseudoalteromonas_porphyrae
Shewanella_benthica
Shewanella_putrefaciens
Shewanella_algae
Shewanella_sp_W3-18-1
Shewanella_sp_38A_GOM-205m
Salinivibrio_siamensis
Salinivibrio_kushneri
Salinivibrio_sp_YCSC6
Salinivibrio_sp_ML290
Salinivibrio_sp_PR5
Vibrio_ordalii
Vibrio_sp_M12-1144
Vibrio_cholerae
Vibrio_mimicus
Vibrio_fluvialis
Vibrio_aphrogenes
Vibrio_anguillarum
Vibrio_navarrensis
Vibrio_metoecus
Vibrio_vulnificus
Vibrio_qinghaiensis
Vibrio_cyclitrophicus
Vibrio_alginolyticus
Vibrio_parahaemolyticus
Vibrio_sp_S234-5
Vibrio_sp_V09_P4A23P171
Vibrio_sp_HA2012

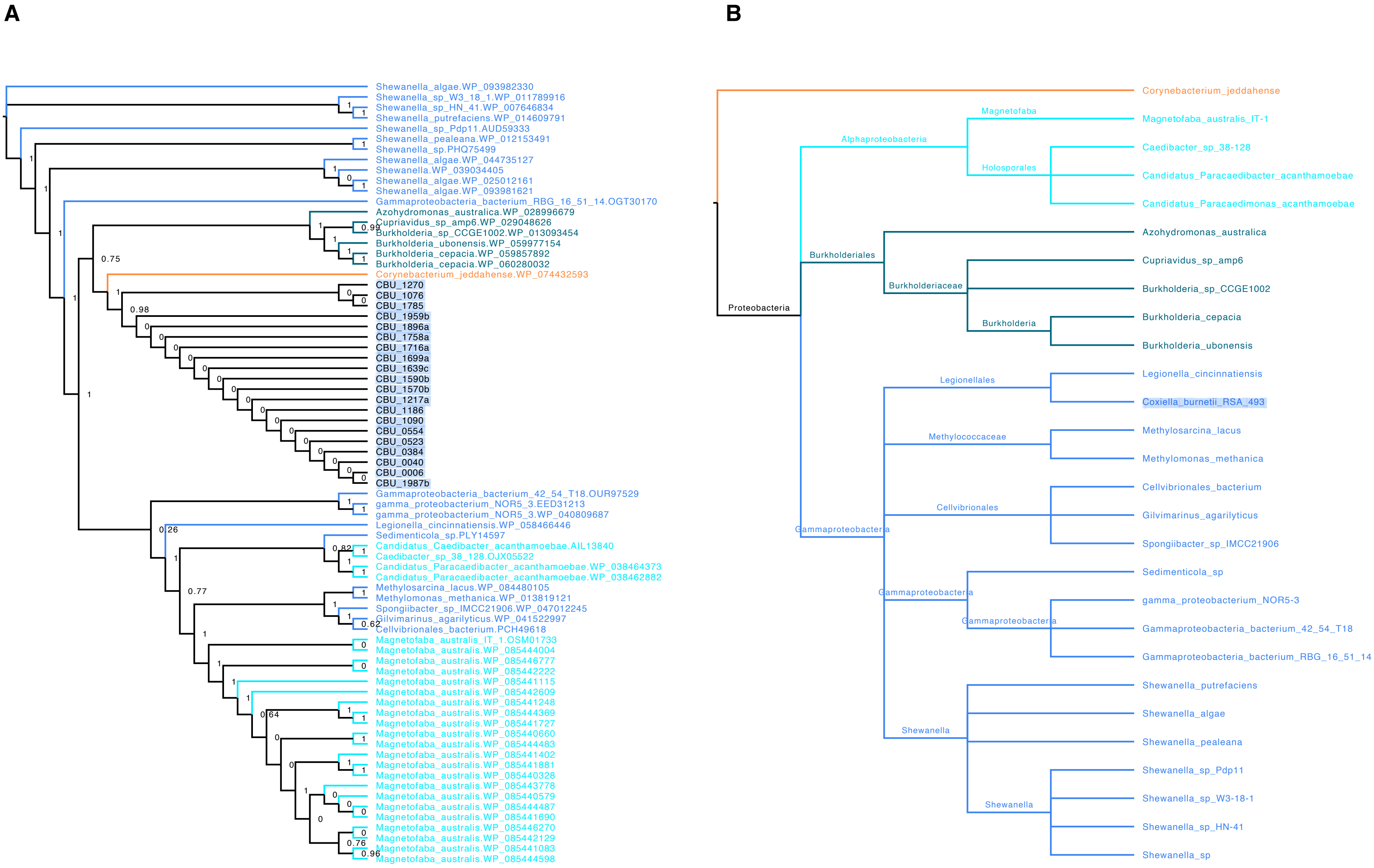
A
Shewanella_algae.WP_093982330
Shewanella_sp_W3_18_1.WP_011789916
Shewanella_sp_HN_41.WP_007646834
Shewanella_putrefaciens.WP_014609791
Shewanella_sp_Pdp11.AUD59333
Shewanella_pealeana.WP_012153491
Shewanella_sp.PHQ75499
Shewanella_algae.WP_044735127
Shewanella.WP_039034405
Shewanella_algae.WP_025012161
Shewanella_algae.WP_093981621
Gammaproteobacteria_bacterium_RBG_16_51_14.OGT30170
Azohydromonas_australica.WP_028996679
Cupriavidus_sp_amp6.WP_029048626
Burkholderia_sp_CCGE1002.WP_013093454
Burkholderia_ubonensis.WP_059977154
Burkholderia_cepacia.WP_059857892
Burkholderia_cepacia.WP_060280032
Corynebacterium_jeddahense.WP_074432593
CBU_1270
CBU_1076
CBU_1785
CBU_1959b
CBU_1896a
CBU_1758a
CBU_1716a
CBU_1699a
CBU_1639c
CBU_1590b
CBU_1570b
CBU_1217a
CBU_1186
CBU_1090
CBU_0554
CBU_0523
CBU_0384
CBU_0040
CBU_0006
CBU_1987b
Gammaproteobacteria_bacterium_42_54_T18.OUR97529
gamma_proteobacterium_NOR5_3.EED31213
gamma_proteobacterium_NOR5_3.WP_040809687
Legionella_cincinnatiensis.WP_058466446
Sedimenticola_sp.PLY14597
Candidatus_Caedibacter_acanthamoebae.AIL13840
Caedibacter_sp_38_128.OJX05522
Candidatus_Paracaedibacter_acanthamoebae.WP_038464373
Candidatus_Paracaedibacter_acanthamoebae.WP_038462882
Methylosarcina_lacus.WP_084480105
Methylomonas_methanica.WP_013819121
Spongiibacter_sp_IMCC21906.WP_047012245
Gilvimarinus_agarilyticus.WP_041522997
Cellvibrionales_bacterium.PCH49618
Magnetofaba_australis_IT_1.OSM01733
Magnetofaba_australis.WP_085444004
Magnetofaba_australis.WP_085446777
Magnetofaba_australis.WP_085442222
Magnetofaba_australis.WP_085441115
Magnetofaba_australis.WP_085442609
Magnetofaba_australis.WP_085441248
Magnetofaba_australis.WP_085444369
Magnetofaba_australis.WP_085441727
Magnetofaba_australis.WP_085440660
Magnetofaba_australis.WP_085444483
Magnetofaba_australis.WP_085441402
Magnetofaba_australis.WP_085441881
Magnetofaba_australis.WP_085440328
Magnetofaba_australis.WP_085443778
Magnetofaba_australis.WP_085440579
Magnetofaba_australis.WP_085444487
Magnetofaba_australis.WP_085441690
Magnetofaba_australis.WP_085446270
Magnetofaba_australis.WP_085442129
Magnetofaba_australis.WP_085441083
Magnetofaba_australis.WP_085444598
B
Corynebacterium_jeddahense
Magnetofaba
Magnetofaba_australis_IT-1
Alphaproteobacteria
Caedibacter_sp_38-128
Holosporales
Candidatus_Paracaedibacter_acanthamoebae
Candidatus_Paracaedimonas_acanthamoebae
Azohydromonas_australica
Burkholderiales
Cupriavidus_sp_amp6
Burkholderiaceae
Burkholderia_sp_CCGE1002
Proteobacteria
Burkholderia
Burkholderia_cepacia
Burkholderia_ubonensis
Legionella_cincinnatiensis
Legionellales
Coxiella_burnetii_RSA_493
Methylosarcina_lacus
Methylococcaceae
Methylomonas_methanica
Cellvibrionales_bacterium
Cellvibrionales
Gilvimarinus_agarilyticus
Gammaproteobacteria
Spongiibacter_sp_IMCC21906
Sedimenticola_sp
Gammaproteobacteria
gamma_proteobacterium_NOR5-3
Gammaproteobacteria
Gammaproteobacteria_bacterium_42_54_T18
Gammaproteobacteria_bacterium_RBG_16_51_14
Shewanella_putrefaciens
Shewanella_algae
Shewanella
Shewanella_pealeana
Shewanella_sp_Pdp11
Shewanella_sp_W3-18-1
Shewanella
Shewanella_sp_HN-41
Shewanella_sp
