## Supplementary figures and images for "Ticks convert pathogenic *Coxiella* into endosymbionts"

### Supplemental Figure 2

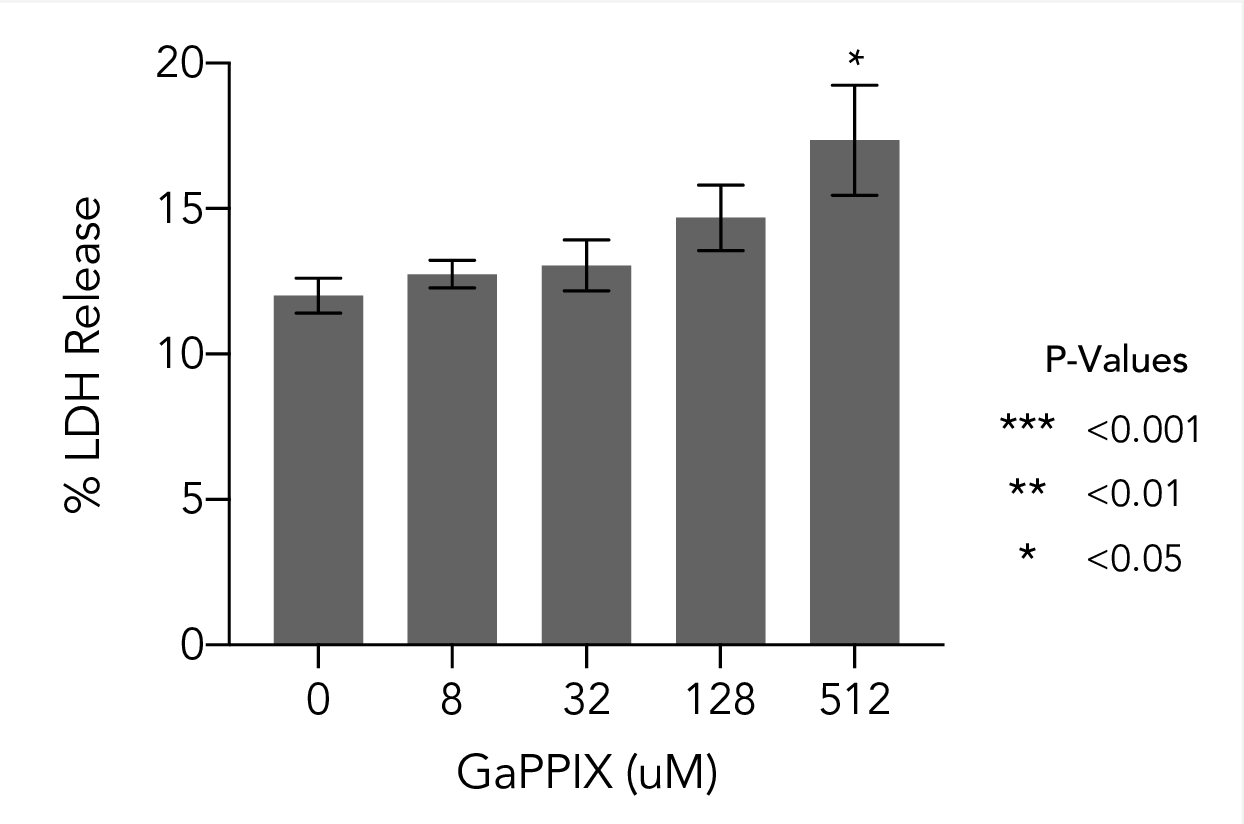

### Supplemental Figure 3

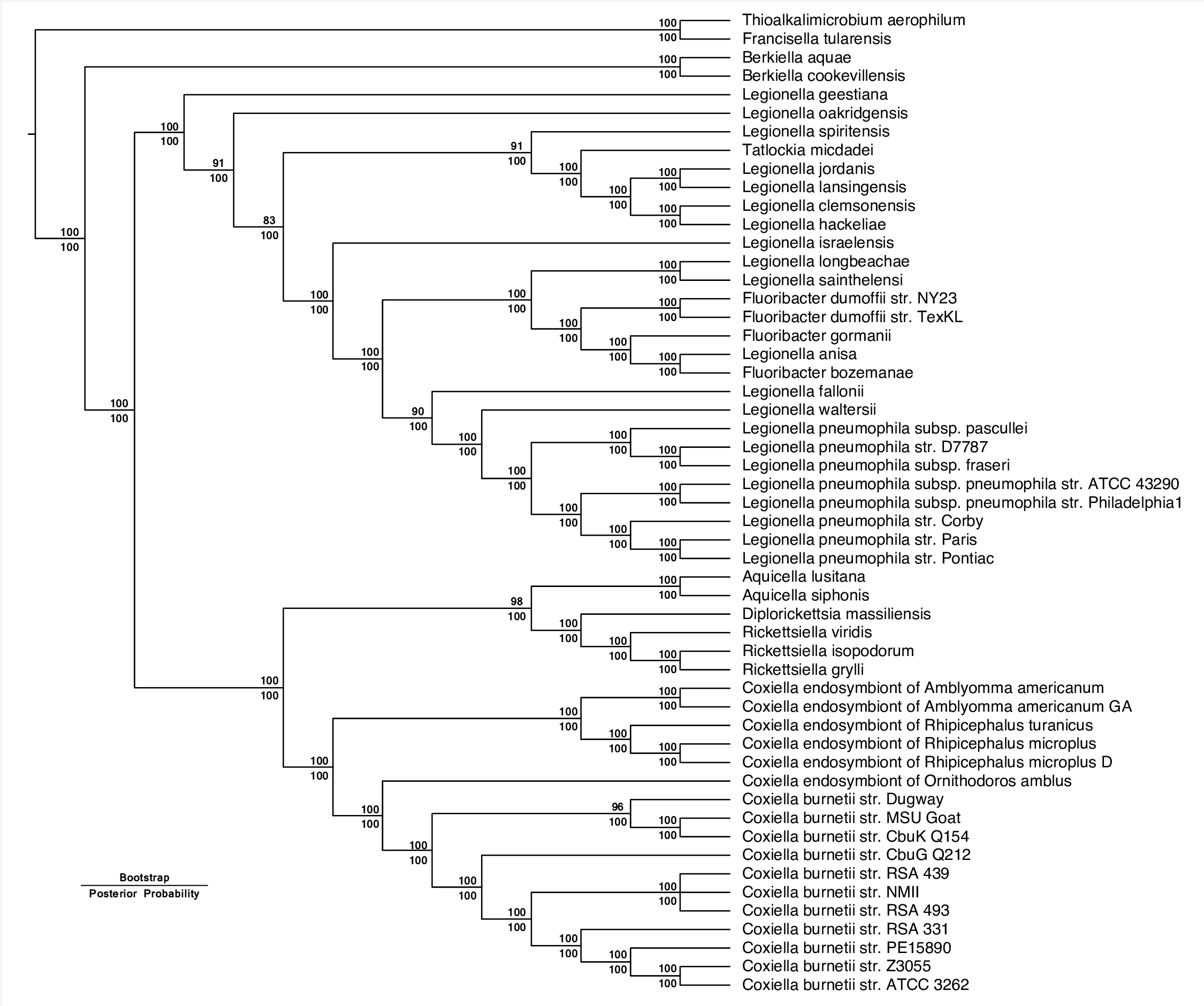
